# Genome wide characterization and expression analysis of CrRLK1L gene family in wheat unravels their roles in development and stress-specific responses

**DOI:** 10.1101/2023.05.24.541849

**Authors:** Nilesh D. Gawande, Subramanian Sankaranarayanan

## Abstract

*Catharanthus roseus* receptor-like kinase 1-like (CrRLK1L) genes encode a subfamily of receptor-like kinases (RLK) that regulate diverse processes during plant growth, development and stress responses. This study aims to provide a comprehensive genome-wide functional characterization of CrRLK1L family in bread wheat (*Triticum aestivum*). The genome of *T. aestivum* encodes 15 *CrRLK1L* family genes that has 43 paralogous copies with three homeologs each, except for *-2-D* and *-7-A*, which were found to be absent. In addition, a frame shift deletion was identified in the Paralog *-2-B*. Chromosomal localization analysis revealed a markedly uneven distribution of *Ta-CrRLK1L* genes across seven different chromosomes, with chromosome 4 housing the highest number of genes while chromosome 6 lacked any CrRLK1L genes. Tissue-specific gene expression analysis revealed distinct expression patterns among the members of the gene family, with certain members exhibiting heightened expression in reproductive tissues. Gene expression analysis under various abiotic and biotic stress conditions unveiled differential regulation of different gene family members. An examination of cis-acting elements in the promoter regions, identified specific elements crucial for plant growth and developmental processes. This comprehensive genome wide analysis and expression study provide valuable insights into the essential functions of CrRLK1L members in wheat.

## Introduction

The plant receptor-like kinases (RLKs) are transmembrane receptor proteins localized in the cell membrane, constituting a well-studied family of kinases. RLKs play a crucial role in regulating various intercellular activities, as well as plant growth and developmental processes, by sensing numerous extracellular cues^1^. RLKs, comprising of an intracellular serine/threonine kinase domain, a transmembrane domain, and a varied extracellular domain, facilitate cellular signaling through interaction with different partners via their transmembrane and juxtamembrane domains. The RLK families are sub-classified based on their N-terminal extracellular domain, which determines their ligand specificity ^2^.

The *Catharanthus roseus* receptor-like kinase 1-like (CrRLK1L) gene family belongs to a specific subfamily of RLKs and is characterized by three distinct domains: an extracellular ligand-binding domain, transmembrane domain, and kinase domain. The extracellular domain of CrRLK1Ls has homology with the carbohydrate binding domain, but it lacks residues important for carbohydrate-rich ligand binding^3^. CrRLK1L genes were first discovered as a novel RLK in Madagascar periwinkle^4^. CrRLK1L gene families have been widely characterized in various plant species, resulting in the identification of diverse members within the gene family.

CrRLK1L genes play multiple crucial roles in plant growth, development, and stress responses^56^. In Arabidopsis, six out of the 17 subfamily members have been identified as regulators of cell growth, cell-cell communication, and cell wall remodelling during vegetative and reproductive development^7^. The Arabidopsis CrRLK1Ls have been clustered into 10 clades, and their functional characterization has shed light into their involvement in plant reproduction. For instance, Arabidopsis CrRLK1Ls, including At-BUPS1 (AT4G39110.1), At-BUPS2 (AT2G21480.1), At-ANX1 (AT3G04690.1), and At-ANX2 (AT5G28680.2), form a complex and that functions in maintaining the pollen tube integrity and preventing premature rupture of the pollen tube before reaching the female gametophyte^8^. Other CrRLK1L members in Arabidopsis include At-FER (AT3G51550.1), At-HERK1 (AT3G46290.1), and At-ANJEA (AT5G59700.1), which are involved in pollen tube reception by the synergid cells and the prevention of polytubey^9^. The most versatile member, FERONIA (FER), named after the Etruscan goddess fertility, acts as a receptor for Rapid Alkalization Factors (RALFs), and regulates numerous plant developmental processes as well as stress responses ^10 11,12,13^. FERONIA and ANJEA are also involved in establishing various pollination barriers through regulation of reactive oxygen species^14,15^. FERONIA is additionally involved in mediating pathogen responses, as loss-of-function FER plants exhibit increased resistance to certain bacterial and fungal pathogens ^16,17,18^. Besides these members, other CrRLK1Ls, like At-MDS1 (AT5G38990.1), At-MDS2 (AT5G39000.1), At-MDS3 (AT5G39020.1), and At-MDS4 (AT5G39030.1), are known to be involved in plant immunity and metal ion stress responses^19,20^.

The common bread wheat, *T. aestivum*, is among the most important cereal crops in the world. Wheat was the first crop to be domesticated and is the major staple food crop grown globally^21^. Belonging to the Triticeae family, wheat encompasses nearly 300 species, including its closest relatives, *Hordeum vulgare* (barley) and *Secale cerelae* (rye). Allopolyploidation through hybridization with species from the *Aegilops* genus was a major breakthrough in the evolution of Triticum species. The divergence between the diploid AA genome species of Triticum, *T. urartu*, and *T. monococcum* occurred less than a million years ago. The first polyploidization event occurred 0.5 million years ago between *T. urartu* (AA genome) and *Aegilops speltoides* (SS genome), which led to the origin of two species, namely, *T. turgidum* (AABB genome) and *T. timopheevii* (AAGG genome). Approximately 10,000 years ago, the second hybridization event occurred between *T. turgidum* (AABB genome) and the wild wheat species *Aegilops tauschii* (DD genome), which gave rise to *T. aestivum* (AABBDD genome)^22^.

Wheat occupies the largest total harvested area among cereal crops, yet its total productivity remains lowest^23^. The detrimental impact of various abiotic and biotic stresses on wheat plants throughout different growth stages results in significant production losses. Therefore, it is crucial to comprehend the effects of these stresses in order to drive advancements in wheat improvement programs. The availability of complete genome sequences, transcriptomics data, and the utilization of biotechnological approaches to unravel gene functions can open up new avenues for enhancing crop improvement.

Previous genome wide analysis of CrRLK1L gene family has been carried out in various plant species, including *Arabidopsis thaliana* (L.), *Oryza sativa L*., *Solanum lycopersicum, Nicotiana Benthamian* and *Gossypium raimondii*. Arabidopsis genome has 17 CrRLK1L encoding genes^7,24,25^ rice has 16^24^, tomato has 24^25^, *Nicotiana benthamiana* has 31^26^ and the diploid cotton species *G. raimondii* has 44 CrRLK1L genes ^27^. All these studies prompted us to further research this gene family in wheat (*T. aestivum*) to unravel their role in plant growth, development and stress responses. In this study, we performed analysis of conserved domains, gene structure, and functional motif analysis of CrRLK1l gene family members, along with inferring their evolutionary relationship with Arabidopsis and other monocot species. Additionally, we examined tissue-specific gene expression patterns and the changes in gene expression in response to abiotic and biotic factors using available transcriptome datasets. Through this research, we aim to gain insights into the possible roles of *Ta-CrRLK1L* genes in abiotic and biotic stress responses, provide a valuable resource for functional characterization and crop improvement.

## Results

### *In silico* analysis identified 15 CrRLK1L*s* gene family members in *T. aestivum*

In total, forty-three genes encoding CrRLK1Ls were identified in the hexaploid genome of *Triticum aestivum*, which consist of fifteen paralogous genes with three homeologous copies each, namely A, B, and D, from their progenitor species, except for *Ta*-*CrRLK1L2* and Ta-*CRLK1L7*, which had missing -*2-D* and -*7-A* copies. A blastn search within the genomes of diploid *T. urartu* and *T. dicoccoides* in the NR database at NCBI did not show high sequence similarity hits for these copies. However, the blastn search for *CrRLK1L2* at IWGSC RefSeq v2.1 and Ensembl plants had a hit with 88% identity within the coding regions of the genes on Chr3A, which is not acceptable for the *T. aestivum* homeologs. To confirm the presence of a -*3-D* copy in the diploid progenitor, a blastn search in the *Ae. tauschii* genome in NCBI data base was done, which also did not show a high sequence similarity hit and suggests that - 2-D may also be missing in *Ae. tauschii. Ta-CrRLK1L2-A* and *2-B* had 789 aa and 814 aa length sequences in the Ensembl database, which were more like transcript variants, and these sequences were corrected using TSA and wheat WGA_v0.4 scaffolds at the IWGSC database. -*7-B* homeolog had 630 aa and missing amino acids in the N-terminal region, which was also confirmed using the IWGSC database and whole genome shotgun (WGS) contigs at the NCBI database. However, the search in the *T. dicoccoides* genome, available in the NR database at NCBI, had full-length copy of the -*7-B homeolog*, which suggests that these changes likely happened after the speciation of T. aestivum. To further analyze the changes at the nucleotide level, the genomic sequence for -*7-B* from *T. dicoccoides* was retrieved from WGS and compared with the *-7-B* genomic sequence of *T. aestivum* using Clustal Omega. Interestingly, it was found that the truncation in the *-7-B* copy of *T. aestivum* was due to a C nucleotide deletion at the 182 nucleotide position in *T. aestivum* species, which caused the frameshift in the amino acid; moreover, the single nucleotide change (C to G) at the 303 nucleotide position in the coding region of *T. aestivum* resulted in a premature stop codon (**Fig. S1**). Sequences for *CrRLK1L*s were confirmed from at least two independent databases that include Ensembl Plants and the NR, TSA, and EST databases at NCBI. The transcript variants for Ta-*CrRLK1L* gene family members, -*11-B* (two variants), -*4-B* (two variants), and -*14-D* (4 variants), were identified from the Ensembl plant database, and the correct variant that codes for FL protein was selected for further analysis **(Table 1 and Table S2)**.

**Table 1.**
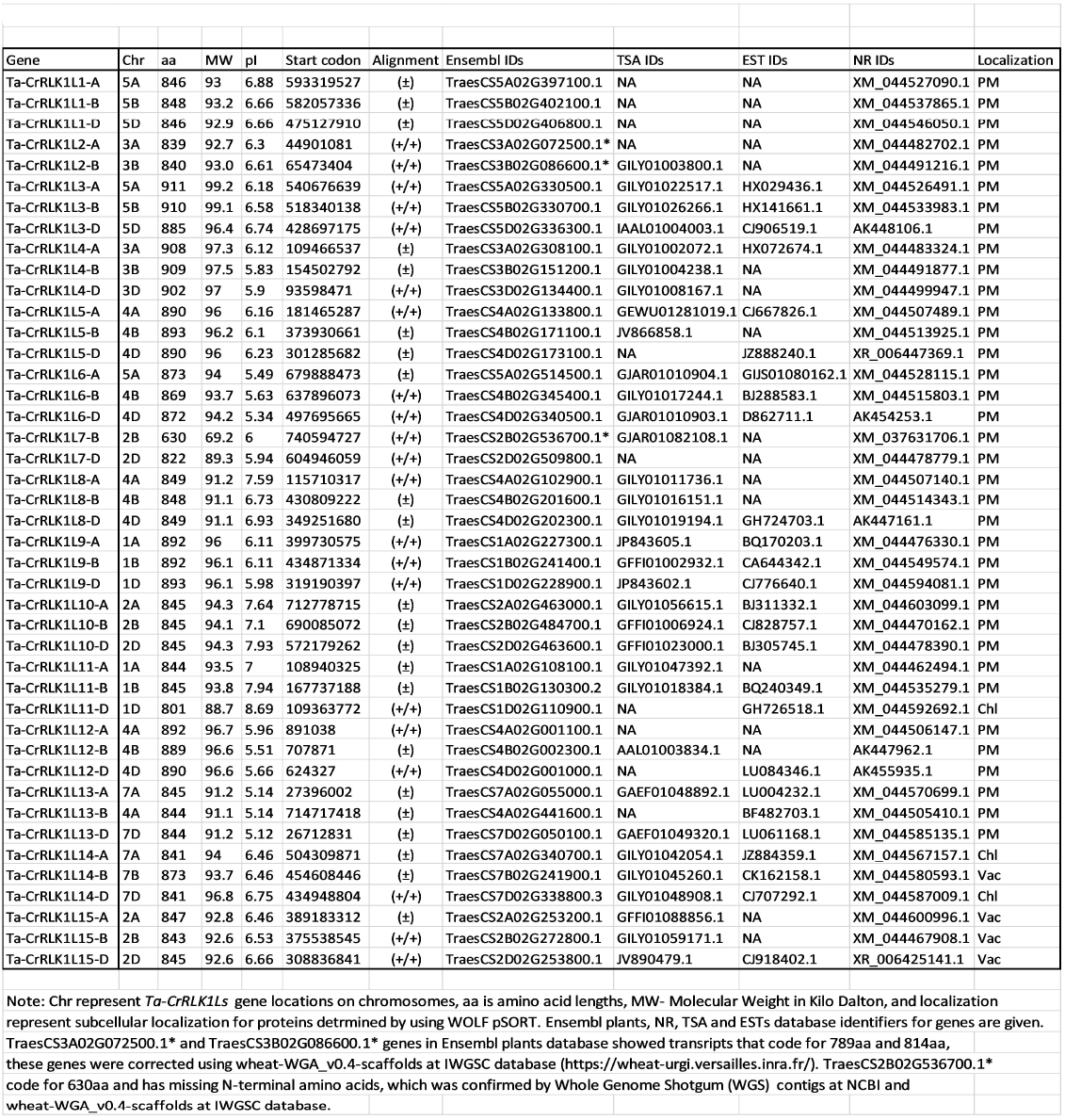
Sequence confirmation for *CrRLK1L* gene families in *Triticum aestivum*

The protein lengths and molecular weights for Ta-CrRLK1L were in the range of 801–911 aa and 88.7-99.2 KDa, respectively. *Ta-CrRLK1L6*-A and -*13-B* were translocated from -4A to 5A and 7B to 4A chromosomes, respectively. Most of the homoeologous copies of *Ta*-*CrRLK1L* showed more than 93.5% similarity in the protein coding regions within their homeologs, except for -*5-B* and -*5-D*, which had 91.0% similarity, respectively. The sub-cellular localization predicted by using Wolf pSORT showed that 36 Ta-CrRLKL1 were localized to the plasma membrane; surprisingly, *-14-B, -15-A, -15-B*, and *-15-D* were found localized to the vacuole, and *-9-D, -12-A*, and *-12-D* were localized to the chloroplast **(Table 1)**.

Blast searches using *T. aestivum CrRLK1L* protein coding sequences in monocot species *Ae. tauschii, B. distachyon*, and *H. vulgare* identified 14, 11, and 15 CrRLK1L encoding genes, respectively. The *CrRLK1L2-D* copy was missing in *Ae. tauschii* (**Table S3**).

### *Ta-CrRLK1Ls* members have highly conserved protein domains and show evolutionary conservation with closely related species

Multiple sequence alignment and conserved domain prediction using SMART showed that *T. aestivum* CrRLK1Ls had highly conserved characteristic domains that included a malectin-like domain (PF12819) and a catalytic domain for serine/threonine protein kinases (SM000220). The lengths of the malectin-like domain in Ta-CrRLK1L varied between 270 and 378 aa, except for the *-7-B* gene copy, which had a 198 aa malectin-like domain. This suggests that the point mutation caused the truncated malectin-like domain in the -7B copy of Ta-CrRLK1L. The lengths for the serine/threonine protein kinase domain ranged from 265 to 281 aa. Sequence analysis using PROSITE showed that *Ta-CrRLK1Ls* were comprised of featured domains that include an ATP-binding region signature (PS00107) and an active site signature for serine/threonine protein kinases (PS00108). The serine/threonine protein kinases had the active site residue (D) that is directly involved in the catalytic functions predicted by PROSITE (**Fig. 1)**. Phylogenetic analysis of CrRLK1Ls using full-length protein sequences showed that more closely related species like *T. aestivum, Ae. tauschii*, and *H. vulagare* were clustered together. The paralogs of Ta-*CrRLK1Ls* were clustered with Arabidopsis CrRLK1Ls, which have been known to be involved in plant reproduction and pollen-pistil interaction in Arabidopsis. At-*FER* was clustered with *Ta-CrRLK1L5, -9*, At-*ANX1* and At-*ANX2* were clustered with *-12, At-HERK2* was clustered with *-10, At-BUPS1* and *At-BUPS2* were clustered with *-13, At-CAP* was clustered with *-1*, and *At-HERK1* and At-*ANJEA* were clustered with *-4* and *-8*. (**Fig. 2 and Fig.S2**), which suggests that these homologs may also have a similar role in plant reproduction.

**Figure 1.**
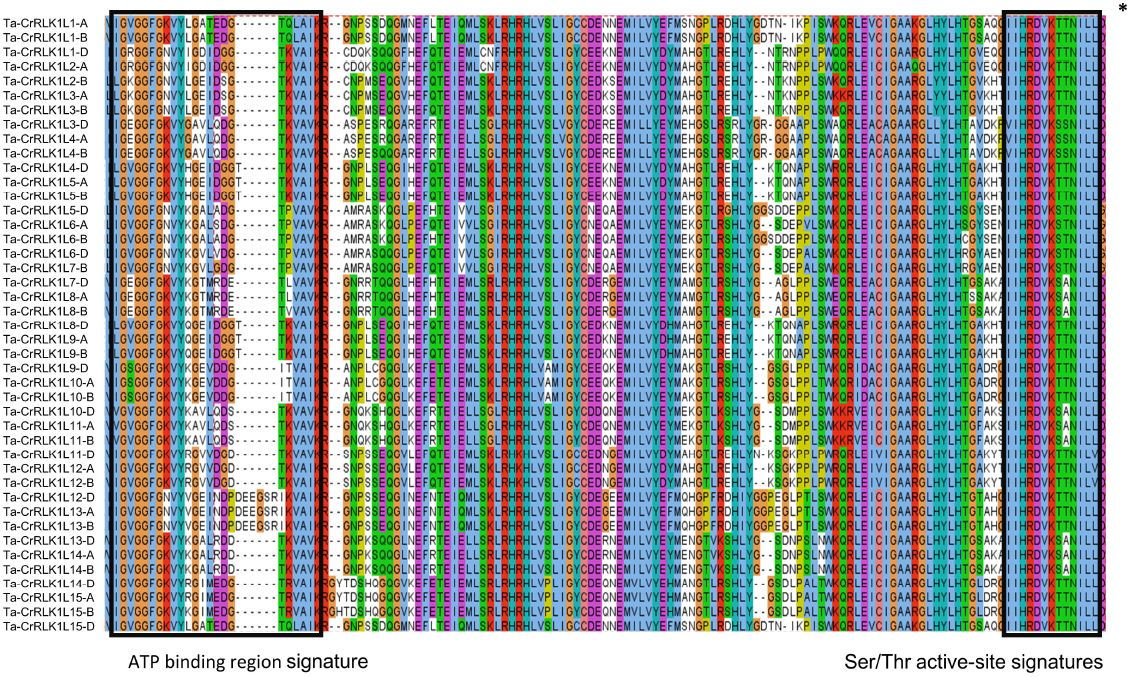
Multiple sequence alignment of Ta-CrRLK1L gene family members. Full length protein sequences were aligned by Clustal W and visualised by Jalview (https://www.jalview.org/). Conserved motifs in the protein kinase domain like protein kinases ATP-binding region signature (PS00107) and serine/threonine kinases active-site signatures (PS00108) determined by PROSITE (https://prosite.expasy.org/) are represented by boxes, and active site residue (D) is marked by an asterix.

**Figure 2.**
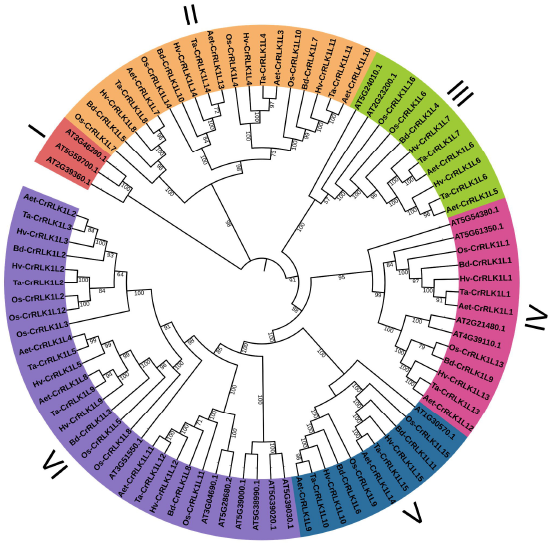
Phylogenetic analysis of CrRLK1L in *T. aestivum* and other species. This analysis involved 88 amino acid sequences. All positions with less than 95% site coverage were eliminated, i.e., fewer than 5% alignment gaps, missing data, and ambiguous bases were allowed at any position. There were a total of 622 positions in the final dataset. Evolutionary analyses were conducted in MEGA11. Phylogenetic stress was visualized by iTOL (https://itol.embl.de/).

### Ta-CrRLK1Ls display varied distribution in seven chromosomes and is absent in chromosome 6

Chromosomes 4A, 4B, and 4D had the highest distribution of CrRLK1L genes and showed the distribution of four genes that include homeologs for *Ta-CrRLK1L5, -6-B* and *-6-D*, and *- 8* and -*12*; however, this included the -*13-B* homeolog, which has translocated from 7B to 4A chromosome (**Fig. 3 and Table 1)**. Chromosomes 3D and 7B had a distribution of single *CrRLK1L* genes that included *-4-D* and *-14-B*, respectively. Chromosome 6 did not have any distribution of *CrRLK1L* genes and hence was eliminated. The intron/exon junctions determined by Splign showed that most *Ta*-*CrRLK1Ls* had no introns and were comprised of single exons, except for the paralogs -*11-D* and homeologs of -*14*, which have a single intron and two exons (**Fig. 4B and Table S4**). *-11-D* copy had the smallest and largest introns of size 124 bp and 1149 bp, respectively. The homeolog of *-14* had shorter first exons of 36 bp, 36 bp, and 111 bp for *-14-A, -14-B*, and *-14-D*, respectively. Motifs predicted by the MEME suite were mostly related to serine/threonine protein kinase domains (**Fig. 4 C and D, and Table S5**).

**Figure 3.**
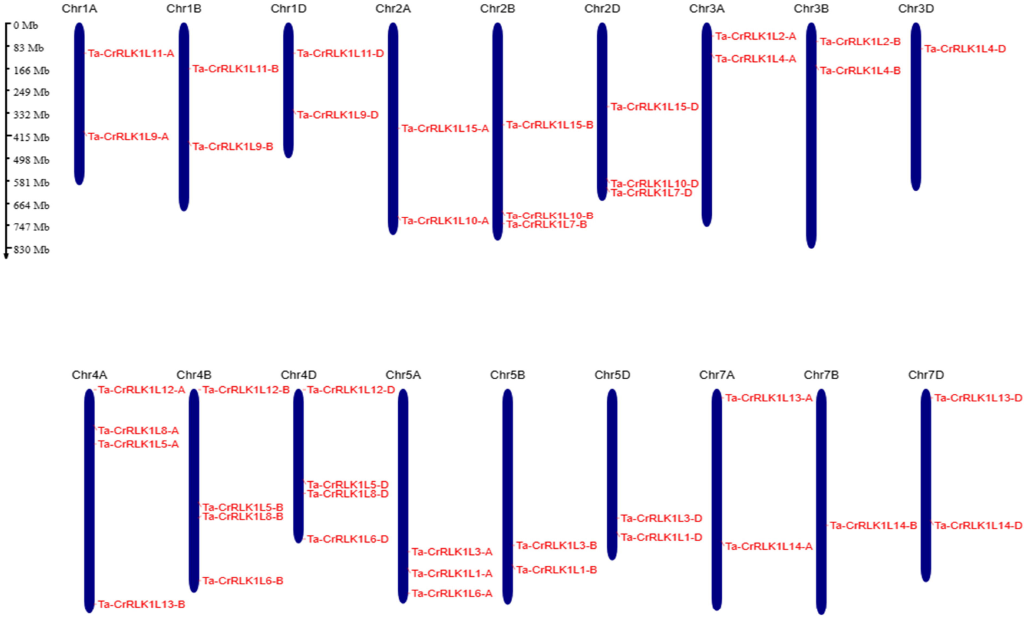
Distribution of CrRLK1L genes across *Triticum aestivum* chromosomes. The distribution of *Ta-CrRLK1L* genes on six chromosomes is given, and gene names are represented in red. Chromosome 6 did not have the distribution of the CrRLK1Ls. Genes were mapped by their physical locations, and the scale bar given is in megabases. Gene positions on the chromosomes were visualized using MG2C v2.1 (http://mg2c.iask.in/mg2c_v2.1/).

**Figure 4.**
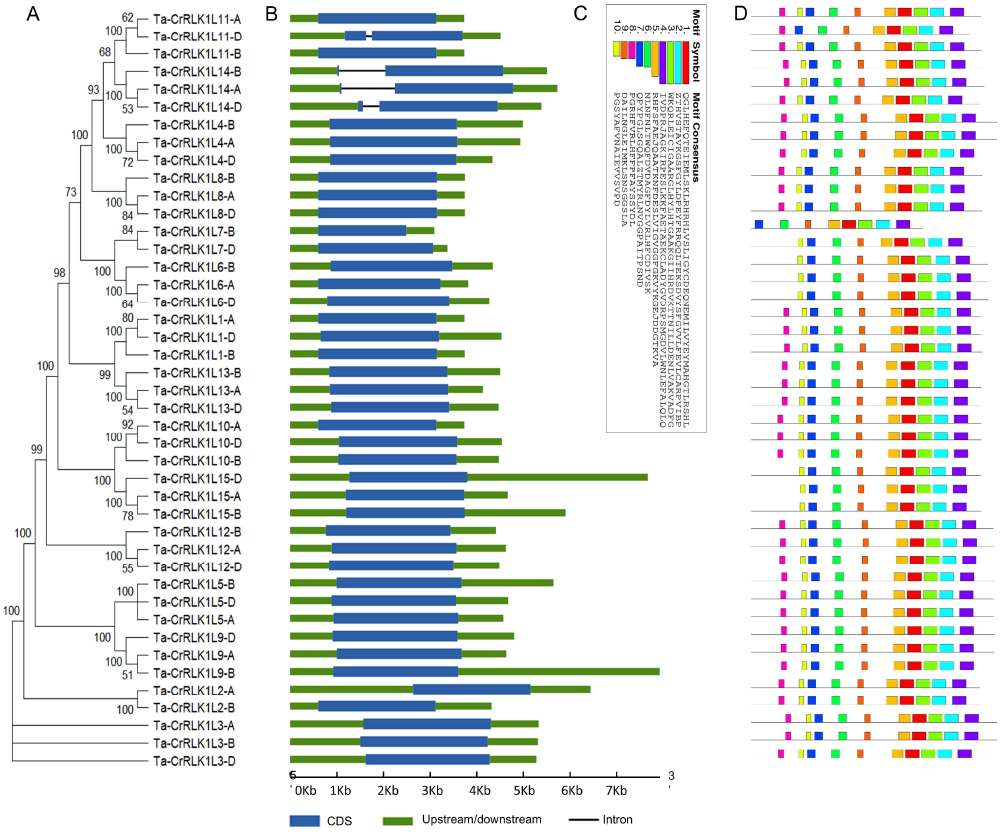
Structural organization of Ta-CrRLK1L gene families in *Triticum aestivum*. A) A phylogenetic tree was constructed using Neighbor-joining method and 1000 bootstrap iterations by MEGA 11 ((https://www.megasofware.net/). B) lntron/exon structure for gene family members was displayed using the Gene Structure Display Server (http://gsds.qao-lab.org/). Blue boxes are the exons, green is the upstream or downstream region, and lines between boxes denote intrans. (C) and (D) are the ten identified motifs in Ta-CrRLKLs protein sequences by using MEME Suite 5.5.2 (https://meme-suite.org/meme/index.html).

### *Ta-CrRLK1L* transcripts show altered expression in different tissues with *CrRLK1L-12* and -*13* highly expressed in the anther

The relative level of gene expression analyzed across a panel of seventy-one tissues at different growth and developmental stages of plants in Azhurnaya spring wheat showed that *CrRLK1Ls* gene families in *T. aestivum* had diverse expression patterns. Gene expression values were in the range of 0 to 160 <u>T</u>ranscripts <u>P</u>er <u>M</u>illion (TPM). It is known that either one or two copies of homeologs have a higher level of expression than other copies^28^, and a similar pattern of expression has been found in *Ta-CrRLK1L* gene family members. *Ta-CrRLK1L*-*12 and* -*13* only showed a higher level of gene expression in anther tissues. The homeologs *CrRLK1L-13* are highly expressed in anther tissues, and out of the three homeologs, *-13-D* had the highest relative level of gene expression with a value of 158.4 TPM, which was also the highest expressed paralog among the Ta-CrRLK1L gene family members. Ta-*CrRLK1L*-12 was another moderately expressed paralog in the anther tissues, and *-12-B* had a higher gene expression level of 61 TPM compared to other homeologs (**Fig. 5a and Table S6**). In phylogenetic analysis, *Ta-CrRLK1L-12* was clustered together with *At-ANX1* and *At-ANX2*, and *Ta-CrRLK1L-13* was clustered with *At-BUPS1 and At-BUPS2* (**Fig. 3**), which are known to be involved in maintaining the pollen tube integrity and prevention of the premature rupture of the pollen tube before reaching the synergid cells (Bordeleau et al., 2022). This suggests that *Ta-CrRLK1L-12* and *-13* may also be involved in the maintenance of pollen tube integrity in wheat. *Ta-CrRLK1L5, -9, and -14* were moderately expressed in the stigma and ovary tissues. Phylogenetic analysis of Ta-CrRLK1L showed that CrRLK1L5, -9, and -14 were clustered together with At-FER, and -14 was clustered with At-HERK1 and At-ANJ, which are known to be involved in receiving the pollen tube at synergid cells during the reproduction^9^. *Ta-CrRLK1L5, -9*, and *-14* expression in the stigma and ovary tissue suggests the possibility for their role in the pollen tube reception during fertilization. *Ta-CrRLK1L3* was highly expressed in the leaves, and *-3-D* had a higher expression compared to its homeologs and was highly expressed in the flag leaf sheath at the ear emergence stage with a value of 118.5 TPM.

**Figure 5.**
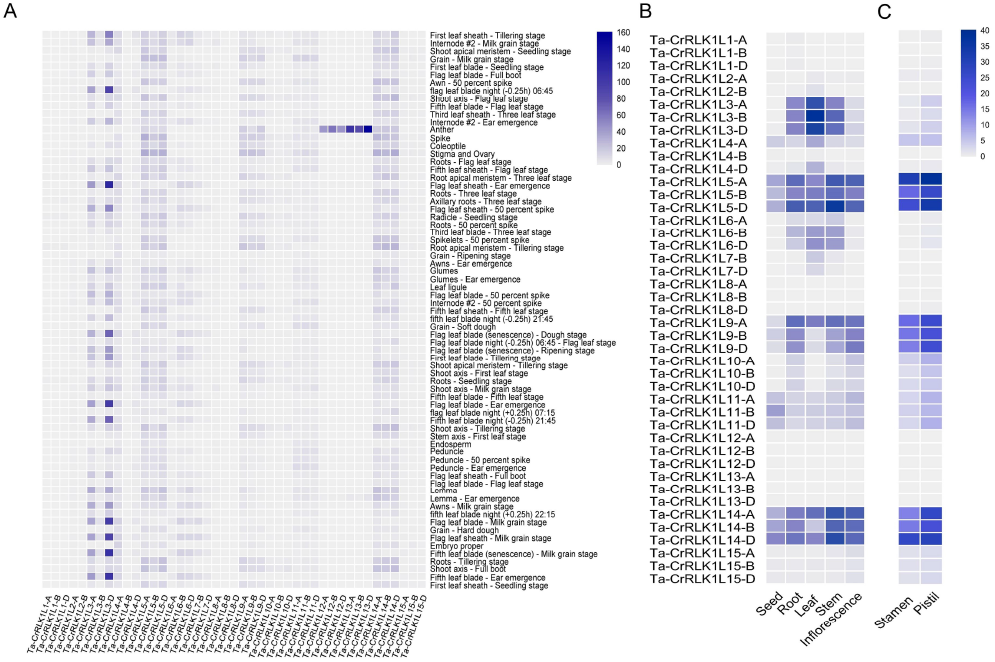
Gene expression analysis of CrRLKll gene families in *Triticum aestivum*. Heatmaps represent the relative level of gene expression analyzed across A) Seventy one tissues in Azhurnaya spring wheat by using the wheat eFP browser (https://bar.utoronto.ca/). Values are expressed as <u>T</u>ranscripts <u>P</u>er <u>M</u>illion (TPM). B) Relative gene expression levels in five tissue types namely seed, root, leaves, stem and inflorescence were analyzed from the transcriptome datasets by Pingault et al. (2015) and C) Gene expression in immature stamen and pistil tissues was analyzed from the transcriptome datasets with Bio Project accession PRJEB36244 from the SRA database at NCBI. Values for B and C are expressed as <u>F</u>ragments <u>P</u>er <u>K</u>ilobase Per <u>M</u>illion (FPKM).

The relative level of gene expression analysed across five tissue types was in agreement with the analysed expression for seventy-one tissue types for *Ta-CrRLK1L3* and showed that it was highly expressed in leaf at whole plant fruit formation (30–50% moisture stage), and the expression was in the range of 29.2 to 36.3 FPKM (**Fig. 5B and Table S7**). In addition, the homeologs of *-3* also had moderate levels of gene expression in the root at the cotyledon emergence stage and in the stem at the two node or internode stage tissues. *Ta-CrRLK1L5* and *-14* were expressed in all five tissue types with different levels of expression, and *-5-B* was least expressed in seed with 8.8 FPKM, and *-5-D* was highly expressed in stem with 34.5 FPKM.

The role of members of the *CrRLK1L* gene families in Arabidopsis has been well studied, and they have been known to act as carriers for the successful pollen-pistil interaction in the plant’s reproduction. *At-ANX1* and *At-ANX2* have been known to function in the maintenance of pollen tube integrity, and *At-FER, At-HERK1, At-HERK2*, and *At-ANJ* are known to function in the pistil, mostly in the synergid cells. These proteins form complexes with each other and carry out the successful plant reproduction processes in Brassicaceae. To determine the possible role of *Ta-*CrRLK1Ls in plant reproduction, the relative levels of gene expression was analysed in the immature stamen and pistil tissues, and it was found that *CrRLK1L5, -9*, and -*14* has a relative higher level of gene expression compared to other gene family members in both stamen and pistil tissues; however, the gene expression levels in pistil tissues were higher than in stamen tissues. *-5-A* and *-14-D* had a higher level of expression than their homeologs and has gene expression values of 38.3 and 29 FPKM in pistil tissues, respectively. This was in agreement with the 71 tissues transcriptome data, where similar paralogs were expressed in stigma and ovary tissues (**Fig. 5C and Table S8**). Similar paralogs were also expressed in stamen tissues at values of 27 and 31 FPKM. Expression of *Ta-CrRLK1L5, -9*, and -*14* genes in both stamen and pistil tissue suggests the dual role of these genes on the male as well as female sides of the pollen-pistil interaction.

### Expression of Ta-CrRLK1Ls are altered in response to environmental stimuli

Most of the *Ta-CrRLK1L* paralogs were upregulated or downregulated in response to one or more abiotic factors. Changes in the level of gene expression in response to cold treatment analyzed in the wheat cultivar Manitou show induction as well as downregulation for the *Ta-CrRLK1L* paralogs. In response to the cold treatments, the paralogs *Ta-CrRLK1L4-B* and *-15-A* were up-regulated by 3.0 and 3.3-fold, respectively. *-9-B* and *-9-D* were other paralogs that were upregulated by 2.14 and 2.28-fold respectively. The paralogs that were downregulated to 15%, 18%, and 21% compared to control conditions included -*3-A, -7-B*, and *-11-A. Ta-CrRLK1L-3-A* had the highest 18% decrease in the gene expression level from 24.69 FPKM to 4.42 FPKM. Other paralogs had a 30–47% decrease in the gene expression level in response to cold stress, which included one of the homeologs of *Ta-CrRLK1L3, -4, -6, -7*, and -*11* (**Fig. 6A and Table S9)**. Surprisingly, *-7-B*, which had a premature stop codon, was the most downregulated paralog; which suggests that the nucleotide level changes in the N-terminal region, which encodes a malectin-like domain, did not affect its response to the cold stress treatment.

**Figure 6.**
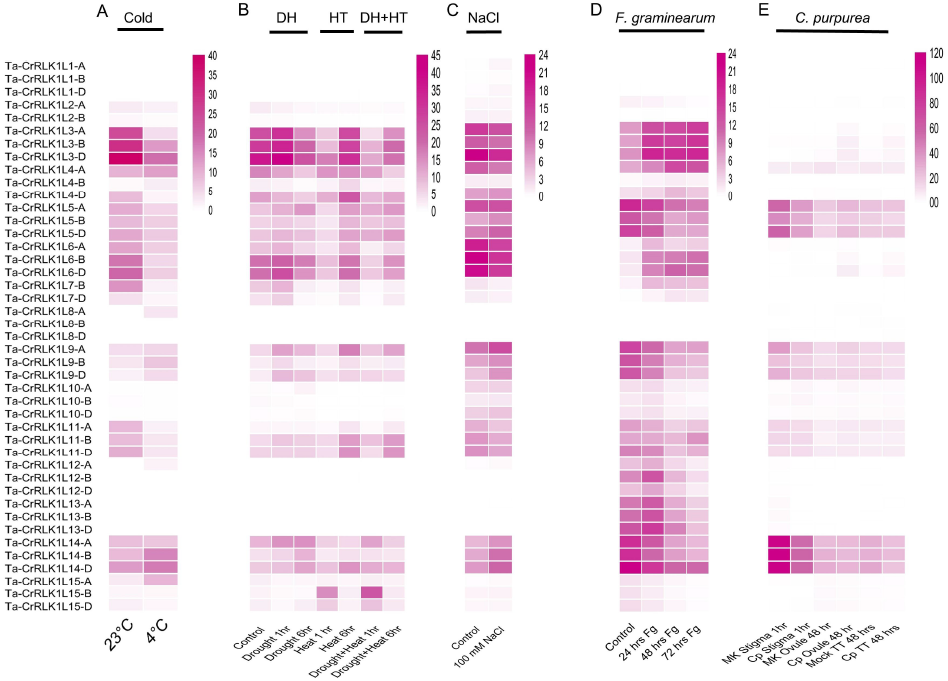
Gene expression analysis in response to abiotic and biotic stresses. The upregulation or downregulation of *Ta-CrRLK1 L* genes in response to abiotic factors like A) cold, B) drought, heat, and combined, and C) salt stress, and biotic factors like infection with D) *F. graminearum* and E) C. *purpurea* were analyzed using transcriptome datasets at NCBI SRA. The scale represents the FPKM values.

The up-regulation or downregulation of gene expression in response to drought, heat, and combined stress was analyzed from the publicly available transcriptome dataset^29^. The homeologs of *-9*, and paralogs *-9-D, -10-A*, and *-14-B* were induced by more than 2.3-folds in response to drought stress of 1 hr and 6 hr, respectively. Other paralogs that were downregulated after 1 hr or 6hr of drought stress were homeologs of *-2, -4, -7, -9, -10*, and *- 15, and -4-B* had the highest downregulation of 25% in the 6 hr drought stress treatment. The paralogs that were upregulated in response to 1 hr or 6 hr of heat stress were -*4-B, -9-A, -10-D, -11*, and *-15-B* and *-15-D*. Downregulation for the paralogs *-2, -3, -4, 6, -7, -9, -10*, and *- 11* in the same treatments was found, and *-10-B* was downregulated to 1.04 to 0.04 FPKM. The upregulation for the paralogs *-4-A, -9-A, 9-D, -11, -15-B*, and -*15-D* was observed in 1 hr or 6 hr combined stress, and the highest expressed paralog *-15-B*, was induced by 13.9-fold. The homeologs of *-7* and *-10* had the highest downregulation, and *-10-B* was downregulated from 1.04 FPKM to 0.02 FPKM. The homeologs *-7-B* and -7*-D* were downregulated to 12% of their control, which had 7.28 and 3.94 FPKM values (**Fig. 6B and Table S10**).

Changes in gene expression analysed in response to 100 mM NaCl treatments in the Chinese spring cultivar using transcriptome datasets from BioProject accession PRJNA522013 showed the upregulation or downregulation of *Ta-CrRLK1L* paralogs in one of the treatments. *Ta-CrRLK1L1* homeologs were induced in response to 100 mM NaCl treatment, and *-1-A* had the highest induction of 5.77-fold compared to *-1-B* and -*1-D*, which were induced by 3.35 and 2.93-fold, respectively; however, the lower expression levels of 0.12, 0.08, and 0.23 FPKM for these homeologs were detected in control treatments, respectively (**Figure 6C and Table S11**). The analysis of upregulated and downregulated gene expression in response to stress conditions suggests that *Ta-CrRLK1L* may have a role under these stress conditions.

### Expression of *Ta-CrRLK1L* transcripts are altered in response to head blight and ergot funguses

Changes in gene expression for *Ta-CrRLK1L* gene family members were analysed by transcriptome datasets for *F. graminearum* infection and *C. purpurea* infection, which cause head blight and ergot disease in wheat, respectively.

In response to *F. graminearum* (*Fg*) inoculation, *Ta-CrRLK1L-7D* was highly induced from 0.08 FPKM in control to 0.81, 1.85, and 1.75 FPKM in 24 hr, 48 hr, and 72 hr post-inoculation treatments. The homeologs of *Ta-CrRLK1L6* and -*7-D* were also upregulated in all three treatments, and *-6D* was induced by a 6.93-fold increase 48 hr post-*Fg* inoculation. However, in response to inoculation after 72 hr, the *-13* homeologs were downregulated by 11% and 10% of their control values, respectively. *Ta-CrRLK1L10*, -*11*, and -*12* homeologs were also downregulated after 72 hr of inoculation, and the downregulation of these genes was in the range of 30–48% of their control values (**Fig. 6D and Table S12**).

*C. purpurea* (*Cp*) infection in the stigma induced *-6-D* and *-15-D* paralogs from 0.06 FPKM in mock to 0.50 and 0.57 FPKM in 1 hr post inoculation treatment, whereas the homeologs *- 12-A and -12-D* were downregulated to 14% and 11% in the treatment compared to their control (**Fig. 6E and Table S13***). Ta-CrRLK1L-3* and *-6* homeologs were induced in response to 48 hr post *Cp* inoculation in the ovule, and -*6-D* and *-3-A* were induced by 10.92 and 10.26-fold,, however, in control conditions, *-3-A* and *-6-D* have lower expression of 0.14 and 0.06 FPKM, respectively. -*6-D and -3-A* were also upregulated in 48 hr post-inoculation ovary transmitting tissue, to a change of 7.04 and 4.26-fold, respectively.

### Various stress-responsive and hormonal responsive cis-acting elements regulating plant development are present in the promoter regions of CrRLK1Ls

Prediction of the cis-acting elements in the putative promoter regions of *Ta-CrRLK1L* genes showed that 54% of the cis-acting elements were comprised of the core promoter and enhancer region elements, including TATA and CAAT boxes. The remaining 46% were divided into three categories that include cis-acting elements involved in hormonal responses, plant developmental processes, and abiotic and biotic stress conditions. These were again divided into subcategories; for example, hormonal responses were divided into abscisic acid, auxin, gibberellin, and salicylic acid respectively. Similarly, other elements were also sub-grouped (**Fig. 7 and Table S14**). Cis-acting elements for light and abscisic acid were found in all *Ta-CrRLK1L* genes. *Ta-CrRLK1L1* had the cis-acting elements for palisade mesophyll cell differentiation, and -*14* had meristem-specific activation elements, which were grouped into cis-acting elements in the plant developmental processes. The paralogs -*9-B, -12-A, -12-D, -13-A, and -13-B* had elements for seed-specific regulation. Cis-acting elements for defense and stress were predicted in all *CrRLK1Ls* except *-10, -11, and -14*.

**Figure 7.**
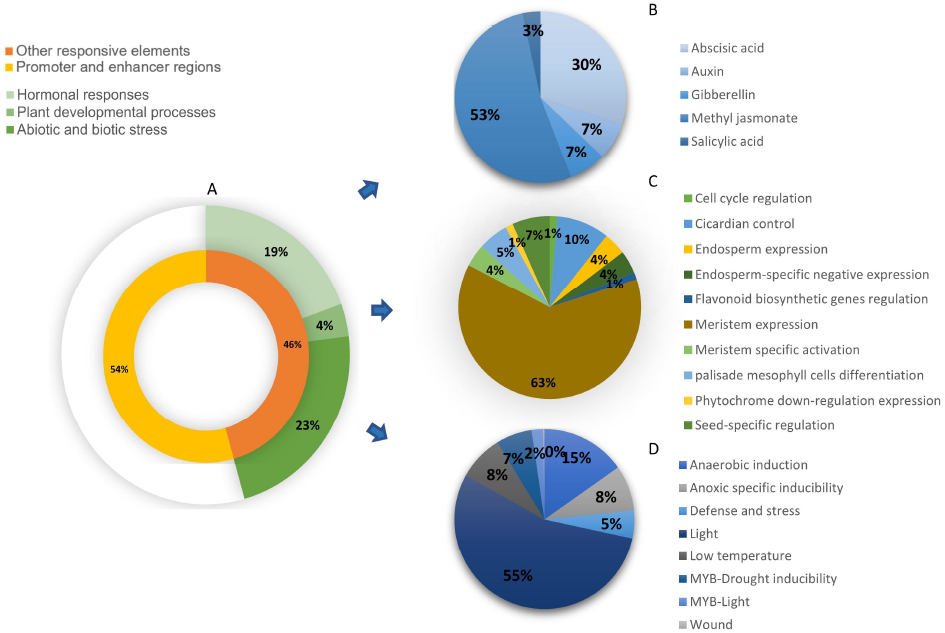
Distribution of the predicted cis-acting elements in Ta-CrRLK1 L gene families. The cis-acting elements predicted by analysis of 2kb upstream regions of genes by PlantCARE (https://bioinformatics.psb.uqent.be/webtools/plantcare/html/) were categorized into A) Promoter and enhancer regions and other responsive elements. Other responsive elements were categorized into three main subgroups that include cis-acting elements involved in B) Hormonal responses, C) Plant developmental processes, and D) Abiotic and biotic stress.

## Discussion

The CrRLK1L gene families in *T. aestivum* was comprised of 15 members, and it total 43 paralogous genes. Most of the members showed more than 93.5% similarity in the coding region of the genes, which has been reported in several studies^28^. The paralogs *-2-D* and *-7-A* in the progenitor species *Ae. tauschii* and *T. urartu* or *T. dicoccoides* did not have high sequence similarity in the coding regions, which suggests that these gene copies may also be absent in the progenitor species. However, the presence of the single nt deletion at the 182 nt position and premature stop codon in the coding region of the *-7-B* copy in the *T. aestivum* genome and the presence of the FL length gene in the *T. dicoccoides* suggest that these changes in *T. aestivum* were recent events and occurred after the speciation of *T. aestivum*. The erroneous sequences for *CrRLK1Ls* from Ensembl Plants databases were corrected by wheat genome shotgun (WGS) contigs sequences at the NCBI database. The reverse translocations in the *T. aestivum* genome are most commonly observed, and these include the translocation between 4AL/5AL and 4AL/7BS^29^, which were observed in our gene family members Ta-CrRLK1L6 and -13. Surprisingly, the homelogs -*13-D* and -*14* had the presence of the introns, which were also present in *T. dicoccoides* when searched by blast in the NCBI WGS assembly.

CrRLK1L gene families in Arabidopsis consist of 17 members, of which FER, ANJ, HERK1, BUPS1/2, and ANX1/2 are the critical players in plant reproduction and have roles in guiding the pollen tube to the synergid cells of the ovule for fertilisation. The FER-ANJ complex acts in the stigma papillae and has an important role in pollen germination, where it is proposed to maintain the ROS levels in the stigma papilla in the absence of the pollen and act as a negative regulator for stigma hydration^30^ Arabidopsis CrRLK1L members BUPS1/BUPS2 that complex with ANX1 and ANX2 act on the pollen tube side and function in the maintenance of pollen tube integrity and prevent its rupture before it reaches the synergid cells^8^. Phylogenetic analysis of CrRLK1L genes showed that ANX1/2 were clustered with Ta-CrRLK1L12, and BUPS1/2 were clustered with Ta-CrRLK1L13, which suggests that these genes are the Arabidopsis homologs for ANX and BUPS in wheat. This was supported by the tissue-specific gene expression analyzed across the seventy-one tissues of wheat, where *Ta-CrRLK1L12* and *Ta-CrRLK1L13* were highly expressed in anther tissues. However, the gene expression analysed by using the transcriptome datasets of immature stamen or pistil tissues around the meiosis of wheat flowers at Zadok stages 41-49 did not expressed these genes, which suggests the specificity of these genes to function in the later stages of plant reproduction, when the pollen germinate on the stigma. *Ta-CrRLK1L5, -9*, and -*14* were also expressed in stigma and ovary tissues in seventy-one tissue types and had a relative level of gene expression in the range 18.25–32 FPKM, 15.12–15.38 FPKM, and 21.16–28.17 FPKM. Similar genes were also expressed in the analysed transcriptome datasets for immature stamen and pistil and in five tissue types transcriptome datasets. The clustering of these genes suggests that *-5* and *-9* belong to the FER, and *-14* belong to HERK, CURVY1, and ANJEA. *Ta-CrRLK1L-1* and *-14* copies showed the presence of specific palisade mesophyll cell differentiation and meristem-specific activation cis-acting elements in their promoter regions, which indicate that these genes may have a role in plant growth and regulation.

FER is a versatile CrRLK1L that functions in plant growth and development, abiotic stress, hormonal signalling, and plant immunity responses^31^. In Arabidopsis, the responses to stress conditions like drought, cold, and heat in the FER mutant, *fer-4*, showed that FER acts as a negative regulator of drought stress and a positive regulator of the cold and heat stress responses^3,32^. To determine the response of *Ta-CrRLK1L* genes, we analyzed the gene expression from the available transcriptome datasets. Abiotic factors included cold, drought, heat, combined stress, and salt stress responses. The cold stress-responsive gene -3-A was clustered with FER, which had the highest downregulation (18% downregulated compared to control) in response to cold stress. Another downregulated gene was the homeolog of -*6-A*, which was clustered with AT5G24010.1 and AT2G23200.1. *Ta-CrRLKL115-A* was induced more than 3.3-fold and clustered with HERK2 in Arabidopsis. At least one of the homeologs of -*3, -4, -6*, and -*9* was downregulated in response to 1 hr of heat and 1 hr of combined stress, whereas -*15-D* was induced in the similar treatments. Among these genes, most were clustered with FER, HERK1, and ANJEA, whereas the upregulated copy *-15-D* was clustered with HERK2. *Ta-CrRLK1L12* and *-13* also responded to *F. graminearum* infections, suggesting that these BUPS1/2 and ANX1/2 homologs may have a role in the head blight caused by *F. graminearum*. HERK2 homolog *Ta-CrRLK1L15* displayed responses to varied stress conditions, including cold, heat, combined drought and heat, and *F. graminearum* infection. Cis-acting element prediction in the promoter region of *-15* had defence and stress-responsive cis-acting elements, and cis-acting elements for abscisic acid responsiveness were found in all the gene copies. The homolog of AT5G24010.1 and AT2G23200.1, *Ta-CrRLK1L6*, was another gene that had wide expression responses to abiotic and biotic factors. Altogether, our results suggest that the CrRLK1L gene family in *T. aestivum* is involved in plant developmental processes and as well as plant adaptation in response to abiotic and biotic stresses.

## Material Methods

### Collection of sequences for *CrRLK1L* gene families in *Triticum aestivum*

Full-length coding and protein sequences for the 16 CrRLK1L from rice were retrieved from the Rice Genome Annotation Project database (http://rice.plantbiology.msu.edu/)^24^. CrRLK1L homologs in wheat, hereafter denoted as *Ta-CrRLK1L*, were searched by using rice FL coding sequences in the NR database at NCBI (https://www.ncbi.nlm.nih.gov/) in *Triticum aestivum* sp. Sequences with high scoring hits and maximum similarity with rice *CrRLK1Ls* were collected, and duplicates were eliminated. The NR sequences were used to search for respective A, B, and D homeologs in the Ensembl plants database (https://plants.ensembl.org/index.html), and genomic, CDS, and protein sequences for Ta-CrRLK1L with high scoring hits and more than 95% identity were collected. Transcript variants that cover the full-length coding sequences were selected. To confirm the homeologous copies, the Ensembl plant-retrieved sequences of *Ta-CrRLK1L* were used to search against the TSA and EST databases at NCBI. The start codon position in the wheat genome for each *Ta-CrRLK1L* homeolog was determined by blastn search of coding regions in the International Wheat Genome Sequencing Consortium (IWGSC) RefSeq v2.1 genome assembly at the WHEAT URGI database (https://wheat-urgi.versailles.inra.fr/). The physical properties, like the molecular weight of the protein and isoelectric point (pI), were calculated using the Expasy Compute pI/MW tool (https://web.expasy.org/compute_pi/). Subcellular localization for the proteins was predicted by WoLF PSORT (https://wolfpsort.hgc.jp/).

### Compilation of gene sequences for CrRLK1L from other plant species

Genes encoding CrRLK1L in closely related species like *Aegilops tauschii* and *Hordeum vulgare* were retrieved from the NCBI database by blastn search using wheat FL coding sequences. *Brachypodium distachyon* sequences were collected from Ensembl plants, and sequences for CrRLK1L gene families in Arabidopsis were retrieved from the TAIR database (https://www.arabidopsis.org/). Rice sequences were collected from the Rice Genome Annotation Project database (http://rice.plantbiology.msu.edu/).

### Conserved domain and phylogenetic analysis

Conserved domains for the CrRLK1L proteins were confirmed by the Simple Modular Architecture Research Tool (SMART) (http://smart.embl-heidelberg.de/) and Batch Conserved Domain (CD)-Search tool (https://www.ncbi.nlm.nih.gov/Structure/cdd/wrpsb.cgi) at NCBI, and active sites in the Ta-CrRLK1L domains were predicted by PROISTE (http://prosite.expasy.org/). Multiple sequence alignment for Ta-CrRLK1Ls were carried out by ClusatlW, and conserved domain features and active sites were visualized using the Jalview program (https://www.jalview.org/)^33^. For phylogenetic analysis, the multiple sequence alignment for protein sequences was carried out by ClustalW (https://www.ebi.ac.uk/Tools/msa/clustalo/), and the evolutionary relationships for CrRLK1L in *T. aestivum* and other five species, namely *Ae. tauschii, A. thaliana, B. distachyon, H. vulgare*, and *O. sativa*, were inferred by the analysis of full-length amino acid sequences in MEGA11^34^. *Ta-CrRLK1L* A homeolog copies were used in the analysis, except for *Ta-CrRLK1L7*, whose D copy was used. In total, eighty eight CrRLK1L protein sequences were used in the analysis. Sequences were aligned by ClustalW with default parameters, and a phylogenetic tree was constructed using the Jukes-Cantor model by the Neighbour-joining method and 1000 bootstrap iterations. The phylogenetic tree was visualised by iTOL (https://itol.embl.de/login.cgi).

### Chromosomal location, gene structure prediction, and motif analysis

Chromosome length for each copy in wheat was collected from the Ensembl Plants database, and start and end positions for *Ta-CrRLK1L* genes were determined from the (IWGSC) RefSeq v2.1 genome assembly. The position of the *Ta-CrRLK1L* genes on the chromosomes was represented by using MG2C v2.1 (http://mg2c.iask.in/mg2c_v2.1/)^35^. Intron/exon junctions were determined by comparing *Ta-CrRLK1L c*DNA sequences from the NR database at NCBI with genomic sequences from Ensembl by using Splign (https://www.ncbi.nlm.nih.gov/sutils/splign/splign.cgi?textpage=online&level=form), and the intron/exon junctions were visualised by Gene Structure Display Server 2.0 (http://gsds.gao-lab.org/). Conserved motifs in CrRLK1L proteins were predicted using MEME Suite 5.5.2 (http://meme-suite.org/tools/meme) with ten motifs as the number of motifs parameter and the GenomeNet Database (https://www.genome.jp/tools/motif/).

### Cis-acting elements prediction

The nucleotide sequences from the upstream 2 Kb regions of the start codons of *Ta*-*CrRLK1L* genes were retrieved from the Ensembl plant database and analyzed for the presence of cis-acting elements. Cis-acting elements were determined by using PlantCARE (https://bioinformatics.psb.ugent.be/webtools/plantcare/html/).

### Gene expression analysis of Ta-CrRLK1L gene families

Tissue-specific gene expression and gene expression in response to stress conditions for Ta-*CrRLK1L* gene families were analysed using *T. aestivum* transcriptome datasets available at SRA (Sequence Read Achieve) repositories in the NCBI database.

To determine the tissue-specific relative gene expression levels, the transcriptome dataset of 71 tissues of wheat cultivar Azhurnaya, available at eFP Browser (http://bar.utoronto.ca/efp_wheat/cgi-bin/efpWeb.cgi) was used^36^ The relative gene expression levels in five tissue types, namely, inflorescence, leaf at the whole plant seed formation stage, root at the cotyledon emergence stage, seed at the fruit ripening stage, and stem, were analysed from the transcriptome datasets available at SRA database at NCBI^37^. Tissue-specific expression in the stamen and pistil tissues was analyzed by using BioProject: PRJEB36244 datasets at SRA repositories (https://www.ncbi.nlm.nih.gov/sra). Differential gene expression in response to abiotic factors like cold^38^ drought, heat, and combined stress^39^ and salinity stress induced by NaCl (Bio Project Accession: PRJEB36244) was carried out by using the transcriptome datasets available at SRA repositories at NCBI. Similarly, the transcriptome datasets from SRA that determine the changes in gene expression in response to infection caused by *Claviceps purpurea* and *Fusarium graminiearum* were used (**Table S1**).

The FPKM (<u>F</u>ragments <u>P</u>er <u>K</u>ilobase <u>P</u>er <u>M</u>illion) changes in gene expression were determined by search of *Ta-CrRLK1L* gene ensemble plant database identifiers against the *T. aestivum* RNA-seq Database (http://ipf.sustech.edu.cn/pub/wheatrna/) at the Plant Public RNA-seq Database^40^, and the search was restricted to SRA accessions of biological replicates for each dataset. FPKM values for the relative level of gene expression and the fold change in response to stress conditions were visualized by TBtools^35^

## Conclusions

We conducted an extensive in silico genome-wide analysis of the *CrRLK1L* gene family in wheat, providing valuable insights into this family’s characteristics in wheat for the first time. In the wheat genome, we identified a total of 15 *CrRLK1L* genes and 43 paralogs. Our analysis revealed a frameshift mutation and a premature stop codon occurrence in the *-7-B* copy, resulting in its truncation. Furthermore, our investigation of the *TaCrRLK1L* genes revealed the presence of the various cis-acting elements involved in specific biological processes, such as plant growth, development, and stress responses. By analyzing publicly available transcriptome data sets, we found diverse expression patterns of most *TaCrRLK1s* in different tissues, including reproductive tissues. Moreover, in response to abiotic or biotic factors, most of the *CrRLK1L* genes were upregulated or downregulated. The data presented in our study will serve as a valuable resource for future functional characterization and validation of CrRLK1L proteins in wheat. Additionally, this knowledge will facilitate the development of targeted strategies for crop improvement, leveraging the potential of the CrRLK1L gene family.

## Acknowledgements

We thank the Indian Institute of Technology Gandhinagar for a post-doctoral fellowship to NG. This work was supported by a DBT Ramalingaswamy Re-entry fellowship grant and start-up grant from Indian Institute of Technology Gandhinagar to SS.

## Author Contributions

NG and SS conceived and designed research; NG performed all the experiments; SS supervised the experiments; NS and SS analyzed the data and wrote the manuscript. Both authors were involved in manuscript discussions and commented on the manuscript.

