## Supplementary Figures for "Genome wide characterization and expression analysis of CrRLK1L gene family in wheat unravels their roles in development and stress-specific responses": Supplementary Figure S1. Nucleotide sequence analysis of Ta-CrRLK7-B copy.docx

```
T_dicoccoides_(WNHA010002628.1)      ATGCGGCGCCAACTCCTCCGTCTCCTTCCCGTCCGATTCCCCCGCCAACATCTTCGTCCC	60
T_aestivum_(CBTL0110686711.1)        ATGCGGCGCCAACTCCTCCGTCTCCTTCCCGTCCGATTCCCCCGCCAACATCTTCGTCCC	60
                                     ************************************************************

T_dicoccoides_(WNHA010002628.1)      CGACGCCGCCTACCTCTCGCCCGCGGGCGCTCCGGCGGTGTCGGCCAGCTCCACCCTGGC	120
T_aestivum_(CBTL0110686711.1)        CGACGCCGCCTACCTCTCGCCCGCGGGCGCTCCGGCGGTGTCGGCCAGCTCCACCCTGGC	120
                                     ************************************************************

T_dicoccoides_(WNHA010002628.1)      CTCCCCGCCAGCTCTGTACGCCGCCGCGCGCGCGGACATCTCGGCCTTCTCGTACCGCCT	180
T_aestivum_(CBTL0110686711.1)        CTCCCCGCCAGCTCTGTACGCCGCCGCGCGCGCGGACATCTCGGCCTTCTCGTACCGCCT	180
```

************************************************************

*****

```
T_dicoccoides_(WNHA010002628.1)      CCCTAGCCCCGCCTCGCCAGACGCGTCGTCATTCCTCGTCCTGCGCCTCCACTTCTTCCC	240
T_aestivum_(CBTL0110686711.1)        C-CTAGCCCCGCCTCGCCAGACGCGTCGTCATTCCTCGTCCTGCGCCTCCACTTCTTCCC	239
                                     * **********************************************************

T_dicoccoides_(WNHA010002628.1)      CTTCTTCCCCGCCACCTCCTCTCAGTATGTCATCAACATCTTGTCCGCGCGCTTCAACGT	300
T_aestivum_(CBTL0110686711.1)        CTTCTTCCCCGCCACCTCCTCTCAGTATGTCATCAACATCTTGTCCGCGCGCTTCAACGT	299
                                     ************************************************************
```

*****

```
T_dicoccoides_(WNHA010002628.1)      TTCGGTCGCCGACGCCTACGCTCTGCTGTCCTCCTTCTCGCCTCCGGCCGCCGGCGTCGT	360
T_aestivum_(CBTL0110686711.1)        TTAGGTCGCCGACGCCTACGCTCTGCTGTCCTCCTTCTCGCCTCCGGCCGCCGGCGTCGT	359
                                     ** *********************************************************

T_dicoccoides_(WNHA010002628.1)      CAAGGAGTTCTTCGTCCCGCGCGACCTGTTCGGTGACCACTTCCACGTCACGTTCACCCC	420
T_aestivum_(CBTL0110686711.1)        CAAGGAGTTCTTCGTCCCGCGCGACCTGTTCGGTGACCACTTCCACGTCACGTTCACCCC	419
                                     ************************************************************

T_dicoccoides_(WNHA010002628.1)      GGACGCCGGCTCCACCGCCTTCGTCAACGCCATCGAGCTGTTCTCGGCCCCGCCGGAGAT	480
T_aestivum_(CBTL0110686711.1)        GGACGCCGGCTCCACCGCCTTCGTCAACGCCATCGAGCTGTTCTCGGCCCCGCCGGAGAT	479
                                     ************************************************************

T_dicoccoides_(WNHA010002628.1)      GCTGTGGAATAGCTCCGTGACGCCCGTGGGAGCCGTGGTGAAGGACGACATGGACCTGTG	540
T_aestivum_(CBTL0110686711.1)        GCTGTGGAATAGCTCCGTGACGCCCGTGGGAGCCGTGGTGAAGGACGACATGGACCTGTG	539
                                     ************************************************************

T_dicoccoides_(WNHA010002628.1)      GCAGCGGCAGCCGCTGGAGACGGTCTATCGCCTCAACGTCGGAGGGCCCAAGGTGACCAT	600
T_aestivum_(CBTL0110686711.1)        GCAGCGGCAGCCGCTGGAGACGGTCTATCGCCTCAACGTCGGAGGGCCCAAGGTGACCAT	599
                                     ************************************************************

T_dicoccoides_(WNHA010002628.1)      TGAGAACGACACGCTGTGGCGGACGTGGCTGCCCGACGGTCCCTACCTCTACGACGCCTC	660
T_aestivum_(CBTL0110686711.1)        TGAGAACGACACGCTGTGGCGGACGTGGCTGCCCGACGGTCCCTACCTCTACGACGCCTC	659
                                     ************************************************************

T_dicoccoides_(WNHA010002628.1)      CGGGCTGTCGGTGGTGAGCAACACCTCCAACCCGATCATCTACGATTCATCGAACGGATA	720
T_aestivum_(CBTL0110686711.1)        CGGGCTGTCGGTGGTGAGCAACACCTCCAACCCGATCATCTACGATTCATCGAACGGATA	719
                                     ************************************************************

T_dicoccoides_(WNHA010002628.1)      CACGAGGGAGGTGGCGCCAGATGTCGTGTACCAGACCCAGCGCATGGCGAACGTGACGGA	780
T_aestivum_(CBTL0110686711.1)        CACGAGGGAGGTGGCGCCAGATGTCGTGTACCAGACCCAGCGCATGGCGAACGTGACGGA	779
                                     ************************************************************

T_dicoccoides_(WNHA010002628.1)      CTTACTGGCGGCGACAACCCCGGGCCTGAACTTCAACCTCACGTGGACGTTCCCGGCGGT	840
T_aestivum_(CBTL0110686711.1)        CTTACTGGCGGCGACAACCCCGGGCCTGAACTTCAACCTCACGTGGACGTTCCCGGCGGT	839
                                     ************************************************************

T_dicoccoides_(WNHA010002628.1)      GAAGGGGTCCCGCTACCTCGTCCGCCTCCACTTCTGCGACTACGAGGTGGTCAGCTCCGT	900
T_aestivum_(CBTL0110686711.1)        GAAGGGGTCCCACTACCTCGTCCGCCTCCACTTCTGCGACTACGAGGTGGTCAGCTCCGT	899
                                     *********** ************************************************

T_dicoccoides_(WNHA010002628.1)      CGTCGGCGTTGGCATCGTCTTCAACGTCTACATCGCGCAGACCATTGGCACTCCAGACCT	960
T_aestivum_(CBTL0110686711.1)        CGTCGGCGTTGGCATCGTCTTCAACGTCTACATCGCGCAGACCATTGGCACTCCAGACCT	959
                                     ************************************************************

T_dicoccoides_(WNHA010002628.1)      CACGCCGAATGCTCGGGCGACTCAGTCGAACGAGGTCTTTTACATGGACTACGCGGCCAG	1020
T_aestivum_(CBTL0110686711.1)        CACGCCGAATGCTCGGGCGACTCAGTCGAACGAGGTCTTTTACATGGACTACGCGGCCAG	1019
                                     ************************************************************

T_dicoccoides_(WNHA010002628.1)      GGCGCCGAGCACCGGGAACCTCACGGTGAGCATCGGCTGGTCGTCGAAAAGGAGCGGAGG	1080
T_aestivum_(CBTL0110686711.1)        GGCGCCGAGCACCGGGAACCTCACGGTGAGCATCGGCTGGTCGTCGAAAAGGAGCGGAGG	1079
                                     ************************************************************

T_dicoccoides_(WNHA010002628.1)      TGGGATACTGAACGGGCTAGAGATTATGAGGCTGCCGCCCGTTGATTTGAGCTCGAGGAG	1140
T_aestivum_(CBTL0110686711.1)        TGGGATACTGAACGGGCTAGAGATTATGAGGCTGCCGCCCGTTGATTTGAGCTCGAGGAG	1139
                                     ************************************************************

T_dicoccoides_(WNHA010002628.1)      GTACGGCAGGACGAAGAGGACCATTGTCATTACGGTGTCGGCAGTGCTCGGCGCCGCCGT	1200
T_aestivum_(CBTL0110686711.1)        GTACGGCAGGACGAAGAGGACCATTGTCATTACGGTGTCGGCAGTGCTCGGCGCCGCCGT	1199
                                     ************************************************************

T_dicoccoides_(WNHA010002628.1)      TCTTGCTTGCGTGGTGCTCTGCTTTTTCGGCGTGCCGTATACGAAGTACAGCGGCTCCGG	1260
T_aestivum_(CBTL0110686711.1)        TCTTGCTTGCGTGGTGCTCTGCTTTTTCGGCGTGCCGTATACGAAGTACAGCGGCTCCGG	1259
                                     ************************************************************

T_dicoccoides_(WNHA010002628.1)      CTGGGCTGAGCAGTTCACGAACCGATGGTCCAGAGAGGGCAAGACCAGCGGGTTGCAGAG	1320
T_aestivum_(CBTL0110686711.1)        CTGGGCTGAGCAGTTCACGAACCGATGGTCCAGAGAGGGCAAGACCAGCGGGTTGCAGAG	1319
                                     ************************************************************

T_dicoccoides_(WNHA010002628.1)      TGTGAGCACGAAGCTGCACATCGCTCTCGCGAAGATCAAGGCCGCCACGGACAACTTCCA	1380
T_aestivum_(CBTL0110686711.1)        TGTGAGCACGAAGCTGCACATCGCTCTCGCGAAGATCAAGGCCGCCACGGACAACTTCCA	1379
                                     ************************************************************

T_dicoccoides_(WNHA010002628.1)      CGAGCGCAACCTCATCGGCGTGGGCGGGTTCGGGAACGTGTACAAGGGCGTGCTCGTTGA	1440
T_aestivum_(CBTL0110686711.1)        CGAGCGCAACCTCATCGGCGTGGGCGGGTTCGGGAACGTGTACAAGGGCGTGCTCGTTGA	1439
                                     ************************************************************

T_dicoccoides_(WNHA010002628.1)      CGGCACGCCAGTGGCGGTGAAGCGCGCCATGCACGCCTCGCAGCAGGGGTTGCCGGAGTT	1500
T_aestivum_(CBTL0110686711.1)        CGGCACGCCAGTGGCGGTGAAGCGCGCCATGCGCGCCTCGCAGCAGGGGTTGCCGGAGTT	1499
                                     ******************************** ***************************

T_dicoccoides_(WNHA010002628.1)      CCAGACGGAGATCGTGGTGCTGTCCGGCATCCGGCACCGGCACCTGGTGTCGCTCATTGG	1560
T_aestivum_(CBTL0110686711.1)        CCAGACGGAGATCGTGGTGCTGTCCGGCATCCGGCACCGGCACCTGGTGTCGCTCATTGG	1559
                                     ************************************************************

T_dicoccoides_(WNHA010002628.1)      GTACTGCAACGAGCAGGCGGAGACGATACTGGTGTACGAATACATGGAGAAAGGCACGCT	1620
T_aestivum_(CBTL0110686711.1)        GTACTGCAACGAGCAGGCGGAGATGATACTGGTGTACGAATACATGGAGAAAGGCACGCT	1619
                                     *********************** ************************************

T_dicoccoides_(WNHA010002628.1)      GCGGAGCCACCTGTACGGTTCCGACGAGCCGGCGTTGTCATGGAAGCAGAGGCTGGAGAT	1680
T_aestivum_(CBTL0110686711.1)        GCGGAGCCACCTGTACGGTTCCGACGAGCCGGCGTTGTCATGGAAGCAGAGGCTGGAGAT	1679
                                     ************************************************************

T_dicoccoides_(WNHA010002628.1)      CTGCATCGGCGCGGCGAGGGGCCTGCACTACCTGCACAGAGGCTACGCGGAGAACATCAT	1740
T_aestivum_(CBTL0110686711.1)        CTGCATCGGCGCGGCGAGGGGCCTGCACTACCTGCACAGAGGCTACGCGGAGAACATCAT	1739
                                     ************************************************************

T_dicoccoides_(WNHA010002628.1)      CCACCGTGACGTCAAGTCGACCAACATCCTCCTCGGGAGCGACGGCGGCAGCACCGGTGG	1800
T_aestivum_(CBTL0110686711.1)        CCACCGTGACGTCAAGTCGACCAACATCCTCCTCGGGAGCGACGGCGGCAGCACCGGTGG	1799
                                     ************************************************************

T_dicoccoides_(WNHA010002628.1)      CGTGATCGCCAAGGTGGCCGACTTCGGGCTGTCGCGCATCGGGCCGTCGTTCGGGGAGAC	1860
T_aestivum_(CBTL0110686711.1)        CGTGATCGCCAAGGTGGCCGACTTCGGGCTGTCGCGCATCGGGCCGTCGTTCGGGGAGAC	1859
                                     ************************************************************

T_dicoccoides_(WNHA010002628.1)      GCACGTGAGCACGGCGGTGAAGGGCAGCTTCGGGTACCTGGACCCGGGGTACTTCAAGAC	1920
T_aestivum_(CBTL0110686711.1)        GCACGTGAGCACGGCGGTGAAGGGCAGCTTCGGGTACCTGGACCCGGGGTACTTCAAGAC	1919
                                     ************************************************************

T_dicoccoides_(WNHA010002628.1)      GCAGCAGCTGACGGACCGGTCGGACGTCTACTCCTTCGGCGTGGTGCTGTTGGAGGTGCT	1980
T_aestivum_(CBTL0110686711.1)        GCAGCAGCTGACGGACCGGTCGGACGTCTACTCCTTCGGCGTGGTGCTGTTGGAGGTGCT	1979
                                     ************************************************************

T_dicoccoides_(WNHA010002628.1)      CTGCGCGCGACCTGTGATCGACCAGAGCCTGGACCACGGCCGGATCAACATCGCCGAATG	2040
T_aestivum_(CBTL0110686711.1)        CTGCGCGCGACCTGTGATCGACCAGAGCCTGGACCACGGCCGGATCAACATCGCCGAATG	2039
                                     ************************************************************

T_dicoccoides_(WNHA010002628.1)      GGCCGTGAGGATGCGCAGGGAAGGGCGGCTCGACAAGATGGCCGACCCGAGGATCGCCGG	2100
T_aestivum_(CBTL0110686711.1)        GGCCGTGAGGATGCGCAGGGAAGGGCGGCTCGACAAGATGGCCGACCCGAGGATCGCCGG	2099
                                     ************************************************************

T_dicoccoides_(WNHA010002628.1)      CGAGGTGGACGAGGAGTCGCTGCTCAAGTTCGCAGAAACCGCTGAGAAGTGCCTGGCGGA	2160
T_aestivum_(CBTL0110686711.1)        CGAGGTGGACGAGGAGTCGCTGCTCAAGTTCGCAGAAACCGCTGAGAAGTGCCTGGCGGA	2159
                                     ************************************************************

T_dicoccoides_(WNHA010002628.1)      GTGCTGGGTGGACCGGCCGTCCATGGGCGACGTGCTGTGGAACCTGGAGTATTGCCTACA	2220
T_aestivum_(CBTL0110686711.1)        GTGCTGGGTGGACCGGCCGTCCATGGGCGACGTGCTGTGGAACCTGGAGTATTGCCTACA	2219
                                     ************************************************************

T_dicoccoides_(WNHA010002628.1)      GCTGCAGGAGACCAATATCACCGGGGACGGACTCGACGACATGGTACCGTCGTCGACGAG	2280
T_aestivum_(CBTL0110686711.1)        GCTGCAGGAGACCAATATCACCGGGGACGGACTCGACGACATGGTACCGTCGTCGACGAG	2279
                                     ************************************************************

T_dicoccoides_(WNHA010002628.1)      CTTGTTGATGGACGAGACCGACTTGAGCATGACCAATGTCGCCGACAGCAAGGTATTCTC	2340
T_aestivum_(CBTL0110686711.1)        CTTGTTGATGGACGAGACCGACTTGAGCATGACCAATGTCGCCGACAGCAAGGTATTCTC	2339
                                     ************************************************************

T_dicoccoides_(WNHA010002628.1)      CCAGCTGAGCGCCCGCGGCGAGGGACGATGATCTGAAGTTGGTGCACCAATTGAACATTT	2400
T_aestivum_(CBTL0110686711.1)        CCAGCTGAGCGCCCGCGGCGAGGGACGATGATCTGAAGTTGGTGCACCAATTGAACATTT	2399
                                     ************************************************************

T_dicoccoides_(WNHA010002628.1)      TCCTTTGGTGGTCCCTGAATCATCTGAACCCCACATGGATGGATGTTGCACAATACTCTG	2460
T_aestivum_(CBTL0110686711.1)        TCCTTTGGTGGTCCCTGAATCATCTGAACCCCACATGGATGGATGTTGCACAATACTCTG	2459
                                     ************************************************************

T_dicoccoides_(WNHA010002628.1)      CAGTTTTTTTTAG	2473
T_aestivum_(CBTL0110686711.1)        CAGTTTTTTTTAG	2472
                                     *************
```

Supplementary Figure S1. Multiple sequence alignment for whole genome shotgun (WGS) contig sequences for *T. aestivum* (WGS accession: CBTL0110686711.1) and *T. dicoccoides* (WNHA010002628.1) was carried out by using Clustal Omega (<https://www.ebi.ac.uk/Tools/msa/clustalo/>). The regions with nucleotide changes in *T. aestivum* are shown by black boxes and changes at 182 and 303 nucleotide positions are marked by red asterix*****.
