## Supplementary Figures for "Genome wide characterization and expression analysis of CrRLK1L gene family in wheat unravels their roles in development and stress-specific responses": Supplementary Figure S2. Phylogenetic analysis of CrRLK1Ls in wheat and other species.docx

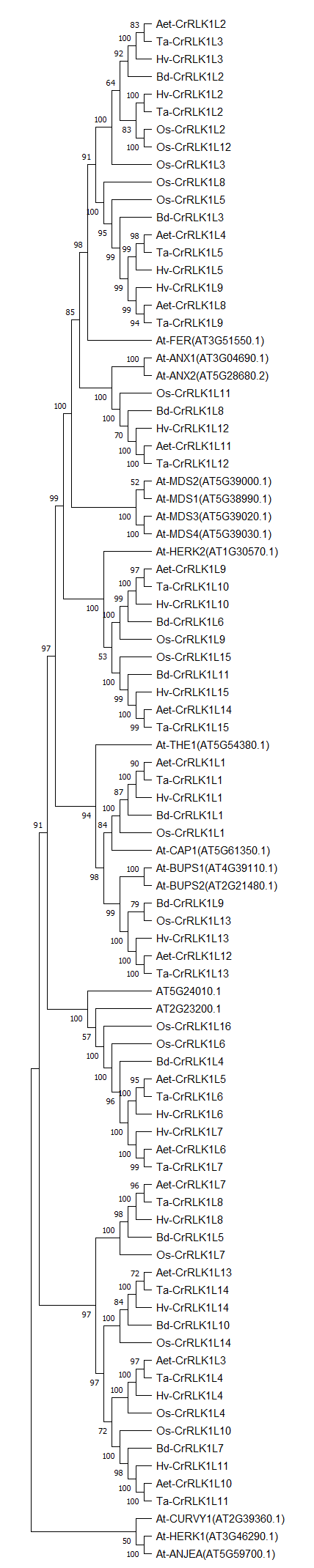

Supplementary Figure S2. Phylogenetic analysis of CrRLK1L gene families in *Triticum aestivum* and other species. The evolutionary history was inferred using the Neighbor-Joining method, JTT model and 1000 bootstraps. This analysis involved 88 amino acid sequences. All positions with less than 95% site coverage were eliminated.
