## Supplementary Tables for "Genome wide characterization and expression analysis of CrRLK1L gene family in wheat unravels their roles in development and stress-specific responses": Supplementary Table S2. Nucleotide and proetin Sequences for T. aestivum CrRLK1L gene families.docx

Table S2. Nucleotide and amino acid sequences for CrRLK1L gene families in *T. aestivum*. Sequences were collected from the Ensembl plants database and *-2-B* and *-2-D* copy was corrected from WGS contigs for *T. aestivum* at NCBI.

Nucleotide sequences:

>Ta-CrRLK1L1-A CDS sequence

ATGCCAGCATTGGCAATTTTGGCCAGAAGCATGGCTCCGTGCAAGAGAGTGCCGATGTTC

TTGATCCTCTTCATCCTTTCCATTACTCGTGTAGCCACCACCAATGCAATTGCATCAAAA

GTAGATCGGTTCGTGCCTCAAGACAACTACCTCCTCAGCTGCGGGGCATCCGCTGCTGTG

CAGGTTGACGATGGCAGGACATTCCGCTCTGATCCTGAGTCGGTGTCATTTCTGTCAACC

CTGACGGACATCAAGATCGCCGCTAAAGCATCCCTGGCTTCTGCTTCACCATTATCCCCA

CTTTACCTTGATGCAAGAGTATTCTCTGATATCTCAACCTATAGCTTCTTCATCTCCCAG

CCTGGTCGCCATTGGATCCGGCTCTACTTCTTACCTATCACCGATAGCCAATACAACCTC

ACCACTGCAACATTTTCCGTGTCCACTGATAGCATGGTCCTTCTCCATGATTTCTCCTTC

ATAGCCAGTCCTCCAAACCCTGTGTTTAGGGAGTATCTTGTGTCAGCACAGGGAGACAAC

TTAAAGATCATCTTTACCCCAAAGAAAAACTCGATAGCATTCATCAATGCTATCGAGGTT

GTCTCAGCCCCACCAAGCCTTATTCCAAATACCACCACCAGAATGGGTCCTCAGGACCAG

TTTGACATATCCAACAATGCATTGCAGGTTGTCTACCGGCTGAACATGGGCGGTGCACTG

GTTACATCGTTTAATGACACACTAGGCAGAACCTGGCAGCCAGATGCACCCTTTCTGAAG

CTTGAGGCAGCAGCAGAGGCAGCTTGGGTTCCTCCTAGAACCATCAAGTACCCTGATGAC

AAGACTCTCACACCCCTCATCGCTCCAGCAAGCATCTACTCAACAGCACAGCAGATGGCA

TCAACAAATATCACAAATGCAAGATTCAACATAACATGGCAAATGGTTGCAGAGCCAGGA

TTCAGGTACCTTATCCGCCTACATTTCAGCGACATTGTCAGCAAGACACTCAATAGCCTC

TACTTCAATGTCTATATTAATGGCATGATGGCTGTTGCCAACCTTGATCTATCGAGCCTG

ACGATGGGGCTTGCAGTAGCCTACTACAAGGACTTGATTGCTGAGTCTTCCAGCATCATC

AATTCCACCCTTCTAGTCCAGGTTGGCCCAAACACAATCGACTCCGGCGACCCCAATGCC

ATCCTTAATGGCCTTGAGATCATGAAGATAAGCAATGAAGCAAGCAGCTTAGATGGCCTT

TTTTCACCAAAAACAAGCTCAGAAGTTAGTAAGACAACACTGACTGGCATAGCATTTGCT

TTGGCAGCAACAGCTGCATTGGCTGTAGTTATATGCTACAGGCGAAACCGTAAACCGGAA

TGGCAGAGGACAAACAGCTTCCATTCTTGGTTTCTTCCACTGAACTCGTCCTCGAGCTTC

ATGAGCAGTTGCAGCAGGCTCTCCAGAAATCGCTTTGGCTCCACAAGGACCAAGAGTGGA

TTTTCGAGCGTGTTTGCATCCAGTGCTTATGGATTGGGACGCTATTTCACCTTCGTCGAA

ATTCAGAAAGCCACGAAAAACTTTGAAGAAAAGGGTGTTATTGGTGTTGGTGGCTTCGGA

AAAGTTTATCTTGGTGCTACTGAAGATGGCACACAGCTGGCTATCAAGCGAGGCAATCCA

TCATCTGATCAAGGTATGAATGAGTTTCTGACTGAAATTCAAATGTTATCTAAACTTCGC

CACCGCCACCTGGTTTCACTCATTGGCTGTTGTGATGAGAACAACGAGATGATATTAGTT

TATGAGTTCATGTCAAATGGTCCACTAAGGGATCATCTGTATGGTGACACAAACATCAAG

CCTATCTCTTGGAAGCAGCGCCTTGAAGTTTGCATTGGGGCAGCAAAGGGTCTGCATTAT

CTTCATACAGGTTCAGCTCAGGGCATAATTCACCGTGATGTCAAGACTACCAACATCCTA

CTTGATGAAAATTTTGTCGCCAAGGTTGCTGATTTTGGCCTATCAAAAGATGCTCCATCC

CTCGAACAAACTCATGTGAGCACTGCTGTCAAAGGAAGCTTTGGGTATCTTGATCCAGAG

TACTTCAGACGTCAACAGCTGACAGATAAGTCTGATGTATACTCTTTTGGTGTGGTACTC

TTTGAAGTGCTGTGTGCAAGGCCAGCCATCAATCCAGCCCTTCCAAGAGACCAAGTGAAT

CTGGCAGAGTGGGCCCGTACATGGCACCGCAAGGGGGAGCTTGGCAAAATAATTGATCCC

AATATTGCAGGACAAATCAGGTCTGATTCACTTGAGATGTTTGCTGAGGCTGCAGAGAAA

TGCCTTGCTGACTATGGAGTCGACCGGCCAACAATGGGAGACGTGCTATGGAAACTTGAA

TTTGCCTTGCAACTTCAAGAGAAGGGTGATGTCGTTGACGGCACCAGTGATGGGATCGCA

ATGAAGAGCTTGGAGGTGACCAATGTGGATAGCATGGAGAAATCTGGTAATGCTATCCCA

TCTTATGTCCAAGGAAGATGA

>Ta-CrRLK1L1-B CDS sequence

ATGCCAGCATTGGCAATTTTGGCCAGAAGCAGCCGCATGGCCGAGTGGGAGAGGGTACCA

ATGTTCTTGATCCTCTTCATCCTTTCCATTACTAGTGTAGCCACCACCAATGCAATTGCA

TCAAAAGTTGATCGGTTCGTGCCTCAAGACAACTACCTCCTCAGCTGTGGGGCATCCGCT

GCTGTGCAGGTTGACGATGGCAGGACATTCCGCTCTGATCCTGAGTCGGTATCGTTTCTG

TCAACCCCGACAGACATCAAGATTGCCGCTAAAGCATCTCTGGCTTCTGCTTCGCCATTA

TCTCCACTTTACCTTGACGCAAGAGTATTCTCTGATATCTCAACCTATAGCTTCTTCATC

TCCCAGCCTGGTCGCCACTGGATCCGTCTCTACTTCTTGCCTATCACCGACACCCAATAC

AACCTCACCACTGCAACATTTTCTGTGTCCACTGAGAGCATGGTTCTCCTCCATGATTTC

TCGTTCATAGCCAGTCCTCCAAACCCTGTGTTTAGGGAGTATCTTGTGTCAGCACAGGGA

GACAACTTAAAGATCATCTTTACTCCAAAGAAAAACTCGATAGCATTCATCAATGCGATC

GAGGTTGTCTCAGCCCCACCAAGCCTTATTCCAAATACCACCACCAGAATGGGTCCTCAG

GACCAGTTTGACATATCCAACAATGCATTGCAGGTTGTCTACCGGCTGAACATGGGCGGT

GCACTGGTTACGTCGTTCAATGACACACTAGGCAGAACCTGGCTGCCAGATGCACCCTTT

CTGAAGCTTGAGGCAGCAGCAGAGGCAGCTTGGGTTCCTCCTAGAACCATCAAGTACCCT

GATGACAAGACTCTCACACCCCTCATCGCTCCAGCAAGCATCTACTCAACAGCACAGCAG

ATGGCCTCAACAAATATCACAAATGCAAAATTCAACATAACCTGGGTAATGGTTGCAGAG

CCGGGATTCAGGTACCTTATCCGCCTACATTTCAGCGACATTGTCAGCAAGACACTCAAT

AGCCTCTACTTCAATGTCTATATTAATGGCATGATGGCTGTTGCCAACCTTGATCTATCG

AGCCTGACAATGGGGCTTGCAGTAGCCTACTACAAGGACTTGATTGCAGAATCTTCCAGC

ATCATCAATTCCACCCTTGTAGTCCAGGTTGGCCCAAGCACAATCGACTCCGGCGACCCC

AATGCCATCCTTAATGGCCTTGAGATCATGAAGATAAGCAATGAAGCAAGCAGCCTAGAT

GGCCTTTTTTCACCAAAAACAAGCTCAGAAGCTAGTAAGAGGACACTGACTGGCATAGCA

TTTGCTTTGGCAGCAACAGCTGCATTGGCTGTGGTTATATGCTACAGGCGAAACCGTAAA

CCGGCATGGCAGAGGACAAACAGCTTCCACTCTTGGTTTCTTCCACTAAACTCGTCCTCG

AGCTTCATGAGCAGCTGCAGCAGGCTCTCCAGAAATCGCTTTGGCTCCACAAGGACCAAG

AGTGGATTTTCGAGCGTGTTTGCATCCAGTGCTTATGGATTGGGGCGCTATTTCACCTTC

GTGGAAATTCAGAAAGCCACGAAAAACTTCGAAGAAAAGGGTGTTATTGGTGTTGGTGGC

TTCGGAAAAGTTTATCTTGGTGCTACTGAAGATGGCACACAGCTGGCAATCAAGCGAGGC

AATCCATCATCTGATCAAGGTATGAATGAGTTTCTGACTGAAATTCAAATGCTATCAAAA

CTTCGCCACCGCCACCTGGTTTCACTCATTGGCTGTTGTGATGAGAACAATGAGATGATA

TTAGTTTATGAGTTCATGTCAAATGGTCCGCTAAGGGATCATCTGTATGGTGACACAAAC

ATCAAGCCTATCTCTTGGAAGCAGCGCCTTGAAGTTTGCATTGGGGCAGCAAAGGGTCTG

CATTATCTTCATACAGGTTCAGCTCAGGGCATAATTCACCGTGATGTCAAGACTACCAAC

ATCCTACTTGATGAAAATTTTATCGCCAAGGTCGCTGATTTTGGCCTATCAAAAGATGCT

CCATCCCTCGAACAAACTCATGTGAGCACTGCTGTCAAAGGAAGCTTTGGGTATCTTGAC

CCAGAGTACTTCAGACGTCAACAGCTGACAGATAAGTCTGATGTATACTCTTTTGGTGTG

GTACTCTTTGAAGTGCTGTGTGCAAGACCGGCCATCAATCCATCTCTTCCAAGAGACCAA

GTGAATCTAGCTGAGTGGGCCCGTACATGGCACCGCAAGGGGGAGCTTGGTAAAATAATT

GATCCCAATATCGCAGGGCAAATCAGGCCTGATTCACTTGAGATGTTTGCTGAGGCTGCT

GAGAAATGCCTTGCTGACTATGGAGTCGACCGGCCAACAATGGGAGATGTGTTATGGAAA

CTTGAATTTGCCTTGCAACTTCAAGAGAAGGGTGATGTTGTTGATGGCGCCAGTGATGGG

ATCCCAATGAAGAGCTTGGAGGTGTCCAATGTGGATAGTATGGAGAAATCTGGTAATGCT

ATCCCATCTTATGTCCAAGGAAGATGA

>Ta-CrRLK1L1-D CDS sequence

ATGCCAGCATTGGCAATTTTGGCCAGAAGCATGGCTGAGTGCAAGAGAGTGCCCATGTTC

TTGATCCTCTTCATCCTTTCCATTACTAGTGTAGCCACCACCAATGCAATTGCATCAAAA

GTAGATCGGTTCGTGCCTCAAGACAACTACCTCCTAAGCTGCGGGGCATCAGCTGCTGTG

CAGGTTGACGATGGCAGGACATTCCGCTCTGATCCTGAGTCGGTATCGTTTCTGTCAACC

CCGACGGACATCAAGATCGCCGCTAAAGCATCCCTGGCTTCTGCTTCACCATTATCCCCA

CTTTACCTTGATGCAAGAGTATTCTCTGATATCTCAACCTATAGCTTCTTCATCTCCCAG

CCTGGTCGCCATTGGATCCGGCTCTACTTCTTACCTATCACCGATAGCCAATACAACCTC

ACCACGGCAACATTTTCCGTGTCCACTGATAGCATGGTCCTTCTCCATGATTTCTCCTTC

ATAGCCAGTCCTCCAAACCCTGTGTTTAGGGAATATCTTGTGTCAGCACAGGGAGACAAC

TTAAAGATCATCTTTACCCCAAAGAAAAACTCGATAGCATTCATCAATGCTATCGAGGTC

GTCTCAGCCCCACCAAGCCTTATTCCAAATACCACCACCAGAATGGGTCCTCAGGACCAG

TTTGACATATCTAACAGTGCATTGCAGGTTGTCTACCGGCTGAACATGGGTGGTGCACTG

GTTACATCGTTTAATGACACACTAGGCAGAACCTGGCAGCCAGATGCACCTTTTCTGAAG

CTTGAGGCAGCAGCAGAGGCAGCTTGGGTTCCTCCTAGAACCATCAAGTACCCTGATGAC

AAGACTCTCACACCCCTCATCGCTCCAGCAAGCATCTACTCGACAGCACAGCAGATGGCC

TCAACAAATATCACAAATGCAAGATTCAACATAACATGGCAAATGGCTGCAGAGCCGGGA

TTCAGGTACCTTATCCGCCTACATTTCAGCGACATTGTCAGCAAGACACTCAATAGCCTC

TACTTCAATGTCTATATTAATGGCATGATGGCTGTTGCCAACCTTGATCTATCGAGCCTG

ACAATGGGGCTTGCAGTAGCCTACTACAAGGACTTGATTGCTGAGTCTTCCAGCATCATC

AATTCCACCCTTGTAGTCCAGGTTGGCCCAAACACAATCGACTCCGGCGACCCCAATGCC

ATCCTTAATGGCCTTGAGATCATGAAGATAAGCAATGAAGCAAACAGCTTAGATGGCCTT

TTTTCACCAAAAACAAGCTCAGAAGTTAGTAAGACGACACTGACTGGCATAGCATTTGCT

TTGGCAGCAACAGCTGCATTGGCTGTAGTTATATGCTACAGGCGAAACCGTAAACCAGCA

TGGCAGAGGACAAACAGCTTCCATTCTTGGTTTCTTCCACTGAACTCGTCCTCAAGCTTC

ATGAGCAGTTGCAGCAGGCTCTCCAGAAATCGCTTTGGCTCCACAAGGACCAAGAGTGGA

TTTTCGAGCGTGTTTGCATCCAGTGCTTATGGATTGGGGCGCTATTTCACCTTCGTCGAA

ATTCAGAAAGCCACGAAAAACTTTGAAGAAAAGGGTGTTATTGGTGTTGGTGGCTTCGGA

AAAGTTTATCTTGGTGCTACTGAAGATGGCACACAGCTGGCTATCAAGCGAGGCAATCCA

TCATCTGATCAAGGTATGAATGAGTTTCTGACTGAAATTCAAATGTTATCAAAACTTCGC

CACCGCCACCTGGTTTCACTCATTGGCTGTTGTGATGAGAACAACGAGATGATATTAGTT

TATGAGTTCATGTCAAATGGTCCACTAAGGGATCATCTGTATGGTGACACAAACATCAAG

CCTATTTCTTGGAAGCAGCGCCTTGAAGTTTGCATTGGGGCAGCAAAGGGTCTGCATTAT

CTTCATACAGGTTCAGCTCAGGGCATAATTCACCGTGATGTCAAGACTACCAACATCCTA

CTTGATGAAAATTTTGTCGCCAAGGTCGCTGATTTTGGCCTATCAAAAGATGCTCCATCC

CTCGAACAAACTCATGTGAGCACTGCTGTCAAAGGAAGCTTTGGGTATCTTGATCCAGAG

TACTTCAGACGTCAACAGCTGACAGATAAGTCTGATGTATACTCTTTTGGTGTGGTACTC

TTTGAAGTGCTGTGTGCAAGACCAGCCATCAATCCAGCCCTTCCAAGAGACCAAGTGAAT

CTGGGAGAGTGGGCCCGTACATGGCACCGCAAGGGGGAGCTTGGTAAAATAATTGATCCC

AATATAGCAGGACAGATCAGGCCTGATTCACTTGAGATGTTTGCTGAGGCTGCTGAGAAA

TGCCTTGCTGACTATGGAGTTGACCGGCCAACAATGGGAGACGTGCTATGGAAACTTGAA

TTTGCCTTGCAACTTCAAGAGAAGGGTGATGTTGTTGACGGCGCCAGTGATGGGATCGCA

ATGAAGAGCTTGGAGGTGACCAATGTGGATAGCATGGAGAAATCTGGTAATGCTATCCCA

TCTTATGTACAAGGAAGATGA

>Ta-CrRLK1L2-A CDS sequence

ATGGTGCTCCCAACCTTACCGGTTACCCTCACATTCCTCACACTGTTAGCCCTCTTGTCG

ATTGCCAAGGCGGCTGATAACAACTCCACAACCTCTGGCCTCATCCTCCTAAACTGCGGA

GAATCTACCCAAGACGATGATGATGGTGGTCGTTCTTGGGATGGGGACACCGGCTCCATA

TTCGCGCCATCAATGAAAGGAGATGCAGCCATTGCTTTAGGTCAACCCCCTTCACTCACC

CCCAGGGTTCCTTATACAACTGCACGCATCTTTACTTCAAATTACACCTATTCCTTCCCT

GTCAGTCCAGGCCGAATGTTCTTACGCCTATACTTCCTTTCAACTGCTTACGAATACTAT

GCTGTCTCAGATGCCGTCTTCGGAGTCACGGCACGGAATCTTGTCCTCTTAAAAGACTTC

AATGCTTTGCAAACAGCTCAGGCGATCACTTCTGCCTACCTTGTGCGTGAATTCTCGGTG

AATGTTTCTTCAGGCAGCTTGGACCTCACCTTTGCACCATCAGCACATCAGTATGGGTCT

TATGCATTTGTGAATGGCATTGAGATTGTGCCCACGCCTGACATCTTCGCAACACCTGAC

ATAAGATTTGTCAGCGGTGATAACACATCTCCATTCACATTCGACGCTGACATGAGCCTC

CAGACTATGTACCGGCTCAATGTTGGGGGCCCAGCCATTTCCCCGAAAGGTGACTCGGGC

TTTTACCGCTCATGGGCCAATGATGCCCCATACATACTTGGTGGCTTTGGGTTGACCTTT

TGGAAAAATGATAATTTGACTATCAGTTATACATCTAGAGTGCCGAATTACACCGCCCCA

GTTGATGTCTATGGTACAGCTCGGTCCATGGGGCCAACTGCACAGATCAACCTGAACTAC

AACCTTACATGGATTTTACCGGTTGATGCGGGTTTCTTTTACCTCCTAAGGTTTCATTTC

TGTGAGATTAAGTATCCTATTACCAAGGTGAATCAGAGGTCGTTCTTCATCTACATCAAC

AACCAGACAGCGCAGGAGCAAATGGATGTCATCTTCCGGAGCGGAGGAATCGGTAGACCA

ACATACACTGAATATGTTATCATGGCTATTGGTTCTGGTCAGGTGGACATGTGGATTGCA

CTTCACCCTGATCTTTCAAGTAAACCACAGTATTCTGATGCAATACTGAATGGTCTCGAG

GTCTTCAAGCTACAGAATTACGGACCGAGCAATCTTGCTGGGCTCAATCCTCCACTTCCG

CAAAAGCCTGATGTGAATCCTAATAGGCTATCTAGCGGTGAAAGAAAAACCAAAGGTGGC

ATACAAGCAACCATCGGTGGTACTGCTGGTGGTTTTGCTTTATTGTTGATTGCCCTTTTC

AGCATGTGTGTTATCTACAGACGGAAGAAGGCAGCGAAGAGTCCCGGCAAGACCGACTAT

GGACATGTGAAGCATCCAACTAAATGCATAAAGTCTACATGTGATCTTGTACGTCATTTC

TCATTTGCTAAAATTCAAGTTGCCACCAAAGACTTTGATGAAGCACTTATCATCGGCAGA

GGCGGTTTCGGGAATGTCTACATCGGCGATATAGATGGAGGGACAAAGGTGGCGATCAAG

CGATGTGACCAGAAATCCCAACAAGGCTTTCATGAGTTCCAGACTGAAATCGAGATGCTG

TGCAATTTCCGCCATCGCCACCTTGTGTCTCTGATTGGCTATTGTGAGGAGAAGAATGAG

ATGATTCTGGTGTATGACTACATGGCTCATGGAACACTGCGTGAGCATTTGTACAACACC

AGGAACCCACCACTGCCGTGGCAGCAGCGCCTTGAGATTTGCATCGGTGCAGCCCGAGGA

CTGCATTACCTCCACACCGGCGTAGAGCAAGGAATCATCCACCGTGACGTCAAGACCACC

AACATCCTACTGGATGATAGGTTAATGGCAAAGGTTTCCGACTTCGGTTTGTTTAAGGCT

AGTCCAGACATTGGCAACACCCACATGAGCACTGCTGTTAAGGGCACCTTTGGATATCTT

GATCTGGAGTACTTCCGGCAGCAGCGTCTCACCAAAAAATCAGATGTGTACTCCTTCGGG

GTTGTGTTGTTTGAGACCCTGTGTGCGCGCCCTGTGATAAATACTGAGCTCCCTTATGAG

CAAGTGAGCTTGCGTGACTGGGTGGTATCTTGCCGAAAGAAAGGTGTACTCGAGGAGATT

GTTGACCCCTGTGTTAAGGAGGAAATCACCCTTGAGTGCTTCAGGATATTTGCAGAGATA

GCGGAGAAATGCGTTGCTGATCGTAGCATAGATAGGCCATCAATGGGTGATGTACTTTGG

AACCTTGAGGTCGCACTCCAGCTTCAGGATAGTGCAAGCTACAACACCAGCTGTGCTGAG

GGTGCATCATCTCTTCAGATCAGCGGAGTGCATTCAGGCAAACCATCAACCAACTCAACA

ATTAGCGTTGCAGCACAGGAAGCCATATTTTCAGATATTGCACATCCAGAAGGCCGATAA

>Ta-CrRLK1L2-B CDS sequence

ATGGTGCTCCCAACCTTACCGGTTACCCTCACATTCCTCACACTGCTAGCTCTCTTGTCG

ATTGCCAAGGCGGCTGATAACAACTCCACAACCTCTGGCCTTATCCTCCTAAATTGCGGA

TCATCAACCCAAAACGATGATGATAGTGGTCGTACTTGGGATGGGGACACCGGCTCCAAA

TTCGCGCCATCAATGAAAGGAGTTGCAGCCATTGCTTTAGGCCAAACCCCTTCACTCACC

CCCAGGGTTCCTTATACAACTGCACGCATCTTTACTTCAAATTACACCTATTCCTTCCCT

GTCAGTCCAGGCCGAATGTTCTTACGCCTATACTTCTTTTCAACTGCTTACGAATACTAT

GCTGTCTCAGATGCCGTCTTCGGAGTCACGTCACGGAATCTTGTCCTCTTAAATGACTTC

AATGCTTTGCAAACAGCTCAGGCGATCACTTCTGCCTACCTTGTGCGTGAATTCTCGGTG

AATGTTTCTTCAGGCAGCTTGGACCTCACCTTTGCACCATCAGCACAACAGTATGGGTCT

TATGCATTTGTGAATGGCATTGAGATTGTGCCCACGCCTGACATCTTCGCAACACCTGAC

ATAAGATTAGTCAGCGGTGATAACACATCTCCATTCACATTCGATGCTGACATGAGCCTC

CAGACTATGTACCGGCTCAATGTCGGGGGCCCAGCCATTTCCACGGAAGGTGACTCGGGC

TTTTACCGCTCATGGGCCAATGATGCCCAATACATACTTGGTGGCTCTGGGTTGACCTTT

TGGAAAAATGATAATTTGACTATCAGTTATACATCTAGAGTGCCGAATTACACCGCCCCA

GTTGATGTCTATGGTACAGCTCGGTCGATGGGGCCAACTGCACAGATCAACCTGAACTAC

AACCTTACATGGATTTTTCCGGTTGATGCAGGTTTCTTTTACCTCCTAAGGTTCCATTTC

TGTGAGATTAAGTATCCTATTACCAAGGTGAATCAGAGGTCATTCTTCATCTACATCAAC

AACCAGACAACGCAGAAGCAAATGGATGTCATCGTCCGGAGCGGAGGAATCGGTAGACCA

ACGTACACTGAATATGTTATCATGGCTATTGGTTCTCGTCAGGTGGACATGTGGATTGCA

CTTCACCCTGATCTTTCAAGTAAACCACAGTATTCGGATGCAATTCTGAATGGTCTCGAG

GTCTTCAAGCTACAGAATTACGGACCGAGCAATCTTGCTGGGCTCAGTCCTCCACTTCCG

CAAAAGCCTGATGTGAATCCTACTAGGCTATCTAATGGTGAAAGAAAATCCAAAGGTGGC

ATACAAGCAATCATCGGTGGTACTACTGGTGGTTTTGCTTTATTGTTGATTGCCCTTTTC

AGCATGTGTGTTATATACAGACGGAAGAAGGTAGCGAAGAGTCCCGGCAAGACCGACTAT

GGACATGTGAAGCATCCAACTAAATGCATAAAGTCTACATGTGATCTTGTACGTCATTTC

TCATTTGCTAAAATTCAAGTTGCCACCAAAGACTTTGATGAAGCACTTATTATCGGCAGA

GGCGGTTTCGGGAATGTCTACATCGGCGATATAGATGGAGGGACAAAGGTGGCAATCAAG

CGATGTGACCAGAAATCCCAACAAGGCTTTCATGAGTTCCAGACTGAAATCGAGATGCTG

TGCAATTTCCGCCATCGCCACCTTGTGTCTCTGATTGGCTATTGTGAGGAGAAGAATGAG

ATGATTCTGGTGTATGACTACATGGCTCATGGAACACTGCGTGAGCATCTGTACAACACC

AGGAACCCACCACTACCGTGGCAGCAGCGCCTTGAGATTTGCATCGGTGCAGCCCAAGGA

CTGCATTACCTCCACACCGGCGTAGAGCAAGGAATCATCCACCGTGACGTCAAGACCACC

AACATCCTACTGGATGATAGGTTAATGGCAAAGGTTTCCGACTTCGGTCTGTCTAAGGCT

AGTCCAGACATTGGAAACACCCACATGAGCACTGCTGTGAAGGGCACCTTTGGATATCTT

GATCCGGAGTACTTCCGGCTGCAGCGTCTCACCAAAAAATCAGATGTGTACTCTTTCGGG

GTCGTGTTGTTTGAGACCCTGTGTGCGCGCCCTGTGATAAACACTGAGCTCCCTTATGAG

CAAGTGAGCTTGCGTGACTGGGCGCTATCTTGCTGGAAGAACGGTGTACTTGAGGAGATT

GTTGACCCCCGTGTTAAGGAGGAAATCACCCCTGAGTGCTTCAGGGTTTTTGCAGAGATA

GCAGAGAAATGTGTAGCTGATCGTAGCATAGAGAGGCCATCAATGGGTGATGTACTTTGG

AACCTTGAGGTCGCACTCCAGCTGCAGCAGGCTAGTGCAAGCTACAACAGCAACCGTGCA

GAGGGTGCTTCATCTCTTCAGATCAGCGCGGTGCATTCAGACAAACCATCCACCAACTCA

ACAATTAGCATCGCAGCACAGGAAGCCATATTTTCAGATATTGCGCATGCAGAAGGCCGA

TAA

>Ta-CrRLK1L3-A CDS sequence

ATGCAAACATCCTGTAGCAAGTTGATCCGATGGAGTCCCCAATTCTTTGATTCCGGTGCA

CCAACCGCCGCAAATTCCAAAATGGCTTTTCCAGCCCTACCAGCTACCCTCACATGCCTC

ACACTGTTAGCTCTCTTGTCGCTGGCCATGGCGGCTGATAACAACTCCACCGGCCTCATC

CTCATAAACTGCGGAGCATCAGTCCAAGAAGACGATGATAATGGTCGTACTTGGGACGGA

GACACCGGCTCCAAGTTCGCGCCATCACTGAAAGGAGTTACAGCCACTGCTCCAAACCAA

GACCCTTCACTCCCCTCCACCGTCCCTTTTATGACCGCGCGCATCTTCGCTTCAAACTAC

ACCTATTCCTTCTCTGTCACCCCAGGCCGCGTGTTCTTACGCCTCTACTTCTATCCGGTT

GCTTATCCAAACTACGCCGTCGCAGATGCCTTCTTCAGTGTCACGACACCGAATCTTGTC

CTCTTAAATGATTTCAATGCTTCGCAAACAGTTCAGGCGATCAGTTCTGCCTACCTTGTG

CGCGAGTTCTCGGTGAATGTTTCTTCAGGCAGCTCCTTGGACCTCACCTTTGCCCCATCA

GCACATCACAATGGTTCTTACGCGTTTGTGAACGGCATTGAGATTGTTTCCACTCCTGAC

ATCTTCACAGCACCTGACACAAGATATGTCGGTGATAACACATCTCCATTCACATTCGAC

TCTGCCATGGCCGTCCAGACTATGTACCGGCTCAATGTCGGGGGCCAAGCCATTTCCCCG

AAAGGTGACTCGGGCTTCTACCGCTCATGGGCCAATGATGCCCCCTACATATTTGGTGGC

TCTGGGGTGACCTTCTCCAAGGATGACAATTTGACTATCACCTATACATCCAAAGTGCCG

AATTACACGGCGCCAGTTGATGTCTATGGTACAGCTCGGTCGATGGGGCCAACTGCACCG

ATCAACCTGAACTACAACCTTACATGGATTTTACCGGTTGATGCGGGGTTCAGTTACCTC

CTGAGGTTCCATTTCTGTGAGATTCAGTATCCTATTACAAAGCAGAATCAGAGGTCCTTC

TTCATCTACATCAACAACCAGACAGCGCAGGAGCAAATGGATGTCATCGTCTGGAGCGGC

GGAATCGGTAGAACAACATACACAGACTATGTTATCCTGACTGCTGGCTCCGGCCAGGTG

GACATGTGGATTGCACTTCACCCTGATCTTTCAAGTAGACCAGAGTATTTTGATGCAATA

CTGAATGGTCTTGAGATCTTCAAGCTACAGAATTACGGAGCATCGAACAATCTTGCCGGG

CTCAATCCTCCACTTCCACAAAAGCCAGCTGATGCCAGTCCCGGCGCGGCATCTGGCAAA

GTGAAATCTGTCGCGGCTATCATAGGTGGAGCTGTTGGTGGTTTCGTAGTGCTGCTGGTC

ACATGTTTTGGCATTTGCATCATCTGCAAACGAAAGAACAAGAGCAAGAAGAAGAAGAAG

ATATCCAAGGATCCTGGTGGTAAATCTGAAGATGGTCACTGGACTCCTCTCACCGAGTAC

AGCGGATCACGATCAGCTATGTCGGGAAACACGGCCACCACCGGGTCGACACTGCCATCC

AACCTCTGCCGCCACTTCACTTTCGCCGAGCTTCAGACCGCCACCAAGAACTTCGACCAG

GCCTTCCTGCTCGGCAAAGGTGGGTTCGGGAACGTGTACCTCGGGGAGATCGACAGCGGC

ACCAAGGTGGCGATCAAGCGGTGCAACCCGATGTCGGAGCAGGGCGTCCATGAGTTCCAG

ACGGAGATCGAGATGCTGTCCAAGCTCCGGCACCGCCACCTCGTGTCCCTCATCGGCTAC

TGCGAGGACAAGAGCGAGATGATCCTGGTGTACGACTACATGGCCCACGGCACGCTCCGG

GAGCACCTGTACAACACCAAGAACCCGCCGCTGTCGTGGAAGCAGCGGCTGGAGATCTGC

ATCGGCGCCGCCCGGGGGCTCTACTACCTGCACACCGGCGTGAAGCACACCATCATCCAC

CGCGACGTCAAGACCACCAACATCCTGCTGGACGACAAGTGGGTCGCCAAGGTGTCCGAC

TTCGGGCTGTCCAAGACGGGGCCCAACATGGACGCCACCCACGTCAGCACCGTCGTCAAG

GGCAGCTTCGGGTACCTGGACCCGGAGTACTTCCGGCGGCAGCAGCTCTCGGAGAAGTCC

GACGTCTACTCCTTCGGCGTCGTGTTGTTCGAGGTGCTCTGCGCGCGCCCGGCGCTGAGC

CCCTCGCTGCCCAAGGAGCAGATCAGCCTCGCCGACTGGGCGCTGCGCTGCCAGAAGCAG

GGCGTGCTCGGCCAGGTCATCGACCCGGTGCTCCAGGGGAAGATCGCGCCCCAGTGCTTC

CTCAAGTTCACGGACACCGCGGAGAAATGCGTGGCCGACCGCAGCGTCGACAGGCCGTCC

ATGGGCGACGTCCTCTGGAACCTCGAGTTCGCGCTCCAGCTGCAGGAGAGCGAGGAGGAC

ACCGGCAGCCTCACGGAGGGGACGCTGTCGTCGTCGGGCGCGTCGCCCCTCGTCATGACC

AGACTGCAGTCGGACGAGCCGTCGATGGACGCAAGCACGACCACGACGAGCACGACCACG

ATGAGCATGACGGGACGGAGCATCGCGAGCATGGACTCGGACGGGCTGACGCCGAGCGCC

GTCTTCTCGCAGATCATGCATCCGGATGGCAGGTGA

>Ta-CrRLK1L3-B CDS sequence

ATGCAAACATCCTGTAGCAAGTTGATCCGATGGAGTCCCCAATTCTTTGATTCTGGTACA

CCAACCACCGCAAAATCCAAAATGGCTTTCCCAGCCCTACCAGTTACCCTCACATGCCTC

ACACTGTTAGCTCTCTTATCACTCGCCATGGCGGCTGATAACAACTCCACTGGCCTCATC

CTCGTAAATTGCGGAGCATCGACCCAAGAAGCTGATGATAGTGGTCGTACTTGGGTCGGG

GACACCGGCTCCAAGTTCGCGCCATTATTGAAAGGAGTTGCAACCACTGCTCCAAACCAA

GACCCTTCACTCCCCTCCACGGTCCCTTTTATGACTGCACGCATCTTCACTTCAAACTAC

ACCTATTCCTTCTCTGTCAACCCAGGCCGCATGTTCTTACGCCTCTACTTCTATCCGGTT

GCTTATGCAAACTATGCCGTCTCAGATGCCTTCTTCAGTGTCACGACACGGAATCTTGTC

CTCTTAAATGATTTCAGTGCTTCGCAAACAGCTCAGGCGATCACTTCTGCCTTCCTTGTG

CGCGAGTTCTCGGTGAATGTTTCTTCAGGATCCTCCTTGGACCTCACCTTTGCCCCATCA

GCACATCGCAATGGTTCTTACGCATTTGTGAACGGCATTGAGATTGTGCCCACTCCTGAC

ATCTTCACAGCACCTGACACAAGATATGTCGGTGATAACACAGCCCCATTCTCATTCGAC

GCTGGCATGGCCGTCCAGACTATGTACCGGCTCAATGTCGGGGGCCAAGCCATTTCCCCG

AAAGGTGACTCGGGCTTCTACCGCTCATGGGCCAATGATGCCCCCTACATATTTGGTGGC

TCTGGGGTGACCTTCTCCAAAGATGATAATTTGACCATCACCTATACATCCAACGTGCCG

AATTACACGGCGCCAGTTGATGTCTATGGTACAGCTCGGTCGATGGGGCCAACTGCACAG

ATCAACCTCAACTACAACCTTACATGGATTTTACCCGTTGATGCGGGGTTCAGTTACCTC

CTAAGGTTCCATTTCTGTGAGATTCAGTATCCTATTACAAAGCAGAATCAGCGGTCCTTC

TTCATCTACATCAACAATCAGACAGCGCAGGAGCAAATGGATGTCATCGTCTGGAGCGGA

GGAATCGGTAGAACAGCATACACAGACTATGTTATCATGGCTGTTGGTTCTGGTCAGGTG

GACATGTGGATTGCACTTCACCCTGATCTTTCAAGTAAACCAGAGTATTTTGATGCAATA

CTGAATGGTCTTGAGATCTTCAAGCTACAGAATTACGGATCACCGAACAATCTTTCTGGG

CTCAATCCTCCACTTCCACAAAAGCCAACTGATGCCAGTCCCGGCGCGGCGTCTGGCAAA

ATGAAATCTGTCGCGGCTATCATAGGTGGAGCTGTTGGTGGTTTCGCAGTGCTGCTGGTC

ACATGTTTTGGCGTTTGCATCATCTGCAAACGAAAGAACAAGAAGAACAAGAAGAAGATA

TCCAAGGATCCTGGTGGTAAATCTGAAGATGGTCACTGGACTCCTCTCACCGAGTATAGC

GGATCACGGTCAGCTATGTCGGGAAACACGGCCACAACCGGGTCGACGCTGCCATCCAAC

CTCTGCCGCCACTTCACTTTCGCCGAGCTTCAGACCGCCACCAAGAATTTCGACCAGGCC

TTCCTGCTCGGCAAAGGTGGGTTCGGGAACGTGTACCTCGGGGAGATCGACAGTGGCACC

AAGGTGGCGATCAAGCGGTGCAACCCGATGTCCGAGCAGGGCGTCCACGAGTTCCAGACG

GAGATCGAGATGCTGTCCAAGCTCCGGCACCGCCACCTGGTGTCCCTCATCGGCTACTGC

GAGGACAAGAGCGAGATGATCCTGGTGTACGACTACATGGCCCACGGCACGCTCCGGGAG

CACCTCTACAACACCAAGAACCCGCCGCTGTCGTGGAAGAAGCGGCTCGAGATCTGCATC

GGCGCCGCCCGGGGCCTCTACTACCTGCACACCGGCGTGAAGCACACCATCATCCACCGC

GACGTCAAGACCACCAACATCCTGCTGGACGACAAGTGGGTCGCCAAGGTCTCCGACTTC

GGGCTGTCCAAGACGGGGCCCAACATGGACGCCACCCACGTCAGCACCGTCGTCAAGGGC

AGCTTCGGGTACCTCGACCCGGAGTACTTCCGGCGGCAGCAGCTCTCGGAGAAGTCCGAC

GTCTACTCCTTCGGCGTCGTGCTGTTCGAGGTGCTCTGCGCGCGCCCCGCGCTGAGCCCC

TCGCTGCCCAAGGAGCAGATCAGCCTCGCCGACTGGGCGCTGCGCTGCCAGAAGCAAGGC

GTGCTCGGCCAGGTCATGGACCCGGTGCTCCAGGGGAAGATCGCGCCCCAGTGCTTCCTC

AAGTTCACGGACACCGCGGAGAAATGCGTGGCCGACCGCAGCGTCGACCGGCCGTCCATG

GGCGACGTCCTCTGGAACCTCGAGTTCGCGCTCCAGCTGCAGGAGAGCGAGGAGGACACC

GGCAGCCTCACTGAGGGGACACTGTCGTCGTCGGGCGCGTCGCCTCTCGTCATGACCAGA

CTGCAGTCGGACGAGCCGTCGACGGACGCAAGCACCACCACGACGACCACGACCACGATG

AGCATGACGGGACGGAGCATCGCGAGCGTGGACTCGGACGGGCTGACGCCGAGCGCCGTT

TTCTCGCAGATCATGCATCCGGATGGCAGGTGA

>Ta-CrRLK1L3-D CDS sequence

ATGGCGTTCCCAGCCCTACCAGTTACCCTCACATGCCTCATACTGTTATCTCTCTTGTCG

CTTGCCATGGCGGCTGATAACAACTCCACTGGCCTCATCCTCGTAAATTGCGGTGCATCA

GTGCAAGGCGACGATGATAGTGGCCGTACTTGGGACGGGGACACCGGCTCCAAGTTCGCG

CCATCATTGAAAGGAGTTGCAGCCACTGCTCCAAACCAAGACCCTTCGCTCCCCTCCACG

GTCCCTTTTATGACCGCACGCATCTTCACTTCAAACTACACATATTCCTTCTCTGTCAAA

CCAGGCCGCATGTTCTTGCGCCTCTACTTCTATCCGGTTGCTTATCCAAACTATGCCGTC

TCAGATGCCTTCTTCAGTGTCACGACGCCGAAACTTGTCCTCTTAAATGACTTCAGTGCT

TCGCAAACAGCTCAGGCGATCACTTCTGCCTTCCTTGTGCGTGAGTTCTCGGTGAATGTT

TCTTCAGGATCCTCCTTGGACCTCACCTTCGCCCCATCTGCACATCGCAATGGTTCTTAC

GCATTTGTGAACGGCATTGAGATTGTGCCCACTCCTGACATCTTCACAGCACCTGACACA

AGAAATGTCGGTGATAACACAGCCCCATTCTCATTCGACACTAGCTCGAGCCTCCAGACT

ATGTACCGGCTCAATGTCGGGGGCCAAGCCATTTCCCCGAAAGGTGACTTGGGGGGCTTC

TACCGCTCATGGGCGAATGACGCCCCGTACATAGCTGGTGGCTCTGGGGTGACCTTCTCC

AAAGATGATAATTTGACCATCACTTATACATCCAAAGTGCCGAAGTACACGGCGCCACCT

GATGTCTATGGTACAGCTCGGTCGATGGGGCCAACTGCACAGATCAACCTCAACTACAAC

CTTACATGGATTTTACCGGTTGATGCGGGGTTCTTTTACCTCCTAAGGTTCCATTTCTGT

GAGATTCAGTATCCTATTATCAAGATCAATCAGAGGTCCTTCTTCATCTACATCAACAAC

CAGACAGCTCAGGAGCAAATGGATGTCATCGTCTGGAGCGGAGGAATCGGTAGAACAACA

TACACGGACTATGTTATCATGGCTGCTGGTTTCGGTCAGGTGGACATGTGGATTGCACTC

CACCCTGATCTTTCAAGCAGACCAGAGTATTTTGATGCAATACTGAATGGTCTGGAGGTC

TTCAAGCTACAGAATTACGGATCACCGAACAATCTTTCTGGGCTCAATCCTCCACTTCCA

CAAAAGCCAGCTGATGCCAGTCCCAGCGCGGCATCTGGCAAAATGAAATCGGTCGCGGCC

ATCATAGGTGGAGCTGTTGGTGGTTTCATAGTGCTACTGGCCGCGTGTTTTGGCGTTTGC

ATCATCTGCAAACGAAAGAACAAGAAGAAGAAGAAGAAGAAGACATCCAAGGATCCTGGT

GGTAAATCTGAAGATGGTCACTGGACTCCTCTCACCGAGTACAGCGGATCACGATCAGCC

ATGTCGGGAAACACGGCCACCACTGGGTCGACACTGCCATCCAATCTCTGCCGCCACTTC

ACTTTCGCCGACCTTCAGACCGCCACCAAGAACTTCGACCAAGCCTTCCTGCTCGGCAAA

GGTGGGTTCGGGAACGTGTACCTCGGGGAGATCGACAGCGGCACCAAGGTGGCGATCAAG

CGGTGCAACCCGATGTCGGAGCAGGGCGTCCATGAGTTCCAGACGGAGATCGAGATGCTG

TCCAAGCTCCGGCACCGGCACCTCGTGTCCCTCATCGGCTACTGCGAGGACAAGAGCGAG

ATGATCCTGGTGTACGACTACATGGCCCACGGCACGCTCCGGGAGCACCTCTACAACACC

AAGAACCCGCCGCTGTCGTGGAAGCAGCGGCTGGAGATCTGCATCGGCGCCGCCCGGGGC

CTCTACTACCTGCACACCGGCGTGAAGCACACCATCATCCACCGCGACGTCAAGACCACC

AACATCCTGCTGGACGACAAGTGGGTCGCCAAGGTCTCCGACTTCGGGCTGTCCAAGACG

GGCCCCAACATGGACGCCACCCACGTCAGCACCGTCGTCAAGGGCAGCTTCGGGTACCTC

GACCCGGAGTACTTCCGCCGGCAGCAGCTCTCGGAGAAGTCCGACGTCTACTCCTTCGGC

GTCGTGCTCTTCGAGGTGCTCTGCGCGCGCCCCGCGCTGAGCCCCACGCTGCCCAAGGAG

CAGATTAGCCTGGCCGACTGGGCGCTGCGCTGCCAGAAGCAGGGCGTGCTCGGCCAGGTC

ATCGACCCGGTGCTCCAGGGGAAGATCGCGCCCCAGTGCTTCCTCAAGTTCACGGACACC

GCGGAGAAATGCGTGGCCGACCGCAGCGTCGACAGGCCGTCCATGGGCGACGTCCTCTGG

AACCTCGAGTTCGCGCTCCAGCTGCAGGAGAGCGAGGAGGACACCGGCAGCCTCACGGAG

GGGACGCGGTCGTCGTCCGGCGCGTCGCCCCTCGTCATGACCAGGCTGCAGTCGGACGAG

CCGTCGACGGACGCAAGCACCACCACGACGTCCACGACCACGATGAGCATGACGGGGCGG

AGCATCGCGAGCATGGACTCGGACGGGCTGACCCCGAGCGCCGTTTTCTCGCAGATCATG

CATCCGGATGGCAGGTGA

>Ta-CrRLK1L4-A CDS sequence

ATGGCCGCGACGGCGAGGCTCCGGCGAGCTCGACCTCGCGGCGTGCTCGGGCTCGTGTCG

GCGCTGCTCGTCTGCGGCGCTGCGGCGTACGCGCCGGAGGACAACTACCTCGTCAGCTGC

GGCTCCTCGCTGGACACGCCCGTGGGCCGGAGGCTCTTCCTCGCCGACGACGGCGGCTCC

GGCTCCGGCGCGGTCACCCTGACGTCCCCTCGCAGCGCCGCGGTGAAGGCCTCGCCGGAC

CTGGTGTCCGGCTTCCGCGACGCCGCGCTGTACCAGAACGCCAGGGTCTTCTCCGCGCCT

TCATCATACTCGTTCGCCATCAGGCGCCGCGGCCGGCACTTCCTCCGCCTCCACTTCTTC

CCCTTCGTGTACCGGAGCTACGACCTCGCCGCCGCGGCCAGGGCGTTCAAGGTGTCCACG

CAGGACGCCGTGCTGCTCGAGGACGGCGTCCCGGCGCCCGAGCCCGGCAACGCGTCGACG

TCATCGTCGCCCCAGCCGGCGCGCGTGGAGTTCCTCCTGGACGTCGCGCGCGACACGCTC

GTGGTCTCGTTCGTGCCGCTCGTCGACGGGGGCATCGCCTTCGTCAACGCCGTCGAGGTC

GTCTCCGCGCCCGACGGCCTCGTCGCCGACGCCGCCGAGTCGTCGACGGGCCGCCCAGAG

CCCATCCCCGCCGCGCTGCCGCTGCAGACGGCCTACCGCCTCAACGTGGGCGGCCCGGCC

GTCGCGCCCGACGACGACGCGCTCTGGCGAGAGTGGACCACCGACCTGCGATTCCTCTCC

CATTCCGTGGCCGACGCGGTGACTCGGGAGGTCCGCTACAACGGGACGCCGAACCGCCTA

CCCGGGCAAGCGACGGCGACCGACGCGCCGGACGTCGTCTACGCCACGGCGAGGGAGCTC

GTGATCAACAGCAGCTCATTCGACGGGCAGAAACAGATGGCGTGGCAGTTCGACGTCGAC

GCCAGCTCCAGCTACTTCATCAGGTTCCACTTCTGCGACATCGTCGGCAAGGCTCCCCAC

CAGCTCCACATCAACGCCTACGTCGACGACGCCAGTCACGCCACCGTGCTGACGGACCTT

GACCTCGCCGCCGTCGGCGATGGTGCGCTGGCGTTCCCGTACTACAAGGACTTCGTGTTG

CCTGCTAGCGAAGCCTCCGGGAAACTCGCCGTCCATGTTGGCCCTCTGGCAAATAAGATC

GTGATGCCCGCTGCCATCCTCAATGGGATTGAGATCATGAAGATGCACCTGAGCGCCGGC

TCTGTCGTCGTCGTCGAGCCGGCGGCGGGGGCAGCCAAGTCGCGTTTCGCCGTCCTTCTT

GGCTCCGTGTGCGGACCGCTCGCTTTCGTGTCCATCGCCGTTGCTCTCGCCATTGTCCTC

AGGAAGAAGAGGAAGAAGGAGGGGGAGGAAGAGGAGGAAAGTGATAAGAAGCAGCCGACG

CCGACGCAGAGCCAGTCGTCCACGCCATGGATGCCACTCCTCGGCCGCCTCAGCGTTCGC

GGCGCCATTGCGTCAGGGTCGTCAAGCTTCACTACTGCCGGTAACACTCCGGGAACCAGC

CCCAGGGCTGCCGCCGCGGTGATGCCGAGCTACCGTTTCCCGCTCGCCGTGTTGCAAGAC

GCGACGCGCAACTTCGACGACAGCCTGATCATCGGAGAGGGAGGGTTCGGCAAGGTGTAC

GGCGCCGTGCTCCAGGACGGCACTAAGGTCGCCGTGAAGCGCGCGAGCCCGGAGTCGCGG

CAGGGCGCGCGGGAGTTCCGCACGGAGATCGAGCTGCTGTCCGGGCTGCGCCACCGCCAC

CTTGTGTCGCTCGTCGGCTACTGCGACGAGCGGGAGGAGATGATCCTGCTGTACGAGTAC

ATGGAGCACGGCTCGCTGCGGAGCCGGCTGTACGGCCGCGGCGGCGCGGCGCCGCTGAGC

TGGGCGCAGCGGCTGGAGGCGTGCGCCGGCGCGGCGAGGGGCCTCCTTTACCTGCACACG

GCCGTGGACAAGCCGGTGATCCACCGCGACGTCAAGTCGTCCAACATCCTGCTGGACGGC

GACCTCACGGGCAAGGTGGCCGACTTCGGGCTGTCCAAGGCCGGGCCGGTGCTGGACGAG

ACGCACGTCAGCACGGCGGTGAAGGGCAGCTTCGGGTACGTCGACCCGGAGTACTGCCGG

ACGAGGCAGCTGACGGCCAAGTCCGACGTGTACTCGCTGGGCGTCGTGCTGCTGGAGGCC

GTCTGCGCGCGGCCCGTCGTCGACCCGAGGCTGCCGAAGCCCATGTCGAACCTGGTGGAG

TGGGGGCTGCACTGGCAGGGCAGGGGCGAGCTGGAGAAGATCGTGGATCGGCGCATCGCC

GCCGCGGCGAGGCCGGCGGCGCTGAGGAAGTACGGCGAGACGGTGGCCAGGTGCCTGGCG

GAGCGCGGCGCCGACCGGCCGGCCATGGAGGACGTGGTGTGGAACCTGCAGTTCGTGATG

CGGCTGCAGGAGGGTGACGGCCTGGACTTCTCCGACGTGAGCAGCCTCAACATGGTGACA

GAGCTCACGCCGCCACCTCGCCGTCAGAGAAGCGCAGTTGATAGCGACGGCCTGGCCCTC

TCCGATGTGAGCAGCCTCAACATGGTTACAGAGCTCACGCCGCCACAAACCGGCAGCGTG

GAGGGAGACGGCGTAGCCGACGACGATTTCACCGATGCATCCATGAGAGGGACCTTCTGG

CAGATGGTCAATGTCCGCAGCAGATGA

>Ta-CrRLK1L4-B CDS sequence

ATGGCCGCGACGGCGAGGCTCCGGCCGGCACGAGCTCGCGGCGTGCTCTGGAGCGTCTCG

GTGTGGCTCGTCTGCGGCGCTGCGGCGTACGCGCCGGAGGACAACTACCTCGTCAGCTGC

GGCTCCTCGCTGGACACGCCGGTGGGCCGGAGGCTCTTCCTCGCCGACGACGGCGGCTCC

GGCTCCGGCTCCGGCGCGGTCACCCTGACGTCCCCTCGCAGCGCCGCGGTGAAGGCCCCG

CCGGACCTGGTGTCCGGCTTCCGCGACGCCGCGCTGTACCAGAACGCCAGGGTGTTCTCC

GCGCCCTCTTCCTACTCCTTCGCCATCAGGCGCCGCGGCCGGCACTTCCTCCGCCTCCAT

TTCTTCCCCTTCGTGTACCGGAGCTACGACCTCGCCGTGGCGGCCAGGGCGTTCAAGGTG

TCCACGCAGGACGCCGTGCTGCTCGAGGACGGCGTCCCGGCGCCCGAGCCCGGCAACGCG

TCGACGTCGACGTCGCCCCAGCCGGCGCGCGTGGAGTTCCTCCTGGACGTCGCGCGCGAC

ACGCTCGTGGTCTCGTTCGTGCCGCTCGTCGACGGGGGCATCGCGTTCGTGAACGCCGTC

GAGCTCGTCTCCGTGCCCGACGACCTCGTCGCCGACGCGGCGGACTCGTCGACGGGCCGG

CCAGAGCCGATCCCCGCCGCGCTGCCGCTGCAGACGGCCTACCACCTCAACGTGGGCGGC

CCGGCCGTCGCGCCCGACGACGACGCGCTCTGGCGAGAATGGACTACCGACCAGCCCCTC

TCCGATCCTAGGGTCGACGCGGTGACTCGGGAGGTTCGTTACAACAGGACGCTGAACCGC

CTGCCCGGGCAAGCGACGGCGACCGACGCGCCGGACATCGTCTACGCCACGGCGAGGGAG

CTCGTGATCAACCGCAGCTCATTCGACGGGCAGAAACAGATGGCGTGGCAGTTCGACGTC

GACGCGGGCTCCAGCTATTTCATCAGGTTCCACTTCTGCGACATCGTCAGCGAGGCTCCC

CACCAGCTCCACATCAACGCCTACGTCGATGACGCCAGTCACGCCACCGTGCTGACGGAC

CTTGACCTCGCCGCCGTCGGCGATGGTGCGCTGGCGTTCCCGTACTACAAGGACTTCGTG

TTGCCTGCTAGCGAAGCGTCCGGGAAACTCGCCGTCCATGTTGGCCCGTTGGCAAATAAG

ATCGTGATGCCCGCCGCCATCCTCAATGGGATTGAGATCATGAAGATGCACCTGAGCGCC

GGCTCTGTCGTCGTCGTAGAGCCGGCGGCGGGGGCAGCCAAGTCGCGTCTCGCCGTCATT

CTTGGCTCCGTGTGTGGAGCGCTCGCTTTCGTGTCCATCGCCATTGCTCTCGCCATTGTC

CTTAGGAAGAAGAAGGGGGAGGGGGAGGAGGGTGTTAAGGAGCAGCCGACGCCGACGCGG

AGCCAGTCGTCCACGCCATGGATGCCACTCCTCGGCCGCCTCAGCGTTCGCGGCGCCATT

GCATCAGGATCGTCAAGCTTCACTACCGCCGGTAACACTCCGGGAGCGAGCCCGAGGGCT

GCTGCTGCTGCCGCCGCCGCGGTGGTGCCAAGCTACCGTTTCCCGCTCGCCATGTTGCAA

GACGCGACGCGCAACTTCGACGACAGCCTGATCATCGGAGAGGGAGGGTTCGGCAAGGTG

TACGGCGCCGTGCTCCAGGACGGCACCAAGGTCGCCGTGAAGCGCGCGAGCCCGGAGTCG

CGGCAGGGCGCGCGGGAGTTCCGCACGGAGATCGAGCTTCTGTCCGGGCTGCGCCACCGC

CACCTGGTGTCGCTCGTCGGCTACTGCGACGAGCGGGAGGAGATGATCCTGCTGTACGAG

TACATGGAGCACGGCTCGCTGAGGAGCCGGCTGTACGGCCGCGGCGGCGCGGCGCCGCTG

AGCTGGGCGCAGCGGCTGGAGGCGTGCGCCGGCGCGGCGAGGGGCCTCCTGTACCTGCAC

ACGGCCGTGGACAAGCCGGTGATCCACCGCGACGTCAAGTCGTCCAACATCCTGCTGGAC

GGCGACCTCACGGGCAAGGTGGCCGACTTCGGGCTGTCCAAGGCCGGGCCGGTGCTCGAC

GAGACGCACGTGAGCACGGCCGTCAAGGGCAGCTTCGGGTACGTCGACCCGGAGTACTGC

CGGACGAGGCAGCTGACGGCCAAGTCCGACGTGTACTCGCTGGGCGTCGTGCTGCTGGAG

GCCGTCTGCGCGCGCCCCGTCGTCGACCCGAGGCTGCCGAAGCCCATGTCGAACCTGGTG

GAGTGGGGGCTGCACTGGCAGGGCAGAGGCGAGCTGGAGAAGATCGTGGACCGGCGCATC

GCCGCCGTGGCGAGGCCGGCGGCGCTGAGGAAATACGGCGAGACGGTGGCCAGGTGCCTG

GCGGAGCGCGGCGCCGACCGGCCGGCCATGGAAGACGTGGTGTGGAACCTGCAGTTCGTG

ATGCGGCTGCAGGAGGGCGACGGCCTGGACTTCTCCGACGTGAGCAGCCTCAACATGGTG

ACAGAGCTCACGCCGCCTCGCCGTCAGAGAAGCGCGGTCGATTGCGACGGCCTGGACTTC

TCCGACGTGAGCAGCCTCAACATGGTTACAGAGCTCACGCCGCCTAAAACCGGCAGCATG

GAAGGAGACGGTGTAGCCGACGATGACGATTTCACAGACGCATCCATGAGAGGGACCTTT

TGGCAGATGGTCAATGTCCGCAGCAGATGA

>Ta-CrRLK1L4-D CDS sequence

ATGGCCGCGACGGCGAGGCTCCGGCCAGCGCGAGCACGCGGCGTGCTCTGGGTCGTCTCG

GTGTTGCTCGTCTGCGGCGCTGCGGCGTACAAGCCTGAGGACAACTACCTCGTCAGCTGT

GGGTCCTCGCTGGACACGCCGGTGGGCCGGAGGCTCTTCCTCGCCGACGACGGCGCCTCC

GGCGCGGTCACCCTGACGTCCCCTCGCAGCGCCGCGGTGAAGGCCCCGCCGGACCTGGTG

TCCGGCTTCCGCGACGCCGCGCTGTACCAGAACGCCAGGGTGTTCTCCGCGCCCTCCTCC

TACTCCTTCGCCATCAGGCGCCGCGGCCGGCACTTCCTCCGCCTCCACTTCTTCCCCTTC

GTGTACCGGAGCTACGACCTCGCCGCGGCGGCCAGGGCGTTCAAGGTGTCCACGCAGGAC

GCCGTGCTGCTCGAGGACGGCATCCCGGCGCCCGAGCCCGGCAACGCGTCGACGTCGACG

TCGCCCCAGCCGGCGCGCCTGGAGTTCCTCCTGGACGTCGCGCGCGACACGCTGGTGGTC

TCGTTCGTGCCGCTCGCCGACGGGGGCATCGCCTTCGTCAACGCCGTCGAGCTCGTCTCC

GTGCCCGACGGCCTCGTCGCCGACGCGGCGGACTCGTCGACGGGCCGGCCGGAGCCCATC

CCCGCCGTGCTGCCGCTGCAGACGGCCTACCGCCTCAACGTGGGCGGCCCGGCCGTCGCG

CCCGACGACGACGCGCTCTGGCGAGAGTGGACCACCGACCTGCGATTCCTCTCCCATTCT

GTGGCCGACGCGGTGACTCGGGAGGTCCGCTACAACGGGATGCTGAACCGGCTGCCCGGG

CAAGCGACGGCGACCGACGCGCCGGACATCGTCTACGCCACGGCGAGGGAGCTCGTGATC

AACGGCAGCTCATTCGACGGGCAGAAACAGATGGCGTGGCAGTTCGACGTCGACACCAGC

TCCAGCTACTTCATCAGGTTCCACTTCTGCGACATCGTCGGCAAGGCTCCCCACCAGCTC

CACATCAACGCCTACGTCGATGACGCCACCGTGAAGCAGGACCTCGACCTCGCCGCCGTC

GGCGATGGTGCGCTGGCGTTCCCGTACTACACGGACTTCGTGTTGCCTGCTAGCGAGGCG

TCCGGGAAACTCGCCGTCCATGTTGGCCCTCTGGCAAATAAGATCGTCATGCCCGCCGCC

ATCCTCAATGGGATTGAGATCATGAAGATGCACCTGAGCGCCGGGTCCGTCGTCGTCGTC

CAGCCGGCGGCGGGGGCAGCCAAGTCGCGTTTCGCCGTCGTTCTTGGCTCCGTGTGTGGA

GCGCTCGCTTTCATATCCGTCGCCGTTGCTCTCGCCATTGTCCTTAGGAAGAAGGAGAAG

GAGAAGGAGGTGGAGGAGGGTGCCAAGGAGCAGCCGACGCCGACGCAGAGCCAGTCGTCC

ACGCCATGGATGCCACTCCTCGGCCGCTTCAGCGTTCGCGGCGCCATTGCGTCAGGATCG

TCAAGCTTCACCACTGCCGGGAACACTCCGGGAGCGAGCCCCAGGGCTGCTGCCGCTGCC

GCGGTGATGCCGAGCTACCGTTTCCCGCTCGCTATGTTGCAAGACGCGACGCGCAACTTC

GACGACAGCCTCATCATCGGAGAGGGAGGGTTCGGCAAGGTGTACGGCGCCGTGCTCCAG

GACGGCACCAAGGTCGCCGTGAAGCGCGCGAGCCCGGAGTCGCAGCAGGGCGCGCGGGAG

TTCCGCACGGAGATCGAGCTGCTGTCCGGGCTGCGCCACCGCCACCTGGTGTCCCTCGTC

GGCTACTGCGACGAGCGGGAGGAGATGATCCTGCTGTACGAGTACATGGAGCACGGCTCG

CTGAGGAGCCGGCTGTACGGCCGCGGCGGCGCGGCGCCGCTGAGCTGGGCGCAGCGGCTG

GAGGCGTGCGCCGGCGCGGCGAGGGGCCTCCTGTACCTGCACACGGCCGTGGACAAGCCG

GTGATCCACCGCGACGTCAAGTCGTCCAACATCCTTCTGGACGGCGACCTCGCGGGCAAG

GTGGCCGACTTCGGGCTCTCCAAGGCCGGGCCGGTGCTCGACGAGACGCACGTCAGCACG

GCGGTGAAGGGCAGCTTCGGGTACGTCGACCCGGAGTACTGCCGGACGAGGCAGCTGACG

GCCAAGTCCGACGTGTACTCGCTGGGCGTCGTGCTGCTGGAGGCCGTCTGCGCGCGCCCC

GTCGTCGACCCGAGGCTGCCGAAGCCCATGTCGAACCTGGTGGAGTGGGGGCTGCACTGG

CAGGGCAGGGGCGAGCTGGAGAAGATCGTGGACCGGCGCATCGCGGCCGCGGCGAGGCCC

GCGGCGCTGAGGAAGTACGGCGAGACGGTGGCCAGGTGCCTGGCGGAGCGGGGCGCCGAC

CGGCCGGCCATGGAGGACGTGGTGTGGAACCTGCAGTTCGTGATGCGGCTGCAGGAGGGC

GACGGCCTGGACTTCTCCGACGTGAGCAGCCTCAACATGGTGACAGAGCTCACGCCGCCT

CGCCGTCAGAGAAGCGCGGTTGATCACGACGGGCTGGACTACTCCGACGTGAACAGCCTC

AACATGGTTACAGAGCTCACGCCGCCTCAAACCGGCAGCGTGGAGGGAGACGGCGAAGCC

GATGATGATTTCACAGACGCATCCATGAGAGGGACCTTCTGGCAGATGGTCAATGTCCGC

AGCAGATGA

>Ta-CrRLK1L5-A CDS sequence

ATGAGAGGAGGCCCGAGATGCGCGCTCCTGCTCCTCGTGGCCGCCGCGGCGCTTGTCCCC

GCGGCGCGGGCGCAGGGGGCGACCGCGCCCGCGCCCTCGGCGGGGGCCCCGTTCGTGCCG

CGGGACGACATCCTGCTCGACTGCGGCGCCACGGGGAAGGGGAACGACACGGACGGCCGC

CAGTGGGACGGCGACGCCGGGTCCAAGTACGCGCCGCCCAACCTCGCCTCGGCCAGCGCC

GGGGCGCAGGACCCCTCCGTGCCGCAGGTGCCCTACCTCACCGCGCGGGTCTCGGCGGCG

CCCTTCACCTACTCCTTCCCGCTCGGCCCCGGCCGCAAGTTCCTCAGGCTGCACTTCTAC

CCGGCCAACTACTCCAACCGCGACGCCGCCGACGCCTTCTTCTCCGTCTCCGTCCCGGCC

GCCAAGGTCACGCTCCTCTCCAACTTCAGCGCCTACCAGACCACCACGGCGCTCAACTTC

GCCTACATCGTGCGCGAGTTCTCCGTCAACGTCACCGGCCAGAACCTCGACCTCACCTTC

ACCCCGGAGAAGGGCCACCCCAACGCCTACGCCTTCATCAACGGCATCGAGGTCGTCTCC

TCCCCTGACCTCTTTGGCATCGCCACGCCGCAATTCGTCACCGGTGATGGCAACAGCCAG

CCATACGAGATGGATCCTGCTGCTGCTCTGCAGACCATGTATCGGCTCAACGTCGGAGGC

CAGGCCATCTCCCCTTCCAAGGACTCCGGCGGGGCTCGGTCATGGGACGACGACACGCCT

TACATCTATGGTGCAGGGGCTGGGGTATCGTACCAGAACGATCCCAATGTCACAATCATC

TACCCTGACAATGTGCCGGGATATGTGGCACCTTCGGATGTCTACGCCACCGCGCGATCA

ATGGGGCCAGACAAGGGTGTAAACATGGCCTACAATCTCACCTGGATATTGCAGGTGGAT

GCTGGGTACCAATACCTTGTGAGGCTCCATTTCTGTGAGATACAATCCCCATTTACTAAA

CCCAATCAGCGGGTGTTCAACATCTACCTCAACAACCAGACTGCCATTGAAGGTGCTGAT

GTGATCCAGTGGGCGGATCCCAATGGTATTGGTACCCCAGTGTACAAGGACTATGTGGTG

AGCACTGTGGGTTCTGGGATTTTGGATTTCTGGGTGGCTCTACACCCAGATGCAGAGACG

AAGCCACAGTACTATGATGCTATTCTCAATGGGATGGAGGTGTTCAAGCTGCAACTTACT

AATGGGAGCCTCGTGGGGCTCAATCCTGTCCCAAGTGCTGATCCACCAGCGCATAGCGGG

TCAGGAGACAAGAAATCTAAAGTCGCGCCTATTGTTGGTGGAGTAATTGGAGGTTTGGCA

GTGCTTGCGCTTGGATATTGCTGCTTCATCTGCAAGCGTCGGAGGAAAGCGGCCAAGGCT

AGCGGCATGAGTGACGGCCATTCTGGTTGGCTGCCATTGTCGCTGTATGGCCATTCACAC

ACTTCAAGCTCAGCCAAGTCGCATGCTACAGGGAGTTATGCTTCATCTTTGCCGTCCAAC

CTGTGCCGCCATTTCTCCTTTGCAGAGATCAAGGCCGCAACCAAAAACTTTGATGAGTCA

CGGATCCTTGGCGTTGGTGGGTTCGGTAAAGTCTACCATGGAGAGATTGACGGGGGCACA

ACTAAGGTGGCTATCAAGCGTGGCAATCCCTTGTCTGAGCAGGGCATACATGAGTTCCAG

ACTGAAATTGAAATGTTGTCAAAGCTCCGGCACCGTCATCTTGTGTCGCTGATTGGTTAC

TGCGAGGAGAAGAACGAGATGATCCTGGTCTATGACTACATGGCTCATGGAACTCTGCGT

GAGCACCTATACAAGACCCAGAATGCACCGCTTAGCTGGAGGCAGCGTTTGGAGATCTGC

ATCGGTGCAGCTCGTGGGCTTCACTACCTTCACACCGGTGCAAAGCACACCATTATCCAC

CGTGATGTGAAGACGACAAACATCCTCCTGGATGAGAAATGGGTTGCCAAGGTTTCAGAT

TTTGGTCTGTCCAAGACTGGGCCATCGATGGATCACACACATGTGAGCACAGTTGTCAAG

GGCAGTTTTGGTTATCTAGATCCTGAATATTTCCGCAGGCAGCAGCTCACTGAGAAATCT

GATGTGTATTCGTTTGGCGTGGTGCTGTTCGAGGTCCTTTGTGCTCGGCCTGCCTTGAAC

CCCACTCTTGCAAAGGAAGAAGTTAGCTTGGCAGAGTGGGCACTGCACTGCCAAAAGAAG

GGAATTCTTGATCAGATTGTTGATCCCTACCTGAAGGGAAAGATTGTTCCGCAGTGCTTC

AAGAAGTTTGCCGAGACGGCTGAGAAGTGTGTTGCCGACAATGGCATCGAGCGCCCTTCG

ATGGGAGATGTGCTTTGGAACTTGGAGTTTGCTCTTCAGATGCAGGAAAGCGCGGAGGAG

AGTGGAAGCATTGGGTGCGGGATGTCAGATGAGGGCACTCCCCTCGTGATGGTTGGAAAG

AAGGATCCCAATGACCCATCAATTGATTCCAGCACCACTACGACCACAACAACTTCCCTA

AGCATGGGTGACCAAAGTGTCGCGAGCATCGATTCGGATGGGCTGACGCCGAGCGCGGTG

TTCTCACAGATCATGAACCCCAAGGGGCGGTGA

>Ta-CrRLK1L5-B CDS sequence

ATGAGAGGAGGCCCGAGATGCGCGCTCCTGCTCCTCGCGGCTCTGGCCGCCGCCGCGGCG

CTTGCCCCCGCGGCGTGGGCGCAGGGGGCGACCGCGCCAGCGCCCTCGGCGGGGCCCCCG

TTCGTGCCGCGGGACGACATCCTGCTCGACTGCGGCGCCACGGGGAAGGGGAACGACACG

GACGGCCGCCAGTGGGACGGCGACGCCGGGTCCAAGTACGCGCCGCCCAACCTGGCCTCG

GCCACCGCCGGGGCGCAGGACCCCTCGGTGCCGCAGGTGCCCTACCTCACCGCGCGGGTC

TCGGCGGCGCCCTTCACCTACTCATTCCCGCTCGGCCCCGGTCGCAAGTTCCTCAGGCTG

CACTTCTACCCGGCCAACTACTCCAATCGCGACGCCGCCGACGCCTTCTTCTCCGTCTCC

GTCCCGGCCGCCAAGGTCACGCTCCTCTCCAACTTCAGCGCCTACCAGACCACCACGGCG

CTCAACTTCGCCTACATCGTACGCGAGTTCTCTGTCAACGTCACCGGCCAGAACCTCGAC

CTCACCTTCACCCCGGAGAAGGGCCACCCCAACGCCTACGCCTTCATCAACGGCATCGAG

GTCGTCTCCTCCCCCGACCTCTTTGACCTCGCCACGCCGCAATTAGTCACCGGTGACGGC

AACAGCCAGCCATACGAGATGGATCCTGCTGCTGCTCTGCAGACCATGTATCGGCTCAAC

GTCGGAGGCCAGGCCATCTCCCCTTCCAAGGACTCCGGCGGGGCTCGGTCATGGGACGAC

GACACGCCTTACATCTATGGTGCAGGGGCTGGGGTATCGTACCAGAACGATCCCAATGTC

ACAATCACCTACCCTGACAATGTGCCGGGATATGTGGCACCTTCGGATGTCTATGCCACG

GCGCGATCAATGGGGCCAGACAAGGGTGTAAACATGGCCTACAATCTCACCTGGATATTG

CAGGTGGATGCTGGGTACCAATACCTTGTGAGGCTCCATTTCTGTGAGATACAATCCCCA

TTTACTAAACCCAACCAGCGGGTGTTCAACATCTACCTCAACAACCAGACTGCCATGGAA

GGTGCTGATGTGATCCAGTGGGCGGATCCCAATGGTATTGGTACCCCAGTATACAAGGAC

TATGTGGTGAGCACTGTTGGTTCTGGGATTATGGATTTCTGGGTGGCTCTACACCCAGAT

GCAGGGACGAAGCCACAGTATTATGATGCTATTCTCAATGGGATGGAGGTGTTCAAGCTG

CAACTTACTAATGGGAGCCTCGTGGGGCTCAATCCTGTCCCAAGTGCTGATCCACCAGCG

CATAGCGGGTCAGGAGACAAGAAATCTAAAGTCGCGCCTATTGTTGGTGGAGTAATTGGA

GGTTTGGCAGTGCTTGCGCTTGGATATTGCTGCTTCATCTGCAAGCGTCGGAGGAAAGCG

GCCAAGGCTAGCGGCATGAGTGACGGCCATTCTGGTTGGCTGCCATTGTCGCTGTATGGC

CATTCACACACTTCAAGCTCAGCCAAGTCGCATGCTACAGGGAGTTATGCTTCATCTTTG

CCGTCCAACCTGTGCCGCCATTTCTCCTTTGCAGAGATCAAGGCCGCAACCAAAAACTTT

GATGAGTCACGGATCCTTGGTGTTGGTGGGTTCGGTAAAGTCTACCATGGAGAGATTGAC

GGGGGCACAACTAAGGTGGCTATCAAGCGTGGCAATCCCTTGTCTGAGCAGGGCATACAT

GAGTTCCAGACTGAAATTGAAATGTTGTCAAAGCTCCGGCACCGTCATCTTGTGTCTCTG

ATTGGTTACTGCGAGGAGAAGAACGAGATGATCCTGGTCTATGACTACATGGCTCATGGA

ACTCTGCGTGAGCACCTATACAAGACCCAGAATGCACCGCTTAGCTGGAGGCAGCGTTTG

GAGATCTGCATCGGTGCAGCTCGTGGGCTTCACTACCTTCACACCGGTGCAAAGCACACC

ATTATCCACCGTGATGTGAAGACGACAAACATCCTCCTGGATGAGAAATGGGTTGCCAAG

GTTTCAGATTTTGGTCTGTCCAAGACTGGGCCATCGATGGATCACACACATGTGAGCACA

GTTGTCAAGGGCAGTTTTGGTTATCTAGATCCTGAATATTTCCGCAGGCAGCAGCTCACT

GAGAAATCTGATGTGTATTCGTTTGGTGTGGTGCTGTTCGAGGTCCTTTGTGCTCGGCCT

GCCTTGAACCCCACCCTTGCAAAGGAAGAAGTTAGCTTGGCAGAGTGGGCACTGCACTGC

CAAAAGAAGGGAATTCTTGATCAGATTGTTGATCCCTACCTGAAGGGAAAGATTGTTCCC

CAGTGCTTCAAGAAGTTTGCCGAGACGGCTGAGAAGTGTGTTGCCGACAATGGCATCGAG

CGCCCTTCGATGGGAGATGTGCTTTGGAACTTGGAGTTTGCTCTTCAGATGCAGGAAAGC

GCGGAGGAGAGTGGAAGCATTGGGTGTGGGATGTCAGATGAGGGCACTCCCCTCGTGATG

GTTGGAAAGAAGGATCCGAATGACCCATCAATTGATTCCAGCACCACTACGACCACAACA

ACTTCCTTAAGCATGGGTGACCAAAGTGTCGCGAGCATTGACTCGGATGGGCTGACGCCG

AGCGCCGTGTTCTCACAGATCATGAACCCCAAGGGGCGGTGA

>Ta-CrRLK1L5-D CDS sequence

ATGAGAGGAGGCCCGAGATGCGCGCTCCTGCTCCTCGTGGCCGCCGCGGCGCTTGTCCCC

GCGGCGCGGGCGCAGGGGGCGACCGCGCCCGCGCCCTCGTCGGGGGTCCCGTTCGTGCCG

CGGGACGACATCCTGCTCGACTGCGGCGCCACGGGGAAGGGGAACGACACGGACGGCCGC

CAGTGGGACGGCGACGCCGGGTCCAAGTACGCGCCGCCGAAGCTCGCCTCAGCCAGCGCT

GGGGCGCAGGACCCATCGGTGCCGCAGGTGCCCTACCTCACCGCGCGGGTCTCGGCGGCG

CCCTTCACCTACTCCTTCCCGCTCGGCCCCGGCCGCAAGTTCCTCAGGCTGCACTTCTAC

CCGGCCAACTACTCCAACCGCAACGCCGCCGACGCCTTCTTCTCCGTCTCCGTCCCGGCT

GCCAAGGTCACGCTCCTCTCCAACTTCAGCGCCTACCAGACCACCACGGCGCTCAACTTC

GCCTACATCGTGCGCGAGTTCTCCGTCAACGTCACCGGCCAGAACCTCGACCTCACCTTC

ACCCCGGAGAAGGGCCACCCCAACGCCTACGCCTTCATCAACGGCATCGAGGTCGTCTCC

TCCCCCGACCTCTTTGACCTCGCCACGCCGCAATTAGTCACCGGTGACGGCAACAGCCAG

CCGTACGAGATGGATCCTGCTGCTGCTCTGCAGACCATGTATCGGCTCAACGTCGGAGGC

CAGGCCATCTCCCCTTCCAAGGACTCCGGCGGGGCTCGGTCATGGGACGACGACACGCCT

TACATCTATGGTGCAGGGGCTGGGGTGTCGTACCAGAACGATCCCAGTGTCGCAATCACC

TACCCTGACAATGTGCCGGGATATGTGGCACCTTCGGATGTCTATGCCACGGCGCGATCA

ATGGGGCCAGACAAGGGTGTAAACATGGCCTACAATCTCACCTGGATATTGCAGGTGGAT

GCTGGGTACCAATACCTTGTGAGGCTCCATTTCTGTGAGATACAATCCCCATACACTAAA

CCCAATCAGCGGGTGTTCAACATCTACCTCAACAACCAGACTGCCATGCAAGGTGCTGAT

GTGATCCAGTGGGCGGATCCCAATGGTATTGGTACCCCAGTGTACAAGGACTATGTGGTG

AGCACTGTGGGTTCTGGGATTATGGATTTCTGGGTGGCTCTACATCCAGATGCAGAAACC

AAGCCACAGTACTATGATGCTATTCTCAATGGGATGGAGGTGTTCAAGCTGCAACTTACT

AATGGGAGCCTCGTGGGGCTCAATCCTGTCCCAAGTGCTGATCCACCAGCGCATAGCGGG

TCAGGAGACAAGAAATCATTAGTCGCGCCTATTGTTGGTGGAGTAATTGGAGGTTTGGCA

GTGCTTGCGCTTGGATATTGCTGCTTCATCTGCAAGCGCCGGAGGAAAGCTGCCAAGGCT

AGCGGCATGAGTGATGGCCATTCTGGTTGGCTGCCGTTGTCGCTGTATGGCCATTCACAC

ACTTCAAGCTCAGCCAAGTCGCATGCTACAGGGAGTTATGCTTCATCTTTGCCGTCCAAC

CTGTGCCGCCATTTCTCCTTTGCAGAGATCAAGGCCGCAACAAAAAACTTTGACGAGTCA

CGGATCCTTGGTGTTGGTGGGTTCGGTAAAGTTTACCATGGAGAGATTGACGGGGGCACA

ACTAAAGTGGCTATCAAGCGTGGCAATCCCTTGTCTGAGCAGGGCATACATGAGTTCCAG

ACTGAAATTGAAATGTTGTCAAAGCTCCGGCACCGTCATCTTGTGTCGCTGATTGGTTAC

TGCGAGGAGAAGAATGAGATGATCCTGGTCTATGACTACATGGCTCATGGAACTCTGCGT

GAGCACCTATACAAGACCCAGAATGCACCGCTTAGCTGGAGGCAGCGTTTGGAGATCTGC

ATCGGTGCAGCTCGTGGGCTTCACTACCTTCACACTGGTGCAAAGCACACCATTATCCAC

CGTGATGTGAAGACGACAAACATCCTCCTGGATGATAAATGGGTTGCCAAGGTTTCAGAT

TTTGGTCTGTCCAAGACTGGGCCATCGATGGATCACACACATGTGAGCACAGTTGTCAAG

GGCAGTTTTGGTTATCTAGATCCTGAATATTTCCGCAGGCAGCAGCTCACTGAGAAATCT

GATGTGTATTCGTTTGGTGTGGTGCTGTTCGAGGTCCTTTGTGCTCGGCCTGCCTTGAAC

CCCACTCTTGCAAAGGAAGAAGTTAGCTTGGCAGAGTGGGCACTGCACTGCCAAAAGAAG

GGAATTCTTGATCAGATTGTTGATCCCTACCTGAAGGGAAAGATTGTTCCGCAGTGCTTC

AAGAAGTTTGCCGAGACGGCTGAGAAGTGTGTTGCCGACAATGGCATCGAGCGCCCTTCG

ATGGGAGATGTGCTTTGGAACTTGGAGTTTGCTCTTCAGATGCAGGAAAGCGCGGAGGAG

AGTGGAAGCATTGGGTGTGGGATGTCAGATGAGGGCACTCCCCTCGTGATGGTTGGAAAG

AAGGATCCCAATGACCCATCAATTGATTCCAGCACCACTACGACCACAACAACTTCCCTA

AGCATGGGTGACCAAAGTGTCGCGAGCATCGACTCGGATGGGCTGACGCCGAGCGCGGTG

TTCTCACAGATCATGAACCCCAAGGGGCGGTGA

>Ta-CrRLK1L6-A CDS sequence

ATGGCCGTCCACGTCGTGCTCCCCCTCCTCCTCCTCCTTCTCGTCGCCACGGTCCTCCCT

TACACCGCCCTCGCCGCCTTCTCCCCGGACTTCAAGATCTTCCTCGCGTGCGGCGCGGGA

GCCGACGTGCCCTTCCCGTCCGACAACCCCGCGCGCACCTTCGTGCGGGACGACGGCTAC

CTCTCGCAGGGGGGCGCCGCCGCGGTGTCTGCCAGTGCCAGCTCCAACGCGGCCTCCCCT

CTGTACGCCGCCGCGCGCGCCGACACCTCGGCCTTCTCCTACCGCCTCACCTACCCTGCC

GCGCCGGACGCGTCGTCGTTCCTCGTCCTGCGCCTCCACTTCTTCCCGTTCGTCCCCGCC

TCCTCCTCCTCCACCAGTCTTTCCTCCGCGCGGTTCACCGTCTCGGTCCTCGACGCCTAC

GCCCTGCTGCCCGCCTTCTCGCCGCCGGCCGACGGCGTCGTCAAGGAGTTCTTCGTCCCG

CGCGGAGCCTCCGGCGGCGACTTCACCGTCAGGTTCGCCCCGGAAGCCGGCTCCTCCGCG

TTCGTCAACGCCGTCGAGCTGTTCTCGGCCCCGCCGGAGCTGCTGTGGAACAACACGGCG

GTGCCGGTGGACCCTGTGGGGAGCAATGACCTGCCCGAGTGGCCGCTGGACGCGCTGGAG

ACGGTGTACCGCCTCAACGTCGGCGGGCCCCTGCTGACCAACGGGAACGACACGCTGTGG

CGGACGTGGCTCCCCGACGACCCCTACCTCTTCGGCGCGCCCGGGCAGTCGGTGGTGAAC

AACACCCCCAGCCCGATCATCTACGCCCCGTCCAACGGTTACACACAGGAGGTGGCGCCG

GACGTGGTGTACAAGACGCAGCGCGCGGCGAACGTGACGGACCTCCTGCAGGCGACAACC

CCGGGCTTCAACTTCAACGTCACGTGGACGTTCCCGGCGGATCAGGGGTCCCGCTACCTC

GTCCGCCTCCACTTCTGCGACTACGAGGTGGTCAGCTCCGTCGTCGGCACTGTCATCGTC

TTCAACGTCTACGTCGCGCAAGCCATTGGCACTCCAGACCTCACGCCGAGTGCTCGGGCG

AGGCAGTCGAACGAGGCCTTCTACATCGACTACGCGGCCATGGCGCCGAGAGCCGGGAAC

CTCACCGTCAGCATCGGCAGGTCGAAGAAAAGCAGCAAAGGCGGCATACTGAACGGCCTG

GAGATCATGAAGCTGCAAACCGTTAATCTGAGCTCGACGGGGTCGCACGGCCGGACGAAG

AGAATCGTCATAATCGTGCTCGCGACGGTGCTCGGCGCCGCCGTCCTTGCTTCCGCGGTG

CTCTGCTTTTTCGTCGTGCGGCGGAGGAAGCGGAGGCAGGTGGCGCCGCCGGGGTCGACG

GAGGACAAGGAGAGCACGCAGCTGCCGTGGTCACCGTACACGCAGGAAGGCATCTCCGGC

TGGGCCGACGAGTCGGCGAACCGGTCCAGCGAGGGCACCACCGCCAGGATGCAGAGGGTG

AGCACCAAGCTGCACATCTCGCTGGCGGAGCTCAAGGCCGCCACGGACAACTTCCACGAC

CGCAACCTCATCGGCGTCGGCGGGTTCGGCAACGTGTACAAGGGCGCGCTCGCGGACGGC

ACGCCCGTGGCTGTGAAGCGCGCCATGCGCGCCTCCAAGCAGGGCCTGCCGGAGTTCCAC

ACGGAGATCGTGGTGCTGTCCGGCATCCGGCACCGGCACCTCGTCTCGCTCATCGGCTAC

TGCAACGAGCAGGCGGAGATGATTCTGGTGTACGAGTACATGGAGAAGGGCACGCTGCGG

GGCCACCTGTACGGCGGCTCCGACGACGAGCCGCCGCTCTCGTGGAAGCAGCGGCTCGAG

ATCTGCATCGGCGCCGCCAGGGGCCTGCACTACCTGCACAGCGGCTACTCCGAGAACATC

ATCCACCGCGACGTCAAGTCCACCAACATCCTCCTCGGCACCGACGGCGGCGGCAGCACC

GGCGGCGGCGCGATCATCGCCAAGGTGGCCGACTTCGGTCTCTCGCGCATCGGGCCGTCG

CTGGGGGAGACGCACGTCAGCACGGCCGTCAAGGGCAGCTTCGGGTACCTCGACCCCGAA

TACTTCAAGACGCAGCAGCTCACGGACCGCTCCGACGTCTACTCCTTCGGCGTGGTGCTC

TTCGAGGTGCTCTGCGCGCGCCCGGTCATCGACCAGAGCCTGGACCGCGACCAGATCAAC

ATCGCCGAGTGGGCCGTCAGGATGCACGGGGAGGGGAAGCTCGACAAGATCGCCGACGCC

AGGATCGCCGGCGAGGTGAACGACAACTCGCTGCGCAAGTTCGCCGAGACGGCCGAGAGG

TGCCTGGCCGACTACGGCGCGGACCGGCCGTCCATGGGCGACGTGCTGTGGAACCTCGAG

TACTGCCTGCAGCTGCAGGAGACGCACGTCAACAGGGACGCGTTCGAGGACAGCGGCGCC

GTCGCCACGCAGCTCCCCGCCGACGTGGTCGTGCCGCGGTGGGTGCCATCGTCCACCAGC

CTGCTGATGATGGACGACGCGGACGAGACGGGCCTGAGCATGACCGACCTCGCCGATAGC

CAGGTCTTCTCCCAGCTGAACGCCCGTGGCGAGGGGCGATGA

>Ta-CrRLK1L6-B CDS sequence

ATGGCCGTCCACGTCGTACCACCCCTCCTCCTCCTCCTCGCCACGGCCCTCCCGTACTCC

GCCCTCGCCGTTTTCTCCCCGGATTTCTCCTTCTTCCTCGCGTGCGGCGCAGGCGCCGAC

GTCACCTTCCCGTCAGACAACCCCACGCGCACCTTCGTGCGGGACGACGGCTACCTCTCG

CAAGGGCGTCCCGCCGCGGTGTCTGCAAACGCCAGCTCCGGCGCGGCCTCCAACCCTCTG

TACGCCGCCGCGCGCGCCGACAGCTCGGCCTTCTCCTACCGCCTCGCGTACCCCGCCACG

GCGGGCGCGTCGTCGTTCCTCGTCCTGCGCCTCCACTTCTTCCCGTTCGTCCCCGCCTCC

TCCTCCACCAGCCTTTCCTCCGCGCGCTTCACCGTCTCGGTCCTCGACGCCTACGCCCTG

CTGCCCGCCTTCTCGCCGCCGGCCGACGGCGTCGTCAAGGAGTTCTTCGTCCCGCGCGGC

GGGTCGAAAGAATTCACCATCAGGTTCAGCCCGGACGCCGGCTCCTCCGCTTTCGTCAAT

GCAGTCGAGCTGTTCCCAGCCCCGCAGCAGCTGCTGTGGAACGGCTCCAACTCGGTGGTG

CCGGTGGGCGTCCTGGGGAACGACGACTTGGCCCAGTGGCAGCTGGACGCGCTGGAGACG

GTGTACCGCCTCAACGTCGGCGGGCCCAAGGTGACCAGGGAGAACGACACGCTCTGGCGG

ACGTGGCTCCCCGACGGCGCCTACCTCTTCGGCGCCCCCGGGCAGTCGGTGGTGAACAAC

ACCTCCAGCCCGATCATCTACAACCCGCCGAACACAAGGGAGGTGGCACCGGACGTGGTG

TACAGGACGCAGCGCGCGGCGAACGTGACGGACTTCCTGCGGGCGACAACGCCGGGCCTG

AACTTCAACGTCACGTGGACGTTCCCGGCGGAGGCAGGGTCCCGCTACCTCGTCCGCCTC

CACTTCTGCGACTATGAGGTGGTCAGCTCCGTCGTCGGTGTTGGCATCGTCTTCAACGTC

TATGTCGCGCAAGCCATTGGCAGCAGAGACCTCGCGCCGAATGCTCAGGCGACTCAGCCG

AACGAGCCCTTATATCTTGACTACGCGGCCACGGCGCCGAGAGCTGGGAACCTCACCGTC

AGCATTGGCACGTCGTCGAAAAGCAGCGGGGGCGGCATACTGAACGGGCTGGAGATCATG

AAGCTGCAATCCGTCGACCTGAGCTCGCCGGGGTCGCATGCCCTGACGAAGAGAAGCATC

ATCATCATCGTGCTCGCGACGGTGCTCGGCGCCGCCGTCCTTGCGTGCGCGGTGCTCTGC

TTTTTCGTCGTGCGGCGGAGGAAGCGCAGACAGGTGGCGCCGCCGGCGTCGAAGGAGGAT

AAGGAGAGCACGCAGCTGCCGTGGTCACCGTACACGCAGGAAGGCATCTCCGGGTGGGCC

GACGAGTCGACGAACCGGTCCAACGAGGGCACGACCGCCAGGATGCAGAGGGTGAGCACC

AAGCTGCACATCTCGCTGCCGGAGCTCAAGGCCGCCACGGACAACTTCCACGAGCGCAAC

CTCATCGGCGTCGGCGGGTTCGGCAACGTGTACAAGGGCGCGCTCTCCGACGGCACGCCC

GTGGCGGTGAAGCGCGCCATGCGCGCCTCCAAGCAGGGCCTGCCGGAGTTTCAGACCGAG

ATCGTGGTGCTGTCCGGCATCCGGCACCGGCACCTGGTGTCGCTCATCGGCTACTGCAAC

GAGCAGGCGGAGATGATCCTGGTGTACGAGTACATGGAGAAGGGCACGCTGCGGAGCCAC

CTGTACGGCTCCGACGAGCCGGTGCTGTCGTGGAAGCAGCGGCTCGAGATCTGCATCGGC

GCCGCCAGGGGCCTGCACTACCTGCACAGCGGCTACTCGGAGAACATCATCCACCGCGAC

GTCAAGTCCACCAACATCCTCCTCGGGACCGACGACGGCGGCAGCACCGGCGGCGGCGCG

ATCATCGCCAAGGTGGCCGACTTCGGGCTGTCGCGCATCGGGCCATCGCTGGGGGAGACG

CACGTCAGCACGGCGGTGAAGGGCAGCTTCGGGTACCTGGACCCTGAGTACTTCAAGACG

CAGCAGCTCACGGACCGCTCCGACGTCTACTCCTTCGGCGTGGTGCTCTTCGAGGTGCTC

TGCGCGCGCCCGGTCATCGACCAGAGCCTCGACCGCGACCAGATCAACATCGCTGAGTGG

GCCGTGAGGATGCACGGGGAGGGGAAGCTCGACAAGATCGCCGACGCCAGGATCGCGGGT

GAGGTGAACGACAACTCGCTGCGCAAGTTCGCCGAGACGGCGGAGAAGTGCCTCGCGGAC

TACGGCGCCGACCGCCCCTCCATGGGCGACGTGCTGTGGAACCTCGAGTACTGCCTCCAG

CTGCAGGAGACACACGTCAACAGGGACGCCTTCGAGGACAGCGGCGCCGTCGCCACGCAG

CTCCCCGCCGACGTCGTCGTGCCGCGCTGGGTGCCGTCGTCGACGAGTCTGCTCATGATG

GACGACGCGGACGAGACGGGACTGAGCATGACCGAGCTCGCCGACAGCCAGGTCTTCTCC

CAGCTCAACGCTCGCGGCGAGGGACGATGA

>Ta-CrRLK1L6-D CDS sequence

ATGGCCGTCCACGTCGTACCACCCCTCCTCCTCCTCCTCCTCGCCACGGCCCTCCCGTAC

ACCGCCCTCGCCGTTTTCTCCCCGGATTTCTCCTTCTTCCTCGCGTGCGGCGCAGGCGCG

GACGTCCCCTTCCCATCCGACAACCCCACGCGCACCTTCGTGCGGGACGACGGCTACCTC

TCGCAAGGGCGCCCCGCCGCGGTGTCTGCCAGTGCCAGCTCCGGCGCGGCCTCCAACCCT

CTGTACGCCGCCGCGCGCGCCGACAGCTCGGCCTTCTCCTACCGCCTCGCGTACCCCGCC

ACGGCGGGCGCGTCGTCGTTCCTCGTCCTGCGCCTCCACTTCTTCCCGTTCGTCCCCGCC

TCCTCCTCCACCAGCCTTTCCTCCGCGCGCTTCACCGTCTCAGTCCTCGACGCCTACGCC

CTCCTGCCTACCTTCTCGCCGCCGGTCGCCGGCGTCGTCAAGGAGTTCTTCGTCCCGCGC

GACGGGTCGAAAGATTTCACCATCAGGTTCACCCCGGACGCCGGCTCCTCCGCGTTCGTC

AACGCCGTCGAGCTGTTCTCGGCCCCGCCGGAGCTGCTGTGGAACAACACGGCGGTACCG

GTGGACCCCGTGGGGAGCAATGACCTGCCCGAGTGGCCGCTGGACGCGCTGGAGACGGTG

TACCGCCTCAACGTCGGCGGGCCCATGGTGACCAAGGAGAACGACACGCTCTGGCGGACG

TGGCTTCCCGACGGCCCCTACCTCTTCGGCGCCCCCGGGCAGTCGGTTGTGAACAGCACC

TCCAGCCCGATCATGTACGACCCGTCCAACGGTTACACACAGGATGTGGCGCCGGACGTG

GTGTACAGGACGCAGCGCGCGGCGAACGTGACGGACCTCCTGGTGGCGACAACGCCGGGC

CTGAACTTCAACGTCACGTGGACGTTCCCGGCGGAGCAGGGGTCCCGCTACCTCGTCCGC

CTCCACTTCTGCGACTATGAGGTGGTCAGCTCCGTCGTCGGTGTTGGCATCGTCTTCAAC

GTCTATGTCGCGCAAGCCATCGGCACTCCAGCCCTCTCGCCAAAGGATCGGGCGAGGCAG

TCGAACGAGGCCTTTTACATGGACTACGCGGCCAGGGCGCCGAGAGCCGGGAACCTCACC

GTCAGCATCGGCTGGTTGCGGCAAAGCAGCGGAGGCGGCATACTCAACGGGCTGGAGATC

ATGAAGCTGCAATCCGCCGACCCGAGCTTGACGGTGTCGCACGGCCTGACGAAGAGAAGC

ATCATCATCATCGTGCTCGCGACGGTGCTCGGCGCCGCCGTCCTTGCGTGCGCGGTGCTC

TGCTTTTTCGTCGTGCGGCGGACGAAGCGGAGGCAGGTGGCGCCGCCGGCGTCGACGGAG

GATAAGGAGAGCACGCAGCTGCCGTGGTCACCGTACACGCAGGAAGGCATCTCCGGCTGG

GCCGACGAGTCGACGAACCGGTCCAGCGAGGGCACCACCGCCAGGATGCAGAGGGTGAGC

ACCAAGCTGCACATCTCGCTGGCGGAGCTCAAGGCCGCCACGGACAACTTCCACGACCGC

AACCTCATCGGCGTCGGCGGGTTCGGCAACGTGTACAAGGGCGCGCTCGCCGACGGCACG

CCCGTGGCGGTGAAGCGCGCCATGCGCGCCTCCAAGCAGGGCCTGCCGGAGTTCCACACG

GAGATCGTGGTGCTGTCCGGCATCCGGCACCGCCACCTCGTGTCGCTCATCGGCTACTGC

AACGAGCAGGCAGAGATGATCCTGGTGTACGAGTACATGGAGAAGGGCACGCTGCGGAGC

CACCTGTACGGCGGCTCCGACGACGAGCCGCCGCTCTCGTGGAAGCAGCGGCTCGAGATC

TGCATCGGCGCCGCCAGGGGCCTGCACTACCTGCACTGCGGCTACTCGGAGAACATCATC

CACCGCGACGTCAAGTCCACCAACATCCTCCTCGGCACCGACGACGGCGGCAGCACCGGC

GGCGGCGCCATCATCGCCAAGGTGGCCGACTTCGGGCTGTCGCGCATCGGGCCATCGCTG

GGGGAGACGCACGTCAGCACGGCGGTGAAGGGCAGCTTCGGGTACCTGGACCCGGAGTAC

TTCAAGACGCAGCAGCTCACGGACCGCTCCGACGTCTACTCCTTCGGCGTGGTGCTCTTC

GAGGTGCTCTGCGCGCGCCCCGTCATCGACCAGAGCCTCGACCGCGACCAGATCAACATC

GCCGAGTGGGCCGTCAGGATGCACGGGGAGGGGAAGCTCGACAAGATCGCCGACGCCAGG

ATCGCCGGCGAGGTCAACGACAACTCGCTGCGCAAGTTCGCCGAGACGGCGGAGAGGTGC

CTGGCTGACTACGGCGCGGACCGGCCGTCCATGGGCGACGTGCTCTGGAACCTCGAGTAC

TGCCTCCAGCTGCAGGAGACGCACGTCAACAGGGACGCCTTCGAGGACAGCGGCGCCGTC

GCCACGCAGCTCCCCGCCGACGTGGTCGTGCCGCGGTGGGTGCCGTCGTCCACGAGCCTG

CTGATGATGGACGACGCGGACGAGGCGGGCCTGAGCATGACCGAGCTCGCCGACAGCCAG

GTCTTCTCCCAGCTCAACGCACGCGGCGAGGGACGATGA

>Ta-CrRLK1L7-B CDS sequence

ATGCTGTGGAATAGCTCCGTGACGCCCGTGGGAGCCGTGGTGAAGGACGACATGGACCTG

TGGCAGCGGCAGCCGCTGGAGACGGTCTATCGCCTCAACGTCGGAGGGCCCAAGGTGACC

ATTGAGAACGACACGCTGTGGCGGACGTGGCTGCCCGACGGTCCCTACCTCTACGACGCC

TCCGGGCTGTCGGTGGTGAGCAACACCTCCAACCCGATCATCTACGATTCATCGAACGGA

TACACGAGGGAGGTGGCGCCAGATGTCGTGTACCAGACCCAGCGCATGGCGAACGTGACG

GACTTACTGGCGGCGACAACCCCGGGCCTGAACTTCAACCTCACGTGGACGTTCCCGGCG

GTGAAGGGGTCCCACTACCTCGTCCGCCTCCACTTCTGCGACTACGAGGTGGTCAGCTCC

GTCGTCGGCGTTGGCATCGTCTTCAACGTCTACATCGCGCAGACCATTGGCACTCCAGAC

CTCACGCCGAATGCTCGGGCGACTCAGTCGAACGAGGTCTTTTACATGGACTACGCGGCC

AGGGCGCCGAGCACCGGGAACCTCACGGTGAGCATCGGCTGGTCGTCGAAAAGGAGCGGA

GGTGGGATACTGAACGGGCTAGAGATTATGAGGCTGCCGCCCGTTGATTTGAGCTCGAGG

AGGTACGGCAGGACGAAGAGGACCATTGTCATTACGGTGTCGGCAGTGCTCGGCGCCGCC

GTTCTTGCTTGCGTGGTGCTCTGCTTTTTCGGCGTGCCGTATACGAAGTACAGCGGCTCC

GGCTGGGCTGAGCAGTTCACGAACCGATGGTCCAGAGAGGGCAAGACCAGCGGGTTGCAG

AGTGTGAGCACGAAGCTGCACATCGCTCTCGCGAAGATCAAGGCCGCCACGGACAACTTC

CACGAGCGCAACCTCATCGGCGTGGGCGGGTTCGGGAACGTGTACAAGGGCGTGCTCGTT

GACGGCACGCCAGTGGCGGTGAAGCGCGCCATGCGCGCCTCGCAGCAGGGGTTGCCGGAG

TTCCAGACGGAGATCGTGGTGCTGTCCGGCATCCGGCACCGGCACCTGGTGTCGCTCATT

GGGTACTGCAACGAGCAGGCGGAGATGATACTGGTGTACGAATACATGGAGAAAGGCACG

CTGCGGAGCCACCTGTACGGTTCCGACGAGCCGGCGTTGTCATGGAAGCAGAGGCTGGAG

ATCTGCATCGGCGCGGCGAGGGGCCTGCACTACCTGCACAGAGGCTACGCGGAGAACATC

ATCCACCGTGACGTCAAGTCGACCAACATCCTCCTCGGGAGCGACGGCGGCAGCACCGGT

GGCGTGATCGCCAAGGTGGCCGACTTCGGGCTGTCGCGCATCGGGCCGTCGTTCGGGGAG

ACGCACGTGAGCACGGCGGTGAAGGGCAGCTTCGGGTACCTGGACCCGGGGTACTTCAAG

ACGCAGCAGCTGACGGACCGGTCGGACGTCTACTCCTTCGGCGTGGTGCTGTTGGAGGTG

CTCTGCGCGCGACCTGTGATCGACCAGAGCCTGGACCACGGCCGGATCAACATCGCCGAA

TGGGCCGTGAGGATGCGCAGGGAAGGGCGGCTCGACAAGATGGCCGACCCGAGGATCGCC

GGCGAGGTGGACGAGGAGTCGCTGCTCAAGTTCGCAGAAACCGCTGAGAAGTGCCTGGCG

GAGTGCTGGGTGGACCGGCCGTCCATGGGCGACGTGCTGTGGAACCTGGAGTATTGCCTA

CAGCTGCAGGAGACCAATATCACCGGGGACGGACTCGACGACATGGTACCGTCGTCGACG

AGCTTGTTGATGGACGAGACCGACTTGAGCATGACCAATGTCGCCGACAGCAAGGTATTC

TCCCAGCTGAGCGCCCGCGGCGAGGGACGATGA

>Ta-CrRLK1L7-D CDS sequence

ATGGCCGTCCGCGGCATACTCCTCGCCCTCCTCCTCGCCATGGTTCTCCCGCGCGCCATC

CTCGCCGCCTTCTCTCCCGGCTTCCAATATTTCCTCGCATGCGGCGCCAACTCCGCCGTC

TCCTTCCCGTCCGATTCCCCCGCCAACATCTTCGTCCCCGACGCCGCCTACCTCTCGCCC

GCGGGCGCTCCGGCGGTGTCCGCCAGCTCCACCCTCGCCTCCCCGCCAGCTCTGTACGCC

GCCGCGCGCGCGGACATCTCGGCCTTCTCGTACCGCCTCCCTAGCCCCGCCTCGCCAGAC

ACGTCGTCATTCCTCGTCCTGCGCCTCCACTTCTTCCCCTCCTTCCCCGCCACCTCCTCT

CAGTATGTCATCAACATCTTGTCCGCGCGCTTCAACGTTTCGGTCGCCGACGCCTACGCT

CTGCTGTCCTCCTTCTCGCCTCCGGCCGCCGGCGTCGTCAAGGAGTTCTTCGTCCCGCGC

GACCTCTTCGATGGCCACTTCCACGTCACGTTCACCCCGGACGCCGGCTCCACCGCCTTC

GTCAACGCCATCGAGCTGTTCTCGGCCCCGCCGGAGATGCTGTGGAATGGCCCCGTGACG

CCGGTGGGAGCCGTGGTGAAGGACGACATGGACCTGTGGCAGCGGCAGCCGCTGGAGACG

GTCTATCGCCTCAACGTCGGCGGGCCCAAGGTGATAATTGAGAACGACACGCTGTGGCGG

ACGTGGCTGCCCGACGGCCCCTACCTCTACGACGCCTCCGGGCTGTCGGTGGTGAGCAAC

ACCTCCAGCCCGATCATCTACGATTCATCGAACGGATACACGAGGGAGGTGGCGCCGGAC

GTCGTGTACCAGACCCAGCGCATGGCGAACGTGACGGACTTACTGGCGGCGACAACCCCA

GGCCTGAACTTCAATCTCACGTGGACGTTCCCGGCGGTGAAGGGGTCCCGCTACCTCGTC

CGCCTCCACTTCTGCGACTACGAGGTGGTCAGCTCCGTCGTCGGCGTCGGCATCGTCTTC

AACGTCTACATCGCACAGGCCATTGGCACTCCAGACCTCACGCCGAATGCTCGGGCGACT

CAGTCGAACGAGGTCTTTTACAAGGACTACGCGGCCAGGGCGCCGAGCGCCGGGAACCTC

ACGGTGAGCATCGGCTGGTCGTCGAAAAGCAGCGGAGGTGGCATACTGAACGGGCTAGAG

ATTATGAGGCTGCCGCCCGTCGATTTGAGCTCGAGGAGGTACGGCAGGACGAAGAGGACC

ATTGTCATTACGGTGTCGGCAGTGCTCGGCGCCGCCGTTCTTGCTTGCGTGGTGCTCTGC

TTTTTCGGCGTGCCGTACACGAAGTACAGTGGCTCCGGCTGGGCTGAGCAGTTCATGAAC

CGATGGTCCAGAGAGCGCAAGACCGGCGGGATGGAGAGTGTGAGCAGGAAGCTGCACATC

GCGCTCGCGAAGATCAAGGCCGCCACGGACAACTTCCACGAGCGTAACCTCATCGGCGTG

GGCGGGTTCGGGAACGTGTACAAGGGCGTGCTCGGTGACGGCACGCCAGTGGCGGTGAAG

CGCGCCATGCGCGCCTCGCAGCAGGGGTTGCCGGAGTTCCAGACGGAGATCGTGGTGCTG

TCCGGCATCCGGCACCGGCACCTGGTGTCGCTCATCGGGTACTGCAACGAGCAGGCGGAG

ATGATACTGGTGTACGAATACATGGAGAAAGGCACGCTGCGGAGCCACCTGTACGGCTCC

GACGAGCCGGCGTTGTCATGGAAGCAGAGGCTGGAGATCTGCATCGGCGCGGCGAGGGGC

CTGCACTACCTGCACAGAGGCTACGCGGAGAACATCATCCACCGTGACGTCAAGTCGACC

AACATCCTCCTCGGGAGCGACGACGGCAGCACCGGTGGCGTGATCGCCAAGGTGGCCGAC

TTCGGGCTGTCGCGCATCGGGCCGTCGTTCGGGGAGACGCACGTGAGCACGGCGGTGAAG

GGCAGCTTCGGGTACCTGGACCCGGGGTACTTCAAGACGCAGCAGCTGACGGACCGGTCG

GACGTCTACTCCTTCGGCGTGGTGCTGTTGGAGGTGCTCTGCGCGCGACCTGTGATCGAC

CAGAGCCTGGACCACGGCCGGATCAACATCGCCGAATGGGCCGTGAGGATGCGCAGGGAA

GGGCGGCTCGACAAGATGGCCGACCCGAGGATCGCCGGCGAGGTGGACGAGGAGTCGCTG

CTCAAGTTCGCAGAGACCGCTGAGAAGTGCCTGGCGGAGGCCTGGGTGGACCGGCCGTCC

ATGGGCGACGTGCTGTGGAACCTGGAGTATTGCCTACAGCTGCAGGAGACCAACATCACC

GGGGACGAACTCGACGACATGGTGCCGTCGTCGACGAGCTTGTTGATGGACGAGACCGGC

TTGAGCATGACCAATGTCGCCGACAGCAAGGTATTCTCCCAGCTGAGCGCCCGCGGCGAG

GGACGATGA

>Ta-CrRLK1L8-A CDS sequence

ATGCCGCCGGTTCCTGACATGCTCGTGCGGCTCCTCGTCGCGTCCGTGCTGCTCGGCGCA

GCCAGTGGCGCGTTTACCCCCGCGGACACCTACCTCGTCCTCTGCGGCACGTCGGCGAGC

GCCACCGTTGCCGCGGGACGGACGTTCGTCGGGGACGCCCGTCTGCCCGCCAAGTCGCTG

GCCGCGCCGCAGAGCGTCGAGGCCAACACGTCGCTGACCGCGGTCGTCCCGTCCGGCGAG

TCGCAGCTTTACCGGTCCGCGCGCGTCTTCACCGCGCCGGCTTCCTACACGTTCGCCGTC

AAGCAGCCCGGCCGGCACTTCGTGCGCCTCCACTTCTTCCCCTTCCCGTACCGGTCCTAC

GACATGGCCGCGGACGCCGCGTTCAACGTGTCCGTGCAGGGCGCGGTGCTCGTCAACGGG

TACGCGCCCAAGAACGGCACGGCGGAGCTCAGGGAGTTCTCGCTGAACGTCACCGGTGCC

ACGCTGGTGATCGCCTTCGCGCCGACGGGGAAGCTCGCGTTCGTGAACGCCATCGAGGTC

GTGTCGGTCCCCGACGAGCTCATCGCCGACACGGCCAGGATGGTGGGCGGGGCCGTCCAG

TACACCGGGCTGTCGACGCAGGCGCTGGAGACGATCCACCGGATCAACATGGGCGTCCCC

AAGATCACGCCCGGCAACGACACGCTGGGGAGGACGTGGCTGCCGGACCAGAGCTTCCAG

CTCAACACCAACCTAGCGCAGCATAAAGACGCCAAGCCCTTGACGATCAAATACGACGAG

AAGTCGGCGCTCTCCTCCGCGTACACGGCGCCGGCGGAGGTCTACGCGACGGCGACGAGG

CTGAGCACGGCGGGCGAGACCAGCACCATCAACGTGCAGTTCAACATCAGCTGGAGGTTC

GACGCCCCGGCCGGGTCGGATTACCTGCTCCGGTTCCACTGGTGCGACATCGTCAGCAAG

GCGGCCATGGGAATGGCCTTCAACGTCTACGTCGGCGGGTCGGTGGTGCTCGAAAACTAC

GAGATTTCGCGTGACACGTTCAACCGGCTGTCCATACCGGTGTACAAGGACTTCCTCCTG

GGCGCCAAGGACGCCAAGGGCGCCATCACCGTGAGCATCGGGTCGTCGACCGAGGACAAC

GCGTTGCCCGACGGCTTCCTCAACGGCCTCGAGATCATGAGGGTAGTCGGGAGCGCCGGC

GCCGGCGCTCCCGCCGCGTCCGCGCGCAGTTCAAAGGTCAAAATCGGGATCATCGCCGGC

TCGGCCGTCTGCGGGGCCACGCTGGTGATGGTGCTCGGGTTCATCGCCTTCAGGACGCTG

CGCGGGAGGGAGCCGGAGAAGAAGCAGCCGTCCGACACCTGGTCGCCCTTCTCGGCGAGC

GCGCTGGGCTCTCGCTCGCGCTCCCGGAGCTTCTCCAAGAGCAACGGGAACACCGTCCTG

CTCGGGCAGAACGGCGCCGGCGCCGGGTACAGGATCCCGTTCGCGGCGCTCCAGGAGGCG

ACCGACGGGTTCGACGAGGCGATGGTCATCGGCGAGGGCGGGTTCGGGAAGGTGTACAAG

GGCACGATGCGCGACGAGACGCTGGTGGCCGTGAAGCGCGGCAACCGGCGGACGCAGCAG

GGGCTGCACGAGTTCCACACGGAGATCGAGATGCTGTCCCGGCTGCGCCACCGCCACCTG

GTCTCGCTCATCGGCTACTGCGACGAGCGCGGGGAGATGATCCTGGTGTACGAGTACATG

GCCATGGGCACGCTGCGGAGCCACCTGTACGGCGCCGGCCTCCCGCCGCTGTCGTGGGAG

CAGAGGCTCGAGGCCTGCATCGGCGCCGCGCGGGGGCTGCACTACCTCCACACCGGCTCC

GCCAAGGCGATCATCCACCGGGACGTCAAGTCGGCCAACATCCTCCTGGACGAGAGCTTC

ATGGCCAAGGTGGCCGACTTCGGGCTGTCCAAGAACGGGCCGGAGCTGGACAAGACGCAC

GTGAGCACCAAGGTGAAGGGCAGCTTCGGGTACCTGGACCCGGAGTACTTCCGGCGGCAG

ATGCTGACGGAGAAGTCGGACGTCTACTCCTTCGGCGTGGTCCTGCTGGAGGTGCTGTGC

GCCCGCACGGTCATCGACCCGACGCTGCCCCGGGAGATGGTGAGCCTGGCCGAGTGGGCG

ACGCCGTGTCTCAGGAACGGTCGGCTCGACCAGATCGTCGACCAGAGGATCGCCGGGACG

ATACGGCCGGGGTCGCTCAAGAAGCTCGCGGACACGGCCGAGAAGTGCCTCGCCGAGTAC

GGGGTGGAGCGGCCCACCATGGGGGACGTGCTCTGGTGCCTCGAGTTCGCGCTGCAGCTG

CAGGTGGGGTCGTCAGACGGCTCGGACGTCGACACCATGTTGCCGCCGGCGCCGCCTGTG

CCCGTGAAAACGCCAGAGGTTCAGCGTAGCCTGTCCGCCGCTACCATGGCGACCGACGCT

GCTGCCATGACCACCAACTTGGGTGATCTAGACGGAATGTCCCTGAGCGGAGTATTCTCG

AAGATGATCAAGAGCGACGAGGTCAGGTGA

>Ta-CrRLK1L8-B CDS sequence

ATGCCGCCGTTTCCTGACATGCTCGTGCGGCTCCTCCTCGCGTCCGTGCTGCTCAGCGCA

GCCAGTGGCGCGTTTACCCCCGCGGACAACTACCTCGTCATCTGCGGCACGTCGGCGAGC

GCCACCGTCGCCCCGAGACGTTCGTTCGTCGGGGACGCCCGCCTGCCCGCCAAGTCGCTG

GCCGCGCCGCAGAGCGTCGAGGCCAACACGTCGCTGACCGCGGTCGTCCCGTCCGGCGAG

TCGGAGCTTTACCGGTCCGCGCGAGTCTTCACCGCACCAGCTTCCTACACGTTCGCCGTC

AAGCAGCCCGGCCGGCACTTCGTGCGCCTCCACTTCTTCCCCTTCTCGTACCGGTCCTAC

GACATGGCCGCGGACGCCGCGTTCAACGTGTCCGTGCAGGGCGCGGTGCTCGTCAACGGG

TACACGCCCAAGAACGGCACGGCAGAGCTCAGGGAGTTCTCCGTGAACGTCACCGGGGGA

ACGCTGGTGATCGCGTTCGCGCCGACGGGGAAGCTCGCGTTCGTGAACGCCATCGAGGTC

GTGTCCGTCCCCGACGAGCTCATCGCCGACACGGCCAGGACGGTGGGCGGCGCCGTCCAG

TACACCGGGCTGTCGACGCAGGCGCTGGAGACGATCCACAGGATCAACATGGGCATTCCC

AAGATCACGCCCGGCAACGACACGCTGGGGAGGACGTGGCTGCCGGACCAGAGCTTCCAG

CTCAACACCAACTTGGCGCAGCACAAAGACGCCAAGCCCTTGACGATCAAATACGACGAG

AAATCGGCACTCTCTTCCCCGTTCACGGCGCCGGCGGAAGTCTACGCGACGGCGACGAGG

CTGAGCACAGCGGGCGAGACCAGCACCATCAACGTGCAGTTCAACATCAGTTGGAGGTTC

GACGCCCCGGCCGGGTCGGATTACCTGCTCCGGTTCCACTGGTGCGACATCGTCAGCAAG

GCGGCCATCGGAATGGCCTTCAACGTCTACGTCGGCGGGTCGGTGGTGCTCGACAACTAC

GAGATCTCGCGTGACACGTTCAACCGGCTATCCATACCGGTGTACAAGGACTTCGTCCTG

GGCGCCAAGGACGCCAAGGGCGCCATCACCGTGAGCATCGGGTCGTCGACCGAGGACAAC

ACATTGCCTGACGGCTTCCTGAACGGCCTCGAGATCATGAGGGTAGTCGGGAGCGCCAGC

GCCGGCGCCGGCGCTCCCGCCGCGTCCCCGCCCAGTTCAAAGGTCAAAATCGGGATCATC

GCCGGCTCGGCCGTTTGCGGGGCCACGCTGGTAACGGTGCTCGGGTTCATCGCCTTCAGG

ATGCTGCGCGGGAGGGAGCCGGAGAAGAAGCAGCCGTCCGACACCTGGTCGCCGTTCTCG

GCGAGCGCGTTGGGCTCTCGCTCGCGCTCTCGGAGCTTTTCCAAGAGCAACGGGAACACC

GTCCTGCTCGGGCAGAACGGCGCCGGCGCCGGGTACAGGATCCCGTTCGCGGCGCTCCAG

GAGGCGACCGGCGGGTTCGACGAGGGGATGGTCATCGGCGAGGGCGGGTTCGGGAAGGTG

TACAAGGGCACGATGCGGGACGAGACGCTGGTGGCCGTGAAGCGCGGCAACCGGCGGACG

CAGCAGGGGCTGCACGAGTTCCACACGGAGATCGAGATGCTGTCCCGGCTGCGCCACCGG

CACCTGGTCTCGCTCATCGGCTACTGCGACGAGCGCGGCGAGATGATCCTCGTGTACGAG

TACATGGCCATGGGCACGCTGCGGAGCCACCTGTACGGCGCCGGCCTCCCGCCCCTGTCG

TGGGAGCAGAGGCTGGAGGCCTGCATCGGCGCCGCGCGGGGGCTGCACTATCTCCACACC

AGCTCCGCCAAGGCGATCATCCACCGGGACGTCAAGTCGGCCAACATCCTCCTCGACGAG

AGCTTCATGGCCAAGGTGGCCGACTTCGGGCTGTCCAAGAACGGGCCGGAGCTGGACGAG

ACGCACGTGAGCACCAAGGTGAAGGGCAGCTTCGGGTACCTGGACCCGGAGTACTTCCGG

CGGCAGATGCTGACGGAGAAGTCGGACGTCTACTCCTTCGGCGTGGTCCTGCTGGAGGTG

CTCTGCGCCCGCACCGTCATCGACCCCACGCTGCCGCGGGAGATGGTGAGCCTGGCAGAG

TGGGCGACGCCGTGTCTCAGAAACGGCCAGCTCGACCAGATCGTCGACCAGAGGATCGCC

GGGACGATACGGCCGGGGTCGCTCAAGAAGCTCGCGGACACGGCCGACAAGTGCCTCGCC

GAGTACGGGGTGGAGCGGCCCACCATGGGGGACGTGCTGTGGTGCCTCGAGTTCGCGCTG

CAGCTGCAGGTGGCGTCCTCAGACGTCTCGGACGCCGACACCATGTTGACGCCGCCCGTG

CCCGTGAAAACGCCCGAGGTTCAGCGTAGCCTGTCCGCCGCTACCGTGGCGACTGACGCT

GCCATGACCACCAACTTGGGTGATCTAGACGGAATGTCCCTGAGCGGAGTATTCTCCAAG

ATGATCAAGAGCGACGAGGTCAGGTGA

>Ta-CrRLK1L8-D CDS sequence

ATGCCGCCGGTTCTTGACATGCTCGTGCGGCTCCTCGTCGCGTCCGTGCTGCTCGGCGCA

GCCAGTGGCGCGTTTACCCCCGCGGACAACTACCTCGTCCTCTGCGGCACGTCGGCGAGC

GCCACCGTCGCCGCCGGACGGACGTTCGTCGGGGACGCGCGTCTGCCGGCCAAGTCGCTG

GCCGCGCCGCAGAGCGTCGAGGCCAACACATCGCGGACCGCGGTCGTCCCGTCCGGCGAG

TCGGAGCTGTATCGGTCCGCACGCGTGTTCACCGCGCCAGCTTCCTACACGTTCGCCGTC

AAGCAGCCCGGCCGGCACTTCGTGCGCCTTCACTTCTTCCCCTTCCCCTACAGGTCTTAC

GACATGGTGGCGGACGCCGCGTTCAACGTGTCCGTGCAGGGCGCGGTGCTCGTCAACGGG

TACACGCCCAAGAACGGCACAGCGGAGCTCAGGGAGTTCTCCGTGAACGTCACCGGGGGC

ACGCTGGTGATCGCGTTCGCGCCGACGGGGAAGCTCGCGTTCGTGAACGCCATCGAGGTC

GTGTCGGTCCCCGACGAGCTCATCGCCGACATGGCCAGGATGGTGGACGGCGCCGTCCAG

TACACCGGGCTGTCGACGCAGGCGCTGGAGACGATCCACAGGATCAACATGGGCGTTCCC

AAGATCACGCCCGGCAACGACACGCTGGGGAGGACGTGGTTGCCGGACCAGAGCTTCCAG

GTCAACACCGACCTAGCGCAGCACAAAGACGCCAAGCCCCTGACGATCAAATACGACGAG

AAATCGGCACTCTCGTCCGCGTACACGGCGCCGGCGGAGGTCTACGCGACGGCGACGAGG

CTGAGCACGGCGGGCGAGACCAGCACCATCAACGTGCAGTTCAACATCAGCTGGAGGTTC

GACGCCCCGGCCGGGTCGGATTACCTGCTCCGGTTCCACTGGTGCGACATCGTCAGCAAG

GCAGCCATGGGAATGGCCTTCAACGTCTACGTCGGCGGGGCGGTGGTGCTCGACAACTAC

GAGATTTCGCGTGACACGTTCAACCGGCTATCCATACCGGTGTACAAGGACTTCCTCCTG

GGCGCCAAGGACGCCAAGGGCGCCATCACCGTGAGCATCGGGTCGTCCACCGAGGACAAC

GCGTTGCCTGACGGCTTCCTCAACGGCCTCGAGATCATGAGTATAGTCGGGAGCGCCGGC

GCCGGCGCTGCCGCCACGTCCCCGCGCAGTTCAAAGGTCAAAATCGGGATCATCGCCGGC

TCGGCCGTCTGCGGGGCCACGCTGGTGATGGTGCTCGGGTTCATCGCCTTCAAGATGCTG

CGCGGGAGGGAGCCGGAGAAGAAGAAGCCGGCCGACGCCTGGTCGCCGTTCTCGGCGAGC

GCGCTGGGCTCTCGCTCGCGCTCCCGGAGCTTCTCCAAGAGCAACGGGAACACCGTCCTG

CTCGGGCAGAACGGCGCCGGCGCCGGGTACAGGATCCCGTTCGCGGCGCTCCAGGAGGCG

ACCGGCGGGTTCGACGAGGGGATGGTCATCGGCGAGGGCGGGTTCGGGAAGGTGTACAAG

GGCACGATGCGCGACGAGACGGTGGTGGCCGTGAAGCGCGGCAACCGGCGGACGCAGCAG

GGGCTGCACGAGTTCCACACGGAGATCGAGATGCTGTCCCGGCTGCGCCACCGGCACCTG

GTCTCGCTCATCGGCTACTGCGACGAGCGCGGCGAGATGATCCTCGTGTACGAGTACATG

GCCATGGGCACGCTGCGGAGCCACCTCTACGGCGCCGGCCTCCCGCCCCTGTCGTGGGAG

CAGAGGCTGGAGGCCTGCATCGGCGCCGCGCGGGGGCTGCACTACCTCCACACCGGCTCC

GCCAAGGCCATCATCCACCGGGACGTCAAGTCGGCCAACATCCTCCTCGACGAGAGCTTC

ATGGCCAAGGTGGCCGACTTCGGGCTGTCCAAGAACGGGCCGGAGCTGGACAAGACGCAC

GTGAGCACCAAGGTGAAGGGCAGCTTCGGGTACCTGGACCCGGAGTACTTCCGGCGGCAG

ATGCTGACGGAGAAGTCGGACGTTTACTCCTTCGGCGTGGTCCTGCTGGAGGCGCTCTGC

GCCCGCACCGTCATCGACCCGACGCTGCCGCGGGAGATGGTGAGCCTGGCGGAGTGGGCG

ACGCCGTGTCTCAGAAACGGCCAGCTCGACCAGATCGTCGACCAGAGGATCGCCGGGACG

ATACGGCCGGGGTCGCTCAAGAAGCTCGCGGACACGGCCGAGAAGTGCCTCGCCGAGTAC

GGGGTGGAGCGGCCCACCATGGGGGACGTGCTCTGGTGCCTCGAGTTCGCGCTGCAGCTG

CAGGTGGGGTCGTCAGACAGCTCGGACGTCGACACCATGTTGCCGCCGGCGCCGCCCGTG

CCCGTGAAAACGCCTGAGGTTCAGCGTCGCCTGTCCGCCGCTACCGTGGCGACTGACGCT

GCTGCCATGACCACCAACTTGGGTGACCTAGACGGAATGTCCCTGAGCGGAGTATTCTCG

AACATGATCAAGAGCGACGAGGTCAGGTGA

>Ta-CrRLK1L9-A CDS sequence

ATGGGAGGAGGCCCGAGATGCGCGCTCCTGCTGCTCGTGGCCGCCGCGGCCGCCGCGCTT

GTCCCCGCGGCGTGGGCGCAGGACCCGACCGCGCCGGCGCCCTCGGGGGCCCCCTTCGTG

CCGCGGGACGACATCCTGCTCGACTGCGGCGCCACGGGGAAGGGCAACGACACGGACGGC

CGGGAGTGGCGTGGCGACGCCGGCTCCAAGTACGCGCCGCCGAACCTCGCCTCCGCCGAC

GCGGGGGCGCAGGACCCCTCGGTGCCGCAGGTGCCCTACCTCACCGCGCGGGTCTCCGCG

GCGCCCTTCACCTACTCCTTCCCGCTCGGCCCCGGCCGCAAGTTCCTCCGCCTGCACTTC

TACCCGGCCAACTACTCCGGCCGCGCCGCCGCCGACGCCTTCTTCTCCGTCTCCGTCCCG

GCAGCCAAGGTCACGCTCCTCTCCAACTTCAGCGCCTACCAGACCGCCACGGCGTTCAAC

TTCGCCTACCTCGTGCGCGAGTTCTCCGTCAACGTCACCGGCCCGACCCTCGACCTCACC

TTCACCCCGGAGAAGGGGCGCCCCAACGCCTACGCCTTCATCAACGGCATCGAGGTCGTC

TCCTCCCCGGATCTCTTCGACCTCGCCACACCGTTCTTCGTCACCGGTGACGGCAACAAC

CAGCCGTTCCCGATGGACCCCGGTGCTGCTCTGCAGACCATGTACCGGCTCAACGTCGGA

GGCCAGGCCATCTCCCCTTCCAAGGACTCCGGCGGGGCTCGGTCATGGGACGACGACACG

CCCTACATTTACGGTGCAGGGGCTGGGGTGTCGTACCCGAACGATCCCAATGTCACAATC

ACCTACCCTCCCAGTGTGCCGGGATATGTGGCGCCACTGGATGTCTATGCCACGGCGCGA

TCAATGGGGATAGACAAGGGTGTGAACTTGGCCTACAATCTCACCTGGATAGTGCAGGTG

GATGCTGGGTTCACATACCTTGTGAGGCTCCATTTCTGCGAGATACAATCCCCAATTGAT

AAGCCGAATCAGCGGGTGTTCAACATTTACCTCAACAACCAGACTGCCGTGGAAGGTGCT

GATGTGCTCCAGTGGGTGGATCCGCGTAGTACCGGTACCCCATTGTACAAGGATTTCGTG

GTCGGCACTGTGGGTTCAGGGATTATGGATTTCTGGGTGGCTCTTCATCCGGATATACGG

AACAAGCCACAGTACTATGATGCTATTCTCAATGGGATGGAGGTGTTCAAGCTGCAACTT

ACTAATGGGAGCCTCGCGGGGCCAAACCCTGTCCCAAGTGCTGATCCAGCGGCGCATACC

GGGCAAGGGAAGAAGAGTTCGCTTGTCGGGCCTATTGCTGGTGGAGTAATTGGAGGTTTG

GCAGTGCTTGCACTTGGATATTGCTGCTTCATTTGCAAGAGGCGGAGGAAAGTGGCCAAG

GATGCCGGCATGAGTGATGGCCATTCTGGTTGGCTGCCGTTGTCGCTGTATGGCAATTCA

CACACTTCAAGCTCAGCCAAGTCGCATGCTACAGGGAGTATTGCTTCATCTTTGCCATCC

AACCTGTGCCGCCATTTCTCCTTTGCAGAGATCAAGGCTGCAACGAAGAACTTTGACGAG

TCACGGATCCTTGGTGTTGGTGGGTTCGGTAAAGTTTACCAGGGAGAGATCGATGGGGGC

ACAACTAAGGTGGCTATCAAGCGTGGCAACCCCTTGTCTGAGCAGGGCATACATGAGTTC

CAGACTGAAATTGAAATGCTGTCAAAGCTCCGCCACCGCCATCTTGTGTCGTTGATTGGT

TACTGCGAGGACAAGAATGAGATGATCCTGGTCTATGACCACATGGCCCATGGAACTCTG

CGTGAGCACCTATACAAGACCCAGAATGCGCCGCTTAGCTGGAGGCAGCGTTTGGAGATC

TGCATTGGTGCAGCCCGTGGGCTGCACTACCTTCACACCGGTGCAAAGCACACCATTATC

CACCGTGATGTGAAGACCACAAACATCCTCCTGGATGAGAAATGGGTCGCCAAGGTTTCA

GATTTTGGTCTGTCCAAGACTGGGCCGTCGATGGATCACACACATGTGAGCACAGTTGTC

AAGGGCAGTTTCGGTTACCTAGATCCTGAATACTTCCGCAGGCAGCAGCTCACCGAGAAG

TCTGATGTGTATTCATTTGGCGTGGTGCTGTTCGAGGTCCTCTGTGCTCGGCCTGCCTTG

AACCCCACTCTTGCAAAGGAAGAAGTCAGCCTGGCAGAATGGGCATTGCACTGCCAGAAG

AAGGGAATTCTGGATCAGATTGTTGATCCCTACCTAAAGGGAAAGATCGTTCCGCAGTGC

TTCAAGAAGTTTGCCGAGACAGCTGAGAAGTGCGTTGCCGACAACGGCATCGAGCGCCCT

TCGATGGGAGATGTGCTTTGGAACCTGGAGTTTGCTCTTCAGATGCAGGAAAGCGCGGAG

GAGAGCGGAAGCTTCGGCTGTGGGATGTCGGATGAGGAGGGCGCTCCCCTGGTGATGGCT

GGAAAGAAGGATCCCAATGACCCGTCAATCGATTCAAGCACCACCACGACCACGACAACT

TCCCTAAGCATGGGCGACCAGAGCGTCGCGAGCATCGACTCGGACGGGCTGACGCCGAGC

GCGGTCTTCTCGCAGATCATGAACCCCAAGGGGCGGTGA

>Ta-CrRLK1L9-B CDS sequence

ATGGGAGGAGGCCCGAGATGCGCGCTCCTGCTGCTCGCCGCGGCCGCCGCTTGCGCGGCG

CTTGTCCCCGCGGCGTGGGCGCAGGGCTCGACCGCGCCCGCGCCCTCGGGGGCCCCCTTC

GTGCCGCGGGACGACATCCTGCTCGACTGCGGCGCCACGGGGAAGGGCAACGACACGGAC

GGCCGGGAGTGGCGCGGCGACGCCAGCTCCAAGTACGCGCCGCCGAACCTCGCCTCGGCC

GACGCGGGGGCGCAGGACCCCTCGGTGCCGCAGGTGCCCTACCTCACTGCGCGGGTCTCC

GCGGCGGCCTTCACCTACTCCTTCCCGCTCGGCCCCGGCCGCAAGTTCCTCCGGCTGCAC

TTCTACCCGGCCAACTACTCCAACCGCGACGCGGCCGACGCCTTCTTCTCCGTCTCCGTC

CCGGCCGCCAAGGTCACGCTCCTCTCCAACTTCAGCGCCTACCAGACCGCCACGGCCCTC

AACTTCGCCTACCTCGTACGGGAGTTCTCCGTCAACGTCACTGGCCCGACCCTCGACCTC

ACCTTCACCCCGGAGAAGGGACGCCCCAACGCCTACGCCTTCATCAACGGCATCGAGGTC

GTCTCCTCTCCGGATCTCTTTGACCTCGCCACACCGTTCTTCGTCACCGGTGACGGCAAC

AACCAGCCATTCCCGATGGATCCCGGTGCTGCTCTGCAGACCATGTATCGGCTCAACGTC

GGAGGCCAGGCGATCTCCCCTTCCAAGGACTCCGGCGGGGCTCGGTCATGGGACGACGAC

ACGCCTTACATTTATGGTGCAGGGGCTGGGGTAACGTACCCGAACGATCCCAATGTCACA

ATCACCTATCCTGACAGTGTGCCGGGATATATGGCGCCTTCGGATGTCTATGCCACGGCC

CGATCAATGGGGATAGACAAGAATGTGAACTTGGCCTACAATCTCACCTGGATAGTGCAG

GTGGATGCTGGGTTCACATACCTTGTGAGGCTCCATTTCTGTGAGATACAATCCCCAATT

GATAAGCCAAATCAGCGGGTGTTCAACATTTACCTCAACAACCAGACTGCCGTGGAAGGT

GCTGATGTGATCCAGTGGGTGGATCCGCTTAGTACTGGCACCCCATTGTATAAGGATTAT

GTGGTCGGCACTGTGGGTTCAGGGATTATGGATTTCTGGGTGGCTCTTCATCCGGATATA

CGGAACAAGCCACAGTACTATGATGCTATTCTCAATGGGATGGAGGTGTTCAAGCTGCAA

CTTAGTAATGGGAGCCTCGCGGGGCCAAACCCTGTCCCAAGTGCTGATCCACAGGCGCAT

ACCGGGCAAGGGAAGAAGAAATCACTAGTCGGGCCTATTGCTGGTGGAGTAATTGGAGGT

TTGGCAGTGCTTGCACTTGGATATTGCTGCTTCATTTGCAAGAGGCGGAGGAAAGCGGCC

AAGGATACTGGCATGAGTGATGGCCATTCTGGTTGGCTGCCGTTGTCGCTGTATGGCAAT

TCACACACTTCAAGCTCAGCCAAGTCGCATGCTACAGGGAGTATTGCTTCATCTTTGCCA

TCGAACCTGTGTCGCCATTTCAGCTTTGCAGAGATCAAGGCTGCAACGAAAAACTTTGAC

GAGTCACGGATCCTTGGTGTTGGTGGGTTCGGTAAAGTTTACCAGGGCGAGATCGATGGG

GGCACAACTAAGGTGGCTATCAAGCGTGGCAATCCCTTGTCTGAGCAGGGTATACATGAG

TTCCAGACTGAAATTGAAATGCTGTCAAAGCTCCGCCACCGCCATCTTGTGTCGCTGATT

GGTTACTGCGAGGACAAGAATGAGATGATCCTGGTCTATGACCACATGGCTCATGGAACT

CTGCGTGAGCACCTATACAAGACCCAGAATGCACCGCTTAGCTGGAGGCAGCGTTTGGAG

ATCTGCATTGGTGCAGCTCGTGGGCTGCACTACCTTCACACCGGTGCAAAGCACACCATT

ATCCACCGTGATGTGAAGACAACAAACATCCTACTGGATGAGAAATGGGTCGCCAAGGTT

TCAGATTTTGGTCTGTCCAAGACTGGGCCATCGATGGATCATACACATGTGAGCACAGTT

GTCAAGGGCAGTTTTGGTTACCTAGATCCTGAATACTTCCGCAGGCAGCAGCTCACCGAG

AAATCTGATGTCTATTCGTTTGGTGTGGTGCTGTTCGAGGTCCTCTGTGCTCGGCCTGCC

TTGAACCCGACTCTTGCAAAGGAAGAAGTTAGCCTGGCAGAATGGGCATTGCACTGCCAG

AAGAAGGGAATTCTGGATCAGATTGTTGATCCCTACCTAAAGGGAAAGATTGTTCCGCAG

TGCTTCAAGAAGTTTGCCGAGACAGCTGAGAAGTGCGTTGCCGACAATGGCATCGAGCGC

CCTTCGATGGGAGATGTGCTTTGGAATTTGGAGTTTGCTCTTCAGATGCAGGAAAGTGCG

GAGGAGAGCGGAAGCTTTGGCTGTGGGATGTCGGATGAGGGCACTCCCCTGGTGATGGCT

GGAAAGAAGGATCCCAATGACCCATCGATCGATTCAAGCACCACCACGACCACGACAACT

TCCCTAAGCATGGGCGACCAAAGTGTCGCGAGCATCGACTCGGACGGGCTGACGCCGAGC

GCCGTGTTCTCACAGATCATGAACCCCAAGGGGCGGTGA

>Ta-CrRLK1L9-D CDS sequence

ATGGGAGGAGGCCCGAGATGCGCGCTCCTGCTGCTCGCCGCGGCCGCCGCCTGCGCGGCG

CTTGTCCCGGCGGCGTGGGCGCAGGCCCCGACAGCGCCGGCGCCCTCGGGGGCTCCCTTC

GTGCCGCGGGACGACATCCTGCTCGACTGCGGCGCCACGGGGAAGGGGAACGACACAGAC

GGCCGGGAGTGGCGCGGCGACGCCGGCTCCAAGTACGCGCCGCCGAACCTCGCCTCGGCC

GACGCGGGGGCGCAGGACCCCTCGGTGCCTCAGGTGCCCTACCTCACCGCGCGGGTCTCC

GCGGCGGCCTTCACCTACTCCTTCCCGCTCGGCCCCGGCCGCAAGTTCCTCCGGCTGCAC

TTCTACCCGGCCAACTACTCCAACCGCGACGCCGCCGACGCCTTCTTCTCCGTCTCCGTC

CCGGCCGCCAAGGTCACGCTCCTCTCCAACTTCAGCGCCTACCAGACCGCCACGGCGCTC

AACTTCGCCTACCTCGTGCGCGAGTTCTCCGTCAACGTCACCGGCCCGACCCTCGACCTC

ACCTTCACCCCGGAGAAGGGACGCCCCAACGCCTACGCCTTCATCAATGGCATCGAGGTC

GTCTCCTCCCCAGATCTCTTTGACCTCGCCACACCGCTCTTCGTCACCGGTGACGGCAAC

AACCAGCCATTCCCGATGGATCCTGGTGCTGCTCTGCAGACCATGTATCGGCTCAACGTC

GGAGGCCAGGCGATCTCCCCTTCCAAGGACTCCGGCGGGGCTCGGTCATGGGACGACGAC

ACGCCTTACATTTATGGTGCAGGGGCTGGGGTAACGTACCCGAACGATCCCAATGTCACA

ATCACCTACCCTGACAATGTGCCGGGATATGTGGCGCCTTCGGATGTCTATGCCACGGCG

CGATCAATGGGGATAGACAAGAATGTGAACTTGGCCTACAATCTCACCTGGATAGTGCAG

GTGGATGCTGGGTTCACATACCTTGTGAGGCTCCATTTCTGTGAGATACAATCCCCAATT

ACTAAGCCGAATCAGCGGGTGTTCAACATTTACCTCAACAACCAGACTGCCGTGGAAGGT

GCTGATGTGATCCAGTGGGTGGATCCGCTTAGTACTGGCACCCCATTGTATAAGGATTAT

GTGGTCAGCACTGTGGGTTCAGGGATTATGGATTTCTGGGTGGCTCTACATCCGAATACA

GGGAGCAAGCCACAGTACTATGATGCTATTCTCAATGGGATGGAGGTGTTCAAGCTGCAG

CTTAGTAATGGGAGCCTTGCGGGGCCAAACCCTGTCCCCAGTGCTGATCCACCGGCGCAT

ACCGGGCAAGAGAAGAAGAATTCACTAGTCGGGCCTATTGCCGGTGGAGTAATTGGAGGT

TTGGTAGTGCTTGCACTTGGATATTGCTGCTTCATTTGCAAGAGGCGGAGGAAAGTGGCG

AAGGATGCCGGCATGAGTGATGGCCATTCTGGTTGGCTGCCGTTGTCGCTGTATGGCAAT

TCACACACTTCAAGCTCAGCCAAGTCGCATGCTACAGGGAGTATTGCTTCATCTTTGCCA

TCCAACCTGTGTCGCCATTTCTCGTTTGCAGAGATCAAGGCTGCAACGAAAAACTTTGAC

GAGTCACGGATCCTTGGTGTTGGTGGGTTCGGCAAAGTTTACCAGGGAGAGATCGATGGG

GGCACAACTAAGGTGGCTATCAAGCGTGGCAATCCCTTGTCTGAGCAGGGTATACATGAG

TTCCAGACTGAAATTGAGATGCTGTCAAAGCTCCGCCACCGCCATCTTGTGTCGCTGATT

GGTTACTGCGAGGACAAGAATGAGATGATCCTGGTCTATGACCACATGGCCCATGGAACT

CTGCGTGAGCACCTATACAAGACCCAGAATGCACCGCTTAGCTGGAGGCAGCGCTTGGAG

ATCTGCATTGGTGCAGCTCGTGGGCTGCACTACCTTCACACCGGTGCAAAGCACACCATC

ATCCACCGTGATGTGAAGACGACAAACATACTCCTGGATGAGAAATGGGTCGCCAAGGTT

TCAGATTTTGGTCTGTCCAAGACTGGGCCGTCGATGGATCACACACATGTGAGCACAGTT

GTCAAGGGCAGTTTTGGATACCTAGATCCTGAATACTTCCGCAGGCAGCAGCTCACCGAG

AAATCTGATGTCTATTCATTTGGTGTGGTGCTGTTCGAGGTCCTCTGTGCTCGGCCTGCC

TTGAACCCCACTCTTGCAAAGGAAGAAGTCAGCCTGGCAGAATGGGCATTGCACTGCCAG

AAGAAGGGAATTCTGGATCAGATTGTTGATCCCTACCTAAAGGGAAAGATTGTTCCTCAG

TGCTTCAAGAAGTTTGCCGAGACAGCTGAGAAGTGTGTTGCCGACAATGGCATCGAGCGC

CCTTCGATGGGAGATGTGCTTTGGAACCTGGAGTTTGCTCTTCAGATGCAGGAAAGCGCG

GAGGAGAGCGGAAGCTTCGGCTGTGGGATCTCGGATGAGGAGGGCACTCCCCTCGTGATG

GCTGGAAAGAAGGATCCCAATGACCCGTCGATCGATTCAAGCACCACCACGACCACGACA

ACTTCCCTAAGCATGGGCGACCAAAGCGTCGCGAGCATCGACTCGGACGGGCTGACGCCG

AGCGCCGTGTTCTCGCAGATCATGAACCCCAAGGGGCGGTGA

>Ta-CrRLK1L10-A CDS sequence

ATGCTCCGGACGAGGATACTCGTGCTGGCTGCTGTGAGCATCGTGTTCGCCAACCTGCAG

TTCTTGAAGGCTCACGGGAGGGAGCTGTTTCTGAGCTGCGGCTCCAACGCCACCGCCGAT

GCCGATGGCCGGAGATGGATCGGCGACATGGCCCCTGACCTGAATTTCACTCTGAGCAGC

CCGGGGATCGCTGCTCTGCTGGCCGGGAGCAGCAATGGCAGTGAAATCATGGCGCCGGTG

TACCGCTCCGCGCGCTTCTTCACCACGACATCTTGGTATGACTTCAGCCTGCTGCCGGGG

AACTACTGCGTCAGGCTGCATTTCTTCCCGTCCGCATTCAGGAATTTCAGTGCAAACGGT

TCAGTGTTTGATGTCGTTGCCAATGACTTCAAGCTGGTGTCCAAGTTTAACGTGTCGGAG

GAGATAGTTTGGAGAAACTCAGTGAGCAATTCGGCCGCCACTGCGGTTGTCAAGGAGTAC

TTTCTTGCAGTCAATGGTTCTCGCCTGCAGATCGAGTTTGATCCAAGGCCCGGTTCATTT

GCATTTGTGAATGCGATCGAGGTGATGCTCACTCCAGATAATTCCTTCAACGGCATGGTG

AACAAAGTTGGTGGTGTGGATGTGCACATTCCTCCTGAATTAAGCGGCCGAGCTGTTGAG

ACTATGTATCGACTGAATATTGGAGGGCCTGCACTTGCATCTTCACATGATCAGTATCTT

CATAGACCATGGTACACTGATGAAGCATTCATGTTTTCTGCCAATGCTGCTTTGATTGTG

TCCAATACTTCAGCCATAAAGTATGTCTCAAGCAATGACTCCTCAATTGCTCCCCTTGAT

GTCTATGAGACCGCGAGAATCATGGGCAACAACATGGTCATGGACAAGAGGTTTAATGTG

ACATGGCGGTTCTTTGTCCACCCCAATTTTGATTACTTGGTCCGCCTTCATTTTTGCGAG

CTTGTCTATGACAAGCCCAGCCAGAGGATCTTCAAGATCTACATCAACAACAAGACAGCT

GCTGAGAACTACGATGTGTACAACAAAGCCGGAGGTATTAACAAGGCATATCATGAGGAC

TACTTTGATAGTTTGCCGCAGCAGGTAGACTCGCTCTGGCTTCAGCTAGGCCCGGACTCC

ATGACCAGTGCTTCAGGTACGGATGCACTTCTCAATGGTTTGGAGATATTCAAGCTCAGC

AGGAGTGGCAACCTTGATTATGTGCTTGGTCATATTGATATGGGCAACAAAAGGGGGCGT

TCCAAGGGTCGGAGCAGGATAGGTTTATGGGAAGAAGTTGGTATTGGCTCGGCCGCTTTT

GTGGCCCTGGTAAGTGTTGCTCTATTCTCATGGTGCTATGTAAGGAGGAAACGAAAAGCT

GTTAACGAGGAGGTCCCTGCTGGTTGGCACCCTCTGGTCCTCCATGAGGCTATGAAAAGC

ACTACAGATGCCCGCGCATCCAAAAAAGCACCCTTGGCACGCAATTCATCTTCCATTGGT

CATAGGATGGGCAGGCGATTCAGCATTGCAGATATTAGAGCTGCCACAAAAAACTTTGAC

GAGTCATTGGTCATTGGTTCTGGAGGTTTTGGCAAGGTTTACAAGGGTGAGGTCGATGAT

GGCATTACAGTCGCAATCAAGCGTGCAAATCCATTATGTGGTCAGGGCCTGAAAGAATTT

GAAACAGAGATCGAGATGCTCTCCAAGCTTAGGCACCGGCACCTTGTTGCAATGATTGGC

TATTGTGAAGAGCAGAAGGAGATGATTCTGGTCTATGAATACATGGCCAAGGGGACATTG

CGAAGCCATCTCTATGGAAGTGGCCTACCACCTTTGACATGGAAGCAACGGATTGATGCC

TGCATTGGTGCGGCCAGGGGCCTTCACTACCTCCACACGGGAGCAGATCGCGGTATAATT

CATAGGGATGTTAAGACTACTAACATCCTGTTGGACAAGAACTTTGTTGCAAAAATAGCA

GATTTTGGGTTGTCGAAAACTGGACCAACACTGGACCAGACCCATGTTAGTACAGCAATC

AGGGGTAGCTTCGGGTATCTTGATCCAGAGTACTTCCGGAGGCAGCAATTGACACAAAAA

TCTGATGTGTATTCTTTTGGTGTGGTTCTCTTTGAAGTTGCTTGTGCCAGGCCGGTTATA

GACCCTTCAGTGCCGAAGGATCAAATCAACTTGGCAGAATGGGCTATGAGATGGCAGCGT

CAGCGTTCGCTGGAAGCAATAGCGGATCCACGGCTGGATGGTGACTACTCGCCAGAATCC

TTGAAGAAGTTTGGTGATATCGCGGAGAAGTGTCTTGCTGATGATGGGAGAACCAGGCCA

TCAATGGGTGAGGTCTTGTGGCACCTGGAGTATGTGCTGCAGCTCCATGAAGCTTACAAA

CGCAACGTGGATTGCGAGTCATTTGGAAGCAGTGAACTGGGGTTCGCTGATATGTCTTTT

AGCATGCCTCACATCAGGGAGGGAGAAGAGGAGCATCACCCAAAGAAATCAGGTATCAGA

GAAGATTCAGCCCCTTGA

>Ta-CrRLK1L10-B CDS sequence

ATGCTCCGGATGAGGATACTCGTGCTGGCTTCTGTGAGCATCGTGTTCGCCAACCTGCAG

TTCTTGAAGGCTCATGGGAGGGAGCTGTTTCTGAGCTGCGGCTCCAACGCCACCGCCGAT

GCCGATGGCCGGAGATGGATCGGCGACATGGCCCCTGACCTGAATTTCACTCTGAGCAGC

CCGGGGATTGCTGCTCTCTTGGCCGGGAGCACCAATGGGAGTGAAATCATGGCACCGGTG

TACCGCTCGGCGCGCTTCTTTACCACGACATCTTGGTATGACATCAGCGTGCTGCCGGGG

AACTACTGTGTCAGGCTGCATTTCTTCCCGTCCGCATTCGGGAATTTCAGTGCAAATGGT

TCAGTGTTTGATGTCGTCGCCAATGAGTTCAAGCTGGTGTCGAAGTTTAACGTTTCGGAG

GAGATTGTTTGGAGAAATTCAGTGAGCAATTCAGCTGCCACTGCGGTTGTCAAGGAGTAC

TTTCTTGCAGTCAATAGTTCTCGCCTGCAGATCGAGTTTGATCCAAGGCCCGGTTCATTT

GCATTTGTTAATGCGATCGAGGTGATGCTCACTCCAGATAATTCCTTCAACAGCACAGTG

AACAAAGTTGGTGGTGTGGATGTGCACATTCCTCCTGAATTAAGCGGCCGAGCTATTGAG

ACCATGTATAGGCTCAACATTGGAGGGCCTGCACTTGCATCTTCACATGATCAGTATCTT

TATAGACCATGGTACACTGATGAAGCCTTCATGTTTTCTGCGAATGCTGCTTTGACCGTG

TCCAATACGTCAGCCATAAAGTATGTCTCAAGCGGCGACTCCTCAATTGCTCCCATCGGT

GTCTATGAGACTGCAAGAATCATGGGCAACAACATGGTCATGGACAAACGGTTCAATGTG

ACATGGCGGTTCGTTGTCCATCCCAATTTCGATTACATGGTCCGTCTTCATTTTTGCGAG

CTTGTCTATGACAAGCCCAGCCAGAGGATCTTCAAGATCTACATCAACAACAAGACAGCT

GCTGAGAACTACGATGTGTATGACAAGGCTGGAGGAATTAACAAGGCATATCATGAGGAC

TACTTTGACAGCTTGCCGCAACAGGTAGACTCACTCTGGCTTCAGCTAGGCCCGGACTCC

ATGACCAGTGCTTCAGGCACGGATGCACTTCTCAATGGTTTGGAGATATTCAAGATCAGC

AGGAGTGGCAACCTTGACTATGTGCTTGGTCATATTGATATGGGCAACAAAAGGGGCCGT

TCCAAGGGTCGGAGCAGGTTAGGTTTATGGGAAGAAGTTGGTATTGGCTCAGCCGCTTTT

GTGGCACTGGCAAGTGTTGCTCTATTCTCATGGTGCTATGTAAGGAGGAAACGGAAAGCT

GTCGACGAGGAGGTCCCTGCTGGTTGGCACCCTCTGGTCCTTCATGAGGCTATGAAAAGC

ACTACAGATGCCCGCGCATCCAAAAAAGCACCATTGGCACGCAATTCATCTTCCATTGGT

CATAGGATGGGCAGACGATTCAGCATTGTAGATATTAGGGCTGCCACAAAGAACTTTGAC

GAGTCATTGGTCATTGGTTCTGGAGGTTTTGGCAAGGTTTACAAGGGTGAGGTCGATGAT

GGCATTACAGTTGCAATCAAGCGTGCAAATCCATTATGTGGCCAGGGCCTGAAAGAATTT

GAAACAGAGATCGAGATGCTCTCCAAGCTTAGGCACCGGCACCTTGTTGCGATGATTGGC

TATTGTGAAGAGCAGAAGGAGATGATTCTGGTCTATGAATACATGGCCAAGGGGACATTG

CGAAGCCATCTCTATGGAAGTGGCCTACCACCTTTGACATGGAAGCAACGGATTGATGCC

TGCATTGGTGCGGCCAGGGGCCTTCACTACCTCCACACAGGAGCAGACCGAGGTATAATT

CATAGGGATGTTAAGACTACTAACATCCTGTTGGACAAGAACTTTGTTGCAAAAATAGCA

GATTTTGGGTTGTCGAAAACTGGACCAACACTGGACCAGACCCATGTTAGTACAGCAATC

AGGGGTAGCTTCGGGTATCTTGATCCAGAGTACTTCCGGAGGCAGCAATTGACACAAAAA

TCCGACGTGTATTCTTTTGGTGTGGTGCTCTTTGAAGTTGCTTGTGCCAGGCCGGTTATA

GACCCTTCAGTGCCGAAGGATCAAATCAACTTGGCAGAATGGGCTATGCGATGGCAGCGT

CAGCGTTCGCTGGAAGCAATAGCGGATCCACGGCTGGATGGTGACTACTCGCCAGAATCC

TTGAAGAAGTTTGGTGATATCGCAGAGAAGTGTCTTGCTGATGATGGGAGAACCAGGCCA

TCAATGGGTGAGGTTTTGTGGCACCTGGAGTATGTGTTGCAGCTCCATGAAGCTTACAAA

CGCAACGTGGATTGCGAGTCGTTTGGAAGCAGTGAACTGGGGTTCGCTGATATGTCTTTT

AGCATGCCTCACATCAGAGAAGGAGAAGAGGAGCATCACCCAAAGAAATCTGGTATCAGA

GAAGATTCAGCCCCTTAA

>Ta-CrRLK1L10-D CDS sequence

ATGCTCCGGATGAGGATACTCGTGCTGGCCGCTGTGAGCATCGTGTTCGCCAACCTGCAG

TTCTTGAAGGCTCACGGGAGGGAGCTGTTTCTGAGCTGCGGCTCCAACGCCACCGCCGAT

GCCGATGGCCGGAGATGGATCGGCGACATGGCCCCTGGCCTGAATTTCACTCTGAGCAGC

CCGGGAATCGCTGCTCTGCTGGCCGGGAGCAGCAATGGCAGTGAAATCATGGCGCCGGTG

TACCGCTCCGCGCGCTTCTTTACCACCACATCTTGGTATGACTTCAGCCTGCTGCCGGGG

AACTACTGCGTCAGGCTGCATTTCTTCCCGTCCACATTCAGGAATTTCAGTGCAAACGGT

TCAGTGTTTGATGTCGTCGCCAATGACTTCAAGCTGGTGTCCAAGTTTAACGTGTCGGAG

GAGATTGTTTGGAGAAACTCAGTGAGCAATTCAGCTGCCACTGCGGTTGTCAAGGAGTAC

TTTCTTGCAGTCAATAGTTCTCGCCTGCAGATCGAGTTTGATCCAAGGCCCGGTTCATTT

GCATTTGTGAATGCGATCGAGGTGATGCTCACTCCAGATAATTCCTTCAACGGCACGGTG

AACAAAGTTGGTGGTGTGGATGCACACATTCCTCCTGAATTAAGCGGCCGAGCTGTCGAG

ACCATGTATCGGCTGAACATTGGAGGGCCTGCACTTGCATCTTCACATGATCAGTATCTT

CATAGACCATGGTACACTGATGAAGCATTCATGTTTTCTGCCAATGTTGCTTTGATTGTG

TCCAATACTTCAGCCATAAAGTATGTCTCAAGCAACGACTCCTCAATTGCTCCCATCGAT

GTCTATGAGACCGCAAGAATCATGGGCAACAACATGGTCATGGACAAGCGGTTCAATGTG

ACATGGCGGTTCTTGGTCCACCCCAATTTTGATTACTTGGTCCGCCTTCATTTTTGTGAG

CTTGTCTATGACAAGCCCAGCCAGAGGATCTTCAAGATCTACATCAACAACAAGACAGCT

GCTGAGAACTATGATGTGTACAACAGGGCCGGAGGTATTAACAAGGCATATCATGAAGAC

TACTTTGATAGTTTGCCGCAGCAGGTAGACTCACTCTGGCTTCAGCTAGGCCCAGACTCC

ATGACCAGTGCTTCAGGTACCGATGCACTTCTCAATGGTTTGGAGATATTCAAGCTCAGC

AGGAGTGGCAACCTTGATTATGTGCTTGGTCATATTGATATGGGCAACAAAAGGGGGCGT

TCCAAGGGTCGGAGCAGGATAGGTTTATGGGAAGAAGTTGGTATTGGCTCGGCCGCTTTT

GTGGCACTGGCAAGTGTTGCTCTGTTCTCATGGTGCTATGTAAGGAGGAAACGAAAAGCT

GTTAACGAGGAGGTCCCTGCTGGTTGGCACCCTCTGGTCCTCCATGAGGCTATGAAAAGC

ACTACAGATGCCCGCGCATCCAAGAAAGCACCCTTGGCACGCAATTCATCTTCCATTGGT

CATAGGATGGGCAGACGATTCAGCATTGCAGATATTAGAGCTGCCACAAAAAACTTTGAC

GAGTCATTGGTCATTGGTTCTGGAGGTTTTGGCAAGGTTTACAAGGGTGAGGTCGATGAT

GGCATTACAGTCGCAATCAAGCGTGCAAATCCATTATGTGGTCAGGGACTGAAAGAATTT

GAAACAGAGATCGAGATGCTCTCCAAGCTTAGGCACCGGCACCTTGTTGCGATGATTGGC

TATTGTGAAGAGCAGAAGGAGATGATTCTGGTCTATGAATACATGGCCAAGGGGACATTG

CGAAGCCATCTCTATGGAAGTGGCCTACCACCTTTGACATGGAAGCAACGGATTGATGCC

TGCATTGGTGCGGCCAGGGGCCTTCACTACCTCCACACGGGAGCAGATCGGGGTATAATT

CATAGGGATGTTAAGACTACTAACATCCTGTTGGACAAGAACTTTGTTGCAAAAATAGCA

GATTTTGGGTTGTCGAAAACTGGACCAACACTGGACCAGACCCATGTTAGTACAGCAATC

AGGGGTAGCTTCGGGTATCTTGATCCAGAGTACTTCCGGAGGCAGCAATTGACACAAAAA

TCTGATGTGTATTCTTTTGGTGTGGTGCTCTTTGAAGTTGCTTGTGCCAGGCCGGTTATA

GACCCTTCAGTGCCGAAGGATCAAATCAACTTGGCAGAATGGGCTATGAGATGGCAGCGT

CAGCGTTCACTGGAAGCAATAGCGGATCCACGGCTGGATGGTGACTACTCGCCAGAATCC

TTGAAGAAGTTTGGTGATATCGCGGAGAAGTGTCTTGCTGATGATGGGAGAACCAGGCCA

TCAATGGGTGAGGTCTTGTGGCACCTGGAGTATGTGCTGCAGCTCCATGAAGCTTACAAA

CGCAATGTGGATTGCGAGTCATTTGGAAGCAGTGAACTGGGGTTTGCTGATATGTCTTTT

AGCATGCCTCACATCAGAGAGGGAGAAGAGGAGCATCACCCAAAGAAATCAGGTATCAGA

GAAGATTCAGCCCCTTGA

>Ta-CrRLK1L11-A CDS sequence

ATGGGCACCACTACCGAGCAGAAGATAGCTCTGCTCCTTCTTGGGACCATCTGGGTTCTT

CTTGGTACCTGCAATGCTGCTGAATTCACCCCTGCAGACAACTACCTCATCAACTGCGGC

TCTACGGTCGACGCTAATCTCAACGATGGGAGGGTCTTCAAAGCAGACAATTCCGGCTCG

ACTATATTGACATCACATCACAGCGTCCCCGCAAACACCTTGCCGGATGCAGTCATAAGT

TCTGACAATCCTGTGCTGTATAAAACTGCAAGGATATTCATTGTGCCGTCGTCCTACTCC

TTCAACATGAAGAGCCGTGGCCGGCATTTTGTCCGGCTACACTTCTTCGGTTTCAGATAC

CAGAGCTATGATCTTGCCGCGGCAAAGTTCAAAGTGTCCACCCAGCATGTTGTGTTACTT

GACAATTTCACTCCGCCGAGCAATTCCTCACTGTTGGTCAGGGAGTATTCGCTGAATATT

ACCGAGGATATGCTGATTCTCTCATTTGTGCCTCTTGGAAACAGCACATCTTTTATCAAT

GCTATTGAAGTAATATCTGTTCCTGATGATCTGATACAGGATTCAGCCCAAACTGTGAAT

CCCTCCGGCCAGTATCTTGGCCTTGCAACGCAGTCATTTCAGACATTCTACAGGATTAAT

GTAGGTGGACGGGAGGTGACCGTTGTCAATGACACACTCTCGCGTTCATGGGATACTGAC

CAGAACTTCTTCATAAATTCCACTACCACTGAACTGTTTGCTTACCAAGGGAAGCTGAAT

TATCAGAAAGGAGCTGCAACCAAGGAGGATGCACCGGACAGTGTGTACAACACTGCAAGG

CGGTTGGCTGTGCAAAATAGAACCAGCCCAGCGTCCAACATGACATGGCAATTTGATGTT

GACGGTCGTTCGAGCTATCTGATCCGGTTCCATTTTTGTGACATAGTGAGCAAGGCAGCA

TATTCCCTCTACTTTGATATTTATGTGGATGGCGGGTTAGCGTTAGAAAATCTTGACCTC

TCTGAAAAAGTTTTTGGTACCTTGGCTGTGCCATACTACACGGAATTTGTCTTGAAGTCA

AGCAATCCTTCTGGTAAGCTAAGTGTCGGCATTGGGCCTTCCAGCTTGAGCAATGTGGCA

CCAGATGGCATCTTGAACGGCCTGGAGATCATGAAGATGAACATCAGTACTGGCACTATT

TATGTTGTGTGGCCACCAGCAACGCCAAAAAGGAAACTGGCTATCATATTGGCCCCTGTT

CTCGGAGGTGTTGGTGCTGTTAGCATTGCTATTATTCTCTGCTTTGTCCTCAGAAGAAAG

AAGGAGAAGAAGCCGCGGCGGGCACCGACAAGTCGACCTTCAAGTTCTTGGTCACCACTT

ACCCTCAATGGCCTTAGCTTCCTTAGCATAGGTACCCGAACAACCAGCCGGACCACTCAT

ACATCTGGGACAAACAGTGATGTAAGCTACCGAATTCCTTTTGCTTTGCTGCAAGTGGCA

ACAAAACACTTCGACGAGCAGATGGTTGTCGGAGTTGGGGGGTTTGGGAAGGTATACAAA

GCAGTTCTGCAGGACAGCACCAAAGTCGCAGTCAAGCGGGGCAACCAGAAGTCCCACCAA

GGGCTCAAAGAATTCCGGACAGAGATCGAGCTGCTGTCGGGGCTGCGACACCGTCACCTC

GTGTCGCTCATCGGATATTGCGATGACCAGAACGAGATGATCTTGGTGTATGAGTACATG

GAGAAAGGCACGCTGAAGAGTCACCTGTATGGCAGTGACATGCCTCCCCTCAGCTGGAAG

AAAAGAGTGGAAATCTGCATAGGGGCTGCAAGGGGGCTCCACTACCTGCACACAGGTTTT

GCAAAGTCGATCATCCATCGTGATGTCAAGTCGGCAAACATTCTCCTCGACGAAAATCTC

ATGGCCAAGGTTTCTGATTTCGGTCTCTCGAAGACGGGACCTGAGTTGGATCAAACACAT

GTTAGCACTGCGGTGAAAGGGAGCTTCGGGTATCTTGACCCTGAGTACTACCGGAGGCAG

AAGCTGACCGACAAGTCGGATGTGTACTCATTCGGCGTGGTCTTGCTGGAGGTGATCTGT

GCGAGGCCAGTCATCGATCCGACGCTTCCAAGAGACATGATCAACCTTGCAGAATGGGCA

ATCAAGTGGCAGAAGAGGGGAGAGCTTGGTCAGATCGTCGATCAACGGATCGCTGGGACA

ATCAGGCCAGAGTCATTGAGGAAGTACGGTGAGACGGTCGAGAAGTGTCTTGTGGATTAC

GGCGTTGACCGCCCTACAATGGGTGATGTCCTGTGGAATCTGGAGTTTGTGCTCCAGCTG

CAGGAGGCCGGCCCGGACATCTCCAATGTCGACAGCATGAATCAGATCTCTGAACTTCCT

TCAGACGCCAGAAGGATGGGCTCTTTGGAGATCGGTACCGCAGACGAAGCAGACGAAGGC

CGCACGCACATGGATTACTCTCAGATGTCGACCAACGATGCCTTCTCGCAGCTGATGAAC

ACTGAAGGGAGGTGA

>Ta-CrRLK1L11-B CDS sequence

ATGGGCACCACTACCGAGCAAAAGATAGCTCTGCTCCTTCTTGGAACCATCTGGGTTCTT

CTTGGTACCTGCAATGCTGCTGAATTCACCCCTGCAGACAACTACCTCATCAACTGCGGC

TCCACGGTCGACGCTAATCTCCACGATGGGAGGGTCTTCAAAGCAGACAATTCCGGCTTG

ACTATATTGACATCACATCACAGCGTCCCCGCAAACACCTTGCCGGATGCAGTCATAAGT

TCTGACAATCCTGTGCTGTATCAAACTTCAAGGATATTCATTGTGCCGTCGTCCTACTCC

TTCAAGATGAAGAGCCGTGGCCGGCATTTTGTCCGGCTACACTTCTTCAGTTTCAGATAC

CAGAGCTATGATCTTGCCGCGGCAAAGTTCAAAGTGTCCACGCAGCATGTTGTGTTACTT

GACAATTTCACTCCGCCGAGCAATTCCTCACCATTGGTCAGGGAGTATTCGCTGAATATT

ACCGAGGATATGCTGATTCTCTCATTTGTGCCTCAGGGAAACAGCACATCTTTCATCAGT

GCTATTGAAGTAATATCTGTTCCTGATGATCTGATACAGGATTCAGCGCAAACTGTGAAT

CCCTCCGGCCAGTATCTCGGCCTTGCAACGCAGTCATTTCAGACATTCTACAGGATTAAT

GTGGGTGGACGGGAGGTGACCGTTGTCAATGACACACTCTCGCGTTCATGGGATACTGAC

CAGAACTTCTTCCTAAATTCCACTACCACTGAACTATTTGCTTACCAAGGGAAGCTGAAT

TATCAGAAAGGAGCTGCAACCAAGGAGGATGCACCGGACAGTGTGTACAACACTGCAAGG

CGGTTGGCTGTGCAAAATAGAACCAGCCCAGCGTCCAACATGACATGGCAATTTGATGTT

GACGGTCGTTCGAGCTATCTGATCCGGTTCCATTTTTGTGACATAGTGAGCAAGGCAGCA

TATTCCCTCTACTTTGATATTTATGTGGATGGCGGGTTAGCGTTAGAAAATCTTGACCTC

TCTGAAAAAGTTTTTGGTACCTTGGCTGTGCCATACTACACGGAATTTGTCTTGAAGTCA

AGCAATCCTTCTGGTAAGCTAAGTGTCGGCGTTGGGCCTTCCAGCTTGAACAATGTGGCG

CCAGATGGCATCTTGAACGGCCTGGAGATCATGAAGATGAACATCAGTACTGGCACTATT

TATGTTGTGTGGCCACCAGCACCGCCAAAAAGGAAACTGGCTATCATATTGGGCTCTGTT

CTCGGAGGTGTTGGTGCTGTTAGCATTGCTATTATTCTCTGCTTTGTCCTCAGAAGAAAG

AAGAAGGAGAAGAAGCCGCGGCGGGCACCGACAAGTCGACCTTCAAGTTCTTGGTCACCA

CTTACCCTCAATGGCCTTAGCTTCCTTACCGTAGGTACCCGAACAACCAGCCGGACCACT

CATACATCTGGGACAAACAGTGATGTAAGCTACCGAATTCCTTTTGCTTTGCTGCAAGTG

GCAACAAAACACTTTGACGAGCAGATGGTTGTCGGAGTTGGGGGGTTTGGGAAGGTATAC

AAAGCAGTTCTGCAGGACAGCACCAAAGTCGCAGTCAAGCGGGGCAACCAGAAGTCCCAC

CAAGGGCTCAAAGAATTCCGGACAGAGATCGAGCTGCTGTCGGGGCTGCGACACCGTCAC

CTCGTGTCGCTCATCGGATATTGCGATGAGCAGAACGAGATGATCTTGGTGTATGAGTAC

ATGGAGAAAGGCACGCTGAAGAGTCACCTGTATGGCAGTGACATGCCTCCCCTCAGCTGG

AAGAAAAGAGTGGAAATCTGCATAGGGGCTGCAAGGGGGCTCCACTACCTGCACACAGGT

TTTGCAAAGTCGATCATCCATCGTGATGTCAAGTCGGCAAACATTCTCCTCGACGAAAAT

CTCATGGCCAAGGTTTCTGATTTCGGTCTCTCGAAGACGGGACCTGAGTTGGATCAGACA

CATGTTAGCACTGCGGTGAAAGGGAGCTTCGGGTATCTTGACCCTGAGTACTACCGGAGG

CAGAAGCTGACCGACAAATCGGATGTGTACTCATTTGGCGTGGTCTTGCTGGAGGTGATC

TGCGCGAGGCCAGTCATCGACCCGACGCTTCCAAGAGACATGATCAACCTTGCAGAATGG

GCAATCAAGTGGCAGAAGAGGGGAGAGCTTGGTCAGATTGTCGATCAACGGATCGCTGGG

ACAATCAGGCCAGAGTCATTGAGGAAGTACGGTGAGACGGTCGAGAAGTGTCTTGCGAAT

TACGGCGTCGACCGCCCTACCATGGGTGATGTACTGTGGAATCTGGAGTTTGTGCTCCAG

CTGCAGGAGGCCGGCCCGGACATCTCCAATGTCGACAGCATGAATCAGATCTCTGAACTT

CCTTCAGACGCCAGAAGGATGGGCTCTCTGGAGATCCGCACCGCAGACGAAGCAGACGAA

AGCCGCACAAACATGGATTACTCTCAGATGTCGACCAACGATGCCTTCTCGCAGCTGATT

AACACTGAAGGGAGGTGA

>Ta-CrRLK1L11-D CDS sequence

ATGGGCACCACTACCGGGCAAAAGACAGCTCTGCTCCTTCTTGGGACCCTCTGGGTTCTT

CTTGGTACCTGCAATGCTGCTGAATTCTCCCCTGCAGACAACTACCTCATCAACTGCGGC

TCCACAGTCGACGCTAATCTCCACGATGGGAGGGTCTTCAAAGCAGACAATTCCGGCTCG

ACTATATTGACATCACATCACAGCGTCCCTGCAAACACCTTGCCGGATGCAGTCATAAGT

TCTGACAATCCTGTGCTGTATCAAACTGCAAGGATATTCATTGTGCCGTCGTCCTACTCC

TTCAACATGAAGAGCCGTGGCCGGCATTTTGTCCGGCTACACTTCTTCGGTTTCAGATAC

CAGAGCTATGATCTTGCCGCGGCAAAGTTCAAAGTGTCCACCCAGCATGTTGTGTTACTG

GACAATTTCACTCCGCCGAGCAATTCCTCACCGTTGGATTCAGCGCAAACTGTGAATCCC

TCCGGCCAGTATCTCGGCCTTGCAACACAGTCATTTCAGACATTCTACAGGATTAATGTG

GGTGGACGGGAGGTGACCGTTGTCAATGACACACTCTCGCGTTCATGGGATACTGACCAG

AACTTCTTCATAAATTCCACTACCACTGAACTGTTTGCTTACCAAGGGAGGCTGAATTAT

CAGAAAGGAGCTGCAACCAAGGAGGATGCACCGGACAGTGTGTACAACACTGCAAGGCGG

TTGGCTGTGCAAAATAGAACCAGCCCAGCGTCCAACATGACATGGCAATTTGATGTTGAC

GGTCGTTCGAGCTATCTGATCCGGTTCCATTTTTGTGACATAGTGAGCAAGGCAGCATAT

TCCCTCTACTTTGATATTTATGTGGATGGCGGGTTAGCGTTAGAAAATCTTGACCTCTCT

GAAAAAGTTTTTGGTACCTTGGCTGTGCCATACTACACGGAATTTGTCTTGAAGTCAAGC

AATCCTTCTGGTAAGCTAAGTGTCGGCATTGGGCCTTCCAGCTTGAACAATGTGGCACTA

GATGGCATCTTGAACGGCCTGGAGATCATGAAGATGAACATCAGTACTGGCACTATTTAT

GTTGTGTGGCCACCAGCAACGCCAAAAAGGAAACTGGCTATCATATTGGGCCCTGTTCTC

GGAGGTGTTGGTGCTGTTAGCATTGCTATTATTCTCTGCTTTGTCCTCAGAAGAAAGAAG

AAGGAGAAGAAGCCGCGGCGGGCACCGACAAGTCGACCTTCAAGTTCTTGGTCACCACTT

ACCCTCAATGGCCTTAGCTTCCTTAGCATAGGTACCCGAACAACCAGCCGGACCACTCAT

ACATCTGGGACAAACAGTGATGTAAGCTACCGAATACCTTTTGCTTTGCTGCAAGTGGCA

ACAAAACACTTCGACGAGCAGATGGTTGTCGGAGTTGGGGGGTTTGGGAAGGTATACAAA

GCAGTTCTGCAGGACAGCACCAAAGTCGCAGTCAAGCGGGGCAACCAGAAGTCCCACCAA

GGGCTCAAAGAATTCCGGACAGAGATCGAGTTGCTGTCGGGGCTGCGACACCGTCACCTC

GTGTCGCTCATCGGATATTGCGATGAGCAGAACGAGATGATCTTGGTGTATGAGTACATG

GAGAAAGGTACGCTGAAGAGTCACCTGTATGGCAGTGACATGCCTCCCCTCAGCTGGAAG

AAAAGAGTGGAAATCTGCATAGGGGCTGCAAGGGGGCTCCACTACCTGCACACAGGTTTT

GCAAAGTCGATCATCCATCGTGATGTCAAGTCGGCAAACATTCTCCTCGACGAAAATCTC

ATGGCCAAGGTTTCTGATTTCGGTCTCTCGAAGACGGGACCTGAGTTGGATCAGACACAT

GTTAGCACTGCGGTGAAAGGGAGCTTCGGGTATCTTGACCCTGAGTACTACCGGAGGCAG

AAGCTGACCGACAAGTCGGATGTGTACTCATTCGGCGTGGTCTTGCTGGAGGTGATCTGC

GCGAGGCCAGTCATCGACCCGACGCTTCCAAGAGACATGATCAACCTTGCAGAATGGGCA

ATCAAGTGGCAGAAGAGGGGAGAGCTTGGTCAGATCGTCGATCAACGGATCGCTGGGACA

ATCAGGCCAGAGTCATTGAGGAAGTACGGTGAGACGGTCGAGAAGTGTCTTGCGGATTAC

GGCGTCGACCGCCCTACCATGGGTGATGTCCTGTGGAATCTGGAGTTTGTGCTCCAGCTG

CAGGAGGCCGGCCCGGACATCTCCAATGTGGACAGCATGAATCAGATCTCTGAACTTCCT

TCAGACGCCAGAAGGATGGGCTCTCTGGAGATCGGTACCGCAGACGAAGGCCGCACGAAC

ATGGATTACTCTCAGATGTCGACCAACGATGCCTTCTCGCAGCTGATGAACACTGAAGGG

AGGTGA

>Ta-CrRLK1L12-A CDS sequence

ATGGCGGCCGCCCGAGGCCGAGGCCGAGGCCGAGGCCGATGCGTCCTTCTCGCTGCCGTC

CTCCTCTTGACGGCGGTGGTCGGCGCAGACATATACAAGCCAACGGACTCCATTCTGGTT

CACTGCGGGTCGGACAAGGACGGGCAGGACGAGGACGGCAGGAAGTGGACCACCGACAAG

GACAGCAAGTGGCTCCCCGACGGCGGCAAGTCCTCCATCATGGGCACCGCCGACGTGGCG

GACCCGTCGCTCCCCTCCCCCGTGCCCTACATGACGGCGCGGGTCTTCCCCAAGGAGACC

GCCTACACCTTCCCCGTGGCCGACGCCGACCGCCACTGGGTGCGCCTCCACTTCTACCCG

GCGGCCTACCACGGCATCCCCGCCGACCACTTCTTCTTCTCCGTCACCACCTCCACCGGC

GTCACGCTGCTGCGCAACTTCAGCGTCTACATCACCGCCAAGGCCCTCACCCAGGCCTAC

ATCATCCGGGAGTTCTCCCTCCCTCCCTCCACCGTCGGCTCGCTCTCCCTCAAATTCACG

CCCACCGCCATGAACAACGCCTCCTACGCCTTCGTCAACGGCATCGAGGTCATCTCCATG

CCCAGCTTCTTCGGCGACCCGGCCACACTGGTCGGCCTCAACGACCAGTCCCTCGACGCC

AGCGCCGCCAACCTGCAGACCATGTACCGGCTCAGCGTCGGCGGCTCCTACATCCCGCCC

GCCAACGACTCCGGCCTGTCCCGCGAGTGGTTCTCCGACACGCCCTACGTCTACGGCGCC

GCCACGGGCGTCACCTTCGAGGCCAACGACACGGTCCCGATCAAGTACCCGGCCCCCGCC

GACGAGTACGCCGCGCCCGTCAGCATCTACGACTCGTTCCGCCACATGGGGCGCGACCCC

AAGATGAACAAGAACAACAACCTCACCTGGGTGTTCGAGGTGGACGGCAACTTCACCTAC

CTCCTCCGCCTCCACTTCTGCTCGCTCATGGAAGACAAGATCAACCAGGTCGTCTTCGCC

ATCCTCCTCAACAACAAGACCGCCACCACCACCGGCAGCGCCGACATCATCGCCTGGGCC

AAGGAGAAGAACCCTGCTAATCCCGGCGCGCCCGGCAAGGGCGTGCCGGTCTTCAAGGAC

TACGCGGTGTTCATGCCCGCCGCCCCGGCGGGCAACGACACCATCCTCTGGCTCACGCTG

CGCCCGGACACCGCCACCAGAACACAGTTCGTCAACGCTTTCCTCAACGGCCTGGAGGTG

TTTAAGGTGAGCGACGCCTCCGGCAACCTGGCCGGCCCAAACCCGGACATCTCCAAGATG

CTGGCGGAGGCCGAGCTGGGGGCCGTGGAGGGGCAGTTCAGGGAGAAGCCGAGCAACGTC

GGGGCGCTCATCGGCGGGGCGGCGGGCGGCGCGGCGGCGTTCGGGCTGGTGGCGGCCGTG

TGCTTCGTGGCGTACCAGAGCAAGAGGAGGAGGGAGCTGAGCAGCAGCCCGTCACACTCC

TCCTCCGGGTGGCTGCCGGTGTACGGCGGGTCGACGAGCGTGAGCAAGTCGTCGGGCGGC

AGGAGCGCGGCGACGCTCAACCCCAACATCACGGCCATGTGCCGGCACTTCTCGCTCCAG

GAGATAAAGTCGGCGACCAAGGGGTTCGACGAGTCGCTGGTGATCGGCGTGGGCGGGTTC

GGCAAGGTGTACCGCGGGGTGGTGGACGGGGACACCAAGGTGGCCGTCAAGCGGAGCAAC

CCGTCGTCGGAGCAGGGGGTACTGGAGTTCCAGACGGAGATCGAGATGCTGTCCAAGCTG

CGGCACAAGCACCTGGTGTCCCTCATCGGCTGCTGCGAGGACAACGGCGAGATGATCCTG

GTGTACGACTACATGGCGCACGGCACGCTGCGGGAGCACCTGTACAACAAGAGCGGCAAG

CCGCCGCTGCCGTGGAGGCAGCGGCTGGAGATCGTCATCGGCGCCGCCCGGGGGCTGCAC

TACCTCCACACGGGCGCCAAGTACACCATCATCCACCGGGACGTCAAGACCACCAACATC

CTGGTGGACGACAAGTGGGTGGCCAAGGTGTCCGACTTCGGCCTCTCCAAGACGGGGCCG

ACGGTGCAGAACCAGACGCACGTGAGCACCATGGTGAAGGGCAGCTTCGGGTACCTGGAC

CCGGAGTACTTCCGGCGGCAGAAGCTGACGGAGAAGTCGGACGTCTACTCGTTCGGCGTG

GTGCTGTTCGAGGTGCTGTGCGGGCGGCCGGCGCTGAACCCGAGCCTGCCGCGGGAGCAG

GTGAGCCTGGCGGACCACGCGCTGAGCTGCCAGCGGAAGGGCACCCTGGAGGAGATCGTG

GACCCGGTGCTGGAGGGGAAGATCGCGCCCGACTGCCTCAAGAAGTTCGCCGAGACGGCG

GAGAAGTGCCTGGCGGACCAGGGCGTGGACCGGCCGTCCATGGGCGACGTGCTGTGGAAC

CTCGAGTTCGCGCTGCAGATGCAGGACACCTTCGACAACGGCGGCAAGCCGCCCGAGGTC

GACGACTACAGCAGCAGCTTCACCATCGCCCAGCCGTCCATGGAGGAGAGCCTGGCGGCC

AACGCCGCGGCGCTCTCGCTCATCAGCGAGGACATGGACGAGGAGGACATTGCCAACTCC

GTCATCTTCTCCCAGATCGCTAAACCAACCGGACGATGA

>Ta-CrRLK1L12-B CDS sequence

ATGGCGGCCGCCCGAGGCCGAGGCGTCCTTCTCGCCGTCCTCCTCTTGACGACGGTGGCC

TTCGCGTTCGTCGGCGCGGACATATACAAGCCAACGGACTCCATTCTGGTTAACTGCGGG

TCGGACAAGGACGGGCAGGACGAGGACGGCAGGAAGTGGACCACCGACAAGGACAGCAAG

TGGCTCCCCGACGGCGGCAAGTCCTCCATCATGGGCACCGCCGACGTGTCGGACCCGTCC

CTCCCCTCCCCCGTGCCCTACATGACGGCGCGGGTCTTCCCTAAGGAGACCGCCTACACC

TTCCCCGTGTCCGACGCCGACCGCCACTGGGTGCGCCTCCACTTCTACCCGGCGGCTTAC

CACGACATCCCCGCCGACCACTTCTTCTTCTCCATCAGCACCTCCACCGGCATCACGCTG

CTGCGCAACTTCAGCGTCTACATCACCGCCAAGGCCCTCACCCAGGCCTACATCGTCCGG

GAGTTCTCCCTCCCTCCCTCCACCGCCGGCTCGCTCTCCCTCAAATTCACGCCCACTGCC

ATGAACAATGCCTCCTACGCCTTCGTCAATGGCATCGAGATCATCTCCATGCCCAACTTC

TTCGGCGACCCGGCCACGCTGGTCGGCCTCGACGACCAGTCCCTCGACGCCAGCGCCGGC

AACCTGCAGACCATGTACCGGCTTAGCGTCGGCGGCTCCTACATCCCGCCCACCAACGAC

TCCGGGCTGACCCGCGAGTGGTTCTCCGACACGCCCTACGTCTACGGCGCCGGCACGGGC

GTCACCTTCGAGGCCAACGACACGATCCCGATCAAGTACCCGGCCCCCGCCGACGAGTAC

GCCGCGCCCGTCAGCATCTATGACACGTTCCGCCACATGGGCCGCGACGCCAACCTGAAC

AAGAACAACAACCTCACCTGGGTGTTCGAGGTGGACGGCAACTTCACCTACCTCCTCCGC

CTCCACTTCTGCTCGCTCATGGAAGACAAGATCAACCAGGTCGTCTTCGCCATCCTCGTC

AACAACAAGACGGCCACCACCACCGGCAGCGCCGACATCATCGCCTGGGCCAAGGAGAAG

AACCCTGCTAATCCCGGCGCGCCCGGCAAAGGCGTGCCGGTCTTCAAGGACTACGCCGTG

TTCATGCCCGCCGCTCCGGCGGGCAACGACACCATCCTCTGGCTCACGCTGCGCCCAGAC

ACCGCCAGTAACCCACAGTTCGTCAACGCTTTCCTCAACGGCCTCGAGATCTTTAAGGTG

AGCGACGCCTCCGGCAACCTGGCCGGCCCAAACCCCGACATTTCTAAGATGCTGGCGGAG

GCCGAGCTGGGGGCCGTGGACGGGCAGTTCAGGGAGAAGCCGAGCAACGTCGGGGCGCTC

ATCGGCGGGGCGGTGGGCGGCGCGGCGGCATTCGGGCTGGTCGCGGCCGTGTGCTTCGTG

GCGTACCAGAGCAAGAGGGGGAGGGAGCTGAGCAGCAGCCCATCGCACTCCTCCTCCAGG

TGGCTGCCGGTGTACGGCAGCTCGCAGACGAGCGTGAGCAAGTCGTCGGGCGGGAGGAGC

GCGATGACGCTGAACCCCAACATCACGGCCATGTGCCGGCACTTCTCGTTCCAGGAGATA

AAGTCGGCGACCAAGGGGTTCGACGAGTCGCTGGTGATCGGCGTGGGCGGGTTCGGCAAG

GTGTACCGGGGGGTGGTGGACGGGGACACCAAGGTGGCCATCAAGCGGAGCAACCCGTCG

TCGGAGCAGGGGGTGCTGGAGTTCCAGACGGAGATCGAGATGCTGTCCAAGCTGCGGCAC

AAGCACCTGGTGTCCCTCATCGGGTGCTGCGAGGACAACGGCGAGATGATCCTGGTGTAC

GACTACATGGCGCACGGCACGCTGCGGGAGCACCTGTACAAGAGCGGCAAGCCGCCGCTG

CCGTGGAGGCAGCGGCTGGAGATCGTGATCGGCGCCGCCCGGGGGCTCCACTACCTCCAC

ACGGGCGCCAAGTACACCATCATCCACCGCGACGTCAAGACCACCAACATCCTCGTCGAC

GAGAAGTGGGTGGCCAAGGTGTCCGACTTCGGGCTGTCCAAGACGGGGCCGACGGTGCAG

AACCAGACGCACGTGAGCACCATGGTGAAGGGCAGCTTCGGGTACCTGGACCCGGAGTAC

TTCCGGCGGCAGAAGCTGACGGAGAAGTCGGACGTCTACTCGTTCGGCGTGGTGCTGTTC

GAGGTGCTGTGCGGGCGGCCGGCGCTGAACCCTAGCCTGCCGCGGGAGCAGGTGAGCCTG

GCGGACCACGCGCTGAGCTGCCAGCGGAAGGGCACCCTGGAGGAGATCATCGACCCGGTG

CTGGAGGGGAAGATCGCGCCCGACTGCCTCAAGAAGTTCGCCGAGACGGCGGAGAAGTGC

CTGGCGGACCAGGGCGTGGACCGGCCGTCCATGGGCGACGTGCTGTGGAACCTCGAGTTC

GCGCTGCAGCAGCAGGACACCTTCGAGAACGGCGGGAAGCCGCCCGAGGTGGACGACTAC

AGCAGCAGCTTCACCATCACCCCGCCGTCCATGGAGGAGAGCCTGGCGGCCAACGCGGCG

GCGCTGTCGCTCATCAGCGAGGACATGGACGAGGAGGACATCGCCAACAGCGTCATCTTC

TCCCAGATCGCAAAACCCACCGGACGATGA

>Ta-CrRLK1L12-D CDS sequence

ATGGCGGCCGCCCGAGGCCGAGGCGTCCTTCTCGCCGTCCTCCTCTTGATGATGGTGGCG

TTTGCGTTCGTCGGCGCAGACATATACAAGCCAACGGACTCCATTCTGGTTCACTGCGGG

TCGGACAAGGACGGGCAAGACGAGGACGGCAGGAAGTGGACCGCCGACAAGGACAGCAAG

TGGCTCCCCGACGGGGGCAAGTCCTCCGTCATGGGCACCGCCGACGTGCCGGACCCGTCG

CTCCCCTCCCCTGTGCCCTACATGACGGCGCGGGTCTTCCCCAAGGAGACCGCCTACACC

TTCCCCGTGGCCGACGCCGACCGCCACTGGGTGCGCCTCCACTTCTACCCGGCGGCCTAC

CACGGCATCCCCGCCGACCACTTCTTCTTCTCCGTCACCACCTCCACCGGCGTCACGCTG

CTCCGCAACTTCAGCGTCTACACCACCGCCAAGGCCCTCACCCAGGCCTACATCGTCCGG

GAGTTCTCCCTCCCTCCCTCCACCACCGGCTCGCTCTCCCTCAAATTCACGCCCACCGCC

ATGAACAACGCCTCCTACGCCTTCGTCAACGGCATCGAGATCATCTCCATGCCCAGCTTC

TTCGGCGACCCGGCCACGCTGGTCGGTCTCGACGACCAGTCCCTCGACGCCAGCGCCGGC

AACCTGCAGACCATGTACCGGCTCAGCGTCGGCGGCTCCTACATCCCTCCCGCCAACGAC

TCCGGGCTGTCCCGTGAGTGGTTCTCCGACACACCCTACGTCTACGGCGCCGCCACGGGC

GTCACCTTCGAGGCCAACGACACGATCCCGATCAAGTACCCGACCCCCGCCGACGAGTAT

GCCGCGCCCGTCAGCATCTACGACTCGTTCCGCCACATGGGGCGCGACCCCAAGATGAAC

AGGAACAACAACCTCACCTGGGTGTTCGAGGTGGACGGCAACTTCACCTACCTCCTCCGC

CTCCACTTCTGCTCGCTCATGGAAGACAAGATCAACCAGGTCGTCTTCGCCATCCTCGTC

AACAACAAGACGGCCACCACCACCGGCAGCGCCGACATCATCGCCTGGGCCAAGGAGAAG

AACCCTGCTAATCCCGGCGCGCCCGGCAAAGGCGTGCCGGTCTTCAAGGACTACGCCGTG

TTCATGCCCGCCGCTCCGGCAGGCAGCGACACCATCCTCTGGCTCACGCTGCGCCCAGAC

ACCGCCACCAATCCACAGTTCGTCAACGCTTTCCTCAACGGCCTGGAGGTCTTTAAGGTG

AGCGACGCCTCCGGCAACCTGGCCGGCCCAAACCCCGACATCTCCAAGATGCTGGCGGAG

GCCGAGCTGGGGGCCGTGGACGGGCAGTTCAGGGAGAAGCCGAGCAACGTCGGGGCGCTC

ATCGGCGGGGCGGCGGGCGGCGCGGCGGCGTTCGGGCTGGTGGCGGCCGTGTGCTTCGTG

GCGTACCAGAGCAAGAGGAGGAGGGAGCTGAGCAGCAGCCCGTCGCACTCCTCCTCCGGG

TGGCTGCCGGTGTACGGCGGCAACTCGCAGACGAGCGTGAGCAAGTCGTCGGGCGGCAGG

AGCGCGGTGACGCTGAACCCCAACATCACGGCCATGTGCCGGCACTTCTCGTTCCAGGAG

ATAAAGTCGGCGACCAAGGGGTTCGACGAGTCGCTGGTGATCGGCGTGGGCGGGTTCGGG

AAGGTGTACCGGGGGGTGGTGGACGGGGACACCAAGGTGGCCATCAAGCGGAGCAACCCG

TCGTCGGAGCAGGGGGTGCTGGAATTCCAGACGGAGATCGAGATGCTGTCCAAGCTGCGG

CACAAGCACCTGGTGTCCCTCATCGGGTGCTGCGAGGACAACGGCGAGATGATCCTGGTG

TACGACTACATGGCGCACGGCACGCTGCGGGAGCACCTGTACAAGAGCGGCAAGCCGCCG

CTGCCGTGGAGGCAGCGGCTGGAGATCGTGATCGGCGCCGCCCGGGGGCTCCACTACCTC

CACACGGGCGCCAAGTACACCATCATCCACCGCGACGTCAAGACCACCAACATCCTGGTG

GACGAGAAGTGGGTGGCCAAGGTGTCCGACTTCGGGCTGTCCAAGACGGGGCCGACGGTG

CAGAACCAGACGCACGTGAGCACCATGGTGAAGGGCAGCTTCGGGTACCTGGACCCGGAG

TACTTCCGGCGGCAGAAGCTGACGGAGAAGTCGGACGTCTACTCGTTCGGCGTGGTGCTG

TTCGAGGTGCTGTGCGGGCGGCCGGCGCTGAACCCGAGCCTGCCGCGGGAGCAGGTGAGC

CTGGCGGACCACGCGCTGAGCTGCCAGCGGAAGGGCACCCTGGAGGAGATCATCGACCCG

GTGCTGGAGGGGAAGATCGCGCCCGACTGCCTCAAGAAGTTCGCCGAGACGGCGGAGAAG

TGCCTGGCGGACCAGGGCGTGGACCGGCCGTCCATGGGCGACGTGCTGTGGAACCTCGAG

TTCGCGCTGCAGCAGCAGGACACCTTCGAGAACGGCGGGAAGCCGCCCGAGGTGGACGAC

TACAGCAGCAGCTTCACCATCACCCCGCCGTCCATGGAGGAGAGCCTGGCGGCCAACGCG

GCGGCGCTCTCGCTCATCAGCGAGGACATGGACGAGGAGGACATCGCCAACAGCGTCATC

TTCTCCCAGATCGCAAAACCCACCGGACGATGA

>Ta-CrRLK1L13-A CDS sequence

ATGGTGCGCCGCGGGGCGCTCCCGCTGGCGCTGCTGGCCGTGCTCGCGACGCTGACGGCC

GTGGCGGGGCAGGGGAAGCCGGTCACGGACAACGGCTCGGGCGGCGGGTCGGGGCCGTCC

AAGTTCACGCCCAAGGACGCCTTCTACATCGACTGCGGCGGCACGGCCGCCGCCGACACC

AAGGACGGCAAGTCCTTCAAGACCGACGCGGAGGCCAACAGCCTGCTCTCCGCCAGGGAC

AACATCAAGGTCGCCGACGACAAGGCCGACGTGCCGTCGCACCTCTACCGCTCCGCGCGG

GTCTTCAAGGAGGAGGCCGTCTACAACTTCCCGCTCACGGCCCCCGGCTGGCACTTCATC

CGGCTCTACTTCTTCCCCATCAAGAGCGGGGAGGCCGACCTCGCGGCGGCCACGTTCGAC

GTGTCCACCGCCGTTAACGTCCTTCTCCACGGCTTCACCCCCGAGGCGAAGGCGGTCATG

AAGGAGTACATCGTCAACGCCACGGAGAACAAGCTCGAGCTCAAGTTCACCCCGCAGTCG

GGCTCGGCGTTCATCAACGCCATCGAGGTCGTCAACGCCCCCGACGAGCTCATCAGCAAG

ACGGCCCTGACGGTGTCGCCGCTAGCCGAGACAAGCGGGTTGTCAGAGGCTGCGTACCAG

GTGGTGTGCCGGCTCAACGTCGGTGGCCCGCCCATCGGCCCCGTGAACGACACGCTCGGC

CGGCAGTGGGAGGACGACGGGCAGTACCTGAACCCCAAGGACGCCGGGACGGAGGTGTCG

GTGCCGACGAGCGCGATCAAGTACCCCGACGCGTTCCCGGCGACCAAGCTCGTGGCACCC

ACGGCGGTGTACGCGACCGCCCGCCACATGGCTGAATCCGGCGTCGCGAACCAGAACTTC

AACGTGTCGTGGAAGGTGGACGTGGACCCGTCGTTCGACTATCTCGTCCGCCTCTTCTTC

GCCGACATCATAAGCACGTCCGCCAACGACCTCTACTTCAACGCGTACATCAACGGCCGC

AAGGCCATCTCCGCCCTGGACCTCTCCACCATCACCGGCGACCTGGCCGCGCCCTACTAC

AAGGACTTCGTGGTGAACTCGTCGGTCAACACCGACGGCCACATTATCATCGGGGTCGGG

CCGCTGGGGCAGGACACGGGCCGCAACGACGCGCTGCTCAACGGCGCGGAGGTGCTCAAG

ATGAGCAACTCGGTGGGCAGCCTGGACGGCGAGTTCGGCGTGGACGGCCGGATGGTGGAC

GACGGCAGCGGCACCCGCAAAGTGGTCGCTGCCGTGGGGTTCGCCATGATGTTCGGCGCC

TTCGCCGGCCTGGGATGCATGGTGGTGAAGTGGCACCGGCGGCCGCAGGACTGGGACCGG

CGCAACAGCTTCTCGTCGTGGCTGCTGCCCATCCACACGGGCCAGTCCTTCTCCAACGGC

AAGGGGTCCAAGAGCGGCTACACCTTCTCCTCCACCGCGGGGCTGGGCCACTTCTTCACC

TTCGCGGAGATGTCAGAGGCGACCAAGAACTTCGACGAGAGCGCCATCATCGGCGTGGGA

GGGTTCGGCAACGTGTACGTGGGCGAGATCAACGACCCCGACGAGGAGGGGTCCAGGATC

AAGGTGGCCATCAAGCGCGGGAACCCGTCGTCGGAGCAGGGCATCAACGAGTTCAACACC

GAGATCCAGATGCTGTCCAAGCTCCGGCACCGCCACCTCGTGTCCCTCATCGGCTACTGC

GACGAGGGCGAGGAGATGATCCTCGTCTACGAGTTCATGCAGCACGGGCCCTTCCGCGAC

CACATCTACGGCGGCCCCGAGGGCCTGCCCACGCTCTCCTGGAAGCAGCGCCTCGAGATC

TGCATCGGCGCCGCCAGGGGCCTCCACTACCTCCACACCGGCACCGCGCACGGGATCATC

CACCGGGACGTCAAGACCACCAACATCCTCCTCGACGAAAAGTTCGTGGCCAAGGTGGCC

GACTTCGGCCTCTCCAAGGACGGCCCCGGGATGAACCAGCTGCACGTCAGCACCGCCGTC

AAGGGCAGCTTCGGGTACCTCGACCCGGAGTACTTCCGGTGCCAGCAGCTGACCGACAAG

TCGGACGTCTACTCCTTCGGGGTGGTGCTGCTGGAGACGCTGTGCGCGCGGGCGCCCATC

GACCCGCAGCTGCCGCGCGAGCAGGTCAGCCTCGCCGAGTGGGGCCTGCAGTGGAAGCGC

AAGGGCCTCATCGAGAAGATCATGGACCCCAACCTCAACGGCAAGGTCAACCCGGAGTCG

CTCGCCAAGTTCGCCGAGACCGCCGAGAAGTGCCTCTGCGAGTTCGGCAGCGACCGCCTC

TCCATGGGCGACGTGCTCTGGAACCTCGAGTACGCGCTGCAGCTGCAGGAGGCCAACCCG

CCCGAGGGCGCCACCGACGCCGACGACGCCGACGCCTCCATCGTCTCCTCCGCCAGCGGC

GTCACCACCGTGCCCGACCAGTCCACCACCTCCGCCAACGAGCTCTTCGCGCAGCTCGCC

GACATGAAGGGGAGATGA

>Ta-CrRLK1L13-B CDS sequence

ATGGTGCGCCGCGGGACGTTCCCGCTGGCGCTGCTGGCCGTGCTGGCGACGCTGACGGCC

GTGGCGGGGCAGGGGAAGCCGGTCACGGACAACGGCTCGGGCGGCGCGTCGGGGCCGGCC

AAGTTCACGCCCAAGGACGCCTTCTACATCGACTGCGGCGGCACGGCCGCCGCCGACACC

AAGGACGGCAAGTCCTTCAAGACCGACGCGGAGGCCAACAGCCTGCTCTCCGCCAGGGAC

AACATCAAGGTCGCCGACGACAAGGCCGACGTGCCGTCGCACCTCTACCGCAGCGCGCGG

GTCTTCAAGGAGGAGGCCGTCTACAACTTCCCGCTCACGGCCCCCGGCTGGCACTTCATC

CGGCTCTACTTCTTCCCCATCAAGAGCGGGGAGGCCGACCTCGCGGCGGCCACGTTCGAC

GTGACCACCGCCGTGAACGTCCTTCTCCATGGCTTCACCGCCGAGGCGAAGGCGGTCATG

AAGGAGTACGTCGTCAACGCCACGGAGAACAAGCTCGAGCTCAAGTTCACCCCGCAGTCG

GGCGCGGCGTTCATCAACGCCATCGAGGTCGTCAACGCCCCTGACGAGCTCATCAGTAAG

ACGGCCCTGACGGTGTCGCCGCTAGCCGAGACAAGCGGGCTGTCAGAGGCTGCCTACCAG

GTGGTGTGCCGGCTCAACGTCGGTGGCCCGCCCATCGGCCCCGTGAACGACACGCTCGGC

CGGCAGTGGGAGGACGACGGGCAGTACCTGAACCCCAAGGAGGCCGGGGCGGAGGTGTCG

GTGCCGACGAGCGCGATCAAGTACCCCGACGCGTTCCCGGCGACCAAGCTCGTGGCACCC

ACGGCGGTGTACGCGACCGCCCGCCACATGGCTGAATCCGGCGTCGCGAACCAGAACTTC

AACGTGTCGTGGAAGGTGGACGTGGACCCGTCGTTCGACTATCTCGTCCGCCTCTTCTTC

GCCGACATTATAAGCACGTCCGCCAACGACCTATACTTCAACGTGTACATCAACGGCCGC

AAGGCCATCTCCGCCCTGGACCTCTCCACCATCACCGGCGACCTGGCCGCGCCCTACTAC

AAGGACTTCGTGGTGAACTCGTCGGTCAATACCGACGGCCACATCATCATCGACGTCGGG

CCGCTAGGGCAGGACACGGGCCGCAACGACGCGCTGCTCAACGGCGCGGAGGTGCTCAAG

ATGACCAACTCGGTGGGCAGCCTGGACGGCGAGTACGGCGTGGACGGCCGGATGGTGGAC

GACGGCAGCGGCACCCGCAAGGTGGTGGCGGCCGTGGGGTTCGCCATGATGTTCGGCGCC

TTCGCCGGCCTGGGATGCATGGTGGTGAAGTGGCATCGGCGGCCGCAGGACTGGGAGCGG

CGCAACAGCTTCTCGTCGTGGCTGCTGCCGATCCACACGGGCCAGTCCTTCAGCAACGGC

AAGTCCAAGAGCGGTTACACCTTCTCCTCCACCGCGGGGCTGGGCCACTTCTTCACCTTC

GCGGAGATGTCAGAGGCGACCAAGAACTTCGCCGAGAGCGCCATCATCGGCGTGGGAGGG

TTCGGCAACGTGTACGTGGGCGAGATCAACGACCCCGACGAGGAGGGCTCCAGGATCAAG

GTGGCCATCAAGCGCGGGAACCCGTCGTCGGAGCAGGGCATCAACGAGTTCAACACCGAG

ATCCAGATGCTGTCCAAGCTCCGGCACCGCCACCTCGTCTCCCTCATCGGCTACTGCGAC

GAGGGCGAGGAGATGATCCTCGTCTACGAGTTCATGCAGCACGGGCCCTTCCGCGACCAC

ATCTACGGCGGGCCCGAGGGCCTGCCCACGCTCTCCTGGAAGCAGCGCCTCGAGATCTGC

ATCGGCGCCGCCAGGGGCCTCCACTACCTCCACACCGGCACCGCGCACGGGATCATCCAC

CGCGACGTCAAGACCACCAACATCCTCCTCGATGACAAGTTCGTGGCCAAGGTGGCCGAC

TTCGGCCTCTCCAAGGACGGCCCCGGGATGAACCAGCTGCACGTCAGCACCGCCGTCAAG

GGCAGCTTCGGGTACCTCGACCCGGAGTACTTCCGGTGCCAGCAGCTGACCGACAAGTCC

GACGTCTACTCATTCGGGGTGGTGCTGCTGGAGACGCTGTGCGCGCGGGCGCCCATCGAC

CCGCAGCTGCCGCGCGAGCAGGTCAGCCTCGCCGAGTGGGGCCTGCAGTGGAAGCGCAAG

GGCCTCATCGAGAAGATCATGGACCCCAACCTCGCCGGCAAGGTCAACCCGGAGTCGCTC

GCCAAGTTCGCCGAGACCGCCGAGAAGTGCCTCTGCGAGTTCGGCAGCGACCGCCTCTCC

ATGGGCGACGTGCTCTGGAACCTCGAGTACGCGCTGCAGCTGCAGGAGTCCAACCCACCC

GAGGGCGCCAGCGACGCCGACGACGCCGACGCCTCCATCGTCTCCTCCGCCAGCGGCGTC

ACCACCGTGCCCGACCAATCCACCACCTCCGCCAACGAGCTCTTCGCGCAGCTCGCCGAC

ATGAAGGGCAGGTGA

>Ta-CrRLK1L13-D CDS sequence

ATGGTGCGCCGCGGGGCGCTCCCGCTTGCGCTGCTGGCCGTGCTCGCGACGCTGACGGCC

GTGGCGGGGCAGGGGAAGCCGGTCACGGACAACGGCTCGGGCGGCGGGGCGGGGCCAGCC

AAGTTCACGCCCAAGGACGCCTTCTACATCGACTGCGGCGGCACGGCCGCCGCCGACACC

AAGGACGGCAAGTCCTTCAAGACCGACGCGGAGGCCAACAGCCTGCTCTCCGCCAGGGAC

AACATCAAGGTCGCCGACGACAAGGCCGACGTGCCGTCGCACCTCTACCGCTCCGCGCGG

GTCTTCAAGGAGGAGGCCGTCTACAACTTCCCGCTCACGGCCCCCGGCTGGCACTTCATC

CGGCTCTACTTCTTCCCCATCAAGAGCGGGGAGGCCGACCTCGCGGCGGCCACGTTCGAC

GTGAGCACCGCCGTCAACGTCCTTCTCCATGGCTTCACCCCCGAGGCGAAGGCGGTCATG

AAGGAGTACATCGTAAACGCCACGGAGAACAAGCTCGAGCTCAAGTTCACCCCGCAGTCG

GGCTCGGCGTTCATCAACGCCATCGAGGTCGTCAACGCCCCCGACGAGCTCATCAGTAAG

ACGGCCCTGACGGTGTCGCCGCTAGCCGAGACAAGCGGGTTGTCAGAGGCTGCGTACCAG

GTGGTGTGCCGGCTCAACGTCGGTGGCCCGCCCATCGGCCCCGTGAACGACACGCTCGGC

CGGCAGTGGGAGGACGACGAGAAGTACCTGAACCCCAAGGAGGCCGGGACGGAGGTGTCG

GTGCCGACGAGCGCGATCAAGTACCCCGACGCGTTCCCGGCGACCAAGCTCGTGGCACCC

ACGGCGGTGTATGCGACCGCCCGCCACATGGCTGAATCCGGCGTCGCGAACCAGAACTTC

AACGTGTCGTGGAAGGTGGACGTGGACCCGTCGTTCGACTATCTCGTCCGCCTCTTATTC

GCCGACATTATAAGCACGTCCGCCAACGACCTCTACTTCAACGTGTACATCAACGGCCGC

AAGGCCATCTCCGCCCTGGACCTCTCCACCATCACCGGCGACCTGGCCGCGCCCTACTAC

AAGGACTTCGTGGTGAACTCGTCGGTCAATACCGATGGCCACATTATCATCGACGTCGGG

CCGCTAGGGCAGGACACGGGCCGCAACGACGCGCTGCTCAACGGCGCGGAGGTGCTCAAG

ATGAGCAACTCGGTGGGCAGCCTGGACGGCGAGTACGGCGTGGACGGCCGGATGGTGGAT

GACGGCAGCGGCACCCGCAAAGTGGTGGCGGCCGTGGGGTTCGCCATGATGTTCGGCGCC

TTCGCCGGCCTGGGATGCATGGTGGTGAAGTGGCACCGGCGGCCGCAGGACTGGGAGCGG

CGCAACAGCTTCTCGTCGTGGCTGCTGCCCATCCACACGGGCCAGTCCTTCAGCAACGGC

AAGTCCAAGAGCGGCTACACCTTCTCCTCCACCGCGGGGCTGGGCCACTTCTTCACCTTC

GCGGAGATGTCAGAGGCGACCAAGAACTTCGACGAGAGCGCCATCATCGGCGTGGGAGGG

TTCGGCAACGTGTACGTGGGCGAGATCAACGACCCCGATGAGGAGGGGTCCAGGATCAAG

GTGGCCATCAAGCGCGGGAACCCGTCGTCGGAGCAGGGCATCAACGAGTTCAACACCGAG

ATCCAGATGCTGTCCAAGCTCCGGCACCGCCACCTCGTGTCCCTCATCGGCTACTGCGAC

GAGGGCGAGGAGATGATCCTCGTCTACGAGTTCATGCAGCACGGGCCCTTCCGCGACCAC

ATCTACGGCGGCCCCGAGGGCCTGCCCACGCTCTCCTGGAAGCAGCGCCTCGAGATCTGC

ATCGGCGCCGCCAGGGGCCTCCACTACCTCCACACCGGCACCGCGCATGGGATCATCCAC

CGCGACGTCAAGACCACCAACATCCTCCTCGACGACAAGTTCGTCGCCAAGGTGGCCGAC

TTCGGCCTCTCCAAGGACGGCCCCGGCATGAACCAGCTGCACGTCAGCACCGCCGTCAAG

GGCAGCTTCGGGTACCTCGACCCGGAGTACTTCCGGTGCCAGCAGCTGACCGACAAGTCC

GACGTCTACTCCTTCGGGGTGGTGCTGCTGGAGACGCTGTGCGCGCGGGCGCCCATCGAC

CCGCAGCTGCCGCGCGAGCAGGTCAGCCTCGCCGAGTGGGGCCTGCAGTGGAAGCGCAAG

GGCCTCATCGAGAAGATCATGGACCCCAACCTCAACGGCAAGGTCAACCCGGAGTCGCTC

GCCAAGTTCGCCGAGACCGCCGAGAAGTGCCTCTGCGAGTTCGGCAGCGACCGCCTCTCC

ATGGGCGACGTGCTCTGGAACCTCGAGTACGCGCTGCAGCTGCAGGAGGCCAACCCGCCC

GAGGGCGCCACCGACGCCGACGACGCCGACGCCTCCATCGTCTCATCCGCCAGCGGCGTC

ACCACCGTGCCCGACCAGTCCACCACCTCCGCCAACGAGCTCTTCGCGCAGCTCGCCGAC

ATGAAGGGGAGATGA

>Ta-CrRLK1L14-A CDS sequence

ATGCCCGCCGCGGGGCGCTCCGGTGGGCCGGGACAGGTCAACATTATGATGGGAAGGAGG

AAGTTGCAAGTGGTGACCTTGGCGATCTTGTGTTTCTGGTCATCTGCTGGGATCTGCAAA

GCACAATCAGTGGATTTCAAGCCTGCCGACAGCTACCTGGTTGACTGTGGGTCTGCCAAG

GGCACGACGGTTCTCGGGAGGGACTTCGCTGCCGATGGGGCAGCTCCGGTGACCGTGGCC

ACCTCCCAAGACATCCTTGCCGGCACCTCGGCCAACGGGGTGTCCTCGTTTGACAACCCG

GTGCTTTACCAGACTGCCCGCATCTTCACGAGCCCGTCATCCTATACTTTTCCTATCCAG

AAGCAGGGGCGGCATTTTGTTCGCCTCTACTTCTACCCCTTCATCTACCAGAGTTATGAT

CTCTCCACCGCCAAGTTCACCGTGTCGACCCAAGATGTGCTCCTGCTCAGTGATTTCCAG

CAGCCGGACAAGACGGCGCCACTGTTCAAGGAATACTCTTTGAACATCACACGTGACCAG

CTTGTCATTTCCTTCAAGCCGTCAAACGGAATTGCATTCATCAATGCAATTGAAGTGATT

TCTGTTCCAGATGATCTCATAGCCGATGTAGCCAATATGGTCAACCCTGTGCAGCAATAC

AGCGGTTTGACTACACAGTCACTGGAGACAGTGTATCGTGTCAACATGGGTGGTCCGAAG

GTCTTCCCGAACAATGATACCCTCTCGAGGACTTGGCAGAAGGATCAGAAGTACATACTG

AACCCCAGTGTGACTAAAACTGCTCAATATGGCAAGGCTATCAACTACAGGAAAGGTGGA

GCAACTCCACTGACGGCCCCTGATATTGTGTACAGTACAGCTACAGAATTGGCGGCTTCA

AACACATCCAACGCACTTTTCAACATGACATGGCAGTTTGATGTGGATGCAGGCTTCAGC

TATCTGATAAGATTTCACTTCTGTGATATAGTCAGCAAGGCACTGAACCAGCTCTACTTC

AATGCATATGTGGGAGGATTCTTTGCACAGCATGATCTTGATCTCTCAGAGCAATCGGTG

AATCAATTGGCTACAGCTATCTATGTTGATGTGGTTCTTTCTTCAAATGATGCGTCGAGC

AAGCTCAGCATCAGTATTGGTCCATCCACCTTGAACAATGCATTGCCTGATGGGATTCTG

AATGGCCTTGAGATTATGAAGATGGGCAGTGGCTCTGGTTCTGCTTTCACTGTTGGGAAT

AACGGTTCAAACAAAAAGTTGCCCATAATTATTGGCTCAGTCCTTGGGGTTGTTGGGCTT

CTGATAATTGTCCTTGTTGTGGTACTGCTTTGCCGGAGGAAGAAGACCGACGACAAGCAG

CACTCCAAGACCTGGATGCCTTTCTCTATCAATGGGCTCACGTCTCTCAGTACAGGAAGT

AGAACTTCTTATGGTACTACACTAACATCAGGTCTGAATGGAAGCTATGGATATCGCTTC

GCCTTCAATGTGCTCCAAGAAGCAACAAACAATTTTGATGAGAGCTGGGTGATTGGAGTT

GGAGGTTTTGGGAAAGTCTACAAGGGAGCTTTGAGGGATGACACAAAGGTTGCCGTGAAG

CGAGGAAACCCCAAGTCCCAGCAAGGTCTGAATGAGTTCCGGACAGAGATCGAACTCCTT

TCGCGTCTGCGTCATCGCCACCTGGTGTCTCTTATTGGGTACTGTGATGAAAGGAATGAG

ATGATCTTGGTCTATGAGTACATGGAGAACGGAACCGTCAAAAGCCACCTGTATGGTTCA

GACAACCCCTCACTCAACTGGAAGCAGCGGCTGGAGATCTGCATTGGAGCAGCAAGGGGG

CTACACTATCTTCATACCGGTTCTGCAAAGGCCATTATCCACCGTGATGTCAAGTCTGCA

AACATCTTGCTTGATGAGAATCTCCTCGCGAAAGTCGCCGACTTTGGGCTATCAAAGACT

GGGCCTGAGCTGGATCAGACTCACGTCAGCACTGCAGTGAAGGGCAGCTTTGGGTACCTC

GACCCTGAATACTTCCGAAGGCAGCAGCTGACCGAGAAGTCAGACGTCTACTCCTTCGGT

GTTGTCATGCTGGAGGTGCTCTGTGCGAGGCCGGTGATCGACCCTTCACTCCCGAGGGAA

ATGGTGAACTTGGCAGAGTGGGGAATGAAGTGGCAGAAGAGAGGGGAGCTGCACCAGATC

GTCGACCAGAAGCTTTCCGGCGCGATCAGGCCGGACTCTCTGAGGAAGTTCGGTGAGACG

GTGGAGAAGTGCCTGGCAGACTACGGCGTGGAGCGGCCGTCGATGGGGGACGTCCTCTGG

AACTTGGAGTATGTCCTGCAGCTCCAGGACGTGGATTCTTCGACCGTGTCGGACGTGAAC

AGCATGAACCGGATCGTCGACCTGTCATCGCAGGTTCAGCACGTCAGTGCCATGGAGAGC

ATCAGCGTGACGATGGCGGAGGACGGAGCTTTGCACGAGCCTGACCACGACCTCTCCGAC

GTGTCGATGAGCCGGGTTTTCTCTCAGCTGATCAAAGCCGAGGGGAGGTGA

>Ta-CrRLK1L14-B CDS sequence

ATGCCCGCCGCGGCGCGCTCCGGTGGACCGGGGCAGGCCAACATTATGATGGGGAGGAGG

AAGTTGCAAGCAGTGACCTTGGCGATCTTGTGTTTCTGGTCATCTGCTGGGGCACAAACA

GTGGATTTCAAGCCTGCCGACAACTACCTGGTTGACTGTGGGTCTGCCAAGGGCACGACG

GTTCTCGGGAGGGACTTCGCTGCCGATGGGGCATCTCCGGTGACAGTGTCAACCTCCCAA

GATATTCTTGCCGGCACCTCGGCCAACGGGGTGTCCTCGTTTGACAACCCGCTGCTTTAC

CAGACCGCCCGCATCTTCACGAGCCCGTCATCCTATACTTTTCCTATCCAGAAGCAGGGG

CGGCATTTTGTTCGTCTCTACTTCTTCCCCTTCATCTACCAGAGTTATGATCTCTCCACC

GCCAAGTTCACTGTGTCGACCCAAGATGTGCTCCTGCTCAGTGATTTCCAGCAGCCGGAC

AAAACTGCGCCGCTGTTCAAGGAATACTCTTTGAACATCACCCGTGACCAGCTTGTCATT

TCCTTCAAGCCGTCAAACGGGATTGCATTCATCAACGCAATTGAAGTGGTTTCTGTTCCA

GATGATCTCATAGCTGATGTAGCCAATATGGTCAACCCTGTGCAGCAGTACAGCGGTTTG

ACTACACAGTCACTGGAGACGGTGTATCGTGTCAACATGGGTGGTCCGAAGGTCTTCCCG

AGCAATGATACCCTCTCGAGGACTTGGCAGAAGGATCAGAAGTACATACTGAACCCCAGT

GTGACCAAAACTGCCCAATATGGCAAGCCTATCAAGTATAGGAAAGGCGGGGCAACTCCA

CTGACGGCCCCAGATATTGTGTACAGTACAGCTACAGAATTGGCGGCTGCAAACACTTCC

AACGCACTTTTCAACATGACATGGCAGTTTGATGTGGATGCAGGCTTCAGCTATCTGATA

AGATTTCACTTCTGTGATATAGTCAGCAAGGCACTGAACCAGCTCTACTTCAATGCATAT

GTGGGAGGCTTCTTTGCACAGCATGATCTTGATCTCTCAGAGCAATCGGTGAATCAACTG

GCTACAGCTATCTATGTTGACGTGGTTCTTTCTTCCAATGATGCATCTAGCAAGCTCAGC

ATCAGTATTGGTCCGTCCACCTTGAACAATGCATTGCCTGATGGGATTCTGAATGGCCTG

GAGATTATGAAGATGGGCAGTGGCTCTGGTTCTGCTTTCACTGTTGGGAATAACGGTTCA

AACAAAAGGTTGCCCATAATTATTGGCTCAGTCCTTGGGGTTGTTGGGCTTCTGATAATT

GTCCTTGTTGTGGTACTGCTTTGCCGGAGGAAGAAGACCGACGACAAGCAGCACTCGAAG

ACCTGGATGCCTTTCTCTATCAATGGGCTCACGTCTCTCAGTACAGGAAGCAGAACTTCC

TATGGTACTACACTAACATCAGGTCTGAATGGAAGCTATGGATATCGGTTTGCCTTCAAT

GTGCTCCAAGAAGCAACAAACAATTTTGATGAGAGCTGGGTGATTGGAGTCGGAGGTTTT

GGGAAAGTCTACAAGGGTGCCTTGAGGGATGACACAAAGGTTGCAGTGAAGCGAGGAAAC

CCCAAGTCCCAGCAAGGTCTCAATGAGTTCCGGACAGAGATCGAGCTCCTTTCACGTCTG

CGTCACCGCCACCTGGTGTCTCTTATTGGGTACTGTGATGAAAGGAATGAGATGATCTTG

GTCTATGAGTACATGGAGAACGGAACCGTCAAGAGCCACCTGTATGGTTCAGACAACCCC

TCACTCAACTGGAAGCAACGCCTGGAGATCTGCATTGGAGCAGCAAGGGGGCTACACTAT

CTTCATACAGGTTCTGCGAAGGCCATTATCCACCGTGATGTCAAGTCTGCAAACATCTTG

CTTGATGAAAATCTCCTTGCCAAAGTCGCCGACTTTGGGCTGTCAAAGACCGGGCCTGAG

CTGGATCAGACTCATGTCAGCACTGCAGTGAAGGGTAGCTTTGGTTACCTTGACCCTGAA

TACTTCCGAAGGCAGCAGCTGACTGAGAAGTCGGACGTCTACTCCTTCGGTGTTGTCATG

CTGGAGGTGCTCTGCGCGAGGCCGGTGATCGACCCTTCGCTCCCGAGGGAAATGGTGAAC

TTGGCAGAGTGGGGAATGAAGTGGCAGAAGAGAGGAGAGCTGCACCAGATCGTCGACCAG

AAGCTTTCCGGCGCGATCAGGCCGGACTCTCTGAGGAAGTTCGGTGAGACGGTGGAGAAG

TGCCTGGCAGACTACGGCGTGGAGCGGCCGTCGATGGGGGACGTCCTCTGGAACTTGGAG

TATGTCCTGCAGCTCCAGGATGTGGATTCTTCGACCGTGTCGGACGTGAACAGCATGAAC

CGGATCGTCGACCTGTCGTCGCAGGTTCAACATGTGGGTGCCATGGAGAGCATCAGCGTG

ACGATGGCGGAGGACGGAGCTTTGCACGAGCCTGACCACGACCTCTCCGACGTGTCGATG

AGCAGGGTTTTCTCACAGCTGATCAAAGCCGAGGGGAGGTGA

>Ta-CrRLK1L14-D CDS sequence

ATGGCAGGGGCCGCCGAGCGCACCTCTAGGCCACCGTCCAGCTCTACTCCCCTTTCCGCT

GCATTTGGCCGGAGGGGAGATCCCAGTCCCACCAGGACTGATTTCTACTTGGCCAACATT

ATGATGGGGAGGAGGAAGTTGCAAGTGGTGACCTTGGCGATCTTGTGTTTCTGGTCATCT

GCTGGGGTCTGCAAAGCACAAACAGTCGATTTCAAGCCTGCAGACAGCTACCTGGTTGAC

TGTGGGTCTACCAAGGGCACGACGGTTCTCGGGAGGGACTTCGCTGCCGATGGGGCATCT

CCGGTGACCGTGTCCACCTCCCAAGATATTCTTGCCGGCACCTCGGCCAACGGGGTGTCC

TCTTTTGACAACCCAGTGCTTTACCAGACCGCCCGCGTCTTCACGAGCCCGTCATCCTAT

ACTTTTCCGATCCAGAAGCAGGGGCGGCATTTTGTCCGTCTCTACTTCTACCCCTTCATC

TACCAGAGTTATGATCTCTCCACTGCCAAGTTCACCGTGTCGACCCAAGATGTGCTCCTG

CTCAGTGATTTCCAGCAGCCGGACAAGACGGCGCCGCTGTTCAAGGAATACTCTTTGAAC

ATCACCCGTGACCAGCTTGTTATTTCCTTCAAGCCGTCAAACGGAATTGCATTCATCAAC

GCTATTGAAGTGGTTTCTGTTCCAGATGATCTCATAGCAGATGTAGCCAATATGGTCAAC

CCTGTGCAGCAGTACAGCGGTTTGACTACACAGTCCCTGGAGACGGTGTATCGTGTTAAC

ATGGGTGGTCCGAAGGTCTTCCCGAACAATGATACCCTCTCGAGGACTTGGCAGAAGGAT

CAGAAGTACATACTGAACCCCAGTGTGACCAAAACTGCTGTATATGGCAAGGCTATCAAG

TACAGGAAAGGCGGGGCAACTCCACTGACGGCCCCAGATATTGTGTACAGTACAGCTACA

GAATTGGCGGCTTCAAACACATCCAACGCACTTTTCAACATGACATGGCAGTTTGATGTG

GATGCAGGCTTCAGCTATCTGATAAGATTTCACTTCTGTGATATAGTCAGCAAGGCACTG

AACCAGCTCTACTTCAATGCATATGTGGGAGGCTTCTTTGCACAGCATGATCTTGATCTC

TCAGAGCAATCGGTGAATCAACTGGCCACAGCTATCTATGTTGACGTGGTTCTTTCTTCC

AATGATGCATCTAGCAAGCTCAGCATCAGTATTGGTCCGTCCACCTTGAACAATGCATTG

CCTGATGGGATTCTGAATGGCCTTGAGATTATGAAGATGGGCAGTGGCTCTGGTTCTGCT

TTCACTGTTGGGAACAACGGTTCAAACAAAAAGTTGCCCATAATTATTGGCTCAGTCCTT

GGGGTTGTCGGGCTTCTGATAATTGTCCTTGTTGTGGTACTGCTTTGCCGGAGGAAGAAG

ACCGACGACAAGCAGCACTCCAAGACCTGGATGCCTTTCTCTATCAATGGGCTCACGTCT

CTCAGTACAGGAAGCAGAACTTCCTATGGTACCACACTAACATCAGGTCTGAATGGAAGC

TATGGATATCGCTTTGCCTTCAATGTGCTCCAAGAAGCAACAAACAATTTTGATGAGAGC

TGGGTGATTGGGGTCGGAGGTTTTGGGAAAGTCTACAAGGGTGCCTTGAGGGATGACACA

AAGGTTGCAGTGAAGCGAGGAAACCCCAAGTCCCAGCAAGGTCTCAATGAGTTCCGGACA

GAGATTGAGCTCCTTTCACGTCTGCGTCACCGCCACCTGGTGTCTCTGATTGGGTACTGT

GATGAAAGGAATGAGATGATCTTGGTCTACGAGTACATGGAGAACGGAACCGTCAAGAGC

CACCTGTATGGTTCAGACAACCCCTCACTCAACTGGAAGCAGCGGTTGGAGATCTGCATT

GGAGCAGCAAGGGGGCTACACTATCTTCATACTGGTTCTGCAAAGGCCATTATCCACCGT

GATGTCAAGTCTGCAAACATCTTGCTTGATGAGAATCTCCTTGCCAAAGTCGCCGACTTT

GGGCTGTCAAAGACTGGGCCTGAGCTGGATCAAACTCATGTCAGCACTGCAGTGAAGGGT

AGCTTTGGGTACCTTGACCCTGAATACTTCCGGAGGCAGCAGCTGACTGAGAAGTCGGAC

GTCTACTCCTTCGGTGTTGTCATGCTGGAGGTGCTCTGCGCGAGGCCGGTGATCGACCCT

TCGCTCCCGAGGGAAATGGTGAACTTGGCAGAGTGGGGGATGAAGTGGCAGAAGAGAGGG

GAGCTGCACCAGATCGTCGACCAGAAGCTTTCCGGCGCGATCAGGCCGGACTCTCTGAGG

AAGTTCGGCGAGACGGTGGAGAAGTGCCTGGCCGACTACGGCGTGGAGCGGCCGTCGATG

GGGGACGTCCTCTGGAATTTGGAGTATGTCCTGCAGCTCCAGGATGTGGATTCTTCGACC

GTGTCGGACGTGAACAGCATGAACCGGATCGTCGACCTGTCGTCACAAGTTCAGCATGTG

GGTGCCATGGAGAGCATCAGCGTGACGATGGCGGAGGACGGAGCTTTGCACGAGCCTGAC

CACGACCTCTCCGACGTGTCGATGAGCAGGGTTTTCTCTCAGCTGATCAAAGCTGAGGGG

AGGTGA

>Ta-CrRLK1L15-A CDS sequence

ATGAATTCCTCCGCCAATTTCCTGTCGATCCTGGTGCTGCTGGTGTTCTTGGCCGCGGGG

AATGCGCGAGCGCAGCCCCAGCCGATCCTCATCAACTGCGCCTCGGATTCCACCACCAGC

GTCGACGCCAGGACATGGATTGGGGATTCTTCCCCTTCCAACAACTTCACGCTCAGCTTC

CCCGGAGCCATCGCCTCGGCGGCTCCGGCTCCGGCTCCGGCTCCGGGAGTTGATGGAGAA

CAAGACCCGTACGGAGATTTGTACAAGACCGCCCGTGTCTTCAACGCCTCCTCCAGCTAC

AGGCTCGCCGTCGCCCCCGGGAGCTACTTCCTCCGCCTCCATTTCAGCCAGCAGTTCGCC

AATCTCGGCGCCCAGGAGCCCATCTTCAGTGTCGCGGCAAATGGCCTGAGGCTGCTCTCC

AAGTTCAGTGTCCACGGGGAGATTTCTTGGAGGGATTCCCAGATCAACTCAACCAGCAGC

GTCATCGTCAAGGAGTACCTTCTCAATGTCACTTCTGGTAAACTGGGCATTGAGTTCACC

CCCGATGAAGGGTCCTTCGCCTTCATCAATGCCATGGAGGTTCTCCCTGTGTCTGGCACC

TCTATTTTTGATTCAGTCAACAAGGTGGATGCTCATGGGTTGAAAGGCCCTTTTAGCCTC

GACGGCGGCGGGATCGAGACCATGTACAGGCTGTGTGTGGGATGCAGAGATGTACTGACG

AGGAAAGAGGATCCAGGATTGTGGAGGAGGTGGGATAAAGATGACCATTTCATATTCTCT

CTGAACGCCGCGAATTCCATCTTCAACTCTTCCAACATAAGTTATGTGTCTGCTGATGAT

CCCACGGTAGCCCCTTTGAGGCTCTATCAGAGTGCAAGGGTGCCAACAGAGAGTTCGGTC

TTGGGAAAGAAGTTCAATGTCTCATGGAGCTTTAACATTGACCCTGGCTTTGATTACTTG

GTCCGGCTGCATTTCTGCGAGCTGCAGTATGACAAGGCCGAGCAACGCAAGTTCAAGATT

TACATAAACAACAAGACCGCTGCAGAGGGCTATGATGTGCTTGCCAGAGCTGGGGGCAAG

AACAAGGCCTTTTATGAAGACTTCCTTGATGCTGCCTCACCGCAGATGGACACTCTTTGG

GTTCAGCTGGGGTCTGAGTCTTCAGCAGGTTCCGCGGCTGCTGATGCTCTTCTCAATGGC

ATGGAGATCTTTAAGGTCAGCCGGGAAGGAAATCTTGCCCATCCAACCGTCAGGATTGGA

GGCTTCAGTGGTGGCACAAGCAAACCAAAACGGAGCCCCAAGTGGGTGCTAATTGGTGCT

GCTTCCGGTCTGATAATTTTTATCGCAATTGCTGCTGCTCTTTATTTATGTTTCAATCTG

CGACGGAAGAAAAATAGTTCAGCCAGCAAAGCCAAGGACAATCCCCATGGTGCTGCACAT

ACCCGTTCTCCAACTCTTCTCACGGCTGGGGCATTTGGGAGCAAAAGGATGGGCAGGCGG

TTCACCATTGCAGAAATCAGAACAGCCACTGTGAACTTTGATGAGTCCTTGGTGATTGGG

GTTGGAGGCTTTGGCAAGGTCTACAGGGGTATAATGGAGGATGGCACTCGGGTGGCAATT

AAGAGGGGTTACACAGATTCTCACCAGGGTCAGGGTGTGAAGGAATTCGAAACTGAGATC

GAGATGCTCTCAAGGTTGCGGCACCGGCACCTTGTGCCCTTGATTGGCTATTGTGATGAG

CAAAACGAGATGGTCTTAGTTTATGAGCACATGGCAAATGGCACATTAAGGAGCCATCTT

TATGGAAGTGACCTTCCTGCTCTTACATGGAAGCAAAGGCTCGAAATATGTATCGGCGCA

GCACGAGGGCTTCACTACCTTCACACTGGGCTTGACAGAGGTATAATCCACAGGGACGTC

AAGACTACCAACATTTTGTTAGACAACAACCTTGTTGCCAAGATGGCAGATTTTGGCATC

TCAAAAGATGGTCCAGCTTTAGATCATACTCATGTTAGTACTGCTGTCAAAGGGAGTTTT

GGTTACCTCGATCCAGAGTACTATAGGAGACAGCAGTTAACGCCAAGTTCAGATGTGTAC

TCTTTTGGTGTCGTGCTGTTTGAAGTGCTGTGTGCTCGACCAGTCATAAATCCAACCCTG

CCAAGAGACCAGATAAACCTTGCTGACTGGGCTCTCAACAGGCAAAGGCACAAGTTACTT

GAGACCATAATCGACCTTCGATTGGATGGAAATTACACACTGGAGTCCATCAGAACATTC

AGCGAGATAGCAGAAAAATGCCTTGCAGATGAGGGGGTGAACCGGCCTTCGATGGGCGAA

GTCCTCTGGCACCTAGAGAGTGCTTTGCAGTTGGAACAAGGTCATCTGCAAAGCACAAAT

GGTGATGGTTGTTCAGACCCTCAACTGAAGCCTTCTGATGTACCTACCCATGTGGCGTGC

ATCAAAGAAGTTGAGCAATCCACTCGTCCAGGCTCCCACGATTCAGATGGGCAAGTTGTC

GATGTCAAGATTGAGGTGCCATGA

>Ta-CrRLK1L15-B CDS sequence

ATGAAATCTTCCGCGAATTTCCTGTCGATCCTGGTGCTGCTGGTGTTCCTGGCCGCGGGG

AATGCGCGAGCGCAGCCCCAGCCGATCCTCATCAACTGCGGCTCGGATTCCACCACCAGC

GTCGATGCCAGGACATGGATTGGGGATTCTTCCCCTTCCAACAACTTCACGCTCAGCTTC

CCGGGAGCCATCGCCTCGGCGGCTCCGGCTCCGGGAGTTGATGGAGAACAGGACCCGTAC

GGAGATTTGTACAAGACCGCCCGTGTCTTCAACGCCTCCTCCAGCTACAGGCTCGCCGTC

GCCCCCGGGAGCTACTTCCTCCGCCTCCATTTCAGCCAGCAGTTCGCCAATCTCGGCGCC

CAGGAGCCCATCTTCAATGTCGCGGCAAATGGCCTGAGGCTGCTCTCCAAGTTCAGTGTC

CACGGAGAGATTTCTTGGAGGGATTCCCAGATCAATTCAACTAGCAGCGTCATCGTCAAG

GAGTACCTTCTCAATGTCACTTCTGGTAAACTGGGCATTGAGTTCACCCCCGATGAAGGA

TCCTTCGCCTTCATCAATGCCATGGAGGTTCTACCTGTGTCTGGCACCTCAATTTTTGAT

TCAGTCAACAAGGTGGACGGTCATGGGTTGAAAGGCCCTTTTAGCCTCGACGGCAGCGGG

ATCGAGACCATGTACAGGCTGTGTGTGGGATGCATCGATGTACTGGCGAGGAAAGAGGAT

CCAGGATTGTGGAGGAGGTGGGATAAAGATGAGCATTTCATATTCTCTCTCAACGCCGCG

AGTTCCATCTTCAACTCTTCCAACATAAGTTATGTGTCTGCTGATGATCCCACAGTAGCC

CCTTTGAGGCTCTATCAGAGTGCAAGGGTGCCAACAGAGAGTTCGGTCTTGGGAAAGAAG

TTCAATGTCTCATGGAGCTTTAACATTGACCCTGGCTTTGATTACTTGGTCCGGCTGCAT

TTCTGCGAGCTGCAGTATGACAAGGCTGAGCAACGCAAGTTCAAGATTTACATAAACAAC

AAGACCGCTGCAGAGGGCTATGATGTGTTTGCCAGAGCTGGAGGGAAGAACAAGGCCTTT

TATGAAGACTTCCTTGATGCTGCCTCACCGCAGATGGACACTCTTTGGGTTCAGCTGGGG

TCTGAGTCTTCAGCAGGTTCCGCGGCTGCTGATGCTCTTCTGAATGGCATGGAGATCTTT

AAGGTCAGCCGGGAAGGAAATCTTGCCCATCCAACCGTCAGGATTGGAGGCATCAGTGGC

GGCGCAAGGAAACCAAAACGGAGCCCCAAGTGGGTGCTAATTGGTGCTGCTTCCGGTCTG

ATAATTTTTATCGCAATTGCTGGTGCTCTTTATTTCTGTTTCAATCTGCAAAGGAAGAAA

AATAGTTCGGCCAACAAAGCCAAGGACAATCTCCATGGTGTTACACATACCCGTTCTCCA

ACTCTTCGCACGGCTGGGGCATTTGGGAGCAAAAGGATGGGCAGGCGGTTCACCATTGCA

GAAATCAGAACAGCCACCGTGAACTTTGATGAGTCCTTGGTGATTGGGGTTGGAGGCTTT

GGCAAGGTCTACAGGGGTATAATGGAGGATGGCACTCGGGTGGCAATTAAGAGGGGTTAT

ACAGATTCTCACCAGGGTCAGGGTGTGAAGGAATTCGAAACTGAGATCGAGATGCTCTCA

AGGTTGCGGCACCGGCACCTTGTGCCCTTGATTGGCTATTGTGATGAGCAAAACGAGATG

GTCTTAGTTTATGAGCACATGGCAAATGGCACATTAAGGAGCCATCTTTATGGAAGTGAC

CTTCCTGCTCTTACATGGAAGCAAAGGCTCGAAATATGTATCGGCGCAGCACGAGGGCTT

CACTACCTTCACACTGGGCTTGACAGGGGTATAATCCACAGGGATGTCAAGACTACCAAC

ATTTTGTTGGACGACAACCTTGTTGCCAAGATGGCAGATTTTGGCATCTCAAAAGATGGT

CCAGCTTTAGATCATACTCATGTTAGTACTGCTGTCAAAGGGAGTTTTGGTTACCTCGAT

CCAGAGTACTATAGGAGACAGCAGTTAACGCCAAGTTCAGATGTGTACTCTTTTGGTGTC

GTGCTGTTTGAAGTGCTGTGTGCTCGACCAGTCATAAATCCAACCCTGCCAAGAGACCAA

ATAAACCTTGCTGACTGGGCTCTCAACAGGCAAAGGCACAGGTTACTTGAGACCATAATC

GACCTTCGATTGGATGGAAATTACACACTGGAGTCCATCAAGATATTCAGCGAGATAGCA

GAAAAATGCCTCGCAGATGAGGGGGTGAACCGGCCTTCGATGGGCGAAGTCCTCTGGCAC

CTAGAGAGTGCTTTGCAGTTGGAACAAGGTCATCCGCAAAGCACAAATGGTGATGGTTGC

TCAGACCCTCAACTGAAGCCTTCTGATGTACCTACCCGTGTGGCGTGCATCAAAGAAGTT

GAGCAATCCACTCGTCCAGGCTCCCACGATTCAGATGGGCAAGTTGTTGATGTCAAGATT

GAGGTGCCATGA

>Ta-CrRLK1L15-D CDS sequence

ATGAAATCCTCCGCCAATTTCCTGTCGATCCTGGTGCTGCTGGTGTTCCTGGCCGCGGAG

AATGCGCGGGCGCAGCCCCAGCCGATCCTCATAAACTGCGGCTCGGATTCCACCACCAGC

GTCGATGCCAGGACATGGATTGGGGATTCTTCCCCTTCCAACAACTTCACGCTCAGCTTC

CCCGGGGCCATCGCCTCGGCGGCTCCGGCTCCGGCTCCGGGAGTTGATGGAGAACAAGAC

CCGTACGGAGATTTGTACAAGACCGCCCGTGTCTTCAACGCCTCCTCCAGCTACAGGCTC

GCCGTCGCCCCCGGGAGCTACTTCCTCCGCCTCCATTTCAGCCAGCAGTTCGCCAATCTC

GGCGCCCAGGAGCCCATCTTCAGTGTCGCGGCAAATGGCCTGAGGCTGCTCTCCAAGTTC

AGCGTCCACGGAGAGATTTCTTGGAGGGATTCTCAGATCAACTCAACGAGCAGCGTCATC

GTCAAGGAGTACCTTCTCAATGTCACTTCTGGTAAACTGGGCATTGAGTTCACCCCCGAT

GAAGGGTCCTTCGCCTTCATCAATGCCATGGAGGTTCTACCTGTGTCTGGCACCTCAATT

TTTGATTCAGTCAACAAGGTGGATGCTCATGGGTTGAAAGGCCCTTTTAGCCTCGACGGC

GACGGGATCGAGACCATGTACAGGCTGTGTGTGGGATGCATCGATGTACTGCCGAGGAAA

GAGGATCCAGGATTGTGGAGGAGGTGGGATAAAGATGAGCATTTCATATTCTCTCTCAAC

GCCGCGAATTCCATCTTCAACTCTTCCAACATAAGTTATGTGTCTGCTGATGATCCCACA

GTAGCCCCTTTGAGGCTCTATCAGAGTGCAAGGGTGCCAACAGAGAGTTCGGTCTTGGGA

AAGAAGTTCAATGTCTCATGGAGCTTTAACATTGACCCTGGCTTTGATTACTTGGTCCGG

CTGCATTTCTGCGAGCTGCAGTATGACAAGGCTGAGCAACGCAAGTTCAAGATTTACATA

AACAACAAGACCGCTGCAGAGAGCTATGATGTGTTTGCCAGAGCTGGGGGCAAGAACAAG

GCCTTTTATGAAGACTTCCTTGATGCTGCCTCACCTCAGATGGACACTCTTTGGGTTCAG

CTGGGGGCTGAGTCTTCAGCAGGTTCCGCGGCTGCTGATGCTCTTCTCAATGGCATGGAG

ATCTTTAAGGTCAGCCGGGAAGGAAATCTTGCCCATCCAACCGTCAGGATTGGAGGCATC

AGTGGTGGTGCAAGCAAACCAAAACGGAGCCCCAAGTGGGTGCTAATTGGTACTGCTTCC

GGTCTGATAATTTTTATCGCAATTGCTGGTGGTCTTTATTTTGGTTTCAATCTGCGACGG

AAGAAAAATAGTTCAGCCAGCAAAGCCAAGGACAATCTCCATGGTGCTACACATACGCGT

TCTCCCACTCTTCGCACAGCTGGGGCATTTGGGAGCAACAGGATGGGCAGGCGGTTCACC

ATTGCAGAAATCAGAACAGCCACCGTGAACTTTGATGAGTCCTTGGTGATTGGGGTTGGA

GGCTTTGGCAAGGTCTACAAGGGTATAATGGAGGATGGCACTCGGGTGGCAATTAAGAGG

GGGCATACAGATTCTCACCAGGGTCAGGGTGTGAAGGAATTCGAAACTGAGATCGAGATG

CTCTCAAGGTTGCGGCACCGGCACCTTGTGCCCTTGATTGGCTATTGTGATGAGCAAAAC

GAGATGGTCTTAGTTTATGAGCACATGGCAAATGGCACATTAAGGAGCCATCTTTATGGA

AGTGACCTTCCTGCTCTTACATGGAAGCAAAGGCTTGAAATATGTATCGGCGCAGCACGA

GGGCTTCACTACCTTCACACCGGGCTTGACAGGGGTATAATCCACAGGGATGTCAAGACT

ACCAACATTTTGTTAGACGACAACCTTGTTGCCAAGATGGCAGATTTTGGCATCTCAAAA

GATGGTCCAGCTTTAGATCATACTCATGTTAGTACTGCTGTCAAAGGGAGTTTTGGTTAC

CTCGATCCAGAGTACTATAGGAGACAGCAGTTAACGCCAAGTTCAGATGTGTACTCTTTT

GGTGTCGTGCTGTTTGAAGTGCTGTGTGCTCGACCAGTCATAAATCCAACCCTGCCAAGA

GACCAAATAAACCTTGCTGACTGGGCTCTGAACAGGCAAAGGCACAGGTTACTTGAGACC

ATAATCGACCTTCGATTGGATGGAAATTACACACTGGCGTCCGTCAAGAAATCCAGCAAG

ATAGCAGAAAAATGCCTGGCAGATGAGGGGGTGAACCGGCCTTCGATGGGCGAAGTCCTC

TGGCACCTAGAGAGTGCTTTGCAGTTGGAACAAGGTCATCCGCAAAGCACAAATGCTGAT

GGTTGTTCAGACCCTCAACTGAAGCCTTCTGATGTACCTACCCGTGTGGCGTGCATCAAA

GAAGATGAGCAATCCACTCGTCCAGGCTCCCACAATTCAGATGGGCAAGTTGTTGATGTC

AAGATTGAGGTGCCATGA

Protein sequences:

>Ta-CrRLK1L1-A

MPALAILARSMAPCKRVPMFLILFILSITRVATTNAIASKVDRFVPQDNYLLSCGASAAVQVDDGRTFRSDPESVSFLSTLTDIKIAAKASLASASPLSPLYLDARVFSDISTYSFFISQPGRHWIRLYFLPITDSQYNLTTATFSVSTDSMVLLHDFSFIASPPNPVFREYLVSAQGDNLKIIFTPKKNSIAFINAIEVVSAPPSLIPNTTTRMGPQDQFDISNNALQVVYRLNMGGALVTSFNDTLGRTWQPDAPFLKLEAAAEAAWVPPRTIKYPDDKTLTPLIAPASIYSTAQQMASTNITNARFNITWQMVAEPGFRYLIRLHFSDIVSKTLNSLYFNVYINGMMAVANLDLSSLTMGLAVAYYKDLIAESSSIINSTLLVQVGPNTIDSGDPNAILNGLEIMKISNEASSLDGLFSPKTSSEVSKTTLTGIAFALAATAALAVVICYRRNRKPEWQRTNSFHSWFLPLNSSSSFMSSCSRLSRNRFGSTRTKSGFSSVFASSAYGLGRYFTFVEIQKATKNFEEKGVIGVGGFGKVYLGATEDGTQLAIKRGNPSSDQGMNEFLTEIQMLSKLRHRHLVSLIGCCDENNEMILVYEFMSNGPLRDHLYGDTNIKPISWKQRLEVCIGAAKGLHYLHTGSAQGIIHRDVKTTNILLDENFVAKVADFGLSKDAPSLEQTHVSTAVKGSFGYLDPEYFRRQQLTDKSDVYSFGVVLFEVLCARPAINPALPRDQVNLAEWARTWHRKGELGKIIDPNIAGQIRSDSLEMFAEAAEKCLADYGVDRPTMGDVLWKLEFALQLQEKGDVVDGTSDGIAMKSLEVTNVDSMEKSGNAIPSYVQGR

>Ta-CrRLK1L1-B

MPALAILARSSRMAEWERVPMFLILFILSITSVATTNAIASKVDRFVPQDNYLLSCGASAAVQVDDGRTFRSDPESVSFLSTPTDIKIAAKASLASASPLSPLYLDARVFSDISTYSFFISQPGRHWIRLYFLPITDTQYNLTTATFSVSTESMVLLHDFSFIASPPNPVFREYLVSAQGDNLKIIFTPKKNSIAFINAIEVVSAPPSLIPNTTTRMGPQDQFDISNNALQVVYRLNMGGALVTSFNDTLGRTWLPDAPFLKLEAAAEAAWVPPRTIKYPDDKTLTPLIAPASIYSTAQQMASTNITNAKFNITWVMVAEPGFRYLIRLHFSDIVSKTLNSLYFNVYINGMMAVANLDLSSLTMGLAVAYYKDLIAESSSIINSTLVVQVGPSTIDSGDPNAILNGLEIMKISNEASSLDGLFSPKTSSEASKRTLTGIAFALAATAALAVVICYRRNRKPAWQRTNSFHSWFLPLNSSSSFMSSCSRLSRNRFGSTRTKSGFSSVFASSAYGLGRYFTFVEIQKATKNFEEKGVIGVGGFGKVYLGATEDGTQLAIKRGNPSSDQGMNEFLTEIQMLSKLRHRHLVSLIGCCDENNEMILVYEFMSNGPLRDHLYGDTNIKPISWKQRLEVCIGAAKGLHYLHTGSAQGIIHRDVKTTNILLDENFIAKVADFGLSKDAPSLEQTHVSTAVKGSFGYLDPEYFRRQQLTDKSDVYSFGVVLFEVLCARPAINPSLPRDQVNLAEWARTWHRKGELGKIIDPNIAGQIRPDSLEMFAEAAEKCLADYGVDRPTMGDVLWKLEFALQLQEKGDVVDGASDGIPMKSLEVSNVDSMEKSGNAIPSYVQGR

>Ta-CrRLK1L1-D

MPALAILARSMAECKRVPMFLILFILSITSVATTNAIASKVDRFVPQDNYLLSCGASAAVQVDDGRTFRSDPESVSFLSTPTDIKIAAKASLASASPLSPLYLDARVFSDISTYSFFISQPGRHWIRLYFLPITDSQYNLTTATFSVSTDSMVLLHDFSFIASPPNPVFREYLVSAQGDNLKIIFTPKKNSIAFINAIEVVSAPPSLIPNTTTRMGPQDQFDISNSALQVVYRLNMGGALVTSFNDTLGRTWQPDAPFLKLEAAAEAAWVPPRTIKYPDDKTLTPLIAPASIYSTAQQMASTNITNARFNITWQMAAEPGFRYLIRLHFSDIVSKTLNSLYFNVYINGMMAVANLDLSSLTMGLAVAYYKDLIAESSSIINSTLVVQVGPNTIDSGDPNAILNGLEIMKISNEANSLDGLFSPKTSSEVSKTTLTGIAFALAATAALAVVICYRRNRKPAWQRTNSFHSWFLPLNSSSSFMSSCSRLSRNRFGSTRTKSGFSSVFASSAYGLGRYFTFVEIQKATKNFEEKGVIGVGGFGKVYLGATEDGTQLAIKRGNPSSDQGMNEFLTEIQMLSKLRHRHLVSLIGCCDENNEMILVYEFMSNGPLRDHLYGDTNIKPISWKQRLEVCIGAAKGLHYLHTGSAQGIIHRDVKTTNILLDENFVAKVADFGLSKDAPSLEQTHVSTAVKGSFGYLDPEYFRRQQLTDKSDVYSFGVVLFEVLCARPAINPALPRDQVNLGEWARTWHRKGELGKIIDPNIAGQIRPDSLEMFAEAAEKCLADYGVDRPTMGDVLWKLEFALQLQEKGDVVDGASDGIAMKSLEVTNVDSMEKSGNAIPSYVQGR

>Ta-CrRLK1L2-A

MVLPTLPVTLTFLTLLALLSIAKAADNNSTTSGLILLNCGESTQDDDDGGRSWDGDTGSIFAPSMKGDAAIALGQPPSLTPRVPYTTARIFTSNYTYSFPVSPGRMFLRLYFLSTAYEYYAVSDAVFGVTARNLVLLKDFNALQTAQAITSAYLVREFSVNVSSGSLDLTFAPSAHQYGSYAFVNGIEIVPTPDIFATPDIRFVSGDNTSPFTFDADMSLQTMYRLNVGGPAISPKGDSGFYRSWANDAPYILGGFGLTFWKNDNLTISYTSRVPNYTAPVDVYGTARSMGPTAQINLNYNLTWILPVDAGFFYLLRFHFCEIKYPITKVNQRSFFIYINNQTAQEQMDVIFRSGGIGRPTYTEYVIMAIGSGQVDMWIALHPDLSSKPQYSDAILNGLEVFKLQNYGPSNLAGLNPPLPQKPDVNPNRLSSGERKTKGGIQATIGGTAGGFALLLIALFSMCVIYRRKKAAKSPGKTDYGHVKHPTKCIKSTCDLVRHFSFAKIQVATKDFDEALIIGRGGFGNVYIGDIDGGTKVAIKRCDQKSQQGFHEFQTEIEMLCNFRHRHLVSLIGYCEEKNEMILVYDYMAHGTLREHLYNTRNPPLPWQQRLEICIGAARGLHYLHTGVEQGIIHRDVKTTNILLDDRLMAKVSDFGLFKASPDIGNTHMSTAVKGTFGYLDLEYFRQQRLTKKSDVYSFGVVLFETLCARPVINTELPYEQVSLRDWVVSCRKKGVLEEIVDPCVKEEITLECFRIFAEIAEKCVADRSIDRPSMGDVLWNLEVALQLQDSASYNTSCAEGASSLQISGVHSGKPSTNSTISVAAQEAIFSDIAHPEGR

>Ta-CrRLK1L2-B

MVLPTLPVTLTFLTLLALLSIAKAADNNSTTSGLILLNCGSSTQNDDDSGRTWDGDTGSKFAPSMKGVAAIALGQTPSLTPRVPYTTARIFTSNYTYSFPVSPGRMFLRLYFFSTAYEYYAVSDAVFGVTSRNLVLLNDFNALQTAQAITSAYLVREFSVNVSSGSLDLTFAPSAQQYGSYAFVNGIEIVPTPDIFATPDIRLVSGDNTSPFTFDADMSLQTMYRLNVGGPAISTEGDSGFYRSWANDAQYILGGSGLTFWKNDNLTISYTSRVPNYTAPVDVYGTARSMGPTAQINLNYNLTWIFPVDAGFFYLLRFHFCEIKYPITKVNQRSFFIYINNQTTQKQMDVIVRSGGIGRPTYTEYVIMAIGSRQVDMWIALHPDLSSKPQYSDAILNGLEVFKLQNYGPSNLAGLSPPLPQKPDVNPTRLSNGERKSKGGIQAIIGGTTGGFALLLIALFSMCVIYRRKKVAKSPGKTDYGHVKHPTKCIKSTCDLVRHFSFAKIQVATKDFDEALIIGRGGFGNVYIGDIDGGTKVAIKRCDQKSQQGFHEFQTEIEMLCNFRHRHLVSLIGYCEEKNEMILVYDYMAHGTLREHLYNTRNPPLPWQQRLEICIGAAQGLHYLHTGVEQGIIHRDVKTTNILLDDRLMAKVSDFGLSKASPDIGNTHMSTAVKGTFGYLDPEYFRLQRLTKKSDVYSFGVVLFETLCARPVINTELPYEQVSLRDWALSCWKNGVLEEIVDPRVKEEITPECFRVFAEIAEKCVADRSIERPSMGDVLWNLEVALQLQQASASYNSNRAEGASSLQISAVHSDKPSTNSTISIAAQEAIFSDIAHAEGR

>Ta-CrRLK1L3-A

MQTSCSKLIRWSPQFFDSGAPTAANSKMAFPALPATLTCLTLLALLSLAMAADNNSTGLILINCGASVQEDDDNGRTWDGDTGSKFAPSLKGVTATAPNQDPSLPSTVPFMTARIFASNYTYSFSVTPGRVFLRLYFYPVAYPNYAVADAFFSVTTPNLVLLNDFNASQTVQAISSAYLVREFSVNVSSGSSLDLTFAPSAHHNGSYAFVNGIEIVSTPDIFTAPDTRYVGDNTSPFTFDSAMAVQTMYRLNVGGQAISPKGDSGFYRSWANDAPYIFGGSGVTFSKDDNLTITYTSKVPNYTAPVDVYGTARSMGPTAPINLNYNLTWILPVDAGFSYLLRFHFCEIQYPITKQNQRSFFIYINNQTAQEQMDVIVWSGGIGRTTYTDYVILTAGSGQVDMWIALHPDLSSRPEYFDAILNGLEIFKLQNYGASNNLAGLNPPLPQKPADASPGAASGKVKSVAAIIGGAVGGFVVLLVTCFGICIICKRKNKSKKKKKISKDPGGKSEDGHWTPLTEYSGSRSAMSGNTATTGSTLPSNLCRHFTFAELQTATKNFDQAFLLGKGGFGNVYLGEIDSGTKVAIKRCNPMSEQGVHEFQTEIEMLSKLRHRHLVSLIGYCEDKSEMILVYDYMAHGTLREHLYNTKNPPLSWKQRLEICIGAARGLYYLHTGVKHTIIHRDVKTTNILLDDKWVAKVSDFGLSKTGPNMDATHVSTVVKGSFGYLDPEYFRRQQLSEKSDVYSFGVVLFEVLCARPALSPSLPKEQISLADWALRCQKQGVLGQVIDPVLQGKIAPQCFLKFTDTAEKCVADRSVDRPSMGDVLWNLEFALQLQESEEDTGSLTEGTLSSSGASPLVMTRLQSDEPSMDASTTTTSTTTMSMTGRSIASMDSDGLTPSAVFSQIMHPDGR

>Ta-CrRLK1L3-B

MQTSCSKLIRWSPQFFDSGTPTTAKSKMAFPALPVTLTCLTLLALLSLAMAADNNSTGLILVNCGASTQEADDSGRTWVGDTGSKFAPLLKGVATTAPNQDPSLPSTVPFMTARIFTSNYTYSFSVNPGRMFLRLYFYPVAYANYAVSDAFFSVTTRNLVLLNDFSASQTAQAITSAFLVREFSVNVSSGSSLDLTFAPSAHRNGSYAFVNGIEIVPTPDIFTAPDTRYVGDNTAPFSFDAGMAVQTMYRLNVGGQAISPKGDSGFYRSWANDAPYIFGGSGVTFSKDDNLTITYTSNVPNYTAPVDVYGTARSMGPTAQINLNYNLTWILPVDAGFSYLLRFHFCEIQYPITKQNQRSFFIYINNQTAQEQMDVIVWSGGIGRTAYTDYVIMAVGSGQVDMWIALHPDLSSKPEYFDAILNGLEIFKLQNYGSPNNLSGLNPPLPQKPTDASPGAASGKMKSVAAIIGGAVGGFAVLLVTCFGVCIICKRKNKKNKKKISKDPGGKSEDGHWTPLTEYSGSRSAMSGNTATTGSTLPSNLCRHFTFAELQTATKNFDQAFLLGKGGFGNVYLGEIDSGTKVAIKRCNPMSEQGVHEFQTEIEMLSKLRHRHLVSLIGYCEDKSEMILVYDYMAHGTLREHLYNTKNPPLSWKKRLEICIGAARGLYYLHTGVKHTIIHRDVKTTNILLDDKWVAKVSDFGLSKTGPNMDATHVSTVVKGSFGYLDPEYFRRQQLSEKSDVYSFGVVLFEVLCARPALSPSLPKEQISLADWALRCQKQGVLGQVMDPVLQGKIAPQCFLKFTDTAEKCVADRSVDRPSMGDVLWNLEFALQLQESEEDTGSLTEGTLSSSGASPLVMTRLQSDEPSTDASTTTTTTTTMSMTGRSIASVDSDGLTPSAVFSQIMHPDGR

>Ta-CrRLK1L3-D

MAFPALPVTLTCLILLSLLSLAMAADNNSTGLILVNCGASVQGDDDSGRTWDGDTGSKFAPSLKGVAATAPNQDPSLPSTVPFMTARIFTSNYTYSFSVKPGRMFLRLYFYPVAYPNYAVSDAFFSVTTPKLVLLNDFSASQTAQAITSAFLVREFSVNVSSGSSLDLTFAPSAHRNGSYAFVNGIEIVPTPDIFTAPDTRNVGDNTAPFSFDTSSSLQTMYRLNVGGQAISPKGDLGGFYRSWANDAPYIAGGSGVTFSKDDNLTITYTSKVPKYTAPPDVYGTARSMGPTAQINLNYNLTWILPVDAGFFYLLRFHFCEIQYPIIKINQRSFFIYINNQTAQEQMDVIVWSGGIGRTTYTDYVIMAAGFGQVDMWIALHPDLSSRPEYFDAILNGLEVFKLQNYGSPNNLSGLNPPLPQKPADASPSAASGKMKSVAAIIGGAVGGFIVLLAACFGVCIICKRKNKKKKKKKTSKDPGGKSEDGHWTPLTEYSGSRSAMSGNTATTGSTLPSNLCRHFTFADLQTATKNFDQAFLLGKGGFGNVYLGEIDSGTKVAIKRCNPMSEQGVHEFQTEIEMLSKLRHRHLVSLIGYCEDKSEMILVYDYMAHGTLREHLYNTKNPPLSWKQRLEICIGAARGLYYLHTGVKHTIIHRDVKTTNILLDDKWVAKVSDFGLSKTGPNMDATHVSTVVKGSFGYLDPEYFRRQQLSEKSDVYSFGVVLFEVLCARPALSPTLPKEQISLADWALRCQKQGVLGQVIDPVLQGKIAPQCFLKFTDTAEKCVADRSVDRPSMGDVLWNLEFALQLQESEEDTGSLTEGTRSSSGASPLVMTRLQSDEPSTDASTTTTSTTTMSMTGRSIASMDSDGLTPSAVFSQIMHPDGR

>Ta-CrRLK1L4-A

MAATARLRRARPRGVLGLVSALLVCGAAAYAPEDNYLVSCGSSLDTPVGRRLFLADDGGSGSGAVTLTSPRSAAVKASPDLVSGFRDAALYQNARVFSAPSSYSFAIRRRGRHFLRLHFFPFVYRSYDLAAAARAFKVSTQDAVLLEDGVPAPEPGNASTSSSPQPARVEFLLDVARDTLVVSFVPLVDGGIAFVNAVEVVSAPDGLVADAAESSTGRPEPIPAALPLQTAYRLNVGGPAVAPDDDALWREWTTDLRFLSHSVADAVTREVRYNGTPNRLPGQATATDAPDVVYATARELVINSSSFDGQKQMAWQFDVDASSSYFIRFHFCDIVGKAPHQLHINAYVDDASHATVLTDLDLAAVGDGALAFPYYKDFVLPASEASGKLAVHVGPLANKIVMPAAILNGIEIMKMHLSAGSVVVVEPAAGAAKSRFAVLLGSVCGPLAFVSIAVALAIVLRKKRKKEGEEEEESDKKQPTPTQSQSSTPWMPLLGRLSVRGAIASGSSSFTTAGNTPGTSPRAAAAVMPSYRFPLAVLQDATRNFDDSLIIGEGGFGKVYGAVLQDGTKVAVKRASPESRQGAREFRTEIELLSGLRHRHLVSLVGYCDEREEMILLYEYMEHGSLRSRLYGRGGAAPLSWAQRLEACAGAARGLLYLHTAVDKPVIHRDVKSSNILLDGDLTGKVADFGLSKAGPVLDETHVSTAVKGSFGYVDPEYCRTRQLTAKSDVYSLGVVLLEAVCARPVVDPRLPKPMSNLVEWGLHWQGRGELEKIVDRRIAAAARPAALRKYGETVARCLAERGADRPAMEDVVWNLQFVMRLQEGDGLDFSDVSSLNMVTELTPPPRRQRSAVDSDGLALSDVSSLNMVTELTPPQTGSVEGDGVADDDFTDASMRGTFWQMVNVRSR

>Ta-CrRLK1L4-B

MAATARLRPARARGVLWSVSVWLVCGAAAYAPEDNYLVSCGSSLDTPVGRRLFLADDGGSGSGSGAVTLTSPRSAAVKAPPDLVSGFRDAALYQNARVFSAPSSYSFAIRRRGRHFLRLHFFPFVYRSYDLAVAARAFKVSTQDAVLLEDGVPAPEPGNASTSTSPQPARVEFLLDVARDTLVVSFVPLVDGGIAFVNAVELVSVPDDLVADAADSSTGRPEPIPAALPLQTAYHLNVGGPAVAPDDDALWREWTTDQPLSDPRVDAVTREVRYNRTLNRLPGQATATDAPDIVYATARELVINRSSFDGQKQMAWQFDVDAGSSYFIRFHFCDIVSEAPHQLHINAYVDDASHATVLTDLDLAAVGDGALAFPYYKDFVLPASEASGKLAVHVGPLANKIVMPAAILNGIEIMKMHLSAGSVVVVEPAAGAAKSRLAVILGSVCGALAFVSIAIALAIVLRKKKGEGEEGVKEQPTPTRSQSSTPWMPLLGRLSVRGAIASGSSSFTTAGNTPGASPRAAAAAAAAVVPSYRFPLAMLQDATRNFDDSLIIGEGGFGKVYGAVLQDGTKVAVKRASPESRQGAREFRTEIELLSGLRHRHLVSLVGYCDEREEMILLYEYMEHGSLRSRLYGRGGAAPLSWAQRLEACAGAARGLLYLHTAVDKPVIHRDVKSSNILLDGDLTGKVADFGLSKAGPVLDETHVSTAVKGSFGYVDPEYCRTRQLTAKSDVYSLGVVLLEAVCARPVVDPRLPKPMSNLVEWGLHWQGRGELEKIVDRRIAAVARPAALRKYGETVARCLAERGADRPAMEDVVWNLQFVMRLQEGDGLDFSDVSSLNMVTELTPPRRQRSAVDCDGLDFSDVSSLNMVTELTPPKTGSMEGDGVADDDDFTDASMRGTFWQMVNVRSR

>Ta-CrRLK1L4-D

MAATARLRPARARGVLWVVSVLLVCGAAAYKPEDNYLVSCGSSLDTPVGRRLFLADDGASGAVTLTSPRSAAVKAPPDLVSGFRDAALYQNARVFSAPSSYSFAIRRRGRHFLRLHFFPFVYRSYDLAAAARAFKVSTQDAVLLEDGIPAPEPGNASTSTSPQPARLEFLLDVARDTLVVSFVPLADGGIAFVNAVELVSVPDGLVADAADSSTGRPEPIPAVLPLQTAYRLNVGGPAVAPDDDALWREWTTDLRFLSHSVADAVTREVRYNGMLNRLPGQATATDAPDIVYATARELVINGSSFDGQKQMAWQFDVDTSSSYFIRFHFCDIVGKAPHQLHINAYVDDATVKQDLDLAAVGDGALAFPYYTDFVLPASEASGKLAVHVGPLANKIVMPAAILNGIEIMKMHLSAGSVVVVQPAAGAAKSRFAVVLGSVCGALAFISVAVALAIVLRKKEKEKEVEEGAKEQPTPTQSQSSTPWMPLLGRFSVRGAIASGSSSFTTAGNTPGASPRAAAAAAVMPSYRFPLAMLQDATRNFDDSLIIGEGGFGKVYGAVLQDGTKVAVKRASPESQQGAREFRTEIELLSGLRHRHLVSLVGYCDEREEMILLYEYMEHGSLRSRLYGRGGAAPLSWAQRLEACAGAARGLLYLHTAVDKPVIHRDVKSSNILLDGDLAGKVADFGLSKAGPVLDETHVSTAVKGSFGYVDPEYCRTRQLTAKSDVYSLGVVLLEAVCARPVVDPRLPKPMSNLVEWGLHWQGRGELEKIVDRRIAAAARPAALRKYGETVARCLAERGADRPAMEDVVWNLQFVMRLQEGDGLDFSDVSSLNMVTELTPPRRQRSAVDHDGLDYSDVNSLNMVTELTPPQTGSVEGDGEADDDFTDASMRGTFWQMVNVRSR

>Ta-CrRLK1L5-A

MRGGPRCALLLLVAAAALVPAARAQGATAPAPSAGAPFVPRDDILLDCGATGKGNDTDGRQWDGDAGSKYAPPNLASASAGAQDPSVPQVPYLTARVSAAPFTYSFPLGPGRKFLRLHFYPANYSNRDAADAFFSVSVPAAKVTLLSNFSAYQTTTALNFAYIVREFSVNVTGQNLDLTFTPEKGHPNAYAFINGIEVVSSPDLFGIATPQFVTGDGNSQPYEMDPAAALQTMYRLNVGGQAISPSKDSGGARSWDDDTPYIYGAGAGVSYQNDPNVTIIYPDNVPGYVAPSDVYATARSMGPDKGVNMAYNLTWILQVDAGYQYLVRLHFCEIQSPFTKPNQRVFNIYLNNQTAIEGADVIQWADPNGIGTPVYKDYVVSTVGSGILDFWVALHPDAETKPQYYDAILNGMEVFKLQLTNGSLVGLNPVPSADPPAHSGSGDKKSKVAPIVGGVIGGLAVLALGYCCFICKRRRKAAKASGMSDGHSGWLPLSLYGHSHTSSSAKSHATGSYASSLPSNLCRHFSFAEIKAATKNFDESRILGVGGFGKVYHGEIDGGTTKVAIKRGNPLSEQGIHEFQTEIEMLSKLRHRHLVSLIGYCEEKNEMILVYDYMAHGTLREHLYKTQNAPLSWRQRLEICIGAARGLHYLHTGAKHTIIHRDVKTTNILLDEKWVAKVSDFGLSKTGPSMDHTHVSTVVKGSFGYLDPEYFRRQQLTEKSDVYSFGVVLFEVLCARPALNPTLAKEEVSLAEWALHCQKKGILDQIVDPYLKGKIVPQCFKKFAETAEKCVADNGIERPSMGDVLWNLEFALQMQESAEESGSIGCGMSDEGTPLVMVGKKDPNDPSIDSSTTTTTTTSLSMGDQSVASIDSDGLTPSAVFSQIMNPKGR

>Ta-CrRLK1L5-B

MRGGPRCALLLLAALAAAAALAPAAWAQGATAPAPSAGPPFVPRDDILLDCGATGKGNDTDGRQWDGDAGSKYAPPNLASATAGAQDPSVPQVPYLTARVSAAPFTYSFPLGPGRKFLRLHFYPANYSNRDAADAFFSVSVPAAKVTLLSNFSAYQTTTALNFAYIVREFSVNVTGQNLDLTFTPEKGHPNAYAFINGIEVVSSPDLFDLATPQLVTGDGNSQPYEMDPAAALQTMYRLNVGGQAISPSKDSGGARSWDDDTPYIYGAGAGVSYQNDPNVTITYPDNVPGYVAPSDVYATARSMGPDKGVNMAYNLTWILQVDAGYQYLVRLHFCEIQSPFTKPNQRVFNIYLNNQTAMEGADVIQWADPNGIGTPVYKDYVVSTVGSGIMDFWVALHPDAGTKPQYYDAILNGMEVFKLQLTNGSLVGLNPVPSADPPAHSGSGDKKSKVAPIVGGVIGGLAVLALGYCCFICKRRRKAAKASGMSDGHSGWLPLSLYGHSHTSSSAKSHATGSYASSLPSNLCRHFSFAEIKAATKNFDESRILGVGGFGKVYHGEIDGGTTKVAIKRGNPLSEQGIHEFQTEIEMLSKLRHRHLVSLIGYCEEKNEMILVYDYMAHGTLREHLYKTQNAPLSWRQRLEICIGAARGLHYLHTGAKHTIIHRDVKTTNILLDEKWVAKVSDFGLSKTGPSMDHTHVSTVVKGSFGYLDPEYFRRQQLTEKSDVYSFGVVLFEVLCARPALNPTLAKEEVSLAEWALHCQKKGILDQIVDPYLKGKIVPQCFKKFAETAEKCVADNGIERPSMGDVLWNLEFALQMQESAEESGSIGCGMSDEGTPLVMVGKKDPNDPSIDSSTTTTTTTSLSMGDQSVASIDSDGLTPSAVFSQIMNPKGR

>Ta-CrRLK1L5-D

MRGGPRCALLLLVAAAALVPAARAQGATAPAPSSGVPFVPRDDILLDCGATGKGNDTDGRQWDGDAGSKYAPPKLASASAGAQDPSVPQVPYLTARVSAAPFTYSFPLGPGRKFLRLHFYPANYSNRNAADAFFSVSVPAAKVTLLSNFSAYQTTTALNFAYIVREFSVNVTGQNLDLTFTPEKGHPNAYAFINGIEVVSSPDLFDLATPQLVTGDGNSQPYEMDPAAALQTMYRLNVGGQAISPSKDSGGARSWDDDTPYIYGAGAGVSYQNDPSVAITYPDNVPGYVAPSDVYATARSMGPDKGVNMAYNLTWILQVDAGYQYLVRLHFCEIQSPYTKPNQRVFNIYLNNQTAMQGADVIQWADPNGIGTPVYKDYVVSTVGSGIMDFWVALHPDAETKPQYYDAILNGMEVFKLQLTNGSLVGLNPVPSADPPAHSGSGDKKSLVAPIVGGVIGGLAVLALGYCCFICKRRRKAAKASGMSDGHSGWLPLSLYGHSHTSSSAKSHATGSYASSLPSNLCRHFSFAEIKAATKNFDESRILGVGGFGKVYHGEIDGGTTKVAIKRGNPLSEQGIHEFQTEIEMLSKLRHRHLVSLIGYCEEKNEMILVYDYMAHGTLREHLYKTQNAPLSWRQRLEICIGAARGLHYLHTGAKHTIIHRDVKTTNILLDDKWVAKVSDFGLSKTGPSMDHTHVSTVVKGSFGYLDPEYFRRQQLTEKSDVYSFGVVLFEVLCARPALNPTLAKEEVSLAEWALHCQKKGILDQIVDPYLKGKIVPQCFKKFAETAEKCVADNGIERPSMGDVLWNLEFALQMQESAEESGSIGCGMSDEGTPLVMVGKKDPNDPSIDSSTTTTTTTSLSMGDQSVASIDSDGLTPSAVFSQIMNPKGR

>Ta-CrRLK1L6-A

MAVHVVLPLLLLLLVATVLPYTALAAFSPDFKIFLACGAGADVPFPSDNPARTFVRDDGYLSQGGAAAVSASASSNAASPLYAAARADTSAFSYRLTYPAAPDASSFLVLRLHFFPFVPASSSSTSLSSARFTVSVLDAYALLPAFSPPADGVVKEFFVPRGASGGDFTVRFAPEAGSSAFVNAVELFSAPPELLWNNTAVPVDPVGSNDLPEWPLDALETVYRLNVGGPLLTNGNDTLWRTWLPDDPYLFGAPGQSVVNNTPSPIIYAPSNGYTQEVAPDVVYKTQRAANVTDLLQATTPGFNFNVTWTFPADQGSRYLVRLHFCDYEVVSSVVGTVIVFNVYVAQAIGTPDLTPSARARQSNEAFYIDYAAMAPRAGNLTVSIGRSKKSSKGGILNGLEIMKLQTVNLSSTGSHGRTKRIVIIVLATVLGAAVLASAVLCFFVVRRRKRRQVAPPGSTEDKESTQLPWSPYTQEGISGWADESANRSSEGTTARMQRVSTKLHISLAELKAATDNFHDRNLIGVGGFGNVYKGALADGTPVAVKRAMRASKQGLPEFHTEIVVLSGIRHRHLVSLIGYCNEQAEMILVYEYMEKGTLRGHLYGGSDDEPPLSWKQRLEICIGAARGLHYLHSGYSENIIHRDVKSTNILLGTDGGGSTGGGAIIAKVADFGLSRIGPSLGETHVSTAVKGSFGYLDPEYFKTQQLTDRSDVYSFGVVLFEVLCARPVIDQSLDRDQINIAEWAVRMHGEGKLDKIADARIAGEVNDNSLRKFAETAERCLADYGADRPSMGDVLWNLEYCLQLQETHVNRDAFEDSGAVATQLPADVVVPRWVPSSTSLLMMDDADETGLSMTDLADSQVFSQLNARGEGR

>Ta-CrRLK1L6-B

MAVHVVPPLLLLLATALPYSALAVFSPDFSFFLACGAGADVTFPSDNPTRTFVRDDGYLSQGRPAAVSANASSGAASNPLYAAARADSSAFSYRLAYPATAGASSFLVLRLHFFPFVPASSSTSLSSARFTVSVLDAYALLPAFSPPADGVVKEFFVPRGGSKEFTIRFSPDAGSSAFVNAVELFPAPQQLLWNGSNSVVPVGVLGNDDLAQWQLDALETVYRLNVGGPKVTRENDTLWRTWLPDGAYLFGAPGQSVVNNTSSPIIYNPPNTREVAPDVVYRTQRAANVTDFLRATTPGLNFNVTWTFPAEAGSRYLVRLHFCDYEVVSSVVGVGIVFNVYVAQAIGSRDLAPNAQATQPNEPLYLDYAATAPRAGNLTVSIGTSSKSSGGGILNGLEIMKLQSVDLSSPGSHALTKRSIIIIVLATVLGAAVLACAVLCFFVVRRRKRRQVAPPASKEDKESTQLPWSPYTQEGISGWADESTNRSNEGTTARMQRVSTKLHISLPELKAATDNFHERNLIGVGGFGNVYKGALSDGTPVAVKRAMRASKQGLPEFQTEIVVLSGIRHRHLVSLIGYCNEQAEMILVYEYMEKGTLRSHLYGSDEPVLSWKQRLEICIGAARGLHYLHSGYSENIIHRDVKSTNILLGTDDGGSTGGGAIIAKVADFGLSRIGPSLGETHVSTAVKGSFGYLDPEYFKTQQLTDRSDVYSFGVVLFEVLCARPVIDQSLDRDQINIAEWAVRMHGEGKLDKIADARIAGEVNDNSLRKFAETAEKCLADYGADRPSMGDVLWNLEYCLQLQETHVNRDAFEDSGAVATQLPADVVVPRWVPSSTSLLMMDDADETGLSMTELADSQVFSQLNARGEGR

>Ta-CrRLK1L6-D

MAVHVVPPLLLLLLATALPYTALAVFSPDFSFFLACGAGADVPFPSDNPTRTFVRDDGYLSQGRPAAVSASASSGAASNPLYAAARADSSAFSYRLAYPATAGASSFLVLRLHFFPFVPASSSTSLSSARFTVSVLDAYALLPTFSPPVAGVVKEFFVPRDGSKDFTIRFTPDAGSSAFVNAVELFSAPPELLWNNTAVPVDPVGSNDLPEWPLDALETVYRLNVGGPMVTKENDTLWRTWLPDGPYLFGAPGQSVVNSTSSPIMYDPSNGYTQDVAPDVVYRTQRAANVTDLLVATTPGLNFNVTWTFPAEQGSRYLVRLHFCDYEVVSSVVGVGIVFNVYVAQAIGTPALSPKDRARQSNEAFYMDYAARAPRAGNLTVSIGWLRQSSGGGILNGLEIMKLQSADPSLTVSHGLTKRSIIIIVLATVLGAAVLACAVLCFFVVRRTKRRQVAPPASTEDKESTQLPWSPYTQEGISGWADESTNRSSEGTTARMQRVSTKLHISLAELKAATDNFHDRNLIGVGGFGNVYKGALADGTPVAVKRAMRASKQGLPEFHTEIVVLSGIRHRHLVSLIGYCNEQAEMILVYEYMEKGTLRSHLYGGSDDEPPLSWKQRLEICIGAARGLHYLHCGYSENIIHRDVKSTNILLGTDDGGSTGGGAIIAKVADFGLSRIGPSLGETHVSTAVKGSFGYLDPEYFKTQQLTDRSDVYSFGVVLFEVLCARPVIDQSLDRDQINIAEWAVRMHGEGKLDKIADARIAGEVNDNSLRKFAETAERCLADYGADRPSMGDVLWNLEYCLQLQETHVNRDAFEDSGAVATQLPADVVVPRWVPSSTSLLMMDDADEAGLSMTELADSQVFSQLNARGEGR

>Ta-CrRLK1L7-B

MLWNSSVTPVGAVVKDDMDLWQRQPLETVYRLNVGGPKVTIENDTLWRTWLPDGPYLYDASGLSVVSNTSNPIIYDSSNGYTREVAPDVVYQTQRMANVTDLLAATTPGLNFNLTWTFPAVKGSHYLVRLHFCDYEVVSSVVGVGIVFNVYIAQTIGTPDLTPNARATQSNEVFYMDYAARAPSTGNLTVSIGWSSKRSGGGILNGLEIMRLPPVDLSSRRYGRTKRTIVITVSAVLGAAVLACVVLCFFGVPYTKYSGSGWAEQFTNRWSREGKTSGLQSVSTKLHIALAKIKAATDNFHERNLIGVGGFGNVYKGVLVDGTPVAVKRAMRASQQGLPEFQTEIVVLSGIRHRHLVSLIGYCNEQAEMILVYEYMEKGTLRSHLYGSDEPALSWKQRLEICIGAARGLHYLHRGYAENIIHRDVKSTNILLGSDGGSTGGVIAKVADFGLSRIGPSFGETHVSTAVKGSFGYLDPGYFKTQQLTDRSDVYSFGVVLLEVLCARPVIDQSLDHGRINIAEWAVRMRREGRLDKMADPRIAGEVDEESLLKFAETAEKCLAECWVDRPSMGDVLWNLEYCLQLQETNITGDGLDDMVPSSTSLLMDETDLSMTNVADSKVFSQLSARGEGR

>Ta-CrRLK1L7-D

MAVRGILLALLLAMVLPRAILAAFSPGFQYFLACGANSAVSFPSDSPANIFVPDAAYLSPAGAPAVSASSTLASPPALYAAARADISAFSYRLPSPASPDTSSFLVLRLHFFPSFPATSSQYVINILSARFNVSVADAYALLSSFSPPAAGVVKEFFVPRDLFDGHFHVTFTPDAGSTAFVNAIELFSAPPEMLWNGPVTPVGAVVKDDMDLWQRQPLETVYRLNVGGPKVIIENDTLWRTWLPDGPYLYDASGLSVVSNTSSPIIYDSSNGYTREVAPDVVYQTQRMANVTDLLAATTPGLNFNLTWTFPAVKGSRYLVRLHFCDYEVVSSVVGVGIVFNVYIAQAIGTPDLTPNARATQSNEVFYKDYAARAPSAGNLTVSIGWSSKSSGGGILNGLEIMRLPPVDLSSRRYGRTKRTIVITVSAVLGAAVLACVVLCFFGVPYTKYSGSGWAEQFMNRWSRERKTGGMESVSRKLHIALAKIKAATDNFHERNLIGVGGFGNVYKGVLGDGTPVAVKRAMRASQQGLPEFQTEIVVLSGIRHRHLVSLIGYCNEQAEMILVYEYMEKGTLRSHLYGSDEPALSWKQRLEICIGAARGLHYLHRGYAENIIHRDVKSTNILLGSDDGSTGGVIAKVADFGLSRIGPSFGETHVSTAVKGSFGYLDPGYFKTQQLTDRSDVYSFGVVLLEVLCARPVIDQSLDHGRINIAEWAVRMRREGRLDKMADPRIAGEVDEESLLKFAETAEKCLAEAWVDRPSMGDVLWNLEYCLQLQETNITGDELDDMVPSSTSLLMDETGLSMTNVADSKVFSQLSARGEGR

>Ta-CrRLK1L8-A

MPPVPDMLVRLLVASVLLGAASGAFTPADTYLVLCGTSASATVAAGRTFVGDARLPAKSLAAPQSVEANTSLTAVVPSGESQLYRSARVFTAPASYTFAVKQPGRHFVRLHFFPFPYRSYDMAADAAFNVSVQGAVLVNGYAPKNGTAELREFSLNVTGATLVIAFAPTGKLAFVNAIEVVSVPDELIADTARMVGGAVQYTGLSTQALETIHRINMGVPKITPGNDTLGRTWLPDQSFQLNTNLAQHKDAKPLTIKYDEKSALSSAYTAPAEVYATATRLSTAGETSTINVQFNISWRFDAPAGSDYLLRFHWCDIVSKAAMGMAFNVYVGGSVVLENYEISRDTFNRLSIPVYKDFLLGAKDAKGAITVSIGSSTEDNALPDGFLNGLEIMRVVGSAGAGAPAASARSSKVKIGIIAGSAVCGATLVMVLGFIAFRTLRGREPEKKQPSDTWSPFSASALGSRSRSRSFSKSNGNTVLLGQNGAGAGYRIPFAALQEATDGFDEAMVIGEGGFGKVYKGTMRDETLVAVKRGNRRTQQGLHEFHTEIEMLSRLRHRHLVSLIGYCDERGEMILVYEYMAMGTLRSHLYGAGLPPLSWEQRLEACIGAARGLHYLHTGSAKAIIHRDVKSANILLDESFMAKVADFGLSKNGPELDKTHVSTKVKGSFGYLDPEYFRRQMLTEKSDVYSFGVVLLEVLCARTVIDPTLPREMVSLAEWATPCLRNGRLDQIVDQRIAGTIRPGSLKKLADTAEKCLAEYGVERPTMGDVLWCLEFALQLQVGSSDGSDVDTMLPPAPPVPVKTPEVQRSLSAATMATDAAAMTTNLGDLDGMSLSGVFSKMIKSDEVR

>Ta-CrRLK1L8-B

MPPFPDMLVRLLLASVLLSAASGAFTPADNYLVICGTSASATVAPRRSFVGDARLPAKSLAAPQSVEANTSLTAVVPSGESELYRSARVFTAPASYTFAVKQPGRHFVRLHFFPFSYRSYDMAADAAFNVSVQGAVLVNGYTPKNGTAELREFSVNVTGGTLVIAFAPTGKLAFVNAIEVVSVPDELIADTARTVGGAVQYTGLSTQALETIHRINMGIPKITPGNDTLGRTWLPDQSFQLNTNLAQHKDAKPLTIKYDEKSALSSPFTAPAEVYATATRLSTAGETSTINVQFNISWRFDAPAGSDYLLRFHWCDIVSKAAIGMAFNVYVGGSVVLDNYEISRDTFNRLSIPVYKDFVLGAKDAKGAITVSIGSSTEDNTLPDGFLNGLEIMRVVGSASAGAGAPAASPPSSKVKIGIIAGSAVCGATLVTVLGFIAFRMLRGREPEKKQPSDTWSPFSASALGSRSRSRSFSKSNGNTVLLGQNGAGAGYRIPFAALQEATGGFDEGMVIGEGGFGKVYKGTMRDETLVAVKRGNRRTQQGLHEFHTEIEMLSRLRHRHLVSLIGYCDERGEMILVYEYMAMGTLRSHLYGAGLPPLSWEQRLEACIGAARGLHYLHTSSAKAIIHRDVKSANILLDESFMAKVADFGLSKNGPELDETHVSTKVKGSFGYLDPEYFRRQMLTEKSDVYSFGVVLLEVLCARTVIDPTLPREMVSLAEWATPCLRNGQLDQIVDQRIAGTIRPGSLKKLADTADKCLAEYGVERPTMGDVLWCLEFALQLQVASSDVSDADTMLTPPVPVKTPEVQRSLSAATVATDAAMTTNLGDLDGMSLSGVFSKMIKSDEVR

>Ta-CrRLK1L8-D

MPPVLDMLVRLLVASVLLGAASGAFTPADNYLVLCGTSASATVAAGRTFVGDARLPAKSLAAPQSVEANTSRTAVVPSGESELYRSARVFTAPASYTFAVKQPGRHFVRLHFFPFPYRSYDMVADAAFNVSVQGAVLVNGYTPKNGTAELREFSVNVTGGTLVIAFAPTGKLAFVNAIEVVSVPDELIADMARMVDGAVQYTGLSTQALETIHRINMGVPKITPGNDTLGRTWLPDQSFQVNTDLAQHKDAKPLTIKYDEKSALSSAYTAPAEVYATATRLSTAGETSTINVQFNISWRFDAPAGSDYLLRFHWCDIVSKAAMGMAFNVYVGGAVVLDNYEISRDTFNRLSIPVYKDFLLGAKDAKGAITVSIGSSTEDNALPDGFLNGLEIMSIVGSAGAGAAATSPRSSKVKIGIIAGSAVCGATLVMVLGFIAFKMLRGREPEKKKPADAWSPFSASALGSRSRSRSFSKSNGNTVLLGQNGAGAGYRIPFAALQEATGGFDEGMVIGEGGFGKVYKGTMRDETVVAVKRGNRRTQQGLHEFHTEIEMLSRLRHRHLVSLIGYCDERGEMILVYEYMAMGTLRSHLYGAGLPPLSWEQRLEACIGAARGLHYLHTGSAKAIIHRDVKSANILLDESFMAKVADFGLSKNGPELDKTHVSTKVKGSFGYLDPEYFRRQMLTEKSDVYSFGVVLLEALCARTVIDPTLPREMVSLAEWATPCLRNGQLDQIVDQRIAGTIRPGSLKKLADTAEKCLAEYGVERPTMGDVLWCLEFALQLQVGSSDSSDVDTMLPPAPPVPVKTPEVQRRLSAATVATDAAAMTTNLGDLDGMSLSGVFSNMIKSDEVR

>Ta-CrRLK1L9-A

MGGGPRCALLLLVAAAAAALVPAAWAQDPTAPAPSGAPFVPRDDILLDCGATGKGNDTDGREWRGDAGSKYAPPNLASADAGAQDPSVPQVPYLTARVSAAPFTYSFPLGPGRKFLRLHFYPANYSGRAAADAFFSVSVPAAKVTLLSNFSAYQTATAFNFAYLVREFSVNVTGPTLDLTFTPEKGRPNAYAFINGIEVVSSPDLFDLATPFFVTGDGNNQPFPMDPGAALQTMYRLNVGGQAISPSKDSGGARSWDDDTPYIYGAGAGVSYPNDPNVTITYPPSVPGYVAPLDVYATARSMGIDKGVNLAYNLTWIVQVDAGFTYLVRLHFCEIQSPIDKPNQRVFNIYLNNQTAVEGADVLQWVDPRSTGTPLYKDFVVGTVGSGIMDFWVALHPDIRNKPQYYDAILNGMEVFKLQLTNGSLAGPNPVPSADPAAHTGQGKKSSLVGPIAGGVIGGLAVLALGYCCFICKRRRKVAKDAGMSDGHSGWLPLSLYGNSHTSSSAKSHATGSIASSLPSNLCRHFSFAEIKAATKNFDESRILGVGGFGKVYQGEIDGGTTKVAIKRGNPLSEQGIHEFQTEIEMLSKLRHRHLVSLIGYCEDKNEMILVYDHMAHGTLREHLYKTQNAPLSWRQRLEICIGAARGLHYLHTGAKHTIIHRDVKTTNILLDEKWVAKVSDFGLSKTGPSMDHTHVSTVVKGSFGYLDPEYFRRQQLTEKSDVYSFGVVLFEVLCARPALNPTLAKEEVSLAEWALHCQKKGILDQIVDPYLKGKIVPQCFKKFAETAEKCVADNGIERPSMGDVLWNLEFALQMQESAEESGSFGCGMSDEEGAPLVMAGKKDPNDPSIDSSTTTTTTTSLSMGDQSVASIDSDGLTPSAVFSQIMNPKGR

>Ta-CrRLK1L9-B

MGGGPRCALLLLAAAAACAALVPAAWAQGSTAPAPSGAPFVPRDDILLDCGATGKGNDTDGREWRGDASSKYAPPNLASADAGAQDPSVPQVPYLTARVSAAAFTYSFPLGPGRKFLRLHFYPANYSNRDAADAFFSVSVPAAKVTLLSNFSAYQTATALNFAYLVREFSVNVTGPTLDLTFTPEKGRPNAYAFINGIEVVSSPDLFDLATPFFVTGDGNNQPFPMDPGAALQTMYRLNVGGQAISPSKDSGGARSWDDDTPYIYGAGAGVTYPNDPNVTITYPDSVPGYMAPSDVYATARSMGIDKNVNLAYNLTWIVQVDAGFTYLVRLHFCEIQSPIDKPNQRVFNIYLNNQTAVEGADVIQWVDPLSTGTPLYKDYVVGTVGSGIMDFWVALHPDIRNKPQYYDAILNGMEVFKLQLSNGSLAGPNPVPSADPQAHTGQGKKKSLVGPIAGGVIGGLAVLALGYCCFICKRRRKAAKDTGMSDGHSGWLPLSLYGNSHTSSSAKSHATGSIASSLPSNLCRHFSFAEIKAATKNFDESRILGVGGFGKVYQGEIDGGTTKVAIKRGNPLSEQGIHEFQTEIEMLSKLRHRHLVSLIGYCEDKNEMILVYDHMAHGTLREHLYKTQNAPLSWRQRLEICIGAARGLHYLHTGAKHTIIHRDVKTTNILLDEKWVAKVSDFGLSKTGPSMDHTHVSTVVKGSFGYLDPEYFRRQQLTEKSDVYSFGVVLFEVLCARPALNPTLAKEEVSLAEWALHCQKKGILDQIVDPYLKGKIVPQCFKKFAETAEKCVADNGIERPSMGDVLWNLEFALQMQESAEESGSFGCGMSDEGTPLVMAGKKDPNDPSIDSSTTTTTTTSLSMGDQSVASIDSDGLTPSAVFSQIMNPKGR

>Ta-CrRLK1L9-D

MGGGPRCALLLLAAAAACAALVPAAWAQAPTAPAPSGAPFVPRDDILLDCGATGKGNDTDGREWRGDAGSKYAPPNLASADAGAQDPSVPQVPYLTARVSAAAFTYSFPLGPGRKFLRLHFYPANYSNRDAADAFFSVSVPAAKVTLLSNFSAYQTATALNFAYLVREFSVNVTGPTLDLTFTPEKGRPNAYAFINGIEVVSSPDLFDLATPLFVTGDGNNQPFPMDPGAALQTMYRLNVGGQAISPSKDSGGARSWDDDTPYIYGAGAGVTYPNDPNVTITYPDNVPGYVAPSDVYATARSMGIDKNVNLAYNLTWIVQVDAGFTYLVRLHFCEIQSPITKPNQRVFNIYLNNQTAVEGADVIQWVDPLSTGTPLYKDYVVSTVGSGIMDFWVALHPNTGSKPQYYDAILNGMEVFKLQLSNGSLAGPNPVPSADPPAHTGQEKKNSLVGPIAGGVIGGLVVLALGYCCFICKRRRKVAKDAGMSDGHSGWLPLSLYGNSHTSSSAKSHATGSIASSLPSNLCRHFSFAEIKAATKNFDESRILGVGGFGKVYQGEIDGGTTKVAIKRGNPLSEQGIHEFQTEIEMLSKLRHRHLVSLIGYCEDKNEMILVYDHMAHGTLREHLYKTQNAPLSWRQRLEICIGAARGLHYLHTGAKHTIIHRDVKTTNILLDEKWVAKVSDFGLSKTGPSMDHTHVSTVVKGSFGYLDPEYFRRQQLTEKSDVYSFGVVLFEVLCARPALNPTLAKEEVSLAEWALHCQKKGILDQIVDPYLKGKIVPQCFKKFAETAEKCVADNGIERPSMGDVLWNLEFALQMQESAEESGSFGCGISDEEGTPLVMAGKKDPNDPSIDSSTTTTTTTSLSMGDQSVASIDSDGLTPSAVFSQIMNPKGR

>Ta-CrRLK1L10-A

MLRTRILVLAAVSIVFANLQFLKAHGRELFLSCGSNATADADGRRWIGDMAPDLNFTLSSPGIAALLAGSSNGSEIMAPVYRSARFFTTTSWYDFSLLPGNYCVRLHFFPSAFRNFSANGSVFDVVANDFKLVSKFNVSEEIVWRNSVSNSAATAVVKEYFLAVNGSRLQIEFDPRPGSFAFVNAIEVMLTPDNSFNGMVNKVGGVDVHIPPELSGRAVETMYRLNIGGPALASSHDQYLHRPWYTDEAFMFSANAALIVSNTSAIKYVSSNDSSIAPLDVYETARIMGNNMVMDKRFNVTWRFFVHPNFDYLVRLHFCELVYDKPSQRIFKIYINNKTAAENYDVYNKAGGINKAYHEDYFDSLPQQVDSLWLQLGPDSMTSASGTDALLNGLEIFKLSRSGNLDYVLGHIDMGNKRGRSKGRSRIGLWEEVGIGSAAFVALVSVALFSWCYVRRKRKAVNEEVPAGWHPLVLHEAMKSTTDARASKKAPLARNSSSIGHRMGRRFSIADIRAATKNFDESLVIGSGGFGKVYKGEVDDGITVAIKRANPLCGQGLKEFETEIEMLSKLRHRHLVAMIGYCEEQKEMILVYEYMAKGTLRSHLYGSGLPPLTWKQRIDACIGAARGLHYLHTGADRGIIHRDVKTTNILLDKNFVAKIADFGLSKTGPTLDQTHVSTAIRGSFGYLDPEYFRRQQLTQKSDVYSFGVVLFEVACARPVIDPSVPKDQINLAEWAMRWQRQRSLEAIADPRLDGDYSPESLKKFGDIAEKCLADDGRTRPSMGEVLWHLEYVLQLHEAYKRNVDCESFGSSELGFADMSFSMPHIREGEEEHHPKKSGIREDSAP

>Ta-CrRLK1L10-B

MLRMRILVLASVSIVFANLQFLKAHGRELFLSCGSNATADADGRRWIGDMAPDLNFTLSSPGIAALLAGSTNGSEIMAPVYRSARFFTTTSWYDISVLPGNYCVRLHFFPSAFGNFSANGSVFDVVANEFKLVSKFNVSEEIVWRNSVSNSAATAVVKEYFLAVNSSRLQIEFDPRPGSFAFVNAIEVMLTPDNSFNSTVNKVGGVDVHIPPELSGRAIETMYRLNIGGPALASSHDQYLYRPWYTDEAFMFSANAALTVSNTSAIKYVSSGDSSIAPIGVYETARIMGNNMVMDKRFNVTWRFVVHPNFDYMVRLHFCELVYDKPSQRIFKIYINNKTAAENYDVYDKAGGINKAYHEDYFDSLPQQVDSLWLQLGPDSMTSASGTDALLNGLEIFKISRSGNLDYVLGHIDMGNKRGRSKGRSRLGLWEEVGIGSAAFVALASVALFSWCYVRRKRKAVDEEVPAGWHPLVLHEAMKSTTDARASKKAPLARNSSSIGHRMGRRFSIVDIRAATKNFDESLVIGSGGFGKVYKGEVDDGITVAIKRANPLCGQGLKEFETEIEMLSKLRHRHLVAMIGYCEEQKEMILVYEYMAKGTLRSHLYGSGLPPLTWKQRIDACIGAARGLHYLHTGADRGIIHRDVKTTNILLDKNFVAKIADFGLSKTGPTLDQTHVSTAIRGSFGYLDPEYFRRQQLTQKSDVYSFGVVLFEVACARPVIDPSVPKDQINLAEWAMRWQRQRSLEAIADPRLDGDYSPESLKKFGDIAEKCLADDGRTRPSMGEVLWHLEYVLQLHEAYKRNVDCESFGSSELGFADMSFSMPHIREGEEEHHPKKSGIREDSAP

>Ta-CrRLK1L10-D

MLRMRILVLAAVSIVFANLQFLKAHGRELFLSCGSNATADADGRRWIGDMAPGLNFTLSSPGIAALLAGSSNGSEIMAPVYRSARFFTTTSWYDFSLLPGNYCVRLHFFPSTFRNFSANGSVFDVVANDFKLVSKFNVSEEIVWRNSVSNSAATAVVKEYFLAVNSSRLQIEFDPRPGSFAFVNAIEVMLTPDNSFNGTVNKVGGVDAHIPPELSGRAVETMYRLNIGGPALASSHDQYLHRPWYTDEAFMFSANVALIVSNTSAIKYVSSNDSSIAPIDVYETARIMGNNMVMDKRFNVTWRFLVHPNFDYLVRLHFCELVYDKPSQRIFKIYINNKTAAENYDVYNRAGGINKAYHEDYFDSLPQQVDSLWLQLGPDSMTSASGTDALLNGLEIFKLSRSGNLDYVLGHIDMGNKRGRSKGRSRIGLWEEVGIGSAAFVALASVALFSWCYVRRKRKAVNEEVPAGWHPLVLHEAMKSTTDARASKKAPLARNSSSIGHRMGRRFSIADIRAATKNFDESLVIGSGGFGKVYKGEVDDGITVAIKRANPLCGQGLKEFETEIEMLSKLRHRHLVAMIGYCEEQKEMILVYEYMAKGTLRSHLYGSGLPPLTWKQRIDACIGAARGLHYLHTGADRGIIHRDVKTTNILLDKNFVAKIADFGLSKTGPTLDQTHVSTAIRGSFGYLDPEYFRRQQLTQKSDVYSFGVVLFEVACARPVIDPSVPKDQINLAEWAMRWQRQRSLEAIADPRLDGDYSPESLKKFGDIAEKCLADDGRTRPSMGEVLWHLEYVLQLHEAYKRNVDCESFGSSELGFADMSFSMPHIREGEEEHHPKKSGIREDSAP

>Ta-CrRLK1L11-A

MGTTTEQKIALLLLGTIWVLLGTCNAAEFTPADNYLINCGSTVDANLNDGRVFKADNSGSTILTSHHSVPANTLPDAVISSDNPVLYKTARIFIVPSSYSFNMKSRGRHFVRLHFFGFRYQSYDLAAAKFKVSTQHVVLLDNFTPPSNSSLLVREYSLNITEDMLILSFVPLGNSTSFINAIEVISVPDDLIQDSAQTVNPSGQYLGLATQSFQTFYRINVGGREVTVVNDTLSRSWDTDQNFFINSTTTELFAYQGKLNYQKGAATKEDAPDSVYNTARRLAVQNRTSPASNMTWQFDVDGRSSYLIRFHFCDIVSKAAYSLYFDIYVDGGLALENLDLSEKVFGTLAVPYYTEFVLKSSNPSGKLSVGIGPSSLSNVAPDGILNGLEIMKMNISTGTIYVVWPPATPKRKLAIILAPVLGGVGAVSIAIILCFVLRRKKEKKPRRAPTSRPSSSWSPLTLNGLSFLSIGTRTTSRTTHTSGTNSDVSYRIPFALLQVATKHFDEQMVVGVGGFGKVYKAVLQDSTKVAVKRGNQKSHQGLKEFRTEIELLSGLRHRHLVSLIGYCDDQNEMILVYEYMEKGTLKSHLYGSDMPPLSWKKRVEICIGAARGLHYLHTGFAKSIIHRDVKSANILLDENLMAKVSDFGLSKTGPELDQTHVSTAVKGSFGYLDPEYYRRQKLTDKSDVYSFGVVLLEVICARPVIDPTLPRDMINLAEWAIKWQKRGELGQIVDQRIAGTIRPESLRKYGETVEKCLVDYGVDRPTMGDVLWNLEFVLQLQEAGPDISNVDSMNQISELPSDARRMGSLEIGTADEADEGRTHMDYSQMSTNDAFSQLMNTEGR

>Ta-CrRLK1L11-B

MGTTTEQKIALLLLGTIWVLLGTCNAAEFTPADNYLINCGSTVDANLHDGRVFKADNSGLTILTSHHSVPANTLPDAVISSDNPVLYQTSRIFIVPSSYSFKMKSRGRHFVRLHFFSFRYQSYDLAAAKFKVSTQHVVLLDNFTPPSNSSPLVREYSLNITEDMLILSFVPQGNSTSFISAIEVISVPDDLIQDSAQTVNPSGQYLGLATQSFQTFYRINVGGREVTVVNDTLSRSWDTDQNFFLNSTTTELFAYQGKLNYQKGAATKEDAPDSVYNTARRLAVQNRTSPASNMTWQFDVDGRSSYLIRFHFCDIVSKAAYSLYFDIYVDGGLALENLDLSEKVFGTLAVPYYTEFVLKSSNPSGKLSVGVGPSSLNNVAPDGILNGLEIMKMNISTGTIYVVWPPAPPKRKLAIILGSVLGGVGAVSIAIILCFVLRRKKKEKKPRRAPTSRPSSSWSPLTLNGLSFLTVGTRTTSRTTHTSGTNSDVSYRIPFALLQVATKHFDEQMVVGVGGFGKVYKAVLQDSTKVAVKRGNQKSHQGLKEFRTEIELLSGLRHRHLVSLIGYCDEQNEMILVYEYMEKGTLKSHLYGSDMPPLSWKKRVEICIGAARGLHYLHTGFAKSIIHRDVKSANILLDENLMAKVSDFGLSKTGPELDQTHVSTAVKGSFGYLDPEYYRRQKLTDKSDVYSFGVVLLEVICARPVIDPTLPRDMINLAEWAIKWQKRGELGQIVDQRIAGTIRPESLRKYGETVEKCLANYGVDRPTMGDVLWNLEFVLQLQEAGPDISNVDSMNQISELPSDARRMGSLEIRTADEADESRTNMDYSQMSTNDAFSQLINTEGR

>Ta-CrRLK1L11-D

MGTTTGQKTALLLLGTLWVLLGTCNAAEFSPADNYLINCGSTVDANLHDGRVFKADNSGSTILTSHHSVPANTLPDAVISSDNPVLYQTARIFIVPSSYSFNMKSRGRHFVRLHFFGFRYQSYDLAAAKFKVSTQHVVLLDNFTPPSNSSPLDSAQTVNPSGQYLGLATQSFQTFYRINVGGREVTVVNDTLSRSWDTDQNFFINSTTTELFAYQGRLNYQKGAATKEDAPDSVYNTARRLAVQNRTSPASNMTWQFDVDGRSSYLIRFHFCDIVSKAAYSLYFDIYVDGGLALENLDLSEKVFGTLAVPYYTEFVLKSSNPSGKLSVGIGPSSLNNVALDGILNGLEIMKMNISTGTIYVVWPPATPKRKLAIILGPVLGGVGAVSIAIILCFVLRRKKKEKKPRRAPTSRPSSSWSPLTLNGLSFLSIGTRTTSRTTHTSGTNSDVSYRIPFALLQVATKHFDEQMVVGVGGFGKVYKAVLQDSTKVAVKRGNQKSHQGLKEFRTEIELLSGLRHRHLVSLIGYCDEQNEMILVYEYMEKGTLKSHLYGSDMPPLSWKKRVEICIGAARGLHYLHTGFAKSIIHRDVKSANILLDENLMAKVSDFGLSKTGPELDQTHVSTAVKGSFGYLDPEYYRRQKLTDKSDVYSFGVVLLEVICARPVIDPTLPRDMINLAEWAIKWQKRGELGQIVDQRIAGTIRPESLRKYGETVEKCLADYGVDRPTMGDVLWNLEFVLQLQEAGPDISNVDSMNQISELPSDARRMGSLEIGTADEGRTNMDYSQMSTNDAFSQLMNTEGR

>Ta-CrRLK1L12-A

MAAARGRGRGRGRCVLLAAVLLLTAVVGADIYKPTDSILVHCGSDKDGQDEDGRKWTTDKDSKWLPDGGKSSIMGTADVADPSLPSPVPYMTARVFPKETAYTFPVADADRHWVRLHFYPAAYHGIPADHFFFSVTTSTGVTLLRNFSVYITAKALTQAYIIREFSLPPSTVGSLSLKFTPTAMNNASYAFVNGIEVISMPSFFGDPATLVGLNDQSLDASAANLQTMYRLSVGGSYIPPANDSGLSREWFSDTPYVYGAATGVTFEANDTVPIKYPAPADEYAAPVSIYDSFRHMGRDPKMNKNNNLTWVFEVDGNFTYLLRLHFCSLMEDKINQVVFAILLNNKTATTTGSADIIAWAKEKNPANPGAPGKGVPVFKDYAVFMPAAPAGNDTILWLTLRPDTATRTQFVNAFLNGLEVFKVSDASGNLAGPNPDISKMLAEAELGAVEGQFREKPSNVGALIGGAAGGAAAFGLVAAVCFVAYQSKRRRELSSSPSHSSSGWLPVYGGSTSVSKSSGGRSAATLNPNITAMCRHFSLQEIKSATKGFDESLVIGVGGFGKVYRGVVDGDTKVAVKRSNPSSEQGVLEFQTEIEMLSKLRHKHLVSLIGCCEDNGEMILVYDYMAHGTLREHLYNKSGKPPLPWRQRLEIVIGAARGLHYLHTGAKYTIIHRDVKTTNILVDDKWVAKVSDFGLSKTGPTVQNQTHVSTMVKGSFGYLDPEYFRRQKLTEKSDVYSFGVVLFEVLCGRPALNPSLPREQVSLADHALSCQRKGTLEEIVDPVLEGKIAPDCLKKFAETAEKCLADQGVDRPSMGDVLWNLEFALQMQDTFDNGGKPPEVDDYSSSFTIAQPSMEESLAANAAALSLISEDMDEEDIANSVIFSQIAKPTGR

>Ta-CrRLK1L12-B

MAAARGRGVLLAVLLLTTVAFAFVGADIYKPTDSILVNCGSDKDGQDEDGRKWTTDKDSKWLPDGGKSSIMGTADVSDPSLPSPVPYMTARVFPKETAYTFPVSDADRHWVRLHFYPAAYHDIPADHFFFSISTSTGITLLRNFSVYITAKALTQAYIVREFSLPPSTAGSLSLKFTPTAMNNASYAFVNGIEIISMPNFFGDPATLVGLDDQSLDASAGNLQTMYRLSVGGSYIPPTNDSGLTREWFSDTPYVYGAGTGVTFEANDTIPIKYPAPADEYAAPVSIYDTFRHMGRDANLNKNNNLTWVFEVDGNFTYLLRLHFCSLMEDKINQVVFAILVNNKTATTTGSADIIAWAKEKNPANPGAPGKGVPVFKDYAVFMPAAPAGNDTILWLTLRPDTASNPQFVNAFLNGLEIFKVSDASGNLAGPNPDISKMLAEAELGAVDGQFREKPSNVGALIGGAVGGAAAFGLVAAVCFVAYQSKRGRELSSSPSHSSSRWLPVYGSSQTSVSKSSGGRSAMTLNPNITAMCRHFSFQEIKSATKGFDESLVIGVGGFGKVYRGVVDGDTKVAIKRSNPSSEQGVLEFQTEIEMLSKLRHKHLVSLIGCCEDNGEMILVYDYMAHGTLREHLYKSGKPPLPWRQRLEIVIGAARGLHYLHTGAKYTIIHRDVKTTNILVDEKWVAKVSDFGLSKTGPTVQNQTHVSTMVKGSFGYLDPEYFRRQKLTEKSDVYSFGVVLFEVLCGRPALNPSLPREQVSLADHALSCQRKGTLEEIIDPVLEGKIAPDCLKKFAETAEKCLADQGVDRPSMGDVLWNLEFALQQQDTFENGGKPPEVDDYSSSFTITPPSMEESLAANAAALSLISEDMDEEDIANSVIFSQIAKPTGR

>Ta-CrRLK1L12-D

MAAARGRGVLLAVLLLMMVAFAFVGADIYKPTDSILVHCGSDKDGQDEDGRKWTADKDSKWLPDGGKSSVMGTADVPDPSLPSPVPYMTARVFPKETAYTFPVADADRHWVRLHFYPAAYHGIPADHFFFSVTTSTGVTLLRNFSVYTTAKALTQAYIVREFSLPPSTTGSLSLKFTPTAMNNASYAFVNGIEIISMPSFFGDPATLVGLDDQSLDASAGNLQTMYRLSVGGSYIPPANDSGLSREWFSDTPYVYGAATGVTFEANDTIPIKYPTPADEYAAPVSIYDSFRHMGRDPKMNRNNNLTWVFEVDGNFTYLLRLHFCSLMEDKINQVVFAILVNNKTATTTGSADIIAWAKEKNPANPGAPGKGVPVFKDYAVFMPAAPAGSDTILWLTLRPDTATNPQFVNAFLNGLEVFKVSDASGNLAGPNPDISKMLAEAELGAVDGQFREKPSNVGALIGGAAGGAAAFGLVAAVCFVAYQSKRRRELSSSPSHSSSGWLPVYGGNSQTSVSKSSGGRSAVTLNPNITAMCRHFSFQEIKSATKGFDESLVIGVGGFGKVYRGVVDGDTKVAIKRSNPSSEQGVLEFQTEIEMLSKLRHKHLVSLIGCCEDNGEMILVYDYMAHGTLREHLYKSGKPPLPWRQRLEIVIGAARGLHYLHTGAKYTIIHRDVKTTNILVDEKWVAKVSDFGLSKTGPTVQNQTHVSTMVKGSFGYLDPEYFRRQKLTEKSDVYSFGVVLFEVLCGRPALNPSLPREQVSLADHALSCQRKGTLEEIIDPVLEGKIAPDCLKKFAETAEKCLADQGVDRPSMGDVLWNLEFALQQQDTFENGGKPPEVDDYSSSFTITPPSMEESLAANAAALSLISEDMDEEDIANSVIFSQIAKPTGR

>Ta-CrRLK1L13-A

MVRRGALPLALLAVLATLTAVAGQGKPVTDNGSGGGSGPSKFTPKDAFYIDCGGTAAADTKDGKSFKTDAEANSLLSARDNIKVADDKADVPSHLYRSARVFKEEAVYNFPLTAPGWHFIRLYFFPIKSGEADLAAATFDVSTAVNVLLHGFTPEAKAVMKEYIVNATENKLELKFTPQSGSAFINAIEVVNAPDELISKTALTVSPLAETSGLSEAAYQVVCRLNVGGPPIGPVNDTLGRQWEDDGQYLNPKDAGTEVSVPTSAIKYPDAFPATKLVAPTAVYATARHMAESGVANQNFNVSWKVDVDPSFDYLVRLFFADIISTSANDLYFNAYINGRKAISALDLSTITGDLAAPYYKDFVVNSSVNTDGHIIIGVGPLGQDTGRNDALLNGAEVLKMSNSVGSLDGEFGVDGRMVDDGSGTRKVVAAVGFAMMFGAFAGLGCMVVKWHRRPQDWDRRNSFSSWLLPIHTGQSFSNGKGSKSGYTFSSTAGLGHFFTFAEMSEATKNFDESAIIGVGGFGNVYVGEINDPDEEGSRIKVAIKRGNPSSEQGINEFNTEIQMLSKLRHRHLVSLIGYCDEGEEMILVYEFMQHGPFRDHIYGGPEGLPTLSWKQRLEICIGAARGLHYLHTGTAHGIIHRDVKTTNILLDEKFVAKVADFGLSKDGPGMNQLHVSTAVKGSFGYLDPEYFRCQQLTDKSDVYSFGVVLLETLCARAPIDPQLPREQVSLAEWGLQWKRKGLIEKIMDPNLNGKVNPESLAKFAETAEKCLCEFGSDRLSMGDVLWNLEYALQLQEANPPEGATDADDADASIVSSASGVTTVPDQSTTSANELFAQLADMKGR

>Ta-CrRLK1L13-B

MVRRGTFPLALLAVLATLTAVAGQGKPVTDNGSGGASGPAKFTPKDAFYIDCGGTAAADTKDGKSFKTDAEANSLLSARDNIKVADDKADVPSHLYRSARVFKEEAVYNFPLTAPGWHFIRLYFFPIKSGEADLAAATFDVTTAVNVLLHGFTAEAKAVMKEYVVNATENKLELKFTPQSGAAFINAIEVVNAPDELISKTALTVSPLAETSGLSEAAYQVVCRLNVGGPPIGPVNDTLGRQWEDDGQYLNPKEAGAEVSVPTSAIKYPDAFPATKLVAPTAVYATARHMAESGVANQNFNVSWKVDVDPSFDYLVRLFFADIISTSANDLYFNVYINGRKAISALDLSTITGDLAAPYYKDFVVNSSVNTDGHIIIDVGPLGQDTGRNDALLNGAEVLKMTNSVGSLDGEYGVDGRMVDDGSGTRKVVAAVGFAMMFGAFAGLGCMVVKWHRRPQDWERRNSFSSWLLPIHTGQSFSNGKSKSGYTFSSTAGLGHFFTFAEMSEATKNFAESAIIGVGGFGNVYVGEINDPDEEGSRIKVAIKRGNPSSEQGINEFNTEIQMLSKLRHRHLVSLIGYCDEGEEMILVYEFMQHGPFRDHIYGGPEGLPTLSWKQRLEICIGAARGLHYLHTGTAHGIIHRDVKTTNILLDDKFVAKVADFGLSKDGPGMNQLHVSTAVKGSFGYLDPEYFRCQQLTDKSDVYSFGVVLLETLCARAPIDPQLPREQVSLAEWGLQWKRKGLIEKIMDPNLAGKVNPESLAKFAETAEKCLCEFGSDRLSMGDVLWNLEYALQLQESNPPEGASDADDADASIVSSASGVTTVPDQSTTSANELFAQLADMKGR

>Ta-CrRLK1L13-D

MVRRGALPLALLAVLATLTAVAGQGKPVTDNGSGGGAGPAKFTPKDAFYIDCGGTAAADTKDGKSFKTDAEANSLLSARDNIKVADDKADVPSHLYRSARVFKEEAVYNFPLTAPGWHFIRLYFFPIKSGEADLAAATFDVSTAVNVLLHGFTPEAKAVMKEYIVNATENKLELKFTPQSGSAFINAIEVVNAPDELISKTALTVSPLAETSGLSEAAYQVVCRLNVGGPPIGPVNDTLGRQWEDDEKYLNPKEAGTEVSVPTSAIKYPDAFPATKLVAPTAVYATARHMAESGVANQNFNVSWKVDVDPSFDYLVRLLFADIISTSANDLYFNVYINGRKAISALDLSTITGDLAAPYYKDFVVNSSVNTDGHIIIDVGPLGQDTGRNDALLNGAEVLKMSNSVGSLDGEYGVDGRMVDDGSGTRKVVAAVGFAMMFGAFAGLGCMVVKWHRRPQDWERRNSFSSWLLPIHTGQSFSNGKSKSGYTFSSTAGLGHFFTFAEMSEATKNFDESAIIGVGGFGNVYVGEINDPDEEGSRIKVAIKRGNPSSEQGINEFNTEIQMLSKLRHRHLVSLIGYCDEGEEMILVYEFMQHGPFRDHIYGGPEGLPTLSWKQRLEICIGAARGLHYLHTGTAHGIIHRDVKTTNILLDDKFVAKVADFGLSKDGPGMNQLHVSTAVKGSFGYLDPEYFRCQQLTDKSDVYSFGVVLLETLCARAPIDPQLPREQVSLAEWGLQWKRKGLIEKIMDPNLNGKVNPESLAKFAETAEKCLCEFGSDRLSMGDVLWNLEYALQLQEANPPEGATDADDADASIVSSASGVTTVPDQSTTSANELFAQLADMKGR

>Ta-CrRLK1L14-A

MPAAGRSGGPGQVNIMMGRRKLQVVTLAILCFWSSAGICKAQSVDFKPADSYLVDCGSAKGTTVLGRDFAADGAAPVTVATSQDILAGTSANGVSSFDNPVLYQTARIFTSPSSYTFPIQKQGRHFVRLYFYPFIYQSYDLSTAKFTVSTQDVLLLSDFQQPDKTAPLFKEYSLNITRDQLVISFKPSNGIAFINAIEVISVPDDLIADVANMVNPVQQYSGLTTQSLETVYRVNMGGPKVFPNNDTLSRTWQKDQKYILNPSVTKTAQYGKAINYRKGGATPLTAPDIVYSTATELAASNTSNALFNMTWQFDVDAGFSYLIRFHFCDIVSKALNQLYFNAYVGGFFAQHDLDLSEQSVNQLATAIYVDVVLSSNDASSKLSISIGPSTLNNALPDGILNGLEIMKMGSGSGSAFTVGNNGSNKKLPIIIGSVLGVVGLLIIVLVVVLLCRRKKTDDKQHSKTWMPFSINGLTSLSTGSRTSYGTTLTSGLNGSYGYRFAFNVLQEATNNFDESWVIGVGGFGKVYKGALRDDTKVAVKRGNPKSQQGLNEFRTEIELLSRLRHRHLVSLIGYCDERNEMILVYEYMENGTVKSHLYGSDNPSLNWKQRLEICIGAARGLHYLHTGSAKAIIHRDVKSANILLDENLLAKVADFGLSKTGPELDQTHVSTAVKGSFGYLDPEYFRRQQLTEKSDVYSFGVVMLEVLCARPVIDPSLPREMVNLAEWGMKWQKRGELHQIVDQKLSGAIRPDSLRKFGETVEKCLADYGVERPSMGDVLWNLEYVLQLQDVDSSTVSDVNSMNRIVDLSSQVQHVSAMESISVTMAEDGALHEPDHDLSDVSMSRVFSQLIKAEGR

>Ta-CrRLK1L14-B

MPAAARSGGPGQANIMMGRRKLQAVTLAILCFWSSAGAQTVDFKPADNYLVDCGSAKGTTVLGRDFAADGASPVTVSTSQDILAGTSANGVSSFDNPLLYQTARIFTSPSSYTFPIQKQGRHFVRLYFFPFIYQSYDLSTAKFTVSTQDVLLLSDFQQPDKTAPLFKEYSLNITRDQLVISFKPSNGIAFINAIEVVSVPDDLIADVANMVNPVQQYSGLTTQSLETVYRVNMGGPKVFPSNDTLSRTWQKDQKYILNPSVTKTAQYGKPIKYRKGGATPLTAPDIVYSTATELAAANTSNALFNMTWQFDVDAGFSYLIRFHFCDIVSKALNQLYFNAYVGGFFAQHDLDLSEQSVNQLATAIYVDVVLSSNDASSKLSISIGPSTLNNALPDGILNGLEIMKMGSGSGSAFTVGNNGSNKRLPIIIGSVLGVVGLLIIVLVVVLLCRRKKTDDKQHSKTWMPFSINGLTSLSTGSRTSYGTTLTSGLNGSYGYRFAFNVLQEATNNFDESWVIGVGGFGKVYKGALRDDTKVAVKRGNPKSQQGLNEFRTEIELLSRLRHRHLVSLIGYCDERNEMILVYEYMENGTVKSHLYGSDNPSLNWKQRLEICIGAARGLHYLHTGSAKAIIHRDVKSANILLDENLLAKVADFGLSKTGPELDQTHVSTAVKGSFGYLDPEYFRRQQLTEKSDVYSFGVVMLEVLCARPVIDPSLPREMVNLAEWGMKWQKRGELHQIVDQKLSGAIRPDSLRKFGETVEKCLADYGVERPSMGDVLWNLEYVLQLQDVDSSTVSDVNSMNRIVDLSSQVQHVGAMESISVTMAEDGALHEPDHDLSDVSMSRVFSQLIKAEGR

>Ta-CrRLK1L14-D

MMGRRKLQVVTLAILCFWSSAGVCKAQTVDFKPADSYLVDCGSTKGTTVLGRDFAADGASPVTVSTSQDILAGTSANGVSSFDNPVLYQTARVFTSPSSYTFPIQKQGRHFVRLYFYPFIYQSYDLSTAKFTVSTQDVLLLSDFQQPDKTAPLFKEYSLNITRDQLVISFKPSNGIAFINAIEVVSVPDDLIADVANMVNPVQQYSGLTTQSLETVYRVNMGGPKVFPNNDTLSRTWQKDQKYILNPSVTKTAVYGKAIKYRKGGATPLTAPDIVYSTATELAASNTSNALFNMTWQFDVDAGFSYLIRFHFCDIVSKALNQLYFNAYVGGFFAQHDLDLSEQSVNQLATAIYVDVVLSSNDASSKLSISIGPSTLNNALPDGILNGLEIMKMGSGSGSAFTVGNNGSNKKLPIIIGSVLGVVGLLIIVLVVVLLCRRKKTDDKQHSKTWMPFSINGLTSLSTGSRTSYGTTLTSGLNGSYGYRFAFNVLQEATNNFDESWVIGVGGFGKVYKGALRDDTKVAVKRGNPKSQQGLNEFRTEIELLSRLRHRHLVSLIGYCDERNEMILVYEYMENGTVKSHLYGSDNPSLNWKQRLEICIGAARGLHYLHTGSAKAIIHRDVKSANILLDENLLAKVADFGLSKTGPELDQTHVSTAVKGSFGYLDPEYFRRQQLTEKSDVYSFGVVMLEVLCARPVIDPSLPREMVNLAEWGMKWQKRGELHQIVDQKLSGAIRPDSLRKFGETVEKCLADYGVERPSMGDVLWNLEYVLQLQDVDSSTVSDVNSMNRIVDLSSQVQHVGAMESISVTMAEDGALHEPDHDLSDVSMSRVFSQLIKAEGR

>Ta-CrRLK1L15-A

MNSSANFLSILVLLVFLAAGNARAQPQPILINCASDSTTSVDARTWIGDSSPSNNFTLSFPGAIASAAPAPAPAPGVDGEQDPYGDLYKTARVFNASSSYRLAVAPGSYFLRLHFSQQFANLGAQEPIFSVAANGLRLLSKFSVHGEISWRDSQINSTSSVIVKEYLLNVTSGKLGIEFTPDEGSFAFINAMEVLPVSGTSIFDSVNKVDAHGLKGPFSLDGGGIETMYRLCVGCRDVLTRKEDPGLWRRWDKDDHFIFSLNAANSIFNSSNISYVSADDPTVAPLRLYQSARVPTESSVLGKKFNVSWSFNIDPGFDYLVRLHFCELQYDKAEQRKFKIYINNKTAAEGYDVLARAGGKNKAFYEDFLDAASPQMDTLWVQLGSESSAGSAAADALLNGMEIFKVSREGNLAHPTVRIGGFSGGTSKPKRSPKWVLIGAASGLIIFIAIAAALYLCFNLRRKKNSSASKAKDNPHGAAHTRSPTLLTAGAFGSKRMGRRFTIAEIRTATVNFDESLVIGVGGFGKVYRGIMEDGTRVAIKRGYTDSHQGQGVKEFETEIEMLSRLRHRHLVPLIGYCDEQNEMVLVYEHMANGTLRSHLYGSDLPALTWKQRLEICIGAARGLHYLHTGLDRGIIHRDVKTTNILLDNNLVAKMADFGISKDGPALDHTHVSTAVKGSFGYLDPEYYRRQQLTPSSDVYSFGVVLFEVLCARPVINPTLPRDQINLADWALNRQRHKLLETIIDLRLDGNYTLESIRTFSEIAEKCLADEGVNRPSMGEVLWHLESALQLEQGHLQSTNGDGCSDPQLKPSDVPTHVACIKEVEQSTRPGSHDSDGQVVDVKIEVP

>Ta-CrRLK1L15-B

MKSSANFLSILVLLVFLAAGNARAQPQPILINCGSDSTTSVDARTWIGDSSPSNNFTLSFPGAIASAAPAPGVDGEQDPYGDLYKTARVFNASSSYRLAVAPGSYFLRLHFSQQFANLGAQEPIFNVAANGLRLLSKFSVHGEISWRDSQINSTSSVIVKEYLLNVTSGKLGIEFTPDEGSFAFINAMEVLPVSGTSIFDSVNKVDGHGLKGPFSLDGSGIETMYRLCVGCIDVLARKEDPGLWRRWDKDEHFIFSLNAASSIFNSSNISYVSADDPTVAPLRLYQSARVPTESSVLGKKFNVSWSFNIDPGFDYLVRLHFCELQYDKAEQRKFKIYINNKTAAEGYDVFARAGGKNKAFYEDFLDAASPQMDTLWVQLGSESSAGSAAADALLNGMEIFKVSREGNLAHPTVRIGGISGGARKPKRSPKWVLIGAASGLIIFIAIAGALYFCFNLQRKKNSSANKAKDNLHGVTHTRSPTLRTAGAFGSKRMGRRFTIAEIRTATVNFDESLVIGVGGFGKVYRGIMEDGTRVAIKRGYTDSHQGQGVKEFETEIEMLSRLRHRHLVPLIGYCDEQNEMVLVYEHMANGTLRSHLYGSDLPALTWKQRLEICIGAARGLHYLHTGLDRGIIHRDVKTTNILLDDNLVAKMADFGISKDGPALDHTHVSTAVKGSFGYLDPEYYRRQQLTPSSDVYSFGVVLFEVLCARPVINPTLPRDQINLADWALNRQRHRLLETIIDLRLDGNYTLESIKIFSEIAEKCLADEGVNRPSMGEVLWHLESALQLEQGHPQSTNGDGCSDPQLKPSDVPTRVACIKEVEQSTRPGSHDSDGQVVDVKIEVP

>Ta-CrRLK1L15-D

MKSSANFLSILVLLVFLAAENARAQPQPILINCGSDSTTSVDARTWIGDSSPSNNFTLSFPGAIASAAPAPAPGVDGEQDPYGDLYKTARVFNASSSYRLAVAPGSYFLRLHFSQQFANLGAQEPIFSVAANGLRLLSKFSVHGEISWRDSQINSTSSVIVKEYLLNVTSGKLGIEFTPDEGSFAFINAMEVLPVSGTSIFDSVNKVDAHGLKGPFSLDGDGIETMYRLCVGCIDVLPRKEDPGLWRRWDKDEHFIFSLNAANSIFNSSNISYVSADDPTVAPLRLYQSARVPTESSVLGKKFNVSWSFNIDPGFDYLVRLHFCELQYDKAEQRKFKIYINNKTAAESYDVFARAGGKNKAFYEDFLDAASPQMDTLWVQLGAESSAGSAAADALLNGMEIFKVSREGNLAHPTVRIGGISGGASKPKRSPKWVLIGTASGLIIFIAIAGGLYFGFNLRRKKNSSASKAKDNLHGATHTRSPTLRTAGAFGSNRMGRRFTIAEIRTATVNFDESLVIGVGGFGKVYKGIMEDGTRVAIKRGHTDSHQGQGVKEFETEIEMLSRLRHRHLVPLIGYCDEQNEMVLVYEHMANGTLRSHLYGSDLPALTWKQRLEICIGAARGLHYLHTGLDRGIIHRDVKTTNILLDDNLVAKMADFGISKDGPALDHTHVSTAVKGSFGYLDPEYYRRQQLTPSSDVYSFGVVLFEVLCARPVINPTLPRDQINLADWALNRQRHRLLETIIDLRLDGNYTLASVKKSSKIAEKCLADEGVNRPSMGEVLWHLESALQLEQGHPQSTNADGCSDPQLKPSDVPTRVACIKEDEQSTRPGSHNSDGQVVDVKIEVP
