## Supplementary Tables for "Genome wide characterization and expression analysis of CrRLK1L gene family in wheat unravels their roles in development and stress-specific responses": Supplementary Table S3. CrRLK1L protein sequences in other species.docx

Supplementary Table S3. Protein sequences used in phylogenetic analysis are given below*. B. distachyon* sequences (Bd) were collected from the Ensembl Plants database, and sequences for *Ae. tauschii* (*Aet*) and *H. vulgare* (*Hv*) were collected from the NR database at NCBI, Arabidopsis sequences (*At*) were retrieved from the TAIR database, and rice sequences (*Os*) were collected from the Rice Genome Annotation Project database (http://rice.plantbiology.msu.edu/) (Nguyen et al., 2014).

>Aet-CrRLK1L1(XP_020194102.1)

MPALAILARSMAECKRVPMFLILFILSITSVATTNAIASKVDRFVPQDNYLLSCGASAAVQVDDGRTFRSDPESVSFLSTPTDIKIAAKASLASASPLSPLYLDARVFSDISTYSFFISQPGRHWIRLYFLPITDSQYNLTTATFSVSTDSMVLLHDFSFIASPPNPVFREYLVSAQGDNLKIIFTPKKNSIAFINAIEVVSAPPSLIPNTTTRMGPQDQFDISNSALQVVYRLNMGGALVTSFNDTLGRTWQPDAPFLKLEAAAEAAWVPPRTIKYPDDKTLTPLIAPASIYSTAQQMASTNITNARFNITWQMAAEPGFRYLIRLHFSDIVSKTLNSLYFNVYINGMMAVANLDLSSLTMGLAVAYYKDLIAESSSIINSTLVVQVGPNTIDSGDPNAILNGLEIMKISNEANSLDGLFSPKTSSEVSKTTLTGIAFALAATAALAVVICYRRNRKPAWQRTNSFHSWFLPLNSSSSFMSSCSRLSRNRFGSTRTKSGFSSVFASSAYGLGRYFTFVEIQKATKNFEEKGVIGVGGFGKVYLGATEDGTQLAIKRGNPSSDQGMNEFLTEIQMLSKLRHRHLVSLIGCCDENNEMILVYEFMSNGPLRDHLYGDTNIKPISWKQRLEVCIGAAKGLHYLHTGSAQGIIHRDVKTTNILLDENFVAKVADFGLSKDAPSLEQTHVSTAVKGSFGYLDPEYFRRQQLTDKSDVYSFGVVLFEVLCARPAINPALPRDQVNLGEWARTWHRKGELGKIIDPNIAGQIRPDSLEMFAEAAEKCLADYGVDRPTMGDVLWKLEFALQLQEKGDVVDGASDGIAMKSLEVTNVDSMEKSGNAIPSYVQGR

>Aet-CrRLK1L2(XP_020194829.1)

MAFPALPVTLTCLILLSLLSLAMAADNNSTGLILVNCGASVQGDDDSGRTWDGDTGSKFAPSLKGVAATAPNQDPSLPSTVPFMTARIFTSNYTYSFSVKPGRMFLRLYFYPVAYPNYAVSDAFFSVTTPKLVLLNDFSASQTAQAITSAFLVREFSVNVSSGSSLDLTFAPSAHRNGSYAFVNGIEIVPTPDIFTAPDTRNVGDNTAPFSFDTSSSLQTMYRLNVGGQAISPKGDLGGFYRSWANDAPYIAGGSGVTFSKDDNLTITYTSKVPKYTAPPDVYGTARSMGPTAQINLNYNLTWILPVDAGFFYLLRFHFCEIQYPIIKINQRSFFIYINNQTAQEQMDVIVWSGGIGRTTYTDYVIMAAGFGQVDMWIALHPDLSSRPEYFDAILNGLEVFKLQNYGSPNNLSGLNPPLPQKPADASPSAASGKMKSVAAIIGGAVGGFIVLLAACFGVCIICKRKNKKKKKTSKDPGGKSEDGHWTPLTEYSGSRSAMSGNTATTGSTLPSNLCRHFTFADLQTATKNFDQAFLLGKGGFGNVYLGEIDSGTKVAIKRCNPMSEQGVHEFQTEIEMLSKLRHRHLVSLIGYCEDKSEMILVYDYMAHGTLREHLYNTKNPPLSWKQRLEICIGAARGLYYLHTGVKHTIIHRDVKTTNILLDDKWVAKVSDFGLSKTGPNMDATHVSTVVKGSFGYLDPEYFRRQQLSEKSDVYSFGVVLFEVLCARPALSPTLPKEQISLADWALRCQKQGVLGQVIDPVLQGKIAPQCFLKFTDTAEKCVADRSVDRPSMGDVLWNLEFALQLQESEEDTGSLTEGTLSSSGASPLVMTRLQSDEPSTDASTTTTSTTTMSMTGRSIASMDSDGLTPSAVFSQIMHPDGR

>Aet-CrRLK1L3(XP_020197218.1)

MAATARLRPARARGVLWVVSVLLVCGAAAYKPEDNYLVSCGSSLDTPVGRRLFLADDGASGAVTLTSPRSAAVKAPPDLVSGFRDAALYQNARVFSAPSSYSFAIRHRGRHFLRLHFFPFVYRSYDLAAAARAFKVSTQDAVLLEDGIPAPEPGNASTSTSPQPARLEFLLDVARDTLVVSFVPLADGGIAFVNAVELVSVPDGLVADAADSSTGRPEPIPAVLPLQTAYRLNVGGPAVAPDDDALWREWTTDLRFLSHSVADAVTREVRYNGMLNRLPGQATATDAPDIVYATARELVINGSSFDGQKQMAWQFDVDTSSSYFIRFHFCDIVGKAPHQLHINAYVDDATVKQDLDLAAVGDGALAFPYYTDFVLPASEASGKLAVHVGPLANKIVMPAAILNGIEIMKMHLSAGSVVVVQPAAGAAKSRFAVVLGSVCGALAFISVAVALAIVLRKKEKEKEVEEGAKEQPTPTQSQSSTPWMPLLGRFSVRGAIASGSSSFTTAGNTPGASPRAAAAAAVMPSYRFPLAMLQDATRNFDDSLIIGEGGFGKVYGAVLQDGTKVAVKRASPESQQGAREFRTEIELLSGLRHRHLVSLVGYCDEREEMILLYEYMEHGSLRSRLYGRGGAAPLSWAQRLEACAGAARGLLYLHTAVDKPVIHRDVKSSNILLDGDLAGKVADFGLSKAGPVLDETHVSTAVKGSFGYVDPEYCRTRQLTAKSDVYSLGVVLLEAVCARPVVDPRLPKPMSNLVEWGLHWQGRGELEKIVDRRIAAAARPAALRKYGETVARCLAERGADRPAMEDVVWNLQFVMRLQEGDGLDFSDVSSLNMVTELTPPRRQRSAVDHDGLDYSDVNSLNMVTELTPPQTGSVEGDGEADDDFTDASMRGTFWQMVNVRSR

>Aet-CrRLK1L4(XP_020147554.1)

MRGGPRCALLLLVAAAALVPAARAQGATAPAPSSGVPFVPRDDILLDCGATGKGNDTDGRQWDGDAGSKYAPPKLASASAGAQDPSVPQVPYLTARVSAAPFTYSFPLGPGRKFLRLHFYPANYSNRNAADAFFSVSVPAAKVTLLSNFSAYQTTTALNFAYIVREFSVNVTGQNLDLTFTPEKGHPNAYAFINGIEVVSSPDLFDLATPQLVTGDGNSQPYEMDPAAALQTMYRLNVGGQAISPSKDSGGARSWDDDTPYIYGAGAGVSYQNDPSVAITYPDNVPGYVAPSDVYATARSMGPDKGVNMAYNLTWILQVDAGYQYLVRLHFCEIQSPYTKPNQRVFNIYLNNQTAMQGADVIQWADPNGIGTPVYKDYVVSTVGSGIMDFWVALHPDAETKPQYYDAILNGMEVFKLQLTNGSLVGLNPVPSADPPAHSGSGDKKSLVAPIVGGVIGGLAVLALGYCCFICKRRRKAAKASGMSDGHSGWLPLSLYGHSHTSSSAKSHATGSYASSLPSNLCRHFSFAEIKAATKNFDESRILGVGGFGKVYHGEIDGGTTKVAIKRGNPLSEQGIHEFQTEIEMLSKLRHRHLVSLIGYCEEKNEMILVYDYMAHGTLREHLYKTQNAPLSWRQRLEICIGAARGLHYLHTGAKHTIIHRDVKTTNILLDDKWVAKVSDFGLSKTGPSMDHTHVSTVVKGSFGYLDPEYFRRQQLTEKSDVYSFGVVLFEVLCARPALNPTLAKEEVSLAEWALHCQKKGILDQIVDPYLKGKIVPQCFKKFAETAEKCVADNGIERPSMGDVLWNLEFALQMQESAEESGSIGCGMSDEGTPLVMVGKKDPNDPSIDSSTTTTTTTSLSMGDQSVASIDSDGLTPSAVFSQIMNPKGR

>Aet-CrRLK1L5(XP_020150974.1)

MAVHVVPPLLLLLLATALPYTALAVFSPDFSFFLACGAGADVPFPSDNPTRTFVRDDGYLSQGRPAAVSASASSGAASNPLYAAARADSSAFSYRLAYPATAGASSFLVLRLHFFPFVPASSSTSLSSARFTVSVLDAYALLPTFSPPVAGVVKEFFVPRDGSKDFTIRFTPDAGSSAFVNAVELFSAPPELLWNNTAVPVDPVGSNDLPEWPLDALETVYRLNVGGPMVTKENDTLWRTWLPDGPYLFGAPGQSVVNSTSSPIMYDPSNGYTQDVAPDVVYRTQRAANVTDLLVATTPGLNFNVTWTFPAEQGSRYLVRLHFCDYEVVSSVVGVGIVFNVYVAQAIGTPALSPKDRARQSNEAFYMDYAARAPRAGNLTVSIGWLRQSSGGGILNGLEIMKLQSADPSLTVSHGLTKRSIIIIVLATVLGAAVLACAVLCFFVVRRTKRRQVAPPASTEDKESTQLPWSPYTQEGISGWADESTNRSSEGTTARMQRVSTKLHISLAELKAATDNFHDRNLIGVGGFGNVYKGALADGTPVAVKRAMRASKQGLPEFHTEIVVLSGIRHRHLVSLIGYCNEQAEMILVYEYMEKGTLRSHLYGGSDDEPPLSWKQRLEICIGAARGLHYLHCGYSENIIHRDVKSTNILLGTDDGGSTGGGAIIAKVADFGLSRIGPSLGETHVSTAVKGSFGYLDPEYFKTQQLTDRSDVYSFGVVLFEVLCARPVIDQSLDRDQINIAEWAVRMHGEGKLDKIADARIAGEVNDNSLRKFAETAERCLADYGADRPSMGDVLWNLEYCLQLQETHVNRDAFEDSGAVATQLPADVVVPRWVPSSTSLLMMDDADEAGLSMTELADSQVFSQLNARGEGR

>Aet-CrRLK1L6(XP_020174759.1)

MAVRGILLALLLAMVLPRAILAAFSPGFQYFLACGANSSVSFPSDSPANIFVPDAAYLSPAGAPAVSASSTLASPPALYAAARADISAFSYRLPSPASPDTSSFLVLRLHFFPFFPATSSQYVINILSARFNVSVADAYALLSSFSPPAAGVVKEFFVPRDLFGGHFHVTFTPDAGSTAFVNAIELFSAPPEMLWNGPVTPVGAVVKDDMDLWQRQPLETVYRLNVGGSKVIIENDTLWRTWLPDGPYLYDASGLSVVSNTSSPIIYNSTNGYTREVAPDVVYQTQRMANVTDLLAATTPGLNFNLTWTFPAVKGSRYLVRLHFCDYEVVSSVVGVGIVFNVYIAQAIGTPDLTPKAWATQSNEVFYMDYAARAPSAGNLTVSIGWSSQSSGGGILNGLEIMRLPPVDLSSRRYGRTKQRTIVITVSAVLGAAVLACVVLCFFGVPYTKYSGSGWAEQFMNRWSRERKTGGMESVSRKLHIALAKIKAATDNFHERNLIGVGGFGNVYKGVLGDGTPVAVKRAMRASQQGLPEFQTEIVVLSGIRHRHLVSLIGYCNEQAEMILVYEYMEKGTLRSHLYGSDEPALSWKQRLEICVGAARGLHYLHRGYAENIIHRDVKSTNILLGSDDGSTGGVIAKVADFGLSRIGPSFGETHVSTAVKGSFGYLDPGYFKTQQLTDRSDVYSFGVVLCARPVIDQSLDQSLINIAEWAVRMRGEGRLDKMADPRIAGEVDEESLLKFAETAEKCLAECWVDRPSMGDVLWNLEYCLQLQETNITGDELDNMRPSSTSLLMDETGLSMTNVADSKVFSQRSACGEGR

>Aet-CrRLK1L7(XP_020194435.1)

MPPVLDMLVRLLVASVLLGAASGAFTPADNYLVLCGTSASATVAAGRTFVGDARLPAKSLAAPQSVEANTSRTAVVPSGESELYRSARVFTAPASYTFAVKQPGRHFVRLHFFPFPYRSYDMVADAAFNVSVQGAVLVNGYTPKNGTAELREFSVNVTGDTLVIAFAPTGKLAFVNAIEVVSVPDELIADMARMVDGAVQYTGLSTQALETIHRINMGVPKITPGNDTLGRTWLPDQSFQVNTDLAQHKDAKPLTIKYDEKSALSSAYTAPAEVYATATRLSTAGETSTINVQFNISWRFDAPAGSDYLLRFHWCDIVSKAAMGMAFNVYVGGAVVLDNYEISRDTFNRLSIPVYKDFLLGAKDAKGAITVSIGSSTEDNALPDGFLNGLEIMSIVGSAGAGAAATSPRSSKVKIGIIAGSAVCGATLVMVLGFIAFKMLRGREPEKKKPADAWSPFSASALGSRSRSRSFSKSNGNTVLLGQNGAGAGYRIPFAALQEATGGFDEGMVIGEGGFGKVYKGTMRDETVVAVKRGNRRTQQGLHEFHTEIEMLSRLRHRHLVSLIGYCDERGEMILVYEYMAMGTLRSHLYGAGLPPLSWEQRLEACIGAARGLHYLHTGSAKAIIHRDVKSANILLDESFMAKVADFGLSKNGPELDKTHVSTKVKGSFGYLDPEYFRRQMLTEKSDVYSFGVVLLEALCARTVIDPTLPREMVSLAEWATPCLRNGQLDQIVDQRIAGTIRPGSLKKLADTAEKCLAEYGVERPTMGDVLWCLEFALQLQVGSSDSSDVDTMLPPAPPVPVKTPEVQRSLSAATVATDAAAMTTNLGDLDGMSLSGVFSNMIKSDEVR

>Aet-CrRLK1L8(XP_020150664.1)

MQSELHTARSRAMGGGPRCALLLLAAAAACAALVPAAWAQAPTAPAPSGAPFVPRDDILLDCGATGKGNDTDGREWRGDAGSKYAPPNLASADAGAQDPSVPQVPYLTARVSAAAFTYSFPLGPGRKFLRLHFYPANYSNRDAADAFFSVSVPAAKVTLLSNFSAYQTATALNFAYLVREFSVNVTGPTLDLTFTPEKGRPNAYAFINGIEVVSSPDLFDLATPLFVTGDGNNQPFPMDPGAALQTMYRLNVGGQAISPSKDSGGARSWDDDTPYIYGAGAGVTYPNDPNVTITYPDNVPGYVAPSDVYATARSMGIDKNVNLAYNLTWIVQVDAGFTYLVRLHFCEIQSPITKPNQRVFNIYLNNQTAVEGADVIQWVDPLSTGTPLYKDYVVSTVGSGIMDFWVALHPNTGSKPQYYDAILNGMEVFKLQLSNGSLAGPNPVPSADPPAHTGQEKKNSLVGPIAGGVIGGLVVLALGYCCFICKRRRKVAKDAGMSDGHSGWLPLSLYGNSHTSSSAKSHATGSIASSLPSNLCRHFSFAEIKAATKNFDESRILGVGGFGKVYQGEIDGGTTKVAIKRGNPLSEQGIHEFQTEIEMLSKLRHRHLVSLIGYCEDKNEMILVYDHMAHGTLREHLYKTQNAPLSWRQRLEICIGAARGLHYLHTGAKHTIIHRDVKTTNILLDEKWVAKVSDFGLSKTGPSMDHTHVSTVVKGSFGYLDPEYFRRQQLTEKSDVYSFGVVLFEVLCARPALNPTLAKEEVSLAEWALHCQKKGILDQIVDPYLKGKIVPQCFKKFAETAEKCVADNGIERPSMGDVLWNLEFALQMQESAEESGSFGCGISDEEGTPLVMAGKKDPNDPSIDSSTTTTTTTSLSMGDQSVASIDSDGLTPSAVFSQIMNPKGR

>Aet-CrRLK1L9(XP_020148915.1)

MLRMRILVLAAVSIVFANLQFLKAHGRELFLSCGSNATADADGRRWIGDMAPGLNFTLSSPGIAALLAGSSNGSEIMAPVYRSARFFTTTSWYDFSLLPGNYCVRLHFFPSTFRNFSANGSVFDVVANDFKLVSKFNVSEEIVWRNSVSNSAATAVVKEYFLAVNSSRLQIEFDPRPGSFAFVNAIEVMLTPDNSFNGTVNKVGGVDAHIPPELSGRAVETMYRLNIGGPALASSHDQYLHRPWYTDEAFMFSANAALIVSNTSAIKYVSSNDSSIAPIDVYETARIMGNNMVMDKRFNVTWRFLVHPNFDYLVRLHFCELVYDKPSQRIFKIYINNKTAAENYDVYNRAGGINKAYHEDYFDSLPQQVDSLWLQLGPDSMTSASGTDALLNGLEIFKLSRSGNLDYVLGHIDMGNKRGRSKGRSRIGLWEEVGIGSAAFVALASVALFSWCYVRRKRKAVNEEVPAGWHPLVLHEAMKSTTDARASKKAPLARNSSSIGHRMGRRFSIADIRAATKNFDESLVIGSGGFGKVYKGEVDDGITVAIKRANPLCGQGLKEFETEIEMLSKLRHRHLVAMIGYCEEQKEMILVYEYMAKGTLRSHLYGSGLPPLTWKQRIDACIGAARGLHYLHTGADRGIIHRDVKTTNILLDKNFVAKIADFGLSKTGPTLDQTHVSTAIRGSFGYLDPEYFRRQQLTQKSDVYSFGVVLFEVACARPVIDPSVPKDQINLAEWAMRWQRQRSLEAIADPRLDGDYSPESLKKFGDIAEKCLADDGRTRPSMGEVLWHLEYVLQLHEAYKRNVDCESFGSSELGFADMSFSMPHIREGEEEHHPKKSGIREDSAP

>Aet-CrRLK1L10(XP_020151474.1)

MGTTTGQKTALLLLGTLWVLLGTCNAAEFSPADNYLINCGSTVDANLHDGRVFKADNSGSTILTSHHSVPANTLPDAVISSDNPVLYQTARIFIVPSSYSFNMKSRGRHFVRLHFFGFRYQSYDLAAAKFKVSTQHVVLLDNFTPPSNSSPLVREYSLNITEDMLILSFVPLGNSTSFINAIEVISVPDDLIQDSAQTVNPSGQYLGLATQSFQTFYRINVGGREVTVVNDTLSRSWDTDQNFFINSTTTELFAYQGRLNYQKGAATKEDAPDSVYNTARRLAVQNRTSPASNMTWQFDVDGRSSYLIRFHFCDIVSKAAYSLYFDIYVDGGLALENLDLSEKVFGTLAVPYYTEFVLKSSNPSGKLSVGIGPSSLNNVALDGILNGLEIMKMNISTGTIYVVWPPATPKRKLAIILGPVLGGVGAVSIAIILCFVLRRKKKEKKPRRAPTSRPSSSWSPLTLNGLSFLSIGTRTTSRTTHTSGTNSDVSYRIPFALLQVATKHFDEQMVVGVGGFGKVYKAVLQDSTKVAVKRGNQKSHQGLKEFRTEIELLSGLRHRHLVSLIGYCDEQNEMILVYEYMEKGTLKSHLYGSDMPPLSWKKRVEICIGAARGLHYLHTGFAKSIIHRDVKSANILLDENLMAKVSDFGLSKTGPELDQTHVSTAVKGSFGYLDPEYYRRQKLTDKSDVYSFGVVLLEVICARPVIDPTLPRDMINLAEWAIKWQKRGELGQIVDQRIAGTIRPESLRKYGETVEKCLADYGVDRPTMGDVLWNLEFVLQLQEAGPDISNVDSMNQISELPSGARRMGSLEIGTADEADEGRTNMDYSQMSTNDAFSQLMNTEGR

>Aet-CrRLK1L11(XP_020184558.1)

MAAARGRGVLLAVLLLMMVAFAFVDADIYKPTDSILVHCGSDKDGQDEDGRKWTADKDSKWLPDGGKSSVMGTADVPDPSLPSPVPYMTARVFPKETAYTFPVADADRHWVRLHFYPAAYHGIPADHFFFSVTTSTGVTLLRNFSVYTTAKALTQAYIVREFSLPPSTTGSLSLKFTPTAMNNASYAFVNGIEIISMPSFFGDPATLVGLDDQSLDASAGNLQTMYRLSVGGSYIPPANDSGLSREWFSDTPYVYGAATGVTFEANDTIPIKYPTPADEYAAPVSIYDSFRHMGRDPKMNRNNNLTWVFEVDGNFTYLLRLHFCSLMEDKINQVVFAILVNNKTATTTGSADIIAWAKEKNPANPGAPGKGVPVFKDYAVFMPAAPAGSDTILWLTLRPDTATNPQFVNAFLNGLEVFKVSDASGNLAGPNPDISKMLAEAELGAVDGQFREKPSNVGALIGGAAGGAAAFGLVAAVCFVAYQSKRRRELSSSPSHSSSGWLPVYGGNSQTSVSKSSGGRSAVTLNPNITAMCRHFSFQEIKSATKGFDESLVIGVGGFGKVYRGVVDGDTKVAIKRSNPSSEQGVLEFQTEIEMLSKLRHKHLVSLIGCCEDNGEMILVYDYMAHGTLREHLYKSGKPPLPWRQRLEIVIGAARGLHYLHTGAKYTIIHRDVKTTNILVDEKWVAKVSDFGLSKTGPTVQNQTHVSTMVKGSFGYLDPEYFRRQKLTEKSDVYSFGVVLFEVLCGRPALNPSLPREQVSLADHALSCQRKGTLEEIIDPVLEGKIAPDCLKKFAETAEKCLADQGVDRPSMGDVLWNLEFALQQQDTFENGGKPPEVDDYSSSFTITPPSMEESLAANAAALSLISEDMDEEDIANSVIFSQIAKPTGR

>Aet-CrRLK1L12(XP_020175506.2)

MVDEPPNIARRRPGQTGLNLAWRHPPPLNPRPFLPPPQVSSAPFPLLRLAPPNQREGERAPPQPGPGHHNNKNQHDDVPGVGRCASRWRRRPRCRLERRHNGSVGVRVTRRSTAPKMVRRGALPLALLAVLATLTAVAGQGKPVTDNGSGGGAGPAKFTPKDAFYIDCGGTAAADTKDGKSFKTDAEANSLLSARDNIKVADDKADVPSHLYRSARVFKEEAVYNFPLTAPGWHFIRLYFFPIKSGEADLAAATFDVSTAVNVLLHGFTPEAKAVMKEYIVNATENKLELKFTPQSGSAFINAIEVVNAPDELISKTALTVSPLAETSGLSEAAYQVVCRLNVGGPPIGPVNDTLGRQWEDDEKYLNPKEAGTEVSVPTSAIKYPDAFPATKLVAPTAVYATARHMAESGVANQNFNVSWKVDVDPSFDYLVRLLFADIISTSANDLYFNVYINGRKAISALDLSTITGDLAAPYYKDFVVNSSVNTDGHIIIDVGPLGQDTGRNDALLNGAEVLKMSNSVGSLDGEYGVDGRMVDDGSGTRKVVAAVGFAMMFGAFAGLGCMVVKWHRRPQDWERRNSFSSWLLPIHTGQSFSNGKSKSGYTFSSTAGLGHFFTFAEMSEATKNFDESAIIGVGGFGNVYVGEINDPDEEGSRIKVAIKRGNPSSEQGINEFNTEIQMLSKLRHRHLVSLIGYCDEGEEMILVYEFMQHGPFRDHIYGGPEGLPTLSWKQRLEICIGAARGLHYLHTGTAHGIIHRDVKTTNILLDDKFVAKVADFGLSKDGPGMNQLHVSTAVKGSFGYLDPEYFRCQQLTDKSDVYSFGVVLLETLCARAPIDPQLPREQVSLAEWGLQWKRKGLIEKIMDPNLNGKVNPESLAKFAETAEKCLCEFGSDRLSMGDVLWNLEYALQLQEANPPEGATDADDADASIVSSASGVTTVPDQSTTSANELFAQLADMKGR

>Aet-CrRLK1L13(XP_045086556.1)

MAGAAERTSRPPSSSTPLSAACGRRGDPSPTRTDFYLANIMMGRRKLQVVTLAILCFWSSAGVCKAQTVDFKPADSYLVDCGSTKGTTVLGRDFAADGASPVTVSTSQDILAGTSANGVSSFDNPVLYQTARIFTSPSSYTFPIQKQGRHFVRLYFYPFIYQSYDLSTAKFTVSTQDVLLLSDFQQPDKTAPLFKEYSLNITRDQLVISFKPSNGIAFINAIEVVSVPDDLIADVANMVNPVQQYSGLTTQSLETVYRVNMGGPKVFPNNDTLSRTWQKDQKYILNPSVTKTAVYGKAIKYRKGGATPLTAPDIVYSTATELAASNTSNALFNMTWQFDVDAGFSYLIRFHFCDIVSKALNQLYFNAYVGGFFAQHDLDLSEQSVNQLATAIYVDVVLSSNDASSKLSISIGPSTLNNALPDGILNGLEIMKMGSGSGSAFTVGNNGSNKKLPIIIGSVLGVVGLLIIVLVVVLLCRRKKTDDKQHSKTWMPFSINGLTSLSTGSRTSYGTTLTSGLNGSYGYRFAFNVLQEATNNFDESWVIGVGGFGKVYKGALRDDTKVAVKRGNPKSQQGLNEFRTEIELLSRLRHRHLVSLIGYCDERNEMILVYEYMENGTVKSHLYGSDNPSLNWKQRLEICIGAARGLHYLHTGSAKAIIHRDVKSANILLDENLLAKVADFGLSKTGPELDQTHVSTAVKGSFGYLDPEYFRRQQLTEKSDVYSFGVVMLEVLCARPVIDPSLPREMVNLAEWGMKWQKRGELHQIVDQKLSGAIRPDSLRKFGETVEKCLADYGVERPSMGDVLWNLEYVLQLQDVDSSTVSDVNSMNRIVDLSSQVQHVGAMESISVTMAEDGALHEPDHDLSDVSMSRVFSQLIKAEGR

>Aet-CrRLK1L14(XP_020201562.1)

MKSSANFLSILVLLVFLAAENARAQPQPILINCGSDSTTSVDARTWIGDSSPSNNFTLSFPGAIASAAPAPAPGVDGEQDPYGDLYKTARVFNASSSYRLAVAPGSYFLRLHFSQQFANLGAQEPIFSVAANGLRLLSKFSVHGEISWRDSQINSTSSVIVKEYLLNVTSGKLGIEFTPDEGSFAFINAMEVLPVSGTSIFDSVNKVDAHGLKGPFSLDGDGIETMYRLCVGCIDVLPRKEDPGLWRRWDKDEHFIFSLNAANSIFNSSNISYVSADDPTVAPLRLYQSARVPTESSVLGKKFNVSWSFNIDPGFDYLVRLHFCELQYDKAEQRKFKIYINNKTAAESYDVFARAGGKNKAFYEDFLDAASPQMDTLWVQLGAESSAGSAAADALLNGMEIFKVSREGNLAHPTVRIGGISGGASKPKRSPKWVLIGTASGLIIFIAIAGGLYFGFNLRRKKNSSASKAKDNLHGATHTRSPTLRTAGAFGSNRMGRRFTIAEIRTATVNFDESLVIGVGGFGKVYKGIMEDGTRVAIKRGHTDSHQGQGVKEFETEIEMLSRLRHRHLVPLIGYCDEQNEMVLVYEHMANGTLRSHLYGSDLPALTWKQRLEICIGAARGLHYLHTGLDRGIIHRDVKTTNILLDDNLVAKMADFGISKDGPALDHTHVSTAVKGSFGYLDPEYYRRQQLTPSSDVYSFGVVLFEVLCARPVINPTLPRDQINLADWALNRQRHRLLETIIDLRLDGNYTLASVKKSSKIAEKCLADEGVNRPSMGEVLWHLESALQLEQGHPQSTNADGCSDPQLKPSDVPTRVACIKEDEQSTRPGSHNSDGQVVDVKIEVP

>At-FER(AT3G51550.1)

MKITEGRFRLSLLLLLLLISAATLISAADYSPTEKILLNCGGGASNLTDTDNRIWISDVK

SKFLSSSSEDSKTSPALTQDPSVPEVPYMTARVFRSPFTYTFPVASGRKFVRLYFYPNSY

DGLNATNSLFSVSFGPYTLLKNFSASQTAEALTYAFIIKEFVVNVEGGTLNMTFTPESAP

SNAYAFVNGIEVTSMPDMYSSTDGTLTMVGSSGSVTIDNSTALENVYRLNVGGNDISPSA

DTGLYRSWYDDQPYIFGAGLGIPETADPNMTIKYPTGTPTYVAPVDVYSTARSMGPTAQI

NLNYNLTWIFSIDSGFTYLVRLHFCEVSSNITKINQRVFTIYLNNQTAEPEADVIAWTSS

NGVPFHKDYVVNPPEGNGQQDLWLALHPNPVNKPEYYDSLLNGVEIFKMNTSDGNLAGTN

PIPGPQVTADPSKVLRPTTRKSKSNTAIIAGAASGAVVLALIIGFCVFGAYRRRKRGDYQ

PASDATSGWLPLSLYGNSHSAGSAKTNTTGSYASSLPSNLCRHFSFAEIKAATKNFDESR

VLGVGGFGKVYRGEIDGGTTKVAIKRGNPMSEQGVHEFQTEIEMLSKLRHRHLVSLIGYC

EENCEMILVYDYMAHGTMREHLYKTQNPSLPWKQRLEICIGAARGLHYLHTGAKHTIIHR

DVKTTNILLDEKWVAKVSDFGLSKTGPTLDHTHVSTVVKGSFGYLDPEYFRRQQLTEKSD

VYSFGVVLFEALCARPALNPTLAKEQVSLAEWAPYCYKKGMLDQIVDPYLKGKITPECFK

KFAETAMKCVLDQGIERPSMGDVLWNLEFALQLQESAEENGKGVCGDMDMDEIKYDDGNC

KGKNDKSSDVYEGNVTDSRSSGIDMSIGGRSLASEDSDGLTPSAVFSQIMNPKGR

>At-HERK1(AT3G46290.1)

MGIEKFETFILISTISILLCICHGFTPVDNYLINCGSPTNGTLMGRIFLSDKLSSKLLTS

SKEILASVGGNSGSDIYHTARVFTEVSSYKFSVTRGRHWVRLYFNPFDYQNFKMGSAKFA

VSSQSHVLLSDFTVTSSKVVKEYSLNVTTNDLVLTFTPSSGSFAFVNAIEVISIPDTLIT

GSPRFVGNPAQFPDMSMQGLETIHRVNMGGPLVASNNDTLTRTWVPDSEFLLEKNLAKSM

SKFSTVNFVPGYATEDSAPRTVYGSCTEMNSADNPNSIFNVTWEFDVDPGFQYYFRFHFC

DIVSLSLNQLYFNLYVDSMVAATDIDLSTLVDNTLAGAYSMDFVTQTPKGSNKVRVSIGP

STVHTDYPNAIVNGLEIMKMNNSKGQLSTGTFVPGSSSSSKSNLGLIVGSAIGSLLAVVF

LGSCFVLYKKRKRGQDGHSKTWMPFSINGTSMGSKYSNGTTLTSITTNANYRIPFAAVKD

ATNNFDESRNIGVGGFGKVYKGELNDGTKVAVKRGNPKSQQGLAEFRTEIEMLSQFRHRH

LVSLIGYCDENNEMILIYEYMENGTVKSHLYGSGLPSLTWKQRLEICIGAARGLHYLHTG

DSKPVIHRDVKSANILLDENFMAKVADFGLSKTGPELDQTHVSTAVKGSFGYLDPEYFRR

QQLTDKSDVYSFGVVLFEVLCARPVIDPTLPREMVNLAEWAMKWQKKGQLDQIIDQSLRG

NIRPDSLRKFAETGEKCLADYGVDRPSMGDVLWNLEYALQLQEAVIDGEPEDNSTNMIGE

LPPQINNFSQGDTSVNVPGTAGRFEESSIDDLSGVSMSKVFSQLVKSEGR

>At-HERK2(AT1G30570.1)

MSKLRKKYLEHLLCVLIFFTYVIGYGEAQSKSFLVDCGSNATTEVDGRTWVGDLSPNKSV

TLQGFDAITASTSKGSSVYAEIYKTARVFDAVLNYTFEGITQGNYFVRLHFSPFAIENHN

VNESSFSVFADGLRLMLDINIAGEIAHKNLILESTGHNATASSLVKEFLLPTGPGKLVLS

FIPEKGSFGFVNAIEIVSVDDKLFKESVTKVGGSEVELGLGGRGIETMYRLNVGGPKLGP

SKDLKLYRTWETDLSYMVIENAGVEVKNSSNITYALADDSPVAPLLVYETARMMSNTEVL

EKRFNISWKFEVDPNFDYLVRLHFCELLVDKQNQRIFRIYINNQTAAGNFDIFAHAGGKN

KGIYQDYLDPVSSKNDVLWIQLGPDSSVGASGDALLSGLEIFKLSKNGNLAHLIRFDSTG

HSVSDSKMRIIWISVGAGIAIIIFFVFLGILVVCLCKKRRSKSDESKNNPPGWRPLFLHV

NNSTANAKATGGSLRLNTLAASTMGRKFTLAEIRAATKNFDDGLAIGVGGFGKVYRGELE

DGTLIAIKRATPHSQQGLAEFETEIVMLSRLRHRHLVSLIGFCDEHNEMILVYEYMANGT

LRSHLFGSNLPPLSWKQRLEACIGSARGLHYLHTGSERGIIHRDVKTTNILLDENFVAKM

SDFGLSKAGPSMDHTHVSTAVKGSFGYLDPEYFRRQQLTEKSDVYSFGVVLFEAVCARAV

INPTLPKDQINLAEWALSWQKQRNLESIIDSNLRGNYSPESLEKYGEIAEKCLADEGKNR

PMMGEVLWSLEYVLQIHEAWLRKQNGENSFSSSQAVEEAPESFTLPACSNQDSSETEQSQ

TGSALHNSA

>At-THE1(AT5G54380.1)

MVFTKSLLVLLWFLSCYTTTTSSALFNPPDNYLISCGSSQNITFQNRIFVPDSLHSSLVL

KIGNSSVATSTTSNNSTNSIYQTARVFSSLASYRFKITSLGRHWIRLHFSPINNSTWNLT

SASITVVTEDFVLLNNFSFNNFNGSYIFKEYTVNVTSEFLTLSFIPSNNSVVFVNAIEVV

SVPDNLIPDQALALNPSTPFSGLSLLAFETVYRLNMGGPLLTSQNDTLGRQWDNDAEYLH

VNSSVLVVTANPSSIKYSPSVTQETAPNMVYATADTMGDANVASPSFNVTWVLPVDPDFR

YFVRVHFCDIVSQALNTLVFNLYVNDDLALGSLDLSTLTNGLKVPYFKDFISNGSVESSG

VLTVSVGPDSQADITNATMNGLEVLKISNEAKSLSGVSSVKSLLPGGSGSKSKKKAVIIG

SLVGAVTLILLIAVCCYCCLVASRKQRSTSPQEGGNGHPWLPLPLYGLSQTLTKSTASHK

SATASCISLASTHLGRCFMFQEIMDATNKFDESSLLGVGGFGRVYKGTLEDGTKVAVKRG

NPRSEQGMAEFRTEIEMLSKLRHRHLVSLIGYCDERSEMILVYEYMANGPLRSHLYGADL

PPLSWKQRLEICIGAARGLHYLHTGASQSIIHRDVKTTNILLDENLVAKVADFGLSKTGP

SLDQTHVSTAVKGSFGYLDPEYFRRQQLTEKSDVYSFGVVLMEVLCCRPALNPVLPREQV

NIAEWAMAWQKKGLLDQIMDSNLTGKVNPASLKKFGETAEKCLAEYGVDRPSMGDVLWNL

EYALQLEETSSALMEPDDNSTNHIPGIPMAPMEPFDNSMSIIDRGGVNSGTGTDDDAEDA

TTSAVFSQLVHPRGR

>At-ANX1(AT3G04690.1)

MSGKTRILFFLTCLSFLLVFPTRSNGQDLALSCGTSEASADQDKKKWEPDTKFLKTGNSI

HATATYQDPSLLSTVPYMTARIFTAPATYEIPIKGDKRHLLRLYFYPSTYTGLNISNSYF

TVEANDVTLLSNFSAAITCQALTQAYLVKEYSLAPTDKDVLSIKFTPSDKYRDAFAFING

IEVIQMPELFDTAALVGFTDQTMDAKTANLQSMFRLNVGGQDIPGSQDSGGLTRTWYNDA

PYIFSAGLGVTLQASNNFRINYQNMPVSIAPADIYKTARSQGPNGDINLKSNLTWMFQID

KNFTYILRLHFCEFQLSKINQKVFNIYINNRTAQADTTPADIIGWTGEKGIPMYKDYAIY

VDANNGGEEITLQMTPSTFGQPEYYDSSLNGLEIFKMDTMKNLAGPNPEPSPMQAEEEVK

KEFKNEKRHAFIIGSAGGVLAVLIGALCFTAYKKKQGYQGGDSHTSSWLPIYGNSTTSGT

KSTISGKSNNGSHLSNLAAGLCRRFSLPEIKHGTQNFDDSNVIGVGGFGKVYKGVIDGTT

KVAVKKSNPNSEQGLNEFETEIELLSRLRHKHLVSLIGYCDEGGEMCLVYDYMAFGTLRE

HLYNTKKPQLTWKRRLEIAIGAARGLHYLHTGAKYTIIHRDVKTTNILVDENWVAKVSDF

GLSKTGPNMNGGHVTTVVKGSFGYLDPEYFRRQQLTEKSDVYSFGVVLFEILCARPALNP

SLPKEQVSLGDWAMNCKRKGNLEDIIDPNLKGKINAECLKKFADTAEKCLNDSGLERPTM

GDVLWNLEFALQLQETADGTRHRTPNNGGSSEDLGRGGMAVNVAGRDDVSDLSSEDNTEI

FSQIVNPKGR

>At-ANX2(AT5G28680.2)

MNEKLRILFSFLCFFYVLLVSPSQSNGQDISLSCGASEPAVDQDKKKWEPDTKFLKTPNT

VHAPATYQDPSLLSTVPYMTSRIFTAPATYEIPVKGDKRHMLRLHFYPSTYTGLNILDSY

FSVAANDLTLLSNFSAAITCQALTQAYLVREYSLAPSEKDVLSIIFTPSDKHPKAFAFIN

GIEVIPMPELFDTASLVGFSDQTSDTKTANLQTMFRLNVGGQDIPGSQDSGGLTRTWYND

APYIFSAGLGVTLQASNNFRIDYQKMPVSTAPADVYKTARSQGPNGDINMKSNLTWMFQV

DTNFTYIMRLHFCEFQLAKINQKVFNIFINNRTAQGDTNPADILGWTGGKGIPTYKDYAI

YVDANTGGGGEEISLQMTPSTFGQPEYYDSQLNGLEIFKIDTMKNLAGPNPKPSPMQANE

DVKKDFQGDKRITAFVIGSAGGVAAVLFCALCFTMYQRKRKFSGSDSHTSSWLPIYGNSH

TSATKSTISGKSNNGSHLSNLAAGLCRRFSLSEIKHGTHNFDESNVIGVGGFGKVYKGVI

DGGTKVAIKKSNPNSEQGLNEFETEIELLSRLRHKHLVSLIGYCDEGGEMCLIYDYMSLG

TLREHLYNTKRPQLTWKRRLEIAIGAARGLHYLHTGAKYTIIHRDVKTTNILLDENWVAK

VSDFGLSKTGPNMNGGHVTTVVKGSFGYLDPEYFRRQQLTEKSDVYSFGVVLFEVLCARP

ALNPSLSKEQVSLGDWAMNCKRKGTLEDIIDPNLKGKINPECLKKFADTAEKCLSDSGLD

RPTMGDVLWNLEFALQLQETADGSRHRTPSNGGGSVDLGGGGGGVTVNISAGESDLGDDL

SSEENSGIFSQIVNPKGR

>At-BUPS1(AT4G39110.1)

MEIRKKPNIFTVLVIDFSSKPSMALLLAILLFLSGPSASAVAAAAVGPATGFKPADDILI

DCGSKSSSKTPDGRVFKSDQETIQYIEAKEDIQVSAPPSDKVASPIYLTARIFREEATYK

FHLTRPGWHWVRLHFLAFPNDKFDLQQATFSVLTEKYVLLHNFKISNNNNDSQAAVQKEY

LVNMTDAQFALRFRPMKSSAAFINAIEVVSAPDELISDSGTALFPVIGFSGLSDYAYQSV

YRVNVGGPLIMPQNDTLGRTWIPDKEFLKDENLAKDVKTTPSAIKYPPEVTPLIAPQTVY

ATAVEMANSLTIDPNFNVSWNFPSNPSFNYLIRLHFCDIVSKSLNDLYFNVYINGKTAIS

GLDLSTVAGNLAAPYYKDIVVNATLMGPELQVQIGPMGEDTGTKNAILNGVEVLKMSNSV

NSLDGEFGVDGRTTGMGKHGMVATAGFVMMFGAFIGLGAMVYKWKKRPQDWQKRNSFSSW

LLPIHAGDSTFMTSKGGSQKSNFYNSTLGLGRYFSLSELQEATKNFEASQIIGVGGFGNV

YIGTLDDGTKVAVKRGNPQSEQGITEFQTEIQMLSKLRHRHLVSLIGYCDENSEMILVYE

FMSNGPFRDHLYGKNLAPLTWKQRLEICIGSARGLHYLHTGTAQGIIHRDVKSTNILLDE

ALVAKVADFGLSKDVAFGQNHVSTAVKGSFGYLDPEYFRRQQLTDKSDVYSFGVVLLEAL

CARPAINPQLPREQVNLAEWAMQWKRKGLLEKIIDPHLAGTINPESMKKFAEAAEKCLED

YGVDRPTMGDVLWNLEYALQLQEAFTQGKAEETENAKPDVVTPGSVPVSDPSPITPSVTT

NEAATVPVPAKVEENSGTAVDEHSGTAMFTQFANLNGR

>At-BUPS2(AT2G21480.1)

MEIRKKPNIPMCLVLDSSSRPFMTLLFTILLFLTGLASAVGAVGGSPTAGFKPADDILID

CGSKSSTKTPEGRVFKSDSETVQYIEAKDDIQVSAPPSDKLPSPIYLTAKIFREEAIYKF

HLTRPGWHWVRLHFFAFPNDKFDLQQATFSVLTEKYVLLHNFKLSNDNNDSQATVQKEYL

LNMTDAQFALRFKPMKGSAAFINGIELVSAPDELISDAGTSLFPVNGFSGLSDYAYQSVY

RVNVGGPLITPQNDTLGRTWTPDKEYLKDENLAKDVKTNPTAIIYPPGVTPLIAPQTVYA

TGAEMADSQTIDPNFNVTWNFPSNPSFHYFIRLHFCDIISKSLNDLYFNVYINGKTAISG

LDLSTVAGDLSAPYYKDIVVNSTLMTSELQVQIGPMGEDTGKKNAILNGVEVLKMSNSVN

SLDGEFGVDGQRASMGKQGMVATAGFVMMFGAFVGLGAMVYKWKKRPQDWQKRNSFSSWL

LPIHAGDSTFMTSKTGSHKSNLYNSALGLGRYFSLSELQEVTKNFDASEIIGVGGFGNVY

IGTIDDGTQVAIKRGNPQSEQGITEFHTEIQMLSKLRHRHLVSLIGYCDENAEMILVYEY

MSNGPFRDHLYGKNLSPLTWKQRLEICIGAARGLHYLHTGTAQGIIHRDVKSTNILLDEA

LVAKVADFGLSKDVAFGQNHVSTAVKGSFGYLDPEYFRRQQLTDKSDVYSFGVVLLEALC

ARPAINPQLPREQVNLAEWAMLWKQKGLLEKIIDPHLVGAVNPESMKKFAEAAEKCLADY

GVDRPTMGDVLWNLEYALQLQEAFSQGKAEAEEVETPKPVAVPAAAPTSPAATTAAASER

PVSQTEEKDDSTVDQHSGTTMFTQFASLNGR

>At-CURVY1(AT2G39360.1)

MINLKLFLELKLCFLITLLCSSHISSVSDTFFINCGSPTNVTVNNRTFVSDNNLVQGFSV

GTTDSNSGDESTLFQTARVFSDESSSTYRFPIEEHGWFLIRIYFLPLVSASQDLTTARFS

VSAQNFTLIREYKPSTTSVVREYILNVTTDSLLLQFLPRTGSVSFINALEVLRLPETLIP

EDAKLIGTQKDLKLSSHAMETVSRVNMGNLSVSRDQDKLWRQWDSDSAYKAHFGTPVMNL

KAVNFSAGGITDDIAPVYVYGTATRLNSDLDPNTNANLTWTFKVEPGFDYFVRFHFCNII

VDPFGFERQIRFDIFVNSEKVRTIDMTEVLNGTFGAPFFVDAVMRKAKSREGFLNLSIGL

VMDVSSYPVSFINGFEISKLSNDKRSLDAFDAILPDGSSSNKSSNTSVGLIAGLSAALCV

ALVFGVVVSWWCIRKRRRRNRQMQTVHSRGDDHQIKKNETGESLIFSSSKIGYRYPLALI

KEATDDFDESLVIGVGGFGKVYKGVLRDKTEVAVKRGAPQSRQGLAEFKTEVEMLTQFRH

RHLVSLIGYCDENSEMIIVYEYMEKGTLKDHLYDLDDKPRLSWRQRLEICVGAARGLHYL

HTGSTRAIIHRDVKSANILLDDNFMAKVADFGLSKTGPDLDQTHVSTAVKGSFGYLDPEY

LTRQQLTEKSDVYSFGVVMLEVVCGRPVIDPSLPREKVNLIEWAMKLVKKGKLEDIIDPF

LVGKVKLEEVKKYCEVTEKCLSQNGIERPAMGDLLWNLEFMLQVQAKDEKAAMVDDKPEA

SVVGSTMQFSVNGVGDIAGVSMSKVFAQMVREETR

>At-MDS2(AT5G39000.1)

MIRHALLIFSILVSTPIVGEGATSTYEPTDVFLFNCGDTSNNVDVSGRNWTAENQKILSS

NLVNASFTAQASYQESGVSQIPYMTARIFRSEFTYSFPVTPGSNFLRLYFYPTRYGSQFN

AVKSFFSVKVNGFTLLNNFSADLTVKASKPQTEFIIKEFIIPVYQTLNLTFTPSLDSLAF

VNGIEIVSIPNRFYSKGGFDDVITNVGSSVDFHIENSTAFETVYRLNVGGKTVGDSGMFR

RWVSDDEIILSESSGISPIVPDIKINYTEKTPSYVAPDDVYATSRSMGNADHPEQNLNFN

LTWLFTVDAGFSYLVRLHFCETLSEVNKEGQRVFSIFIENQTATLEMDVFRMSGGSWIPM

YLDYTVIAGSGSGRRHDLRLDLHPLVSINPKYYDAILNGVEILKMNDPDGNLAGPNPDPL

VSPDLIPNRATPRIRKNKSHILPITLAVVGSLVVLAMFVVGVLVIMKKKKKSKPSTNSSW

CPLPHGTDSTNTKPAKSLPADLCRRFSIFEIKSATNDFEDKLIIGVGGFGSVYKGQIDGG

ATLVAVKRLEITSNQGAKEFETELEMLSKLRHVHLVSLIGYCDEDNEMVLVYEYMPHGTL

KDHLFRRDKTSDPPLSWKRRLEICIGAARGLQYLHTGAKYTIIHRDIKTTNILLDENFVT

KVSDFGLSRVGPTSASQTHVSTVVKGTFGYLDPEYYRRQVLTEKSDVYSFGVVLLEVLCC

RPIRMQSVPPEQADLIRWVKSNYRRGTVDQIIDSDLSADITSTSLEKFCEIAVRCVQDRG

MERPPMNDVVWALEFALQLHETAKKKNDNVESLDLMPSGEVGTTTDGEDDLFSRTTGHVG

KSTTTDDSVLVVGDERSGSSWGVFSEINEPKAR

>At-MDS3(AT5G39020.1)

MNCNVLFLLSVLVSVTAGVTAAYHPTDVFLFNCGDTSNNVDNSGRNWTVESRQILSSNLV

NASFTSEASYQKAGVSRIPYMKARIFRSEFTYSFPVTPGSIFLRLYFYPTQYKSGFDAVN

SFFSVKVNGFTLLRNFNADSTVQASIPLSNSLIKEFIIPVHQTLNLTFTPSKNLLAFVNG

IEIVSMPDRFYSKGGFDNVLRNVSSDVDFQIDNSTAFESVHRLNVGGQIVNEVDDSGMFR

RWLSDDSFGNSGSIVNVPGVKINYTEKTPAYVAPYDVYATSRLMGNSSNLMFNLTGMFLT

VDAGYNYLVRLHFCETLPQVTKAGQRVFSIFVEDKMAKKETDVIRLSGGPRIPMYLDFSV

YVGFESGMIQPELRLDLVPLKDTNQTYYDAILSGVEILKLNDSDGNLARPNPELLVSTDS

TPDDSNVTPPIKGKPHVLVIILIVVGSVIGLATFIVIIMLLIRQMKRKKNKKENSVIMFK

LLLKQYIYAELKKITKSFSHTVGKGGFGTVYRGNLSNGRTVAVKVLKDLKGNGDDFINEV

TSMSQTSHVNIVSLLGFCYEGSKRAIISEFLEHGSLDQFISRNKSLTPNVTTLYGIALGI

ARGLEYLHYGCKTRIVHFDIKPQNILLDDNFCPKVADFGLAKLCEKRESILSLIDTRGTI

GYIAPEVVSRMYGGISHKSDVYSYGMLVLDMIGARNKVETTTCNGSTAYFPDWIYKDLEN

GDQTWIIGDEINEEDNKIVKKMILVSLWCIRPCPSDRPPMNKVVEMIEGSLDALELPPKP

SRHISTELVLESSSLSDGQEAEKQTQTLDSTII

>At-ANJEA(AT5G59700.1)

MGGEKFGFLIWILSIPCLIFLCYGYVPVDNYLINCGSSTNVTVTSRVFISDNLASNFLTS

PNEILAASNRNSNSDIYQTARIFTGISKYRFSVARGRHWIRLHFNPFQYQNFQMVSAKFS

VSSETHVLLSDFTVSSRVMKEYSLNVATDHLELTFTPSGDSFAFLNALEVVSVPDTLFSG

DPSFAGSPGKFQGLSWQALETVYRVNMGGPRVTPSNDTLSRIWEPDSEFLVEKNLVKSVS

KIASVDYVPGFATEETAPRTVYGTCTEMNSADNPSSNFNVTWDFDVDPGFQYFLRFHFCD

IVSKALNQLYFNLYVDSMDVVENLDLSSYLSNTLSGAYAMDFVTGSAKLTKRIRVSIGRS

SVHTDYPTAILNGLEIMKMNNSKSQLSIGTFLPSGSSSTTKKNVGMIIGLTIGSLLALVV

LGGFFVLYKKRGRDQDGNSKTWIPLSSNGTTSSSNGTTLASIASNSSYRIPLVAVKEATN

SFDENRAIGVGGFGKVYKGELHDGTKVAVKRANPKSQQGLAEFRTEIEMLSQFRHRHLVS

LIGYCDENNEMILVYEYMENGTLKSHLYGSGLLSLSWKQRLEICIGSARGLHYLHTGDAK

PVIHRDVKSANILLDENLMAKVADFGLSKTGPEIDQTHVSTAVKGSFGYLDPEYFRRQQL

TEKSDVYSFGVVMFEVLCARPVIDPTLTREMVNLAEWAMKWQKKGQLEHIIDPSLRGKIR

PDSLRKFGETGEKCLADYGVDRPSMGDVLWNLEYALQLQEAVVDGDPEDSTNMIGELPLR

FNDYNHGDTSVNFSVAKEGRFDEEESSVDDSSGVSMSKVFSQLIKSEGR

>At-CAP1(AT5G61350.1)

MGGDFRHFSSHVSLLLLFLLIVKSSSSFTPADNYLIDCGSSDETKLSDGRNFKSDQQSVA

FLQTDEDIKTSVDSIPITDSNASTLPLYLTARIFAGKSTYSFYISRPGRHWIRLHFYPLN

HPLYNLTNSVFSVTTDTTVLLHDFSAGDTSSIVFKEYLIYAAEKLSLYFKPHKGSTAFIN

AVEIVSVPDELVPDSASSVPQAPDFKGLSSFSLEILHRINIGGDLISPKIDPLSRTWLSD

KPYNTFPEGSRNVTVDPSTITYPDGGATALIAPNPVYATAEEMADAQTSQPNFNLSWRMS

VDFGHDYFIRLHFCDIVSKSLNDLIFNVFINKLSAISALDLSSLTSALGTAYYADFVLNA

STITNGSILVQVGPTPNLQSGKPNAILNGLEIMKLNNAAGSLDGLFGVDGKYKGPIGGMS

SKKLAIAGIGFVMALTAFLGVVVLLVRWQRRPKDWQKQNSFSSWLLPLHASHSSYISSKG

GSTSRRMSIFGSKKSKSNGFSSFFSNQGLGRYFPFTELQTATQNFDENAVCGVGGFGKVY

IGEIDGGTQVAIKRGSQSSEQGINEFQTEIQMLSKLRHRHLVSLIGFCDENKEMILVYEY

MSNGPLRDHLYGSKENDPNPIPTLSWKQRLEICIGSARGLHYLHTGAAQGIIHRDVKTTN

ILLDENLVAKVSDFGLSKDAPMDEGHVSTAVKGSFGYLDPEYFRRQQLTDKSDVYSFGVV

LFEVLCARPVINPQLPREQVNLAEYAMNLHRKGMLEKIIDPKIVGTISKGSLRKFVEAAE

KCLAEYGVDRPGMGDVLWNLEYALQLQEASAQVDLSEDKTTMNIEMDLIPGEEMQSPSHS

IP

>At-MDS1(AT5G38990.1)

MICHVLVIFTILVSAVVDATASYEPTDVFLINCGDTSNNMDYSGRNWTTENPKFMSSNAV

DDASFTSSASYQESGIPQVPYLKARIFRYDFTYSFPVSPGWKFLRLYFYPTRYGSDFDAV

KSFFSVNVNRFTLLHNFSVKASIPESSSLIKEFIVPVNQTLDLTFTPSPNSLAFVNGIEI

ISMPDRFYSKGGFDDVVRNVGRDVDFEIDNSTAFETVYRVNVGGKVVGDVGDSGMFRRWL

SDEGFLLGINSGAIPNITGVKINYTDKTPAYVAPEDVYTTCRLMGNKDSPELNLNFNLTW

LFEVDAGFAYIVRLHFCETQPEVNKTGDRVFSIFFGYQLAMREMDVFRLSGGFRLPMYLD

FKVLVDADGTSQRPSLRVDLTPYKEDYPTYYDAILSGVEILKLSNSDGNLAGLNPIPQLS

PPPQSITPLKGKGKSSHVLPIIIAVVGSAVALAFFVLVVVLVVMKRKKKSNESSVDTTNK

PSTNSSWGPLLHGTGSTNTKSASSLPSDLCRRFSIYEIKSATNDFEEKLIIGVGGFGSVY

KGRIDGGATLVAVKRLEITSNQGAKEFDTELEMLSKLRHVHLVSLIGYCDDDNEMVLVYE

YMPHGTLKDHLFRRDKASDPPLSWKRRLEICIGAARGLQYLHTGAKYTIIHRDIKTTNIL

LDENFVAKVSDFGLSRVGPTSASQTHVSTVVKGTFGYLDPEYYRRQILTEKSDVYSFGVV

LLEVLCCRPIRMQSVPPEQADLIRWVKSNFNKRTVDQIIDSDLTADITSTSMEKFCEIAI

RCVQDRGMERPPMNDVVWALEFALQLHETAKKKNDNVESLDLMPSGEVGTTTDGEDDLFS

RTTGHVGKSTTTDDSVLVVGDERSGSSWGVFSEINEPKAR

>At-MDS4(AT5G39030.1)

MICFILFVFSFLVSVSATAPYKPDDVFLINCGETDVPFDNHGRTWTQEEKNILPKNSDNA

SFSSVVSYKEESGIPQVPYMTARIFRSDFTYSFPVSPGWKFLRLYFYPTSYKSGFDAVNS

FVSVTVNDFTLLQNFSADLTVKASIPESKSLIKEFIVPVYLTLNLTFRPSNNSLAFVNGI

EIVSMPDRFYSKGGFDDLITNVGSLIDFEIDNSTASETVHRLNVGGHMVDEVNDSGMFRR

WLSDDYEFLIGGVSPYMPDVNISYTEKTPAYVAPAYVYSTCRMMGNAQDTYLNLNFNLTW

LFTVDAGFSYLVRLHFFEKYLNKANQRVFSIFLGNQMAREEMDVIRLSGGPRIPIYLDFR

IYVGSESGPRPDLRLDLHPLVKDNPEYYEAILNGVEILKLNNSGNLAIIQDNELKPNPPL

SSNLTPNHVTQQIKGKSSHLLVKIFIAVGPGTGLATFVVVLMLWMRQMKRKNRKEERVVM

FKKLLNMYTYAELKKITKSFSYIIGKGGFGTVYGGNLSNGRKVAVKVLKDLKGSAEDFIN

EVASMSQTSHVNIVSLLGFCFEGSKRAIVYEFLENGSLDQFMSRNKSLTQDVTTLYGIAL

GIARGLEYLHYGCKTRIVHFDIKPQNILLDGNLCPKVSDFGLAKLCEKRESVLSLMDTRG

TIGYIAPEVFSRMYGRVSHKSDVYSFGMLVIDMIGARSKEIVETVDSAASSTYFPDWIYK

DLEDGEQTWIFGDEITKEEKEIAKKMIVVGLWCIQPCPSDRPSMNRVVEMMEGSLDALEI

PPKPSMHISTEVITESSSLSDGGEDV

>AT5G24010.1

MAFPINLTQTLLFFFCPLLHLSFAAFTPTDNYLINSGSNTNTSFFTTRSFLSDSSEPGSS

FLSTDRSISISDTNPSPDSPVLYNTARVFPVGGSYKFQVTTKGTHFIRLHFAPFKASRFN

LRSAKFRVLINGFSVINSFSTSSVVVKEFILKIDDPVLEISFLPFKASGFGFVNAVEVFS

APKDYIMDQGTKLVIPNSAQIFSNLSSQVLETVHRINVGGSKLTPFNDTLWRTWVVDDNY

LLLRAAARRAWTTHSPNYQNGGATREIAPDNVYMTAQEMDRDNQELQARFNISWGFQVDE

KRVLHLVRLHFCDIVSSSLNQLYFNVFINEYLAFKDVDLSTLTFHVLASPLYIDFVAESD

RSGMLRISVGPSDLSNPARVNALLNGVEIMRILSPVSSEVVSGKRNVVWIVVGSVLGGFV

FLSLFFLSVLCLCRRKNNKTRSSESTGWTPLRRFRGSSNSRTTERTVSSSGYHTLRISFA

ELQSGTNNFDRSLVIGVGGFGMVFRGSLKDNTKVAVKRGSPGSRQGLPEFLSEITILSKI

RHRHLVSLVGYCEEQSEMILVYEYMDKGPLKSHLYGSTNPPLSWKQRLEVCIGAARGLHY

LHTGSSQGIIHRDIKSTNILLDNNYVAKVADFGLSRSGPCIDETHVSTGVKGSFGYLDPE

YFRRQQLTDKSDVYSFGVVLFEVLCARPAVDPLLVREQVNLAEWAIEWQRKGMLDQIVDP

NIADEIKPCSLKKFAETAEKCCADYGVDRPTIGDVLWNLEHVLQLQESGPLNIPEEDYGD

VTDPRTARQGLSNGSNIERDYGDGTSGIISSTQVFSQLMTNAGR

>AT2G23200.1

MENFCFQDSVSLFITIMVLVLLPRLSLSDTSTYTRPENFYVNCGSDSNVFYGGQTFVGDT

NSSTNSVSFTNKGTEVINDQSSVAPEIYRTVRIFRHPSSYKFKLDSLGLHFVRLHFSVVF

SRADLLTARFTVSATSGSNHHLKSFSPQNLTNTPRVEEFLLMMNSLEFEIRFVPDHSSLA

LINAIEVFSAPDDLEIPSASDKNLHTIYRLNVGGEKITPDNDTLGRTWLPDDDDFLYRKD

SARNINSTQTPNYVGGLSSATDSTAPDFVYKTAKAMNRSSNEQVGMLMNVTWSFKVKSNH

RHFIRIHFSDILSNLSNSDSDFYLFVNGYWRVDVKPSEQPRLASPFFKDVVNVSDGSGLL

NISIGTKEANKDAGFLNGLEMMEVLSKSGSDYSNRSSSRVHIITGCAVAAAAASALVFSL

LFMVFLKRRRSKKTKPEVEGTVWSPLPLHRGGSSDNRPISQYHNSPLRNLHLGLTIPFTD

ILSATNNFDEQLLIGKGGFGYVYKAILPDGTKAAIKRGKTGSGQGILEFQTEIQVLSRIR

HRHLVSLTGYCEENSEMILVYEFMEKGTLKEHLYGSNLPSLTWKQRLEICIGAARGLDYL

HSSGSEGAIIHRDVKSTNILLDEHNIAKVADFGLSKIHNQDESNISINIKGTFGYLDPEY

LQTHKLTEKSDVYAFGVVLLEVLFARPAIDPYLPHEEVNLSEWVMFCKSKGTIDEILDPS

LIGQIETNSLKKFMEIAEKCLKEYGDERPSMRDVIWDLEYVLQLQMMTNRREAHEEDSTA

INSGGSLVAPRLMVSDSFSTNSIFQNGDESKNRFGFTDSSETRVFSQLKISDAR

>Hv-CrRLK1L1(XP_044984910.1)

MPALAILARSMVQCKRVPMFLILFILSITRVVTTDAIGSKVERFVPQDNYLLSCGASAAVQVDDGRTFRSDPESVSFLSTPTDIKITAKASLASASPLSPLYLDARVFSDISTYSFFISQPGRHWIRLYFLPITDSQYNLTTATFSVSTDSMVLLHDFSFIASPPNPVFREYLVSAQGDNLKIIFTPKKNSIAFINAIEVVSAPPSLIPNTTTRMGPQDQFDISNNALQVVYRLNMGGALVTSFNDTLGRTWQPDAPFLKLEAAAEAAWVPPRTIKYPDDKTLTPLIAPPSIYSTAQQMASTNITNARFNITWVMVAETGFRYLIRLHFSDIVSKTLNGLYFNVYINGMMAVANLDLSSLTMGLAVAYYKDLIAESSSIINSTLVVQVGPNTIDSGEPNAILNGLEIMKISNEASSLDGLFSPKTSSEVSKTTLTGIAFALAATAAFAVVICYRRNRKPAWQRTNSFHSWFLPLNSSSSFMSSCSRLSRNRFGSTRTKSGFSSVFASSAYGLGRYFTFIEIQKATKNFEEKGVIGVGGFGKVYLGATEDGTQLAIKRGNPSSDQGMNEFLTEIQMLSKLRHRHLVSLIGCCDENNEMILVYEFMSNGPLRDHLYGDTNIKPLSWKQRLEVCIGAAKGLHYLHTGSAQGIIHRDVKTTNILLDENFVAKVADFGLSKDAPSLEQTHVSTAVKGSFGYLDPEYFRRQQLTDKSDVYSFGVVLFEVLCARPAINPALPRDQVNLAEWARTWHRKGELGKIIDPNIAGQIRPDSLDMFAEAAEKCLADYGVDRPTMGDVLWKLEFALQLQEKGDVVDGASDGIPMKSLEMSNVDSMEKSGNAIPSYVQGR

>Hv-CrRLK1L2(XP_044974727.1)

MVLPTLPVTLIFLTLLAFLSIAMAADSNSMASGLILLNCGASGQGDDDIGRTWDGDISSKFAPLLEGVAANAAYEDPSLPSMVPYMTARIFTSNYTYSFPVTAGRVFLRLYFYPIAYGNYVVSDAFFGVTAGNLVLLNGFSASQTAQATSSAYLVREYSVNVSSGSLDLTFAPSTHQTGSYAFVNGIEIVPTPDIFTTADTKFVSGNHTDLFKFTADTGFQTMYRINVGGPYISPKDDSGFYRSWINDAPYRYDDSGVTFSKDDNVTIRYTPTVPNYTAPVDVYASARSMGQNPHVNPNFHLTHKLNYNLTWILPVDAGFFYLLRFHFCEIEYPITKVNQRVFFININNHAAQQKVDVILWSGGIGRTAYRDYAIMATGSSMVDLWIALEADFSDQPEFTDVILNGLEVFKLQGYGTNNLAGLNPPLPQKPSGARKYKGDKLAAIWGTTGGFALILIALTITCVISRQKKVGKSSFKTDCRHLNRPTECRESTCDLVRRFSFAEIQLVTKDFDEAFIIGRGGFGNVYSGEIDGRTKVAIKRFNQKSQQGFHEFQTEIEMLCNFRHRHLVSLIGYCEEKNEMILVYDYMAHGTFREHLYNTGNPPLPWQQRLEICIGAARGLHYLHTGTEQGIIHRDVKTTNILLDDRLMAKVSDFGLSKASPDIDNSHMSTVVKGTFGYLDPEYFRLQRLTKKSDVYSFGVVLFETLCARPVINTELPYEQVSLRDWALSCRKKGVLEEIVDPCVKEEITPQCFRTFAEIAEKCVADRSIDRPSMGDVLWNLEVALQLQDSASYNSSCAEGASSLQTSGVHSGKPSTNSTISVSAQEAIFSDIVHPEGR

>Hv-CrRLK1L3(XP_044950518.1)

MASPTLPVTLTCLTLLALLSLAMAADNNNSTGLILVNCGASGQEDDDSGRTWGEDTGSKFAPSLKGVAANAQRQDSSLPSTVPFMTARIFTSNYTYSFPVTPGRMFLRLYFYPTDYPNFAASDSSFSVSVSTPNLVLLDGFNASQTVQAISSAFLVREFSVNVSSGSTLDLTFAPSAHQNGSYAFVNGIEIVPTPDIFTAPDTRTVGDDTSPYTFDTAMAVQTMYRLNVGGQAISPKGDSGFYRSWTNDAPYIAGGSGVTFSKDDNLTITYTSKVPKYTAPADVYGSARSMGTTAQVNLNYNLTWILPVDAGFSYLLRFHFCEIQYPITKINQRSFFIYINNQTAQEQMDVIVWSGGIGRTTYTDYVTMAVGAGQVDMWIALHPDLSSKPEYFDAILNGLEVFKLQNNGSPNNLSGLNPPLPQRPPDAIPSAPSAGGKSVGAIVGAAVGGFAVLLVACFGVCIICKRKNNKKKKISKEPGGKSEDGHWTPLTEYSGSRSTMSGNTATTGSTLPSNLCRHFTFAELQTATKNFDQAFLLGKGGFGNVYLGEVDSGTKVAIKRCNPMSEQGVHEFQTEIEMLSKLRHRHLVSLIGYCEDKSEMILVYDYMAHGTLREHLYSTKNPPLSWKKRLEICIGAARGLYYLHTGVKHTIIHRDVKTTNILLDDKWVAKVSDFGLSKTGPNMDATHVSTVVKGSFGYLDPEYFRRQQLSEKSDVYSFGVVLFEVLCARPALSPSLPKEQISLADWALRCQKQGVLGQIIDPMLQGRIAPQCFVKFTETAEKCVADRSVDRPSMGDVLWNLEFALQLQESDEDTSSLTDGMLSSSGASPLVMTRLQSDEPSTDASTTTTSTTTMSMTGRSIASVDSDGLTPSTVFSQLMRPGGR

>Hv-CrRLK1L4(XP_044975236.1)

MAAAARHRPARARGALWIVSVLLVCVAAAYTPEDNYLVSCGSSLDTPVGRRLFLADDGGSGAVTLTSPRSAAVKASPDLVSGFRDAALYQNARVFSAPSSYSFAIKRRGRHFLRLHFFPFVYRSYDLAAAARAFKVSTQDAVLLEDGIPAPEPGNASSSPQPARVEFLLDVERDTLVVSFVPLIDGGIAFVNAVEVVSVPDNLVTDAAATTADSSSGRPELNPAALPLQTAYRVNVGGPVVAPDDDALWREWTTDQPLSDPRVDAVTREVRYNRTLNRLPGQATVTDAPDIVYATARELVITNISMDGQKQMAWQFDVDTRSSYFIRFHFCDIVGNASHQLRMNAYVDDATVKQDLDLAAIGNGALAFPYYTDFVLSASAASGKLAVHVGPRENKIVSPAAILNGIEIMKMHLSAGSVVVVEPAAKAAKSRLAVLLGSVCGAFAFVTIAVALAIVLRKKKNEKEEKEGDKEQPTPTQSQSSTPWMPLLGRFSVRSAIASGSSSFTTAGNTPGASPRAAAAAAAAVMPSYRFPLAMLQDATRNFDDSLVIGEGGFGKVYGAVLQDGTKVAVKRASPESRQGAREFRTEIELLSGLRHRHLVSLVGYCDEREEMILLYEYMEHGSLRSRLYGRSASPLSWAQRLEACAGAARGLLYLHTAVDKPVIHRDVKSSNILLDGDLTGKVADFGLSKAGPVLDETHVSTAVKGSFGYVDPEYCRTRQLTAKSDVYSLGVVLLEAVCARPVVDPRLPKPMSNLVEWGLHWQGRGELEKIVDRRIAAAARPAALRKYGETVARCLAERAADRPTMEDVVWNLQFVMRLQEGDGLDFSDVSSLNMVTELRPPRRQRNSVDCDGLDLSDVNSLKLVTEQTQPQTGSVEGDGVADDDFTDASMRGTFWQMVNVRSR

>Hv-CrRLK1L5(XP_044981625.1)

MRGPPRCALLLLLAAAALVPAARAQGAAAPAPSAGAAFVPRDDILLDCGATGKGNDTDGRQWDGDAGSKYAPPNLASASAGAQDPSVPQVPYLTARVSAAPFTYSFPLGPGRKFLRLHFYPANYSNRDAADAFFSVSVPAAKVTLLSNFSAYQTITALNFAYLVREFSVNVTGQNLDLTFTPEKGRPNAYAFVNGIEVVSSPDLFDLATPLFVTGDGNNQPFPMDPAAALQTMYRLNVGGQAISPSKDSGGARSWDDDTPYIYGAGAGVSYQNDPNVTITYPDNVPGYVAPSDVYATARSMGPDKGVNLAYNLTWILQVDAGYQYLVRLHFCEIQSPFTKPNQRVFSIYLNNQTAMKGADVILWADPNGIGTPVYKDYVVSTVGSGTMDFWVALHPDVETKPQYYDAILNGMEVFKLQLTNGSLVGLNPLPSPDPPVNSGSGNKKSIVVPIVGGVVGGLAVLALGYCCFICKRRRKAAKASGMSDGHSGWLPLSLYGHSHTSSSAKSHATGSYASSLPSNLCRHFSFAEIKAATKNFDESRILGVGGFGKVYHGEIDGGTTKVAIKRGNPLSEQGIHEFQTEIEMLSKLRHRHLVSLIGYCEEKNEMILVYDYMAHGTLREHLYKTQNAPLSWRQRLEICIGAARGLHYLHTGAKHTIIHRDVKTTNILLDEKWVAKVSDFGLSKTGPSMDHTHVSTVVKGSFGYLDPEYFRRQQLTEKSDVYSFGVVLFEVLCARPALNPTLAKEEVSLAEWALHCQKKGILDQIVDPYLKGKIVPQCFKKFAETAEKCVADNGIERPSMGDVLWNLEFALQMQESAEESGSIGCGMSDEGTPLVMVGKKDPNDPSIDSSTTTTTTTSLSMGDQSVASIDSDGLTPSAVFSQIMNPKGR

>Hv-CrRLK1L6(XP_044983157.1)

MAVHVLLPLLLLLLVPTVLPYIALAASSPDFTIFLACGAGANISFPSDNPARTFVPDAGYFSPARAPAVSAGASSSAASPLYAAARAGSSDFSYRLTYPDTAGASSFLVLRLHFFPFVPASSSTSLSSARFTVSVLDAYALLRNFSPPADGIVKEFFLPRGRSGDFTVRFSPDAGSSAFVNAVELFPAPPELLWNGSTSVVPVGVLEGTGLPQWQLAALETVYRLNVGGPMVTRENDTLWRTWLPDGPYLFGAPGQSVVNNTSSPIIYNPPTTREVAPDVVYRTQRAANVTDLMRATTPGLNFNVTWTFPAEAGSRYLVRLHFCDYEVVSSVVGVGIVFNVYVAQALATLDLSPEDRARQPNEAFYVDYAAMAPRAGNLTVSIGWSPKSSGGGILNGLEIMKLQSANLSSPRPHGLTKKTIIVIVLATVLGAAVLACAVLCFFVVLRRKRRQVAPPASTEDKESTQLPWSPYTQEGVSGWADESTNRSSEGTTARMQRVSTKLHISLAEVKAATDNFHDRNLIGVGGFGNVYKGALADGTPVAVKRAMRASKQGLPEFQTEIVVLSGIRHRHLVALIGYCNEQAEMILVYEYMEKGTLRSHLYGSDEPTLSWKQRLEICIGAARGLHYLHCGYSENIIHRDVKSTNILLGTDDHGGGSASGGAAIIAKVADFGLSRIGPSLGETHVSTAVKGSFGYLDPEYFKTQQLTDRSDVYSFGVVLFEVLCARPVIDQSLDRDQINIAEWAVRMHGEGKLDKIADARIAGEVNENSLRKFAETAEKCLAEYGADRPSMGDVLWNLEYCLQLQETHVNRDAFEDSGAVATQLPADVVVPRWVPSSTSMLMMDDADETGLSMTEIADSQVFSQLNARGEGR

>Hv-CrRLK1L7(XP_044970778.1)

MAVHGRLLVLLLATVLPRAALAALSPGFQFQFFLACGANYSVSFPSNDFPTNTFVPDDAYLSPASAPAVSARFTPYSRPALHAAARADISAFSYRFPSPASPYTPPFVVLRLHFFPFFQATSSQYVINIFSARFNVSVSGHEYALMSSFSPPVNGAVKEFFVPRDLSGGDFHVTFTPDAGSSAFVNAIELFSAPLEMLWCGSVTPVGAVVKDDMDLWQRQPLETVYRLNVGGPEVTIENDTLWRTWLPDGPYLYDARGKSVVSNTSSPIIYDTSNGYTREVAPDVVYQTQRMANVTDWLLATTPGLNFNLTWTFPAVKGSRYLVRLHFCDYEVVSSVVGVSIVFNVYIAQAIGTPDLMPNDRATVSNKAFYMDYAAKAPSTGNLTVSIGLVLKSNGGGILNGLEIMRLPPVDLSSRRSNGQTKRTVLITVAVVLGAAVLACVALCLFGVPYTKYSASGWAEQWTNRWFGEGETSGMESVSRKLHIPLAKIKAATDSFHERNLIGVGGFGNVYKGVLSDGTPVAVKRAMRASQQGLPKFQTEIVVLSGIRHQHLVSLIGYCNEQAEMILVYEYMEKGTLRSHLYGSDEPALSWKQRLEICIGAARGLHYLHRGYAENIIHRDVKSTNILLGSDGGSTGGVITKVADFGLSRIGPSFGETHVSTAVKGSFGYLDPGYFKTQQLTDRSDVYSFGVVLLEVLCARPVIDQSLDHSMINIAEWAMRMRREGRLDKMADPRIAGEVDEESLLKFVETAEKCLADCWVDRPSMGDVLWNLEYCMQLQEMNVIGDEHDNMVPSSTSLLLDDTGLSMTNVADNKVSPARARDEAR

>Hv-CrRLK1L8(XP_044981971.1)

MPPLVDMLLLASVLLGAASSAVAGADNYLVVCGTSASATVAPGRTFAGDARLPAKSLAAPQSVEANTSLTAAVPSGESELYRSARVFTAPASYTFAVKQPGRHFVRLHFFPFAYQSYDMAADAAFNVSVQGAVFLNGYTNKNGTAELREFSVNVTGATLVIAFTPTGKLAFVNAIEVVPLPDELIADTASMVDRAVQYTGLSTQALETIHRINMGIPKITPGNDTLGRTWLPDQGFQLNANLAQHKDAKPLTIKYDEKSALSSAFTAPAEVYATATKLSTAGEVSTINVQFNISWRFDAPAGSDYLLRFHFCDIVSKAAIGMAFNVYVGGSVVLKNYEISRDTFNRLSIPVYKDFLLGAEDAKGTITVSIGSSTDDNALPDGFLNGLEIMRLVGSAGAGAAAASPRSSKVKIGIIAGSAVCGATLIMVLGFIAFRTLRRTEPEKKPSDTWSPFSASALGSRSRSRSFSKSSGNTVMLGQNGAGAGYRIPFAALQEATGGFDEGMVIGEGGFGKVYKGTMRDETLVAVKRGNRRTQQGLHEFHTEIEMLSRLRHRHLVSLIGYCDERGEMILVYEYMAMGTLRSHLYGAGLPPLSWEQRLEACIGAARGLHYLHTGSAKAIIHRDVKSANILLDDTFMAKVADFGLSKNGPELDKTHVSTKVKGSFGYLDPEYFRRQMLTEKSDVYSFGVVLLEVLCARTVIDPTLPREMVNLAEWATPCLRNGQLDQIVDQRIAGTIRPGSLKKLADTADKCLAEYGVERPTMGDVLWCLEFALQLQMGSSDGSETDTMLPPPVPGKTPLVQRSLSTATVPTDDAAMTTNLGDLEGMSMSGVFSKMIKSDEVR

>Hv-CrRLK1L9(XP_044973269.1)

MRGGPRCALLLLLLAACAALVPAAWAQGGGDAPAPAAPFVPRDDILLDCGATGKGDDTDGRQWAGDAGSKYAPPNLASAAAGAQDPSVPQVPYLTARVSAAPFTYSFPLGPGRKFLRLHFYPANYSDRNAADAFFSVSVPAAKVTLLSNFSAYQTTTALNFAYIVREFSVNVTGPTLDLTFTPEKARRNAYAFINGIEVVSSPDLFDLATPFFVTGDANNQPFPMDPGAALQTMYRLNVGGQAISPSKDSGGARSWDDDTPYIYGAGAGVSYPNDPNITITYPDNVPGYVAPLDVYATARSMGPDKDVNLAYNLTWIVQVDAGFTYLVRLHFCEIQSPITLPNQRVFNIYLNNQTAQTGADVIQWVDPKSIGTPVYKDYVVSTVGSGIMDFWVALHPDTGNKPQYYDAILNGLEVFKLQLSNGSLAGPNPVPSADPPVHTGQGKKSSLVGPIAGGVIGGLALLALGYCCLICKRRRKTAKDTGMSDGHSGWLPLSLYGNSHTSSSDKSHATGSIASSLPSNLCRHFSFAEIKAATKNFDESRILGVGGFGKVYQGEIDGGTTKVAIKRGNPLSEQGIHEFQTEIEMLSKLRHRHLVSLIGYCEDKNEMILVYDHMAHGTLREHLYKTQNAPLSWRQRLEICIGAARGLHYLHTGAKHTIIHRDVKTTNILLDEKWVAKVSDFGLSKTGPSMDHTHVSTVVKGSFGYLDPEYFRRQQLTEKSDVYSFGVVLFEVLCARPALNPTLAKEEVSLAEWALHCQKKGILDQIVDPYLKGKIVPQCFKKFAETAEKCVADNGIERPSMGDVLWNLEFALQMQESAEESGSFGCGMSDEGTPLVMPGKKDPNDPSIDSSTTTTTTTSISMGDQSVASIDSDGLTPSAVFSQIMNPKGR

>Hv-CrRLK1L10(XP_044967870.1)

MLQMRLLVLAAVSIVFANLQFLKAHGRELFLSCGSNATADADGRRWIGDMAPDLNFTLSSPGIAALLAGGSNGSEIMAPVYRSARFFTTTSWYDFSLLPGNYCVRLHFFPSTFRNFSASSSVFDVVANEFKLVSKFNVTEEIVWRNSVSNSAATALVKEYFLAVNTSRLQIEFDPRPGSFAFVNAIEVVLAPDNSFNDTVNKVGGVDVHIPPELSGRAVETMYRLNIGGPALASSHDQHLHRPWYTDEAFMFSANAALTVSNTSAIKYVSSNDSSIAPIDVYETARIMGNNMVMDKRFNVTWRFFVHPNFDYLVRLHFCELVYDKPSQRIFKIYINNKTAAENYDVYDRAGGINKAYHEDYFDSLPQQVDSLWLQLGPDSMTSASGTDALLNGLEIFKLSRSGSLDYVLGHIDVGNKRGRSKGRSRIGLWEEVGIGSAAFVVLASVALFSWCYVRRKRKAVDEEVPAGWHPLVLHEAMKSTTDARASKKSPLARNSSSIGHRMGRRFSIADIRAATKNFDESLVIGSGGFGKVYKGEVDDGITVAIKRANPLCGQGLKEFETEIEMLSKLRHRHLVAMIGYCEEQKEMILVYEYMAKGTLRSHLYGSGLPPLTWKQRIDACIGAARGLHYLHTGADRGIIHRDVKTTNILLDKNFVAKIADFGLSKTGPTLDQTHVSTAIRGSFGYLDPEYFRRQQLTQKSDVYSFGVVLFEVACARPVIDPSVPKDQINLAEWAMRWQRQRSLEAIADPRLDGDYSPESLKKFGDIAEKCLADDGRTRPSMGEVLWHLEYVLQLHEAYKRNVDCESFGSSELGFADMSFSMPHIREGEEEHHPKKSDIREDSAP

>Hv-CrRLK1L11(XP_044966102.1)

MGTTSEQKIALLLLGTIWVLLGTCNAEFTPADNYLINCGSTVDVHVPGQGFFRADNSGSTILKSDHNVAANTLPDAVISSDNPVLYQTARIFSVPSSYSFNMKSRGRHFVRLHFFGFRYQSYDLAVAKFKVSTQDVVLLDNFTPPSNSSPLVREYSLNITEDKLILTFVPLGNSTSFINAIEVISVPDDLIRDSAQTVNPSGQYLGLTTQSFQTFYRINVGGREVTAANDTLSRSWDTDQNFFLNSTTTELFAYQAKLNYQKGAATKEDAPDSVYNTARRFAVQNRTSLVSNMTWQFDVDGSSSYLIRFHFCDIVSKAAYSLYFDIYVDGRLALENVDLSERVLGTLAVPYYMEFVLKSSDPSGKLSVGIGPSSLNNVAPDGILNGLEIMKMDISTGTVYVVWPPGTPNRKLAIILGTVLGGVGAVSIAIILCFVLRRKKKEKKPRRAPTSRPSSSWSPLTLNGLSFLSTGTRTTSRTTLTSGTNSDASYRIPFALLQVATKHFDEQMVVGVGGFGKVYKAVLQDSTKVAVKRGNQKSHQGLREFRTEIELLSGLRHRHLVSLIGYCDEQNEMILVYEYMEKGTLKSHLYGSDMPPLSWKKRVEICIGAARGLHYLHTGFAKSIIHRDVKSANILLDENLMAKVSDFGLSKTGPELDQTHVSTAVKGSFGYLDPEYYRRQKLTDKSDVYSFGVVLLEVICARPVIDPTLPRDMINLAEWAIKWQKRGELGQIVDQRIAGTIRPESLRKYGETVEKCLADYGVDRPTMGDVLWNLEFVLQLQEAGPDVSNVDSMNQISELPPDTRRMGSLEIGTADESPTNMDYSQMSTNDAFSQLMNTEGR

>Hv-CrRLK1L12(XP_044983735.1)

MAAARGRGVLLAVLLAVALALVSVGADIYKPTDSILVDCGSDKEGQDEDGRKWTSDKDSKFLPDGGKSSITATADINDPSLPSSVPYMTARVFPKETAYTFPVSDADRHWVRLHFYPASYHDIPADHFFFSVTTSTGVTLLRNFSVYVTAKALTQGYIVREFSLPPSTTGSLSLKFTPTAMNNASYAFVNGIEILSMPNIFADPAQLVGLGDQTLDASAGNMQTMYRLTAGGSYIPATKDSGLSREWFPDTPYIYGAATGVTYEANDTVPIKYPSPAAEYLAPQSVYDTSRHMGRDGNVNRVNNLTWAFEVDGNFTYLVRLHFCSLMEDKINQVVFSILVNNKTATTTGSADIIAWAKEKNPDSPGKGVPVIRDYAVFTAAAPAGADSVLWVTLRPDLTTTPQFVNAFLNGLEVFKTSDASSNLAGPNPDISKMLAEAEAEADDVEGEFRERASNVGALIGGAAGGAAAFGLVAGLCFVAYQSKKRKELSNSRSHSSSGWLPVYGGGGNSQTSVSKSSGGRSAVTLNPNITAMCRHFSLQEIKSATKGFDESLVIGVGGFGKVYRGVVDGDTKVAIKRSNPSSEQGVLEFQTEIEMLSKLRHKHLVSLIGCCEDNGEMILVYDYMGHGTLREHLYKSGKPPLLWRQRLEILIGAARGLHYLHTGAKYTIIHRDVKTTNILVDDKWVAKVSDFGLSKTGPTVQNQTHVSTMVKGSFGYLDPEYFRRQKLTEKSDVYSFGVVLFEVLCARPALNPSLPREQVSLADHALSCQRRGTLEEIIDPVLEGKVAPDCLKKFAETAEKCLSDQGVDRPSMGDVLWNLEFALQMQDTFDNGGKPPEVDDYSSSFTITPPSMEESLAANAAALSLISEDMDEEDIANSVVFSQLTHPTGR

>Hv-CrRLK1L13(XP_044958511.1)

MVRRGALPLALLAALATLTAVSGQGKPVTDNGAGSSQAKFTPKDAFYIDCGGTAAADTKDGKSFKTDAEANNLLSAKDAIKVADDKADVPSHLYRTARVFKAEAIYNFPLTAPGWHFIRLYFFPIKSGEADLAAATFDVTTAANVLLHSFTAEAKAVMKEYVINATENKLELKFTPASGSAAFINAIEVVNAPDQLLNDKALTVSPLAEISGLSEAAYQVVCRLNVGGPVIGPVNDTLGRQWDEDGPYLNPKEAGVEVSVSPNVIKYPETYPVSKLVAPTAVYTTARHMANSGVANQNFNVSWKVDVDASFDYLVRLFFADIISDTMNDLYFNVYINERKAISGLDLSTITGDMAAPFYKDFVVNSSVDTDGHIIFAVGPMGQDTGRIDALLNGAEVLRISNSVGSLDGEFGVDGRMVDDGSGTRKVVAAVGFAMMFGAFAGLGCMVVKWHRRPQDWERRNSFSSWLLPIHTGQSFTNGKGSKSGYTVSSTAGLGHFFSFAEMQEATKNFDDSAIIGVGGFGNVYVGEINDPDEEGSRIKVAIKRGNPSSEQGINEFNTEIQMLSKLRHRHLVSLIGYCDENEEMILVYEFMQHGPFRDHIYGGPEGLPTLSWKQRLEICIGAARGLHYLHTGTAHGIIHRDVKTTNILLDEKFVAKVADFGLSKDGPGMNQLHVSTAVKGSFGYLDPEYFRCQQLTDKSDVYSFGVVLLETLCARAPIDPQLPREQVSLAEWGMQWKRKGLIEKIMDPNLAGKVNPESLAKFAETAEKCLCEFGSDRLSMGDVLWNLEYALQLQEANPPEGATDADDADASIVSSSSGVSTVPDASTTSANELFAQLADMKGR

>Hv-CrRLK1L14(KAE8784173.1)

MPAAGRSGGPGQVNIMTGRRKLHVVTLAILCFWSSAGVCKAQTVDFKPADSYLVDCGSAKGTTVLGRNFAADGASPVTVSTSQDILAGTSANGVSSFDNPVLYQTARIFTSPSSYTFPIQKQGRHFVRLYFYPFIYQTYDLSTAKFTVSTQDVLLLSDFQQPDKTAPLFKEYSLNITRDQLVISFKPSNGIAFINAIEVVSVPDDLIADVANMVNPVQQYSGLTTQSLETVYRVNMGGPKVFPSNDTLSRTWQKDQKYILNPSVTKTAQYGKAINYRNGGATPLTAPDIVYSTATELAASNTSNALFNMTWQFDVDAGFSYLIRFHFCDIVSKALNQLYFNAYVGGFFAQHDLDLSEQSVNQLATAIYVDVVLSSNDASSKLSISIGPSTLNNALPDGILNGLEIMKMGSGSGSAFTVGNNGSNKKLPIIIGSVLGVVGLLIIILVVVLLCRRKKTDDKQHSKTWMPFSINGLTSLSTGSRTSYGTTLTSGLNGSYGYRFAFNVLQEATNNFDENWVIGVGGFGKVYKGALRDDTKVAVKRGNPKSQQGLNEFRTEIELLSRLRHRHLVSLIGYCDERNEMILVYEYMENGTVKSHLYGSDNPSLNWKQRLEICIGAARGLHYLHTGSAKAIIHRDVKSANILLDENLLAKVADFGLSKTGPELDQTHVSTAVKGSFGYLDPEYFRRQQLTEKSDVYSFGVVMLEVLCARPVIDPSLPREMVNLAEWGMKWQKRGELHQIVDQKLSGAIRPDSLRKFGETVEKCLADYGVERPSMGDVLWNLEYVLQLQDADSSTVSDVNSMNRIVDLSSQVQHVGAMESISVTMAEDGASHEPDHDLSDVSMSRVFSQLIKAEGR

>Hv-CrRLK1L15(XP_044966011.1)

MESSAKFLLMLVAFLTAGNARAQPQPVLINCGSDSATNADARIWIGDSSPSSNFTLSFPGAVATAAPGGQDPYGDLYKTARLFNASSSYRLAVAPGSYFLRLHFSQLFANPGAQEPIFSVAANGLKLLSKFSVHGEISWRDSQINSTSSVIVKEYLLNVTSGKLGIEFTPDEGSFAFINAMEVLPVSGTPIFDSVNKVGAHGLKGPLSLDRGGIETMYRLCVGCIDVLPRKEDPGLWRKWDSDEHFIFSLNAARSIFNSSNISYVSADDPTLAPLRLYQAARVPTESSVLGKKFNVSWSFNIDPGFDYLVRLHFCELEYDKAEQRKFKIYINNKTAAEGYDVFARAGGKNKAFYEDFLDAASPQMDTLWVQLGAESSAGSAAADALLNGMEIFKVSRDGNLAHPTVRIGGISGGVSKPKRSPKWVLIGAASGLIFFIAIIGAVYFCFNLQRKKNSSANKAKDNLHGATHTRSPTLRTAGAFGSNRMGRRFTIAEIRTATLNFDESLVIGVGGFGKVYKGKMEDGTRVAIKRGHTESHQGQGVKEFETEIEMLSRLRHRHLVPLIGYCDEQNEMVLVYEHMANGTLRSHLYGSDLPALTWKQRLEICIGAARGLHYLHTGLDRGIIHRDVKTTNILLDNNLVAKMADFGISKDGPALDHTHVSTAVKGSFGYLDPEYYRRQQLTPSSDVYSFGVVLFEVLCARSVINPTLPRDQINLADWALNRQRHKLLETIIDLRLEGNYTLESIKKFSEIAEKCLADEGVNRPSMGEVLWHLESALQLQQGHPQSTNGDDCSDSQAQPSDVPIRIKEAEQSTRPGSHDSDGQVVDVKIEVP

>Os-CrRLK1L1

MLSLQKPSKTPFVPASSALFNPESIFLPSHERSPSSCYRPMLLIPTLAKMARSMLGWKRV

PLFSILLILSITNIATTYAIASQADRFVPRDNYLLSCGAPAAVQLDDGRTFRSDPDSASF

LSTPVDIKITAKNSLASGAPSSQLYLTSRVFSDISTYSFFISQPGHHWIRLHFLPIPDDH

YNLTTATFSVSTDDMVLLHDFSFIATPPNPVLREYIVATQGDTLKIIFTPKKDSIAFINA

IEVVSAPPSLIPNTTTGMAPQGQLDISNNALQVVYRLNMGGPLVTAFNDTLGRIWLPDAP

FLKLQAAANAAWVPPRTIKYPDDKTNTPLIAPANIYSTAQQMASTNTSDARFNITWEMVT

EPGFSYFVRLHFCDIVSKALNSLYFNVYINGMMGVLNLDLSSLTVGLAVPYYRDFIIDSS

SIINSTLIVQIGPGTTDTSNPNAILNGLEIMKISNQENSLDGLFSPKRSSQLGKKTMTGI

GLAMAVMAAALAVVMCCRRRHRPGWQKTNSFQSWFLPLNSTQSSFMSTCSRLSSRNRFGS

TRTKSGFSSIFASSAYGLGRYFTFVEIQKATKNFEEKAVIGVGGFGKVYLGVLEDGTKLA

IKRGNPSSDQGMNEFLTEIQMLSKLRHRHLVSLIGCCDENNEMILVYEFMSNGPLRDHLY

GGTDIKPLSWKQRLEISIGAAKGLHYLHTGAAQGIIHRDVKTTNILLDENFVAKVADFGL

SKAAPSLEQTHVSTAVKGSFGYLDPEYFRRQQLTEKSDVYSFGVVLFEVLCARPAINPTL

PRDQVNLAEWARTWHRKGELNKIIDPHISGQIRPDSLEIFAEAAEKCLADYGVDRPSMGD

VLWKLEFALQLQEKGDIVDGTSNQFPMKSLEVTSGDSMEKSGNVVPSYVQGR

>Os-CrRLK1L2

MMHPSLLATTQWVTLSTLLSIAIAADNYFSSSSPIFLNCGASAMQLDSNNRSWDGDTSST

FAPSVKGLAARASYQDPSLPSLVPYMTSRIFISNYTYSFPVIPGRMFVRLHFYPVAYGNY

ASRDAYFGVTTNNLTLLDNFNASQTALAAKYAYILREFSLNVTSGSLDLTFFPSTQNGSY

AFVNGIEIVPTPDIFTTLSPIPPTNGNPDPSDIDSMISFQTMYRLNVGGMTISPQGDSMF

YRSWENDSPYIYGSAFGVTFSKDSNVTITYPSTMPNYIAPADVYGTARSMGPIAQINLHY

SLTWILPVDAGFYYLLRFHFCEIEYPITKVNQRSFFIYINNQTVQEQMDVIVWSGGIGIT

TYTDYVIVTVGSGQMDLWVALHPDLSSGPEYYDAILNGLEVFKLQDIGKKSLAGLNPPLP

PQPKSDVNPKGVSGGGKSKGAVPASIRGAMGSTATMLIACFSVCIICRLKKVAKHSFMTD

KKCMTYRTEFYHSPSNLCRNFTFDEIQVATRNFDESLLLGRGGFGDVYRGEIDNNGENVA

IKRSNPLSVQGVHEFQTEIELLSKLRYCHLVSLIGYCKEKNEMILVYEYMAQGTLREHLY

NSNKPSLPWKQRLKICIGAARGLHYLHMGANQTIIHRDVKTANILLDDKWVAKVSDFGLS

KANPDIDSTHVSTVVKGTFGYLDPEYYRRKQLTQKSDVYSFGVVLFEILCARPAVNIELP

EEQASLRDWALSCQKKGMLGKIIDPHLHGEISPPCLRMFADCAKQCVADRSIDRPLMSDV

LWSLEAALKLQENAENNKKFSEATTSSKRTPDLITIMGTDKPSTYSTMSITGQKIIFSDM

MHPQGR

>Os-CrRLK1L3

MMHPSLLATIQWLTLSALLSIAMAADNNSTASAPIFLNCGASGVQPDSYNRSWDGDASSK

FAPSVKGNVARASYQDPSLPSPVPYMTARFFTSNYTYSFPVSPGRMFVRLHFYPTNYNGN

LDSANAYFGVTTNNLILLDNFNASQTALATSSAYFFREFSVNVTSSSLKLTFAPSTRNGS

YAFVNGIEIVPTPDIFTTPTPTSANGGDNVQYGIDPVMGLQTMYRLNVGGQPISPQGDSG

FYRSWDNDSPYIYGAAYGVTFSKDGNVTIKYPNTEPNYTAPVAVYATARSMGPTAQINLN

YNLTWILPVDAGFTYLLRFHFCEIQYPITKVNQRSFFIYINNQTAQNQMDVIVWSGGIGR

TTYTNYVVTTVGSGQTDLWVALHPDLSSKPEYFDAILNGLEVFKLQDLGRNNLAGLNPPL

PPKPGVNPNGGSSRGKSKSVAPAAIGGAVGGLAVLLIACVGLCIICRRKKKVAKDTGKSD

EGRWTPLTDFTKSQSATSGKTTNTGSHSMLPANLCRHFSFAEIQAATNNFDKSFLLGKGG

FGNVYLGEIDSGTRVAIKRGNPLSEQGVHEFQNEIEMLSKLRHRHLVSLIGYCEDRNEMI

LVYDYMAHGTLREHLYNTKNPPLSWKQRLEICIGAARGLYYLHTGAKQTIIHRDVKTTNI

LLDDKWVAKVSDFGLSKAGPNVDNTHVSTVVKGSFGYLDPEYFRRQQLTEKSDVYSFGVV

LFEVLCARNALSPSLPKEQVSLADWALRCQKKGVLGEIIDPLLKGKIAPQCFLKFAETAE

KCVADRSVDRPSMGDVLWNLEFALQLQESTEDSSSLTEGTSASTSPLVVARLHSDEPSTD

VTTTTTTTTSLSITDRSIASVESDGLTPSNIFSQLMTPDGR

>Os-CrRLK1L4

MIKLRSALGVLEILSVLCISLVAAYTPVDNYLISCGSSVDTPVGQRLFVADDSGTVVLTS

PASDAVKASPSAVSGLRDDAAMYQSARVFKAPSSYSFRIRDPGRHFVRLHFFPFVYLGYD

LATASFKVSTQDAVLLDGFAPAAAARGNASTTTTTATAAAVCEEFLLDVARDTLVVTFVP

LAGRLAFVNAIEVVSVPDDLIGAADSSLSTSDSTGQQLNPAVMPLQTVYRVNVGGQAVAP

DSDTLWREWTSDQQLLVGPAMTKGVSYNRTPNYLPGQATANDAPAIVYATGRELIIMTNS

TDDGMKQMAWQFDVGRSASYLIRFHFCDIVSSVPGRLHMNAYVDSSNAIQDLDLSAIGNG

TLAFPYYRDFVLAASTPSGKLAVYVGSTSQKITTPAAILNGLEIMRILTTAGNVAVVEPT

MPPGTKKKNNLAVVLGSVCGAFGFVSVAAALVIVLRRKEEKEELRTPTTSQPSTAWMPLL

GRISFRSAPPSAVGSRSPSFTIDTNANTPGGGATPGMAAAASSSPSYRFPFAALQDATGN

FDEGLVIGEGGFGKVYAAVLQDGTKVAVKRANPESRQGAREFRTEIEMLSGLRHRHLVSL

IGYCDEQDEMILLYEYMEHGSLRSRLYGGGAATATATALSWAQRLEACAGAARGLLYLHT

ATAKPVIHRDVKSSNILLDDGLTAKVADFGLSKAGPDMDETHVSTAVKGSFGYVDPEYVR

TRKLTAKSDVYSFGVVLLEALCARPVVDPRLPKPMVNLVEWGLHWQRRDELEKIVDRRIA

GTVRPAALRKYGETVARCLADRGADRPAMEDVVWSLQFVARLQEVDGLDASDVSSLNMVH

QLMPPTSLHARQRSAGESETGRTDADEDSSVVDDDYTDASMRGIFWQMVNVRGR

>Os-CrRLK1L5

MGSSRFVLLLLLLLAVAACVARGQGGGNSSSAAAPAPAAGAGPFVPRDDILLDCGATGKGNDTDGRVWSGDAGSKYAPASLGSASAAGQDPSVPQVPYLTARVSAAPFTYSFPLGAGRKFLRLHFYPANYSSRDAADARFSVSVPAANVTLLSNFSAYQTATALNFAYIVREFSVNVTTPTMELTFTPEKGHPNAYAFVNGIEVVSSPDLFDISTPNLVTGDGNNQPFPIDAGTALQTMYRLNVGGQAISPSKDTGGYRSWDDDSPYVFGAAFGVSYPKDDNVTIAYPSNVPEYVAPVDVYATARSMGPDKNVNLAYNLTWIMQVDAGFTYLVRLHFCEIQYPITMINQRVFNIYINNQTAFQGADVIAWTNNNGIGSPVYQDFVVTTVGSGAMDLWVALYPDVQAKPQYYDAILNGLEVFKLPLSNGSLAGLNPVPTVEPSLDGGAVKKSSVGPIVGGVIGGLVVLALGYCCFMICKRRSRVGKDTGMSDGHSGWLPLSLYGNSHSSGSAKSHTTGSYASSLPSNLCRHFSFAEIKAATNNFDESLLLGVGGFGKVYRGEIDGGVTKVAIKRGNPLSEQGVHEFQTEIEMLSKLRHRHLVSLIGYCEEKNEMILVYDYMAHGTLREHLYKTKNAPLTWRQRLEICIGAARGLHYLHTGAKHTIIHRDVKTTNILLDEKWVAKVSDFGLSKTGPSMDHTHVSTVVKGSFGYLDPEYFRRQQLTEKSDVYSFGVVLFEVLCARPALNPTLAKEEVSLAEWALHCQKKGILDQIVDPHLKGKIAPQCFKKFAETAEKCVSDEGIDRPSMGDVLWNLEFALQMQESAEDSGSIGCGMSDEGTPLVMPGKKDPNDPSIESSTTTTTTTSISMGDQSVASIDSDGLTPSAVFSQIMNPKGR

>Os-CrRLK1L6

MLCFEPNKSVVSGRGGIGKCDDPISVHSIVHSTIPVFHHALPSGRCPIFLRVHFPVPDST

HHSTASHSSFPLPSNTTTRTVRTLVRFRLILSWTRQWRHDHASTTASMATVIVILLLLPL

LPSTALAAFPYFLACGAASNVSFPGDSPARTFVPDAPFLSSAGRVPAVTSTGSNTIPPLY

AAARAAGSGFSYSFADPDTATVNVSRVLRLHFFPFTSSSSVNLSSASFSVSVRDAYTLLS

SFSPPRDGVVKEYFVPGDGSGEFRVKFTPDAGSTAFVSAIELFPAPPELLWRRPVKPVGA

LVDSVDVNAWPQQALETVYRLNVGGSKVTAANDTLWRTWLPDDPYFSSPRGLSQVNSTST

PIIYGTSIGYTREVAPDSVYKTQRAMNMASQQLFLTPGPFNLTWTFALPPPAPGSDSDYL

VRLHWCDYSLVSSVVATGIVFDVYVAQRLASKDLDRNAADAAEQPNEAFYLDYAATAPTT

GNLTISIGKSDKSDAGGMLNGLEIMKLRRADNLNSAGSHGRRKKILIGTLSAALGVAVLA

CALLCLLAVLRRRRQAPTPAPEEKESTQLPWSQHTQDGSSWVDMSNASGAGMTGGLHRMS

MQLNISLADITAATENFNERNLIGVGGFGNVYSGVLRDGTRVAVKRAMRASKQGLPEFQT

EIEVLSRIRHRHLVSLIGYCNEQSEMILVYEYMEKGTLRSHLYGSEEPPLSWKQRLEICI

GAARGLHYLHTGYSENIIHRDVKSTNILLGDAFIAKVADFGLSRIGPSFGETHVSTAVKG

SFGYLDPEYFKTQQLTDRSDVYSFGVVLFEVLCARTVIDQSLERDEINLAEWAVSLQQKG

ELAKITDPRIAGQVNGNSLRKFAETAEKCLADYGLDRPSMGDVLWNLEYCLQLQETHVNR

DAFEDSGAVATQFPADVVVPRWVPSSTSFLMDDSVTDSGIANSKAFSQLSSGDGR

>Os-CrRLK1L7

MATVLEMLVQLAVVVTVLCAAVRAYTPADSYLFLCGTSGNATVDGRTFVGDAGLPASVLM

APQSTEANMPANQVTGAGDDSPALYQSARVFTAPANYAFSAKPGRHFVRLRFFPFRYQSY

DLAADAAFNVSVQGVVFVDGYTPKNGTAVVREFSVNITGRALVIAFTPTGKKVAFVNAIE

VVSHPDELIGDTAPMVNPRNQSQYTGLTAKALETVHRINMGEPKVTPNNDTLWRTWLPDW

TFLHESSFAAHNQVSPAMIKYQSGYATSLTAPSAVYTTVTELNTTAAMVGNTQAQLNLTW

KFNAPAVSDYLLRLHLCDIVSKATLGVVFNVYVGQWRVLQDYESSGDTFSLLATPLYKDF

VLAASDAAKGTITVSIGSSTATNALPGGFLNGLEIMRIVGSTGSIDGATSPRGSKIKTGI

IAGSAVGGAVLAIALGCVAVRMLRRKKKPVKQPSNTWVPFSASALGARSRTSFGRSSIVN

VVTLGQNGAGAGAGYRFPFAALQEATGGFEEEMVIGVGGFGKVYRGTLRDGTQVAVKRGN

RLSQQGLNEFRTEIELLSQLRHRHLVSLIGYCDERGEMILVYEYMAKGTLRSHLYGSDLP

PLPWKQRLEACIGAARGLHYLHTGSAKAIIHRDVKSANILLDDGFMAKVADFGLSKTGPE

LDKTHVSTAVKGSFGYLDPEYFRRQMLTEKSDVYSFGVVLLEVLCARAVIDPTLPREMVN

LAEWATRRLRDGELDRIVDQKIAGTIRPDSLKKFADTAEKCLAEYGVERPSMGDVLWCLE

YALQLQVASPDSSVTTLQRSSSISSVVTDATVSANLGDLDGMSMKRVFSKMLKSEEEGRR

KMH

>Os-CrRLK1L8

MMVSSRFVAVLLLVALAPAARGQGGGGGNSSAPAASPPGPFVPRDNILLDCGATGQANDT

DGRLWTGDTGSKYLPANLAAAAATAQDPSVPQVPYLTARFSAAPFTYSFPVGAGRKFLRL

HFYPANYSNRNAADALFSVSIPDPNITLLSNFSAYQTALALNFDYLVREFSVNVTASTLD

LTFTPEKGHPNAFAFVNGIEVVSSPDLFGSSNPMEVTGDGSGTPFPIDAGTAMQTMYRLN

VGGNAISPSKDTGGYRSWEDDTPYIPFASFGVSYANDTNVPINYPDSIPQYVAPADVYST

ARSMGPDNNVNLQYNLTWAMQVDAGYQYLVRLHFCEIQSGISKINQRTFDIYINNQTAFS

GADVIAWSTGLGIPVYKDFVVFPMGSGPMDLWVDLHPNVKNKPQYYNAILNGMEVFKLQL

TNGSLAGLNPVPSIVPTASGGNSGKKSSVGPIIGGVIGGLVVLALGCCCFFVICKRRQRA

GKDSGMSDGHSGWLPLSLYGNSHTSSSAKSHTTGSHASSLPSNLCRHFSFVEIKAATNNF

DESLLLGVGGFGKVYRGEIDGGATKVAIKRGNPLSEQGVHEFQTEIEMLSKLRHRHLVSL

IGYCEEKNEMILVYDYMAHGTLREHLYKTQNAPLSWRQRLDICIGAARGLHYLHTGAKHT

IIHRDVKTTNILLDEKWVAKVSDFGLSKTGPTMDHTHVSTVVKGSFGYLDPEYFRRQQLT

DKSDVYSFGVVLFEVLCARPALNPTLAKEEVSLAEWALHCQKKGILDQIVDPHLKGKIAP

QCFKKFAETAEKCVSDQGIDRPSMGDVLWNLEFALQMQESAEESGSLGCGMSDDSTPLVI

VGKKDPNDPSIESSTTTTTTTSISMGEQSVASIDSDGLTPSAVFSQIMNPKGR

>Os-CrRLK1L9

MRLLALAVASIVLANLHLLGVHGRDLLLSCGSNATVDAGGRRWIGDMAPGLNFTLSSPGI

AASQAGSSNGNEIFGLVYHSARFFSTASWYNFSVLPGNYCLRLHFFPYTFGNFSGNDSLF

DVTANDFKLVSKFNVSEEIVWRSTVSNSAINAVVKEYFLLVGSRGLQVEFDPSPGSFAFV

NAIEVMLTPDNLFNDTVNKVGSAGNGQLPLGLSNRGLETMYRLNVGGHALNSSSDQYLHR

PWYTDEAFMFSANAAQIVSNTSSVSYLSNNDSSISPIDVYETARIMSNNMVVDKRFNVSW

RFYVHPNFDYLVRLHFCELFYDKPNQRVFKIYINNKTAAEDYDVYVRAGGINKAYHEDYF

DNLPQQVDSLWLQLGPDSLTSASGTDPLLNGLEIFKLSRNGNLAYVLGHIDMGNQRGISK

DRNRKILWEEVGIGSASFVTLTSVVLFAWCYVRRKRKADEKEAPPGWHPLVLHEAMKSTT

DARAAGKSPLTRNSSSIGHRMGRRFSISEIRAATKNFDEALLIGTGGFGKVYKGEVDEGT

TVAIKRANPLCGQGLKEFETEIEMLSKLRHRHLVAMIGYCEEQKEMILVYEYMAKGTLRS

HLYGSDLPPLTWKQRVDACIGAARGLHYLHTGADRGIIHRDVKTTNILLDENFVAKIADF

GLSKTGPTLDQTHVSTAVKGSFGYLDPEYFRRQQLTQKSDVYSFGVVLFEVACGRPVIDP

TLPKDQINLAEWAMRWQRQRSLDAIVDPRLDGDFSSESLKKFGEIAEKCLADDGRSRPSM

GEVLWHLEYVLQLHEAYKRNNVDCESFGSSELGFADMSFSLPHIREGEEEHHSKPSSIRE

DPDT

>Os-CrRLK1L10

MDSTFRKLKLVLALVGIITWIIGTCNAKFTPADNYLVNCGSTVDATVGQRVFVADNSQSI

VLTTPQSQSIAARTTLNSVSGFDNAELFQTARIFTAPSSYSFKMRSSGRHFVRLYFFPFL

YQSYDLASSKFKVSTEDVVLIDNFPQPSNSISVVMEYSLNITRDRLILTFVPEGNSTSFV

NAIEVVSVPDDLITDSAQLLGVGQYLGLAAQPLQTFHRINVGGPKVTAENDTLARTWFAD

QSFFRNPTVAQAVTYQERLNYKDGSATQDDAPDSVYNTARRLVGQRNASSTPNMTWEFNV

DGRSSYLIRFHFCDIVSKAAFQLYFDVYVYNFSAAKDLDLSAREFGTLAAPFYMDIVLPS

SDPSGNLTVSIGPSSLPNATPDGILNGLEIMKMNFSSGSVYVVKPPSAAKQQLPIILGSV

LGGIGAAIIVVVLCVVFRRKKKMKKPQTPLTSRPSSSWTPLSLNALSFLSTGTRTTSRTT

YTSGTNSDTSYRIPFVVLQEATNHFDEQMVIGVGGFGKVYKAVLQDSTKVAVKRGNQKSH

QGIREFRTEIELLSGLRHRHLVSLIGYCDERNEMILVYEYMEKGTLKGHLYGGDQPPLSW

KKRLEICIGAARGLHYLHTGFAKSIIHRDVKSANILLDENLMAKVSDFGLSKTGPEFDQT

HVSTAVKGSFGYLDPEYYRRQKLTDKSDVYSFGVVLLEVICARPVIDPTLPRDMINLAEW

AIKWQKRGELDQIIDKRIAGTIRPESLRKYGETVEKCLAEYGVERPTMGDVLWNLEFVLQ

LQEAGPDMSNIDSMNQISELPSNAQRISSLEISTADESRTAMDYSQMSTSNAFSQLINTE

GR

>Os-CrRLK1L11

MASRRHVLIAALIMVGVLEFANADKYKPTESILVNCGSDKEGQDIDGRKWLSDKDSKWLI

DGEKSSIMANADFQDPSLPSPVPYMTARVFTKETMYNFSVGEERHWVRLHFYPASYHDLP

AENFFFSVSTSTGITLLKNFSVYITAKALSQAYIIREFTLPPSTTGSLSLIFTPTAMNNA

SYAFVNGIEIISMPNIFSQAAASVDIAGNEVSTTDSSLQTIYRLNVGGSYVAPTNDSGLS

RDWYDDTPYIYGAAVGVTYQANDTVQIKYPKNDPDAEYAAPASVYLTSRSMGPDPKVNKN

YKLTWVFEVDGNFTYIVRLHFCELLLSKPNQRVFDILINNKTAQSGADVIGWGGQFVPVY

KDYATIMPGGAGDKVLWVQLMPNVGSGSEFFDSLLNGLEIFKMSDSSGNLAGPNPDPSKL

LEEAESSAQGKFKSKPSNLKATVIGGAAGGAAAFGIVAAICIVVYQSKKRKVLNNSASHS

SGWLPVYGGNSHTSTSKSSGGRSAALINPNITAMCRHFSFGEIKSATKNFDESLVIGVGG

FGKVYRGVVDGDTKVAIKRSNPSSEQGVLEFQTEIEMLSKLRHKHLVSLIGCCEDEGEMI

LVYDYMAHGTLREHLYKGGKPALSWKQRLEITIGAARGLHYLHTGAKYTIIHRDVKTTNI

LVDEKWVAKVSDFGLSKTGPTAMNQTHVSTMVKGSFGYLDPEYFRRQQLTEKSDVYSFGV

VLFEVLCARPALNPSLPREQVSLADHAMSCQRKGTLHDIIDPLLNGKIAPDCLKKFAETA

EKCLADHGVDRPSMGDVLWNLEFALQMQETFENGGKTEGADSTSDSTTTSVADSMAANAA

ALSLISEDMDEEDIANSVVFSQLVRPTGR

>Os-CrRLK1L12

MIHPSLLTTIQWVALSTLILITIAADNYSSSSSPIFLNCGASTMQLDINNRSWEGDTRSK

FASAMNGIAASATYQDPSLPSLVPYMTSRIFISNYTYSFPISPGRIFVRLYFYPVAYGYY

ASEDAYFGVKTNNLILLDNFNASQTAQAANYAYILREFSLNVTLGSLDLTFFPSTQNGSY

AFVNGIEIVPTPDIFTTRTPTHNTEGNLDPSDIDSMTSFQTMYRLNVGGQAIIPQGDSRF

YRSWEDDSPYIYGAAFGVTFGKDSNVTITYPGTMPNYTAPADVYATARSMGPNWQINLNY

NLTWILSVDAGFYYLLRFHFCEIQYPITKMNQRSFFIYINNQTVQDQMDVIRWSGGIGMA

TYADYLIVTVGSGQMDLWVALHPDLSSRPQYYDAILNGLEVFKLWDIGKKNLAGLNPPLP

PQPKTDVNPKGVSGGGKLKAAVPAAICAVVVLITACFCVCIICRRKKVAKHSGKTDKKCL

TYQTELYKSPSNLCRNFTFHEMQIATSSFDETLLLGRGGFGDVYRGEIDNGTTVAIKRSN

PLSLQGVHEFQTEIETLSKVRHGHLVSLIGYCQEKNEMILVYEYMARGTLREHLYSTKRP

PLPWKERLKICIGAARGLYYLHTGPKETIIHRDVKTANILLDDKWVAKVSDFGLSKVNPD

IDATHVSTVVKGTFGYFDPEYFRLKQLTQRSDVFSFGVVLFEILCARPPVNTELPEEQVS

LREWALSCKKIGTLGEIIDPYLQGEIAPDCLKKFADCAEQCVADRSIDRPEMGDVLRNLE

VALKMQECAENNSKFSEETTSSKTTPDMMTIMDTDKQSTYSTMSITGQRTIFSDMMDPQA

R

>Os-CrRLK1L13

MRRRGRMIPPALLLAAAVAAALATAVSGQGRPVTESGAQTAPTPSTFTPKDNFLIDCGST

SPVTTGGKVYKTDAQSNSLLSAKDAIKVATTDADVPSPLYLTARIFRDEAVYSFPLTVPG

WHFVRLYLFPLKNSDFDLTTATFTVSTDTNVLLHSFTAENKPVMKEFLVNATENHLAVKF

YPLKGSAAFINAIEVVNAPDELITDMAMGIAPVGEMTGLAEAAYQVVYRINVGGPAIAPD

KDTLGRQWDVDAPYVQSKEAVKDVSVPVGNIKFPDGTSKLVAPAQVYASCAKMADAGVGS

PSFNMSWKMEVDPAFGYLVRLFFADIVSKSMNDLYFNVFVNGRKAISGLDLSTVTGELSA

AYYKDIVVNSSIATDKLSIQVGPMGEDTGRVDALLSGVEVLKMSNSVGSLDGEFGVDGKK

ADDGSGSRKAVAAVGFAMMFGAFAGLGAMAVKWYKRPQDWERRNSFSSWLLPIHTGQSFT

TSKGGSSKSGYTFSSTLGLGRFFSFAEIQAATKNFEESAIIGVGGFGNVYIGEIDDGTKV

AVKRGNPQSEQGINEFNTEIQMLSKLRHRHLVSLIGYCDENAEMILVYEYMHNGPFRDHI

YGKDLPALTWKQRLEICIGAARGLHYLHTGTAQGIIHRDVKTTNILLDDNFVAKVSDFGL

SKDGPGMNQLHVSTAVKGSFGYLDPEYFRCQQLTDKSDVYSFGVVLLETLCARPPIDPQL

PREQVSLAEWGMQWKRKGLIEKIMDPKLAGTVNQESLNKFAEAAEKCLAEFGSDRISMGD

VLWNLEYALQLQDANPPEGADKPADHDGAGAAPATSSGSGVSTVPDVSTTAAGEMFAQLA

DMKGR

>Os-CrRLK1L14

MPAARRSGGRLTEEVNMMVALSGRKRRLQAATMVALCFLSSICVSTAQFKPADNYLVDCGSSKSTTLGTRTFAADGAAPVKVDTSLEILAGTSANGVASFDNSALYQTARIFTSPSSYTFPIQKQGRHFVRLYFFAFAYQSYDLSTAKFTVSTQEMLLLSDFQQPDKTAPLFKEYSLNITQDKLIISFKPSNGIAFINAIEVVSVPDDLIGDSAPMVNPMQQYSGLSTQPLETVYRVNMGGPKVTADNDTLSRTWVTDKKYLVNPSVTREVNGGKVNYMKGGGSTPLIAPDIVYSTATELAASNTTNALFNMTWQFDVDSGFSYLIRFHFCDIVSKALNQLYFNAYVGSFYAQHDIDLSIQSMNQLATAIYLDVVLSSNDASNKLSISIGPSTLNNALPDGILNGLEVMKMSSGSGSAFTVGSSGSNKNLGVIIGSVLGAVGILIIVLVIVLLCRKKKTLEKQHSKTWMPFSINGLTSLSTGSRTSYGTTLTSGLNGSYGYRFAFSVLQEATNNFDENWVIGVGGFGKVYKGVLRDDTKVAVKRGNPKSQQGLNEFRTEIELLSRLRHRHLVSLIGYCDERNEMILVYEYMEKGTLKSHLYGSDNPSLNWKQRLEICIGAARGLHYLHTGSAKAIIHRDVKSANILLDENLLAKVADFGLSKTGPELDQTHVSTAVKGSFGYLDPEYFRRQQLTEKSDVYSFGVVLLEVLCARPVIDPTLPREMVNLAEWGMKWQKRGELHQIVDQRVSGSIRPDSLRKFGETVEKCLADYGVERPSMGDVLWNLEYVLQLQDADSSTVSDVNSMNRIVELPSQVQNIGALESISVTMAEAGASHEPDHDLSDVSMSRVFSQLIKAEGR

>Os-CrRLK1L15

MKMESFAWLLLILVLFSILEDVRGSKSKPILINCGSDSTTDVDGRRWIGDSSPKNFTLSL

PGTVATAPDSDGKETYGDLYKNARIFNASSSYKFIVAAAGSYFLRLHFSQLPTNFSTKES

LFDVSANGLKLVSKFNVPAEIYLRNSKINSTSRAIVKEYLLNVTSSNLEIEFSPDAESFA

FINAMEIVPVSGNSVFDSVNKVGGYGLKGPFSLGDSAVETMYRICVGCGKIESKEDPGLW

RKWDSDENFIFSMSAARAISNSSNISYVSSDDSTSAPLRLYETARVTTESSVMDKKFNVS

WSFNVDPDFDYLVRLHFCELEYDKAEQRKFKIYINNKTAAENYDVFAKAGGKNKAFHEDF

LDAASPQMDTLWVQLGSESSAGPAATDALLNGMEIFKVSRNGNLAHPTVRIGGFNSAMGK

PKRSPKWVLIGAAAGLVIFVSIVGVIFVCFYLRWKKKTSANKTKDNPPGWRPLVLHGATT

PAANSRSPTLRAAGTFGSNRMGRQFTVAEIREATMNFDDSLVIGVGGFGKVYKGEMEDGK

LVAIKRGHPESQQGVKEFETEIEILSRLRHRHLVSLIGYCDEQNEMILVYEHMANGTLRS

HLYGTDLPALTWKQRLEICIGAARGLHYLHTGLDRGIIHRDVKTTNILLDDNFVAKMADF

GISKDGPPLDHTHVSTAVKGSFGYLDPEYYRRQQLTQSSDVYSFGVVLFEVLCARPVINP

ALPRDQINLAEWALKWQKQKLLETIIDPRLEGNYTLESIRKFSEIAEKCLADEGRSRPSI

GEVLWHLESALQLHQGLLQSANTDDLSQPELKLSDASCNLGCIEEVEESCRAGSQDVNEE

YVDVKIEVP

>Os-CrRLK1L16

MAAIVLLLFLVVGLMPVSNGQTTPFSPRFSVYLACGAGGNVVVTSDSPQRTFVPDDGELS

GKSARFSNPDASPPSPLYAAARAGTSGFSYRLSYAADAAPDGNTTLVLRLHFFPFASQSG

DLLSARFSVSAMGRYVLLPPSFSPPRAGVVREFLLPSDGSGEFDVAFTPESGGLAFVNAI

ELFPAPQELLWKFPLTAVNTDVSPSHQALETLYRLNVGGPTVTPTGDTMWRTWLPDDSYL

SPATVSAVASIQGQIIFDRAQGYTQMVAPDAVYKSQRTTNSTTSNVTWTFAVDGNSSYVV

RLHFCAFEELSSVIGEGVDFNVYLMQAMGTRELKAKDYATLSSPTQAFYMDYVAVVPTAG

ENLTVSIGRAASSDSKKAILNGLEIMKLRAVDMTPASSSGKTSKVVVVAVTAAVLGAAVL

AGVALCVLLVRRRQRRATLPVPEEEEKESVGTPWSPFTPDGEGSFGSAVVTPRRMNMKLH

IPLAEIMVATGDFDDANILGVGGFGNVYRGVLRDGTRVAVKRAKRASRQGFPEFQTEILV

LSSIRHRHLVSLIGYCNERSEMILVYELMAHGTLRSHLYGSDAAAATPPPLSWKQRLEIC

IGAAKGLHYLHTGHSDNIIHRDVKSTNILLGDGFVAKVADFGLSRVGPSTGQTHVSTAVK

GSFGYLDPEYFKTRQLTDRSDVYSFGVVLFEVLCARPAIDQSLPPDEINLAEWAMQWSRR

GRFDKIVDPAVAGDASTNSLRKFAETAGRCLADYGEQRPSMGDVVWNLEYCLQLQESQPS

TETALDLDDSGAHLPRDIVVARRVAPLAPDASADAAGDDMSWSETASFTATGNVFSQIMS

RDGR

>Bd-CrRLK1L1(KQK13083)

MPTFAILAESMVEWKRVPMFLIIFILSITGVATTNAIASKTDRFVPQDNYLLSCGASAAV

PLDDGRTFRSDPDSVSFLSTPTDIKIAAKASLASASPLSPLYLTARVFSDISTYSFFISQ

PGRHWIRLYFSPIPESQYNLTTATFSVSTDNMVLLHDFSFIASPPTPILREYLVAVQGDN

LKIVFTPKKNSVAFVNAIEVVSVPPSLIPNTTTRMGPQDQFDISNNALQVIYRLNMGGAL

VTSFNDTLGRTWLPDAPFLKIEAAAEAAWVPPRTIKYPDDKTITPLIAPANIYSTAQKMA

SANITDARFNITWEMAADPGFRYLIRLHFSDIISKTLNSLYFNVYINGMMGVSNLDLSSL

TMGLAVAYYKDFIADSSSIINSTLVVQVGPSTTDSGNPNAILNGLEIMKISNEASSLDGL

FSPKTSSQVSKRTLTGIGLALVVTAALAVVICCRRSHRPEWQKTNSFHSWFLPLNSSHSS

FMSSCSRLSRNRFGSTRTKSGFSSIFASSAYGLGRYFTFAEIQKATKNFEEKGVIGVGGF

GKVYLGSIEDGTKLAIKRGNPSSDQGMNEFLTEIQMLSKLRHRHLVSLIGCCDENNEMIL

VYEYMSNGPLRDHLYGDTNIKPLSWKQRLEVSIGAAKGLHYLHTGAAQGIIHRDVKTTNI

LLDENFVAKVADFGLSKAAPSLEQTHVSTAVKGSFGYLDPEYFRRQQLTEKSDVYSFGVV

LFEVLCARPAINPALPRDQVNLAEWARSWHRKGELNKIIDPHIAGQIRPDSLEMFAEAAE

KCLADYGVDRPSMGDVLWKLEFALQLQEKGDVVEGSNDGIPMKSLEMSNVDNMEKSANVI

PSYVQGR

>Bd-CrRLK1L2(KQK06923)

MGFPILLVIILTLLSLILLVVVADNNSTASGHVALNCGASGQNNDDNGRTWDGDTSSKFA

PLVKGVTAPASYQHPSLPSTVPFMTARIFTSNYTYSFPVSAGRMFVRLYFYPIAYGNYAV

SDAFFSVTTRNLSLLNDFNASQTAQAINVAYLVREFSLNVSSGSLDLTFAPSKHWNGSYA

FVNGIEIVPTPDIFTTADTRFVNGGTPAPFQINTDRGFQTMHRLNVGGQAIPPKDDLSFY

RSWANDSPYIFGGSGVAFSRDNNLTIKYTSTVPNYTAPIGVYGTARSMGTNAQVNLNYNL

TWILPVDAGFFYLLRFHFCEIQYPITKVNQRSFFIYINNQTAQRQMDVIAWSGGIGRTAY

TDYVIITTGSGQVDMWVALYPDLSSKPEYYDAILNGLEVFKLQDYGKNNLAGLNPPLPQK

PDVNPNGPSREGNSRGTVLAAICGAIGGFAVLLICFGVCIACRRNKKISKDSDKSDDGCW

TPLADYSRSRSGNSGNTATTGSHASLPSNLCRHFSFAEVQAATNNFDQAFLLGKGGFGNV

YLGEIDSGTKLAIKRCNPMSEQGVHEFQTEIEMLSKLRHRHLVSLIGYCEDKNEMILVYD

YMAHGTLREHLYKTKNPPLSWKQRLEICIGAARGLHYLHTGVKQTIIHRDVKTTNILLDD

KWVAKVSDFGLSKTGPNVDNTHVSTVVKGSFGYLDPEYFRRQQLSEKSDVYSFGVVLFEV

LCARPALSPSLPKEQVNLADWALHCQKKGILGQIIDPLLQGKISPQCFVKFAETAEKCVA

DHSIDRPSMSDVLWNLEFVLQLQESAEDNSSLTGGMSSSDVSSPLVPTRLQSDEPSTETT

TTTTASTMSITQRSIASAESAGLTPSTIFSQLMNPDGR

>Bd-CrRLK1L3(KQK21820)

MGRSPRCVLLLLLAACAAALVPAAGAQGGGNSTAPAPAAAGSAPFVPRDDILLDCGATGN

GNDTDGREWGGDAGSKYAPANLGSAIAGAQDPSVPQVPYLTARVSAAPFTYSFPLGPGRK

FLRLHFYPANYSNRNAADAFFSVTVPAAKVTLLSNFSAYQTSTALNFAYLIREFSVNVTG

QTLDLTFTPEKGHPNAYAFINGIEVVSSPDLFDLSTPELVMGDGNNQPYTMEAGTALQTM

YRLNVGGQAISPSKDTGGYRSWDDDTPYIWGAGAGVSYQNDANVTITYPDNVPGYVAPTD

VYATARSMGPDKDVNLAYNLTWIMQVDAGFFYLVRLHFCEIQSPITKPNQRVFDIYINNQ

TAMAGADVILWASPNGIGSPVYKDYVVNTMGSGTMDFWVALHPDVTQKPQYFDAILNGME

VFKLQQSNGSLVGLNPVPSAEPLVDGGSGKKKSTVGPIVGGVVGGLAVLALGYCFIVICK

RRRRAGKDAGMSDGHSGWLPLSLYGNSHTSGSAKSHTTGSYASSLPSNLCRHFSFAEIKA

ATKNFDESLILGVGGFGKVYRGEVDGGTTKVAIKRGNPLSEQGIHEFQTEIEMLSKLRHR

HLVSLIGYCEEKNEMILVYDYMAHGTLREHLYKTQNAPLSWRQRLEICIGAARGLHYLHT

GAKHTIIHRDVKTTNILLDEKWVAKVSDFGLSKTGPSMDHTHVSTVVKGSFGYLDPEYFR

RQQLTEKSDVYSFGVVLFEVLCARPALNPTLAKEEVSLAEWALHCQKKGILDQIVDPYLK

GKIVPQCFKKFAETAEKCVADNGIERPSMGDVLWNLEFALQMQESAEESGSIGCGMSDEG

TPLVMVGKKDPNDPSIDSSTTTTTTTSISMGDQSVASIDSDGLTPSAVFSQIMNPKGR

>Bd-CrRLK1L4(KQK23907)

MAVVLLALLLLAATLLLAPSPAALAAFSPSFECFLACGAADNLSFPSDSPARVFVPDAAF

LSPATTPALASSQASSSALYAAARGSSSEFSYSIPCSSPATFLVLRLHFFPLPATPSSSA

RFAVSVRHGAYALTLLQSFSPPPAGVVKEFFLPDAGSNGELRVTFAPASGSSAFVNALEL

FPAPPELLWNENSPYPYTPVGTAADNNATATWPQQALETLHRLNVGGPTVNSTLDTLWRT

WLPDDAFLYGGIAQTTVGSSITPVFDPDNGYTREVAPDVVYKTQRFANVTDYMLATNPGI

SFNVTWTFPAVPGSGGYLVRLHFCDYDMVSSVVGVGIVFDVYVAQAVAARDLKLAELGKR

VPSQAFYFDYAAMAPSAGNLTVSIGKARSTGGMILNGLEIMKLLPLSAVSSHEGMPKRTI

VIAALASVLGAAVLACSVLCLVVLMRRRKRRMRPAPEKASTTMPPWSPFRGGSSWVVDQS

TDHSGEGTGMQRVISTKLHISLSEIRAATEGFHERNLIGVGGFGNVYKGALHDGTPVAVK

RAMRASKQGLPEFQTEIVVLSGIRHRHLVSLIGYCDDQAEMILVYEYMEHGTLRSHLYGF

DDDDDNSEPLSWKQRLEICIGAARGLHYLHTGYSENIIHRDIKSTNILLGSEDGVLVAKV

ADFGLSRIGPSFGETHVSTAVKGSFGYLDPEYFKTQQLTDRSDVYSFGVVLFEMLCARPV

IDQSLDRDQINIAEWAVRMHGQGQLGKIVDPRMAMAAGGVDENSLRKFAETAEKCLADYG

VDRPSMGDVLWNLEYCLQLQETHVSRDAFEDSGAVTATRLPAGVVVPRWVPASTMDDVDD

TGMSIGMSVVADSKVFSQLSAGGEGR

>Bd-CrRLK1L5(KQK22263)

MPPHLDLLLLWLLVVSAPVLASAATPSTAFVPADNYLVICGTSGSATDTAGRTFVGDGRL

PASALAAPQSVEANASLSSSNGDEQALYQSARIFTAPASYTFAIKKPGRHFVRLHFFPFR

YQSYDLAAAAAFKVFVQGAVFVDGSYTPKNGTVVVKEFSVNVTGGSLVIAFTPTGKLAFV

NAIEVVSLPDDLIADTAAMAGSARGLYTGLSARALETVHRINMGAPKITPANDTLWRTWL

PDQSFQLDSSLALAEHKEVLPSAIKYTPVATPWTAPVGVYATATKQSTSGGTSTINVQFN

VTWRFGAVAAGSDYLLRFHFCDIVSKAATGLAFNVYVGAWLVLDNYEYSRDTINTLAVPV

YKDFVLGAKDVKGGNITVSIGSSTVGVSNVSPDGFLNGLEIMRVLGSAGAGAEPSKRSSK

VKTWIIAGSAVGGAAVAMALAFIAFRMLCRKRGKPEKKASNSTLSPFSASALGSRSRSSG

KKSNGNTIVLGQNGLGAGYRIPLAVLQEATSGFGEAMVIGEGGFGKVYKGTLPDETPVAV

KRGSRKTLQAMQEFRTEIEMLSRMRHRHLVSLIGYCDARDEMILVYEYMAMGTLRSHLYG

ADDLPPLTWEQRLEACIGAARGLHYLHTSSATAVIHRDVKSSNILLDETLMAKVADFGLS

KAGPELDKTHVSTKVKGSFGYLDPEYFRRQMLTEKSDVYSFGVVLLEVLCARAVIDPTLP

REMVNLAEWAMQWLKKGEVDRIVDQRIAGTIRPQSLKKLADTAEKCLAEYGVERPTMGDV

LWCLEFALQLQVASPDDSVIDGMPLAPVATPQVQRIQSIASVATDTAMTANLGDLDGMSM

SGVFSKMVKSEEVR

>Bd-CrRLK1L6(PNT61865)

MFQMRLLVLAAVSIVFANLQFLKAHGRELLLSCGSNATVDADGRRWVGDMAPDLNFTLSS

PGIAALLAGGSNASEIFGPVYRSARLFTTTSWYDFSVLPGNYCIRLHFFPSTFGNFSANS

SVFDVVANDFKLVSKFNVSEEILWRSSVSNLAVTAIVKEYFLAVSTQRLQIEFDPSRGSF

AFVNAIEVMLTPDNSFNDTVHKVGGGDGYLPPGLSSRGVETMYRLNIGGPALASSSDQYL

HRPWYTDEAFMFSANAALTVSNTSAIRYLSSNDSSIAPIGVYETARIMSNNMVVDKRFNV

TWRFFVHPNFDYLVRLHFCELVYDKPSQRIFKIYINNKTAAENYDVYVRAGGINKAYHED

YFDSLPQQVDSLWIQLGPDSMTSASGTDALLNGLEIFKLSRNGELDYVLGHIDMGNQRGP

SKGKRKINIWEEVGIGSASFVMLASVALFSWCYVRRKRKAAEKEAPPGWHPLVLHEAMKS

TTDARASSKSPLARNSSSIGHRMGRRFSISDIRSATKNFDETLVIGSGGFGKVYKGEVDE

GTTVAIKRANPLCGQGLKEFETEIEMLSKLRHRHLVAMIGYCEEQKEMILIYEYMAKGTL

RSHLYGSDLPPLTWKQRLDACIGAARGLHYLHTGADRGIIHRDVKTTNILLDKNFVAKIA

DFGLSKTGPTLDQTHVSTAIRGSFGYLDPEYFRRQQLTQKSDVYSFGVVLFEVACARPVI

DPTLPKDQINLAEWAMRWQRQRSLEAIMDPRLDGDYSPESLKKFGDIAEKCLADDGRTRP

SMGEVLWHLEYVLQLHEAYKRNLDCESFGSSELGFADMSFSMPHIREGEEERQSKQSGIR

EDSDT

>Bd-CrRLK1L7(KQK07346)

MGTATNERKLAVVFLWSISVLSGTCNAEFTPADNYLINCGSTVDASVGRRVFEADNSKST

ILTSHQSVAANTFLDSVPASDYAVLYQTARIFGVPSSYSFKMKSRGRHFVRLHFFSFKYQ

SYDLAVAKFKVSTQDAVLLDNFTPPSNSSPVVREYSLNITRGMLILTFVPLGNSTSFINA

IEVISVPDDLILDLGQSVNPIRQYAGLAAQPFQTFYRINVGGRKVTADNDTLWRSWDTDQ

SFFLNSTTTQTVTYEGKLNYQRGAATEEDAPDSVYNTARRLVAQNNTASASNMTWQFNVD

RRASYLIRFHLCDIVSKAMAALYFDVYVDRWSAAEDLDLSEKGFGTLAVPYYTDVILESS

DPSGKLSVSIGPSSLNNVAQDGILNGLEIMKMNISTGTVEIVQPPPSQKRELPIILGSIL

AVCAATAVAILCFVLRRKKNKKPQTASTSRTSSAWTPLTLNGISFLSTGTRTTSRTTLTS

GTNGDATYQIPFVVLQEATNHFDEQMIIGVGGFGKVYKAVLQDGTKVAVKRGNHKSHQGI

KEFRTEIELLSGLRHRHLVSLIGYCNEHNEMILVYEYMEKGTLKGHLYGSDIPALSWKKR

VEICIGAARGLHYLHTGFAKSIIHRDVKSANILLDENLMAKVSDFGLSKTGPELDQTHVS

TAVKGSFGYLDPEYYRRQKLTDKSDVYSFGVVLLEVICARPVIDPSLPREMINLAEWASK

WQKRGELDQIVDQRIAGTIRPESLRKYGETVEKCLAEYGVDRPTMGDVLWNLEFVLQLQE

SGPDITNIDSMNQISELPSEARRVGSLEISTADESHTNINYSQMSTNDAFSQLMNTEGR

>Bd-CrRLK1L8(KQK06928)

MRARQHLFLAELLLVISWVLLFCDAEKYEPTETILVDCGSEKDGQDAQGRKWVMDKDSKW

LGDGGKSSMMAAADAQDPSLPSPVPYMSARVFTKEAVYTFPVADADRHWVRLHFYPAAYH

DLPAEQFFFSVSTASGITLLRNFSVYITVKALSQAYIVREFTLPPSTTGSISLKFTPTAM

NNASYAFVNGIEIISMPNIFAEPATLVGLDSQTVDLAAGSLQTMYRLNVGGAYVASTNDS

GLSREWFDDTPYIYGAATGVTFEPNDTFPIKYPSPEGEFAAPADVYITSRSMGPDGRVNK

NNNLTWVFEVDANFTYVLRLHFCGLRVDKVNQVVFDIYINNKTAQDNADIIGWSSAKDVP

VFKDYAVFMPDMPGDKILWLALHPDVDSKPQFFDAILNGLEIFKMSDGSGNLAGPNPDPS

KMLMESEVEQGKFRAKPSNLQATLIGGAAGGAAALGIVAAICLVVYQTKKNRALSSSPSH

SSGWLPVYGGNSHTNASSGSRSAALNPNITAMCRHFSFPEIKSATKNFDEGLVIGVGGFG

KVYKGVVDGDTKVAIKRSNPSSEQGVMEFQTEIEMLSKLRHKHLVSLIGCCEDDGEMILV

YDYMAHGTLREHLYKSGKPPLLWKQRLEIVIGAARGLHYLHTGAKYTIIHRDVKTTNILV

DEKWVAKVSDFGLSKTGPTAQNQSHVSTMVKGSFGYLDPEYFRRQQLTEKSDVYSFGVVL

FEVLCARPALNPSLPREQVSLADHALSCQRKGTLQDIVDPLLKGKIAPDCMKKFAETAEK

CLADHGVDRPSMGDVLWNLEFALQMQETFENGGKPEGGDSVGSSSSGSTPPSMADSMAAN

AAALSLICEDMDEEDIANSVVLSQLVRPTGR

>Bd-CrRLK1L9(KQK19638)

MRRGAVPLALALATLAMLAVSVSGQGRPVIDTSGGMTEALPSKFTPKDAFFIDCGGTNPV

TVEGKAFKTDAQANQLLAAQDAIRASVDKADSVSSPVYLTARIFKEEAVYNFPLAVPGWH

FIRLYFFPLKNPDSDLAAATFSVTTDTNVLLHSFTADPKPTMKEYLINATENHLEIKFTP

LKGSAAFINGIEVVNGPDELITDTALAVLPFAEMSGLSEAAYQVIYRLNVGGPGISPGND

TLGRQWDNDEKYVQSKEMVKDVSVPTNTIKYPDTFPVSKLVAPMLVFASAAKMADMDSTV

SNANFNVTWKLDVDPSFDYFVRLFFADIISKSANDLYFNVYIDGRKAISGLDLSGITGDL

AVPYYKDFVVNSSITADGHLSIQIGPLGQDTGRIDALLNGAEVFKMSNSVGSLDGEFGVD

GRKADDGSGGRKVVAVVGFAMMFGAFAGLGAMVVKWHKRPQDWQRRNSFSSWLLPIHTGQ

SFSNGKGSKSGYTFSSTGGLGRFFSFAEMQEATKNFDESAIIGVGGFGNVYVGEIDDGTK

VAIKRGNPQSEQGINEFNTEIQMLSKLRHRHLVSLIGYCDENAEMILVYEYMHYGPFRDH

IYGGDGNLPALSWKQRLEICIGAARGLHYLHTGTAQGIIHRDVKTTNILLDENFVAKVAD

FGLSKDGPGMDQLHVSTAVKGSFGYLDPEYFRCQQLTDKSDVYSFGVVLLETLCARAPID

PQLPREQVSLAEWGLQWKRKGLIEKIMDPKLAGKVNEESLNKFAETAEKCLAEFGSDRIS

MGDVLWNLEYALQMQEQNPPEGAAAGEGGDGDSLDAGISSSSIATTSSSSSGINTVPDAS

TTSAGELFAQLADMKGR

>Bd-CrRLK1L10(KQK21184)

MPGRRRKLQVAALLLLFCVCSSAGIICRAQLVDQFKPADSYLVNCGSAKGTTVSERNFAA

DGAAPLTVSTPQEILAGTSANGVSSFDNSALYQTARIFTGPSSYTFPINKQGRHFVRLYF

FPFIYQSYDLSTAKFTVSTQDVLLLSDFQQPDKTAPLFKEYSLNITRDQLIISFKPSNGI

AFVNAIEVVSVPDDLIADVAQMVNPVQQYSGLSTQSLETVYRVNMGGPKVTPNNDTLSRT

WLNDQKFIMNPSVTKKVVYGKTIKYKNGGASSLTAPDVVYSTATELAASNTSNALFNMTW

QFDVDAGFSYLIRFHFCDIVSKALNQLYFNAYVGGFFAQHDLDLSEQSMNQLATAIYVDV

VLSSNDASSKLSISIGPSTLNNAFPDGILNGLEVMKMGSGSGSAFTVGSSGSKKILAVII

GSVIGVIGLLVIVLLLVLLCRRKKTDDKQHSKTWMPFSINGLTSLSTGSRTSYGTTLTSG

LNGSLGYRFAFNVLQEATNNFDENWVIGVGGFGKVYKGVLRDDTKVAVKRGNPKSQQGLN

EFRTEIELLSRLRHRHLVSLIGYCDERNEMILVYEYMENGTVKSHLYGSDNPSLNWKQRL

EICIGAARGLHYLHTGSAKAIIHRDVKSANILLDENFLAKVADFGLSKTGPELDQTHVST

AVKGSFGYLDPEYFRRQQLTEKSDVYSFGVVMLEVLCARPVIDPTLPREMVNLAEWGMKW

QKRGELHQIVDQRLSSTIRPDSLRKFGETVEKCLADYGVERPSMGDVLWNLEYVLQLQDA

DSSTVSDVNSMNRIVDLSSQVQHVGALESISMTMAEAGASHEPDHDLSDVSMSRVFSQLI

KAEGR

>Bd-CrRLK1L11(KQK21036)

MKSSTNLLLILVVFLAAENARAQPGPILINCGSDSSSAVVDGRRWIGDSSPSKNFTLSFP

GTIALAAAAPGVDGEEEPYGDLYKTARVFNSSSSYNLGVAAGSYFLRLHFSQLFANLSAE

EPIFDVAANGLKLLSRFSVTGEISWRDSQINSTSKVIVKEYLLNFTSGKLGIEFRPDEGS

FAFVNAMEVVPVSGSSIFDSVNKVGGYGLKGPFSLADGGIETMYRLCVGCSDIARKEDPG

LWRKWDSDEHFIFSLNAAHTITNSSNISYASADDSTLAPLRLYETAKVTTESSVVEKKFN

VSWSFIIDPGFDYLVRLHFCELEYDKAEQRKFKIYINNKTAAESYDVFARAGGKNKAFHE

DFLDVASPQMDTLWVQLGSESSAGAAATDALLNGMEIFKVSREGNLAHPTVRIGGISGGT

RKPKRSPKWVLIGAATGLIVFIAIVGAVYICFCLQRKKRSSANKTKNPPGCQPLALHGSA

NTRSPSLRTAGTLGSSQLGRRFTIAEIRTATQNFDESLVIGVGGFGKVYKGKMESGTLVA

IKRGHTESQQGQGVKEFETEIEMLSRLRHRHLVPLIGYCDERNEMILVYEHMANGTLRSH

LYGSDLPALTWNQRLEICIGAARGLHYLHTGLDRGIIHRDVKTTNILLNGNLVAKMADFG

ISKDGPALDHTHVSTAVKGSFGYLDPEYYRRQQLTPSSDVYSFGVVLLEVLCARPVINPT

LPRDQINLAEWALNCQRQQLLETIIDPRLDGNYTLESMKTFSKIAEKCLADEGVNRPSMG

EVLWHLESALQLHQGHLHADCDEVLSGPELTPSDASITVTHIREAEESTRAARDANDEVV

DVKIEVP
