## Supplementary Tables for "Genome wide characterization and expression analysis of CrRLK1L gene family in wheat unravels their roles in development and stress-specific responses": Supplementary Table S4. Intron-exon determination of CrRLK1L genes in T. aestivum.docx

Supplementary Table S4. Intron/exon lengths CrRLK1L genes in *T. aestivum*

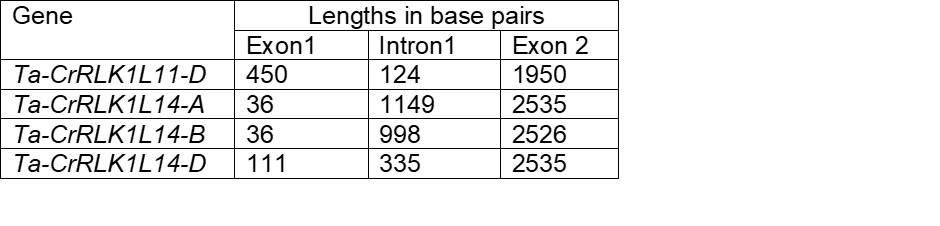

Note: Intron/Exon lengths for *CrRLK1Ls* were determined by comparison of NR database retrieved cDNA sequences for *CrRLK1L* genes and genomic sequences from Ensembl Plants database.
