## Supplementary Tables for "Genome wide characterization and expression analysis of CrRLK1L gene family in wheat unravels their roles in development and stress-specific responses": Supplementary Table S5. Conserved motif prediction in CrRLK1L by MEME suite.docx

Supplementary Table S5. Details of the conserved motifs predicted by MEME suite

| Sr No. | Motif sequence | Pfam/NCBI-CDD ID | Description of the motif |
| --- | --- | --- | --- |
| 1 | QGLHEFQTEIEMLSKLRHRHLVSLIGYCDEQNEMILVYEYMAHGTLRSHL | PF07714 | Protein tyrosine and serine/threonine kinase |
| 2 | ZTHVSTAVKGSFGYLDPEYFRRQQLTEKSDVYSFGVVLFEVLCARPVIBP | PF07714 | Protein tyrosine and serine/threonine kinase |
| 3 | WKQRLEICIGAARGLHYLHTGAAKGIIHRDVKTTNILLDENLVAKVADFG | PS00108 | Serine/Threonine protein kinases active-site signature |
| 4 | IVDPRJAGKIRPESLRKFAETAEKCLADYGVDRPSMGDVLWNLEFALQLQ | 271060 | Catalytic domain of the Serine/Threonine kinase |
| 5 | RHFSFAEJQAATKNFDESLVIGVGGFGKVYKGEJDDGTKVA | cd14061 | Catalytic domain of the Serine/Threonine Kinases |
| 6 | NLNFNLTWQFDVDAGFSYLVRLHFCDIVSKA | pfam12819 | Malectin like carbohydrate-binding protein of the ER |
| 7 | LPREQVNLAEWALRWQRKGEL | ND | ND |
| 8 | PYPGLSAAALZTMYRLNVGGPAISPSNDT | ND | ND |
| 9 | GSYAFVNAIEVVSAPDELIAD | ND | ND |
| 10 | PGRHFLRLHFFPFAYSSYDLA | ND | ND |

Note: Motifs predicted by MEME suite were searched for their details at Genome Net database. Pfam or NCBI Conserved Domain Database (CDD) ID for the IDs are given. ND denotes the motifs that did not find hits in any of the protein databases.
